# Type-II kinase inhibitors that target Parkinson’s Disease-associated LRRK2

**DOI:** 10.1101/2024.09.17.613365

**Authors:** Nicolai D. Raig, Katherine J. Surridge, Marta Sanz-Murillo, Verena Dederer, Andreas Krämer, Martin P. Schwalm, Lewis Elson, Deep Chatterjee, Sebastian Mathea, Thomas Hanke, Andres E. Leschziner, Samara L. Reck-Peterson, Stefan Knapp

## Abstract

Aberrant increases in kinase activity of leucine-rich repeat kinase 2 (LRRK2) are associated with Parkinson’s disease (PD). Numerous LRRK2-selective type-I kinase inhibitors have been developed and some have entered clinical trials. In this study, we present the first LRRK2-selective type-II kinase inhibitors. Targeting the inactive conformation of LRRK2 is functionally distinct from targeting the active-like conformation using type-I inhibitors. We designed these inhibitors using a combinatorial chemistry approach fusing selective LRRK2 type-I and promiscuous type-II inhibitors by iterative cycles of synthesis supported by structural biology and activity testing. Our current lead structures are selective and potent LRRK2 inhibitors. Through cellular assays, cryo-electron microscopy structural analysis, and in vitro motility assays, we show that our inhibitors stabilize the open, inactive kinase conformation. These new conformation-specific compounds will be invaluable as tools to study LRRK2’s function and regulation, and expand the potential therapeutic options for PD.

## Introduction

Parkinson’s disease (PD) is a progressive neurodegenerative disorder affecting over 10 million people globally (**1**). While most PD cases are idiopathic with no known cause, approximately 10 % are familial, linked to specific gene mutations. Mutations in leucine-rich repeat kinase 2 (LRRK2) are a common cause of familial PD (*2*–*5*). Significantly, LRRK2 kinase hyperactivity has also been described in idiopathic PD cases (**6**), making LRRK2 a key target for PD research and therapeutic intervention.

LRRK2 is a large, multidomain protein. At its N-terminus, LRRK2 contains armadillo (ARM), ankyrin (ANK) and leucine-rich repeat (LRR) domains. Its catalytic, C-terminal portion is made up of Ras of complex (ROC), C-terminal of ROC (COR), kinase and WD40 domains (**Figure 1 A**). Most pathogenic mutations in LRRK2 are located within the catalytic portion, termed LRRK2^RCKW^ for <u>R</u>OC, <u>C</u>OR, <u>K</u>inase, <u>W</u>D40. Of these, the G2019S mutation is the most common variant associated with PD. This mutation is located in the kinase active site and leads to an aberrant increase in kinase function (*7*–*9*). The vast majority of PD-linked mutations, located both within the kinase and at distal domains, share this common feature of increasing LRRK2 kinase activity (**10**).

**Fig. 1.**
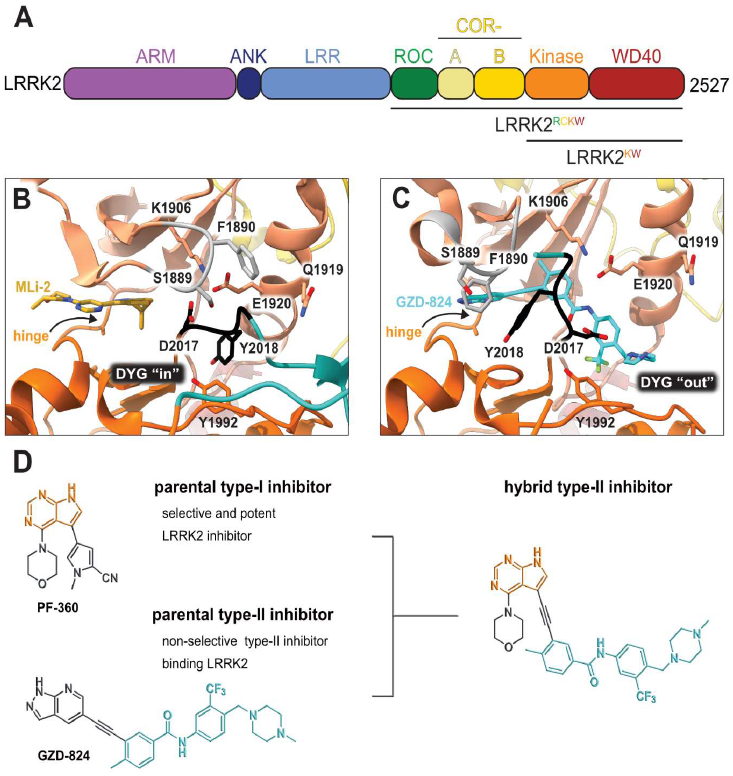
LRRK2 type-II inhibitor design strategy. **(A)** Schematic domain structure of LRRK2. The three constructs used in this study are indicated: full-length LRRK2, LRRK2^RCKW^ and LRRK2^KW^; **(B)** and **(C)** Close up of the inhibitor binding pocket from cryo-EM maps and models of LRRK2^RCKW^ bound to the type-I inhibitor MLi-2 (PDB: 8TXZ) (B) and type-II inhibitor GZD-824 (PDB: 8TZE) (C). Key residues and features are labelled. Both structures are shown in the same view, aligned through the C-lobe of the kinase. Dark orange: C-lobe; light orange: N-lobe; black: DYG motif; grey: G-loop; green: activation loop. **(D)** Scheme depicting our hybrid design strategy to develop potent and selective type-II inhibitors for LRRK2.

Kinases are highly ‘druggable’ proteins, with over 80 kinase inhibitors currently clinically approved worldwide (**11**). Given LRRK2’s status as a key, actionable therapeutic target, considerable efforts have been made in recent years to develop high affinity, highly specific LRRK2 kinase inhibitors. The first generation of selective LRRK2 inhibitors such as LRRK2-IN-1, CZC-25146 and GNE-7915, was published in 2011 (*12*–*14*). In 2015, the next generation of LRRK2 inhibitors, which included PF-360 and MLi-2, were disclosed (*15, 16*). These next generation inhibitors have been transformative for the field, possessing high affinity for LRRK2 and exceptional selectivity profiles. MLi-2 is regarded as the gold standard chemical tool compound for LRRK2 inhibition and has contributed significantly to building our understanding of LRRK2’s mechanism and function at the molecular, cellular, and organismal scales. Despite the excellent pharmacological properties of these published inhibitory compounds and two clinical trials (clinicaltrials.gov) for LRRK2 kinase inhibitors currently in progress, no LRRK2-targeting kinase inhibitor has been approved for clinical use.

All LRRK2 kinase inhibitors published to date are type-I inhibitors, which bind to the kinase active site and stabilize an active-like/ closed conformation (**Figure 1 B**). In contrast, while broad-spectrum type-II inhibitors including GZD-824 and Rebastinib efficiently target LRRK2 (*17*–*19*), there are currently no published highly selective type-II LRRK2 inhibitors that could be used to study the effects of targeting the LRRK2 inactive conformation. Thus, generating LRRK2-specific type-II kinase inhibitors is a critical need for probing the cellular function of LRRK2 and for the further development of LRRK2 therapeutics. Type-II inhibitors stabilize the inactive/ open kinase conformation (**Figure 1 C**), reaching into the far back pocket of the kinase active site and disrupting a critical salt-bridge by releasing the regulatory αC helix (*20, 21*). These disruptions prevent the Y from the DYG motif in the LRRK2 kinase active site (or F in the DFG motif commonly found in other kinases) from docking into the back pocket and placing the D in its catalytically-competent position (the “DYG in” conformation), instead placing the DYG into the “DYG out” conformation (**Figure 1 C**)(*20, 21*). Recent cryo-EM structures from our group and others of LRRK2 bound to selective type-I and non-selective type-II inhibitors have revealed the key structural differences in engaging the two main classes of kinase inhibitor, paving the way for the rational design of new inhibitory compounds (*18, 22*).

Recent studies have highlighted the significance of these contrasting conformational changes within the LRRK2 kinase domain in terms of LRRK2 activation and subcellular localization (*19, 23*–*25*). While the vast majority of LRRK2 is predicted to be cytosolic, LRRK2 activation requires it to be recruited to membranes where its substrate, Rab GTPases, are located (*26*–*28*). In addition, under certain conditions LRRK2 forms filamentous associations with microtubules (*29, 30*) that block the movement of dynein and kinesin motor proteins *in vitro* (**19**). Type-I inhibitors enhance LRRK2’s ability to block motor motility, whereas type-II inhibitors can restore motor protein movement in the presence of LRRK2 (**19**). Type-I inhibitors also induce dephosphorylation of the LRRK2 biomarker phosphorylation site S935, a response not elicited by type-II inhibitors (**31**). Potentially pathogenic histopathological changes to the lungs of mice and non-human primates have also been described upon LRRK2 depletion or pharmacological inhibition with type-I inhibitors (*32, 33*). Whether these effects are related to the specific mode of action of type-I kinase inhibitors is unknown, but taken together these data reinforce the importance of developing LRRK2-specific type-II kinase inhibitors both as potential therapeutics against PD and as chemical probes to explore the functional significance of LRRK2’s different conformational states in cells and *in vivo*.

Here, we set out to design type-II inhibitory compounds with high affinity for LRRK2 and significantly improved selectivity profiles compared to the currently available broad-spectrum type-II kinase inhibitors that can act on LRRK2. We used a combinatorial chemistry approach for the compound design, integrating the hinge-binding properties of well-characterized and selective type-I LRRK2 inhibitors with the binding mode of promiscuous type-II inhibitors known to associate with LRRK2 (**Figure 1 D**). Here we present the development and characterization of the first LRRK2 selective type-II inhibitors, named RN277 and RN341, as cellular tools targeting the LRRK2 inactive state.

## Results

### Compound design, synthesis and evaluation

#### MLi-2 hybridization series

The type-I kinase inhibitor MLi-2 is the gold standard chemical tool compound for LRRK2 due to its outstanding kinome selectivity and high affinity (picomolar range) (**16**). Therefore, to develop a LRRK2-specific type-II inhibitor we chose to first use a hybrid design strategy that incorporates the specificity and excellent pharmacokinetic properties of MLi-2 with type-II allosteric DYG-out binding moieties of established, broad-spectrum type-II inhibitors. Specifically, our design was based on combining the hinge-binding indazole of MLi-2 with the DYG-out pocket binding moieties from GZD-824 and Rebastinib, two broad-spectrum type-II inhibitors known to bind LRRK2. Additionally, we screened a library of published type-II inhibitors to identify those that bind to LRRK2^KW^ (a <u>K</u>inase and <u>W</u>D40 domain construct of LRRK2) with high affinity. Using differential scanning fluorometry (DSF) we identified Foretinib as an additional broad-spectrum type-II LRRK2 ligand (Supporting Information **Table S1**), suitable for use as a parental type-II inhibitor in our hybrid approach. This initial synthesis strategy using MLi-2 as the parental type-I compound led to the generation of eight candidate LRRK2-directed hybrid type-II inhibitors.

Next, we evaluated the potency and selectivity of these new compounds. We performed on-target screening by measuring the thermostability increase of LRRK2^KW^ in the presence of the new compounds using DSF (Supporting Information **Table S2**). We also determined the *in vitro* IC_50_ (median inhibitory concentration) values for each of the new compounds by purifying LRRK2^RCKW^ and measuring its kinase activity in the presence of our new inhibitors using the *PhosphoSense*^*®*^ activity assay from AssayQuant (Supporting Information **Table S2**). Finally, we estimated the kinome selectivity of each compound by measuring the thermostability increase of approximately 100 representative kinases in the presence of the compounds via DSF (Supporting Information **Table S2**). Only one compound (**52**, Supporting Information **Table S2**) of our first series of MLi-2 inspired hybrid compounds showed high thermal stabilization of LRRK2^KW^ and inhibited LRRK2 kinase activity. Compound **52** increased the melting temperature of LRRK2^KW^ (Δ*T*_M_) by 10 K, suggesting strong binding, and inhibited LRRK2 kinase activity with a sub-micromolar IC_50_ value. However, the selectivity of compound **52** was unfavorable, demonstrated by significant Δ*T*_M_ values for 27 % of the 100 kinases we screened in our initial DSF selectivity panel (Supporting Information **Table S2**).

#### PF-360 hybridization series

Since most of the MLi-2-like hybrid type-II inhibitors demonstrated inadequate LRRK2 binding, we exchanged the hinge binding moiety to the 7*H*-pyrrolo[2,3-*d*]pyrimidine, inspired by PF-360 (**Figure 1 D**), another type-I LRRK2 binding compound with a favorable selectivity profile (**15**). This moiety was fused with the DYG-binding moieties of type-II inhibitors GZD-824 and Foretinib to give rise to two potential LRRK2 inhibitors, **1** and **2**.

We evaluated the potency and selectivity of these two compounds using the assays previously described (**Table 1**). In terms of on-target binding, we found that compound **2** bound more strongly (higher ΔT_m_) to LRRK2^KW^ than compound **1** (**Table 1**). **2** also demonstrated strong inhibition of LRRK2 kinase activity with an IC_50_ value of 185 nM (**Table 1**). However, as observed for the MLi-2 inspired compound **52**, the selectivity values of both **1** and **2** were still unfavorable (**Table 1**). Given the potent on-target binding and inhibitory activity of compound **2**, we next synthesized a larger set of PF-360-Foretinib inspired inhibitors.

**Table 1.**
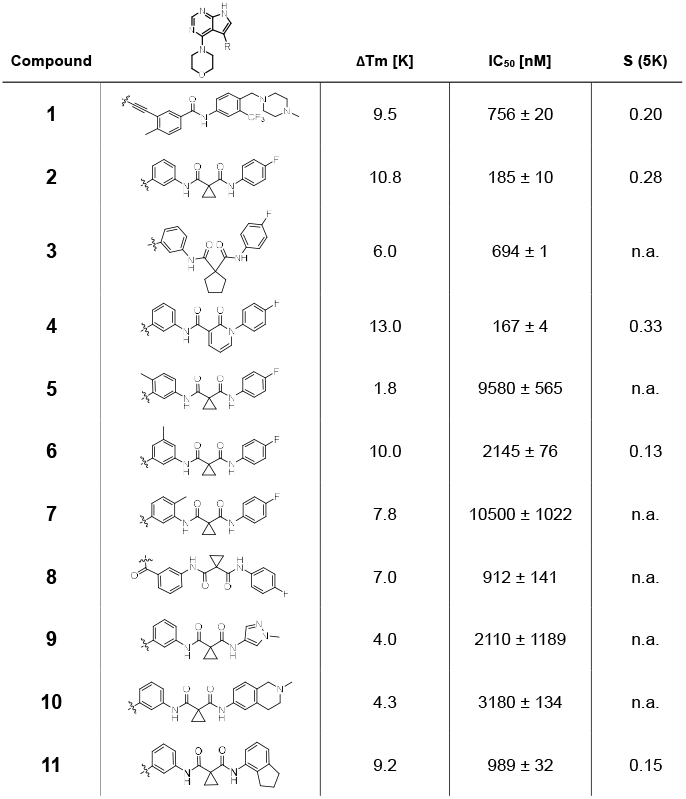
First PF-360 hybridization series. Compound structures, thermal shift data and *in vitro* IC_50_ values of PF-360 inspired type-II hybrid compounds. The selectivity was calculated as the number of kinases showing a thermal shift greater than 5 K divided by the total number of kinases screened.

| Compound | | $\Delta T_m$ [K] | IC<sub>50</sub> [nM] | S (5K) |
| --- | --- | --- | --- | --- |
| **1** |  | 9.5 | 756 ± 20 | 0.20 |
| **2** |  | 10.8 | 185 ± 10 | 0.28 |
| **3** |  | 6.0 | 694 ± 1 | n.a. |
| **4** |  | 13.0 | 167 ± 4 | 0.33 |
| **5** |  | 1.8 | 9580 ± 565 | n.a. |
| **6** |  | 10.0 | 2145 ± 76 | 0.13 |
| **7** |  | 7.8 | 10500 ± 1022 | n.a. |
| **8** |  | 7.0 | 912 ± 141 | n.a. |
| **9** |  | 4.0 | 2110 ± 1189 | n.a. |
| **10** |  | 4.3 | 3180 ± 134 | n.a. |
| **11** |  | 9.2 | 989 ± 32 | 0.15 |

In this set of **2**-derived analogues, major changes to the inhibitor geometry were introduced. These changes led to weaker inhibition or less selective compounds (**Table 1**) and we therefore focused the structure activity relationship (SAR) analysis on compound **2**. We hypothesize that the terminal phenyl ring of **2** would be positioned in the far back pocket of the kinase active site, similar to the crystal structures of the parental inhibitor Foretinib with other kinases (PDB: 6I2Y, 5IA4). This region of the binding pocket is less conserved in kinases compared to the ATP binding site, which may allow for improved selectivity by introducing diverse moieties at this position (*20, 21*). We systematically alternated the substitution pattern of the terminal phenyl ring by synthesizing 16 new hybrid type-II inhibitors.

By evaluating the potency and selectivity of these compounds (**Table 2**) we found that the fluorine in *para* position (R^3^) of the phenyl ring is beneficial for LRRK2 binding (**12**-**16**). The addition of substituents in *meta* position (R^2^/R^4^) (**17**-**21**) was well accepted, with IC_50_ values in the low nanomolar range (**Table 2**). With a bromine in *meta* position (**19**) the stabilization of LRRK2^KW^ significantly increased. The addition of a fluorine substituent in *ortho* position (R^1^) (**23**) was accepted and the selectivity increased dramatically to S (5K) = 16 %. We therefore combined more potent *meta* substitutions (R^2^/R^4^) with more selective *ortho* (R^1^) substituted derivatives to produce two regio isomers, **25** and **26** (named RN222). This new compound, RN222 (**26**), showed both increased inhibition of LRRK2^RCKW^ kinase activity and improved selectivity (IC_50_ = 134 nM and S (5K) = 14 % **Table 2**).

**Table 2.** Second PF-360 hybridization series. Compound structures, thermal shift data and *in vitro* IC_50_ values of compound 2-derived hybrid type-II inhibitors. The selectivity was calculated as the number of kinases showing a thermal shift greater than 5 K divided by the total number of kinases screened.

| Compound | R<sup>1</sup> | R<sup>2</sup> | R<sup>3</sup> | R<sup>4</sup> | $\Delta T_m$ [K] | IC<sub>50</sub> [nM] | S (5K) |
| --- | --- | --- | --- | --- | --- | --- | --- |
| **2** | H | H | F | H | 10.8 | 185 ± 10 | 0.28 |
| **12** | H | H | H | H | 9.3 | 350 ± 25 | 0.26 |
| **13** | H | H | Cl | H | 9.5 | 717 ± 14 | 0.37 |
| **14** | H | H | Me | H | 9.1 | 156 ± 33 | 0.22 |
| **15** | H | H | Et | H | 9.1 | 960 ± 33 | 0.23 |
| **16** | H | H | OMe | H | 4.3 | 1100 ± 217 | 0.10 |
| **17** | H | F | F | H | 10.3 | 253 ± 33 | 0.32 |
| **18** | H | Cl | F | H | 12.3 | 152 ± 44 | 0.36 |
| **19** | H | Br | F | H | 14.0 | 264 ± 19 | 0.31 |
| **20** | H | Me | F | H | 13.3 | 110 ± 11 | 0.42 |
| **21** | H | OMe | F | H | 9.3 | 331 ± 59 | 0.28 |
| **22** | H | tBu | H | H | 12.0 | 886 ± 63 | 0.22 |
| **23** | F | H | F | H | 9.2 | 159 ± 0.2 | 0.16 |
| **24** | Me | H | F | H | 8.5 | 1190 ± 231 | 0.12 |
| **25** | F | H | F | Br | 9.0 | 818 ± 56 | 0.10 |
| **26**<br>(RN222) | F | Br | F | H | 10.7 | 134 ± 5 | 0.14 |
| **27** | H | Br | H | F | 10.2 | 1960 ± 652 | 0.09 |

However, despite significant improvements compared to our earlier series of hybrid type-II inhibitors, the selectivity of **26** (RN222) still fell short of our criteria for a LRRK2-specific compound. Therefore, another back pocket type-II binding motif was tested. Since the PF-360 hinge binding moiety with the linking phenyl ring in position 5 of the 7*H*-pyrrolo[2,3-*d*]pyrimidine provided successful on-target association with LRRK2^RCKW^ across most compounds in our second series of inhibitors, we kept this hinge binding scaffold and combined it with the DYG-binding motif of a different type-II inhibitor, Rebastinib.

#### PF-360-Rebastinib hybrid series

The initial compound **28** obtained from this chemical series, named RN129, significantly outperformed all previously designed and tested hybrid type-II inhibitors both in stabilizing LRRK2^KW^ and in inhibiting LRRK2 kinase activity (IC_50_= 59.7 nM) (**Table 3**). Unfortunately, RN129 (**28**) was not only by far the most potent LRRK2 inhibitor designed using our hybridization approach, but also the most promiscuous one, showing high thermal stabilization of 49 % of the kinases screened.

**Table 3.** Second PF-360-Rebastinib hybrid series. Compound structures, thermal shift data and *in vitro* IC_50_ values of PF-360-Rebastinib inspired type-II hybrid compounds. The selectivity was calculated as the number of kinases showing a thermal shift greater than 5 K divided by the total number of kinases screened.

| Compound | R<sup>1</sup> | R<sup>2</sup> | R<sup>3</sup> | $\Delta T_m$ [K] | IC<sub>50</sub> [nM] | S (5K) |
| --- | --- | --- | --- | --- | --- | --- |
| **28**<br>(RN129) |  |  | H | 20.0 | 59.7 ± 2.9 | 0.49 |
| **29** |  |  | H | 21.0 | 20.2 ± 0.01 | 0.36 |
| **30**<br>(RN277) |  |  | Cl | 18.0 | 135 ± 2 | 0.10 |
| **31** |  |  | Cl | 18.0 | 82.4 ± 19 | 0.15 |
| **32**<br>(RN341) |  |  | SEt | 19.9 | 296 ± 39 | 0.10 |
| **33** |  |  |  | 16.0 | 4.96 ± 0.11 | 0.14 |
| **34** |  |  |  | 12.5 | 7.21 ± 2.1 | 0.05 |
| **35** |  |  |  | 1.0 | n.d. | n.a. |
| **36** |  |  |  | 0.0 | 2820 ± 1703 | n.a. |
| **37** |  |  |  | 1.5 | n.d. | n.a. |

To elucidate the binding mode of RN129 (**28**) and to inform future design strategies, we obtained a co-crystal structure of RN129 (**28**) with one of its potent off-targets, CDC-like kinase 3 (CLK3), a kinase that has been crystallized routinely in our group (**Figure 2 A**). Our structure revealed that RN129 (**28**) has a canonical type-II binding mode that has not been described for CLK3 previously. As expected, the 7*H*-pyrrolo[2,3-*d*]pyrimidine moiety of RN129 (**28**) formed two hydrogen bonds with the hinge of CLK3. The linking phenyl ring formed a π-π-interaction with the phenylalanine F236. The urea was located between the αC-helix and the DFG-motif, with the bulky tail of the molecule extending into the far back-pocket. The hinge of CLK3 contains a leucine residue (L238) that is also present in LRRK2, whereas most human kinases have larger phenylalanine or tyrosine residues at this position. Therefore, we hypothesized that developing the inhibitor towards this residue would improve selectivity. Since RN129 (**28**) has a large molecular weight of 588 Da, we aimed to reduce the molecular weight of the inhibitor before specifically targeting it towards the leucine within the hinge region. Instead of the quinoline in the far back-pocket we installed a toluene (**29**), which resulted in slight improvements in potency and selectivity compared to RN129 (**28**) (**Table 3**). Next, a chlorine atom was installed at position 2 (R^3^) of the 7*H*-pyrrolo[2,3-*d*]pyrimidine hinge binder to produce hybrid compound RN277 (**30**). Using the assays previously described we demonstrated that this compound was dramatically more selective (S (5K) = 10 %) than its predecessors, while maintaining excellent on-target affinity with ΔTm = 18 K and IC_50_ = 135 nM (**Table 3**).

**Fig. 2.**
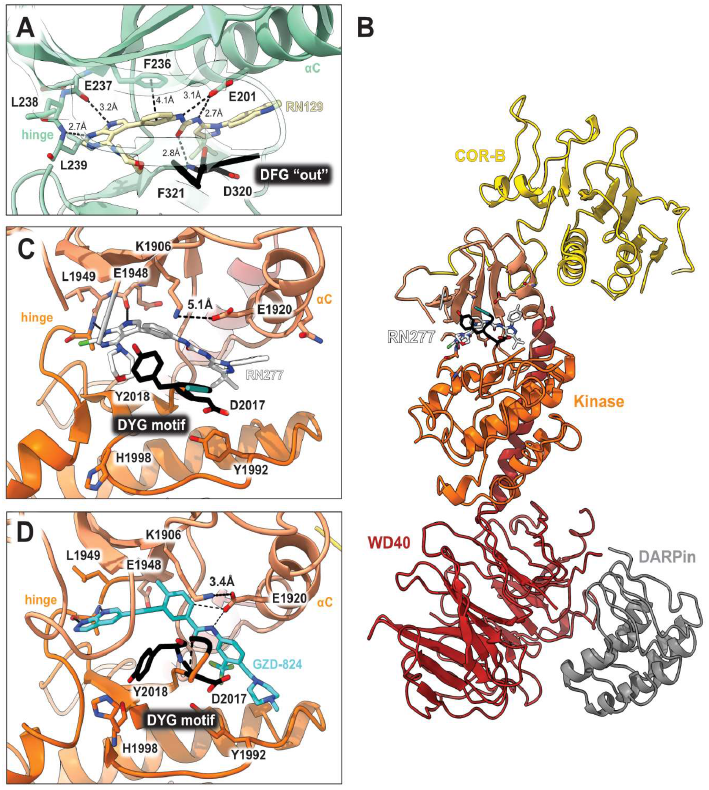
Co-crystal structure of RN129 bound to CLK3 and cryo-EM structure of RN277 bound to LRRK2RCKW. **(A)** The co-crystal structure of RN129 (28) with CLK3 highlighting the type-II binding mode and interactions between the protein and inhibitor (PDB: 9EZ3); **(B)** Ribbon diagram of the atomic model of LRRK2^RCKW^:RN277:E11 DARPin complex (PDB: 9DMI); **(C)** and (**D)** Close ups of the active sites of the cryo-EM structures of LRRK2^RCKW^:RN277 (**C**) and LRRK2^RCKW^:GZD824 (PDB: 8TZE) (**D**).

The high selectivity of RN277 (**30**) caused by the introduced chlorine in 2-position (R^3^) of the 7*H*-pyrrolo[2,3-*d*]pyrimidine was exploited by exchanging R^3^ with more space demanding moieties. Concurrently, the morpholine at the heterocycles 4-position was replaced by (di)methylamines, to save on the inhibitors molecular weight. Our resulting lead compound, the hybrid type-II inhibitor **32** (RN341, IC_50_ = 296 nM) was designed with a medium sized thiol ether residue at the 7*H*-pyrrolo[2,3-*d*]pyrimidine 2-position (R^3^). This compound maintained good on-target affinity (although slightly decreased compared to RN277) with high thermal stabilization of LRRK2^KW^ (ΔTm = 20 K), and an improved selectivity profile. RN341 (**32**) stabilized the same number of screened off-target kinases as RN277 (**30**, S (5K) = 10 %), but the degree of thermal stabilization of off-targets was lower (Supporting Information **Data S1**). Taken together, DSF data, IC_50_ values, and observed LRRK2-selectivity of RN277 (**30**) and RN341 (**32**) indicate that both inhibitors have significant potential as tool compounds for specifically targeting and inhibiting LRRK2 kinase activity. The synthesis of both derivatives is outlined in **Scheme 1**.

**Scheme 1.**
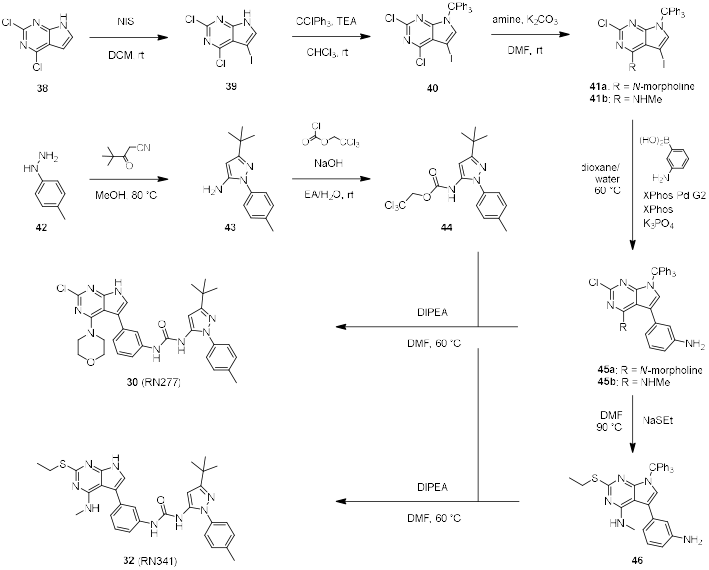
Synthesis of RN277 and RN341. The convergent synthesis route of compound 30 (RN277) and 32 (RN341). The detailed procedures and analytics are shown in the supplementary information. rt: room temperature, NIS: N-Iodosuccinimide, DCM: dichloromethane, TEA: tri-ethylamine, DMF: dimethylformamide, XPhos: dicyclohexyl[2′,4′,6′-tris(propan-2-yl)[1,1′-biphenyl]-2-yl]phosphane, EA: ethyl acetate, DI-PEA: N,N-diisopropylethylamine.

#### RN277 is a type-II inhibitor of LRRK2

To address how the new type-II inhibitor RN277 (**30**) binds to the kinase domain of LRRK2, we determined its structure by single particle cryo-EM, using purified recombinant LRRK2^RCKW^. The sample was prepared in the presence of guanosine diphosphate (GDP) as a ligand for the GTPase (ROC) domain. We solved this structure bound to a LRRK2-specific designed ankyrin repeat protein (DARPin) (**34**). This small protein binds specifically to the WD40 domain of LRRK2 and decreases the preferential orientation of un-ligated LRRK2 on cryo-EM grids, improving particle distribution of LRRK2^RCKW^ (**34**). The map resolution allowed us to distinguish the molecular details of the kinase, especially in its active site. The map had an overall resolution of 3.3 Å, with local resolution reaching 3 Å around the kinase active site (Supporting Information **Figure S1** and **Table S3**). We then performed two focused refinements on different parts of the map: one comprised the ROC and COR-A domains and the other the COR-B, kinase, and WD40 domains. We built molecular models into each of those locally refined maps, which had better resolutions than the corresponding portion of the global map, and combined them into the final model (**Figure 2 B, C**).

Our structure of LRRK2^RCKW^ bound to RN277 (**30**) revealed all the features of a type-II inhibitor binding mode, including an open kinase conformation with the catalytic triad DYG motif “out”, αC “out”, a broken R-spine, and a disordered activation loop (**Figure 2 B, C**). However, we also identify some features which differ from the previously reported structure of LRRK2^RCKW^ bound to the broad-spectrum type-II inhibitor GZD-824 (**18**). First, the distance between K1906 and E1920 is 5.1 Å in the RN277 (**30**) structure, while it is 3.4 Å in the GZD-824 structure, the latter corresponding to an electrostatic interaction (**Figure 2 C, D**). Second, there was no well-defined density for the G-rich loop in the N-lobe of the kinase in the RN277 (**30**) structure, while this motif was well resolved in the GZD-824 map. This suggests that the loop is flexible when LRRK2^RCKW^ is bound to RN277 (**30**). Third, the Y2018 of the DYG triad showed different conformations in the two structures (**Figure 2 C, D**). Finally, H1998 showed a single rotamer in the RN277 (**30**) structure, in contrast to the two coexisting rotamers we observed in LRRK2^RCKW^ bound to GZD-824 (**Figure 2 C, D**). Overall, our cryo-EM structure validated the type-II inhibitor binding mode of our new LRRK2-targeting compound RN277 (**30**) and identified features that differed from the interaction observed in the structure of GZD-824 with LRRK2.

#### Kinome selectivity profiling of RN341

As mentioned, the DSF selectivity screens of RN277 (**30**) and RN341 (**32**) showed the same number of kinases stabilized (> 5 K) by both compounds. However, the degree of stabilization favors RN341 (**32**) as the more selective hybrid type-II inhibitor (Supporting Information **Data S1**). To understand how the selectivity of our lead compound RN341 (**32**) for LRRK2 increased compared to commercially available type-II inhibitors, we measured the DSF thermal stabilization of its broad-spectrum mother compound, the type-II kinase inhibitor Rebastinib. The DSF selectivity screen for Rebastinib (**Figure 3 A**) revealed that 37 kinases were stabilized (> 5 K) across almost all kinase families. In contrast, RN341 (**32**) showed only ΔT_m_ values of >5K for 10 kinases, indicating a significantly improved selectivity profile (**Figure 3 B**). The kinome selectivity of RN341 (**32**) was further characterized in a ^33^PanQinase^TM^ activity assay screen of 350 kinases (ReactionBiology) at two different inhibitor concentrations (1 µM and 10 µM). The waterfall plot of this screen (**Figure 3 C**) showed that even at the high concentration of 10 µM, for RN341 (**32**) only 10 kinases were inhibited with less than 30 % residual activity remaining. At 1 µM RN341 (**32**) only Threonine Tyrosine Kinase (TTK) showed inhibition less than 50 % of control activity. Surprisingly, RN341 (**32**) was not detected as a LRRK2 inhibitor in the ^33^PanQinase^TM^ screen. LRRK2 is a large and unstable protein *in vitro*, and it is likely that the kinase was inactive in this selectivity panel. We confirmed the potent activity of RN341 (**32**) on LRRK2 in multiple orthogonal assay formats. The potential major off-target kinases (with ΔTm >5 K or % control kinase activity < 22 % at 10 µM RN341) from both the DSF-screen and the ^33^Pan-Qinase^TM^ screen were validated in dose response assays using a cellular nanoBRET^®^ assay (**35**) (**Figure 3 D**). The determined EC_50_ values of RN341 (**32**) indicate that of the 12 off-target kinases identified, only 4 (STK10, MAPK14, JNK2 and TTK) were inhibited by RN341 (**32**) with an EC_50_ in the low micromolar range (< 5µM).

**Fig. 3.**
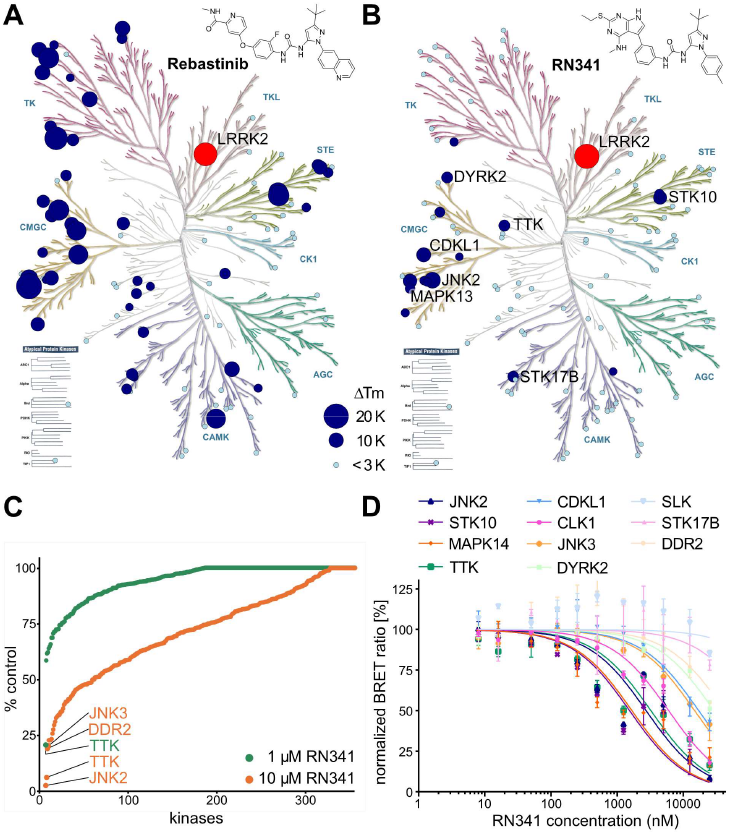
Kinome selectivity of RN341. **(A)** Kinome phylogenetic tree, with 96 kinases screened in the DSF assay against Rebastinib highlighted in blue. The 18.5 K ΔTm shift of LRRK2^KW^ is highlighted in red. For all screened kinases, the bubble size correlates with the degree of ΔTm shift, as indicated in the legend; **(B)** Kinome phylogenetic tree, with 103 kinases screened in the DSF assay against RN341 highlighted in blue. The 20 K ΔTm shift of LRRK2^KW^ is highlighted in red. The bubble size for each kinase correlates with the ΔTm shifts, as indicated in the legend (as in **A**). Kinases with ΔTm > 6 K are labeled; **(C)** Waterfall plots of the ReactionBiology ^33^PanQinase^TM^ screen of RN341 at 1 µM and 10 µM against 350 wild-type kinases. Kinases with decreased activity in the presence of RN341 to < 22 % of the control value are labeled; **(D)** Off-target validation from both screens via *in cellulo* nanoBRET assay in 2 biological replicates, error bars ± sd, EC_50_ (JNK2) = 2.7 µM, EC_50_ (STK10) = 1.5 µM, EC_50_ (MAPK14) = 1.7 µM, EC_50_ (TTK) = 3.2 µM, EC_50_ (CDKL1) = 17 µM, EC_50_ (CLK1) = 6.0 µM, EC_50_ (JNK3) = 15 µM, EC_50_ (DYRK2) = >20 µM, EC_50_ (SLK) >20 µM, EC_50_ (DDR2) >20 µM, EC_50_ (STK17B) = >20 µM.

#### In vitro and in cellulo inhibition of LRRK2 kinase activity by RN277 and RN341

The best characterized physiological substrates of LRRK2 are a subset of small Rab GTPases (**36**). To investigate the ability of RN277 (**30**) and RN341 (**32**) to inhibit LRRK2 kinase activity towards its physiological substrates *in vitro*, we incubated a dilution series of the compounds with recombinant LRRK2^RCKW^ and Rab8a, and quantified Rab8a phosphorylation (pT72) by mass spectrometry. As anticipated, increasing quantities of each compound resulted in decreased Rab8a phosphorylation, indicative of inhibited LRRK2 kinase activity. RN277 (**30**) displayed a slightly better inhibitory effect (IC_50_ = 70 nM) compared to RN341 (**32**, IC_50_ = 110 nM) (**Figure 4 A, B**).

**Fig. 4.**
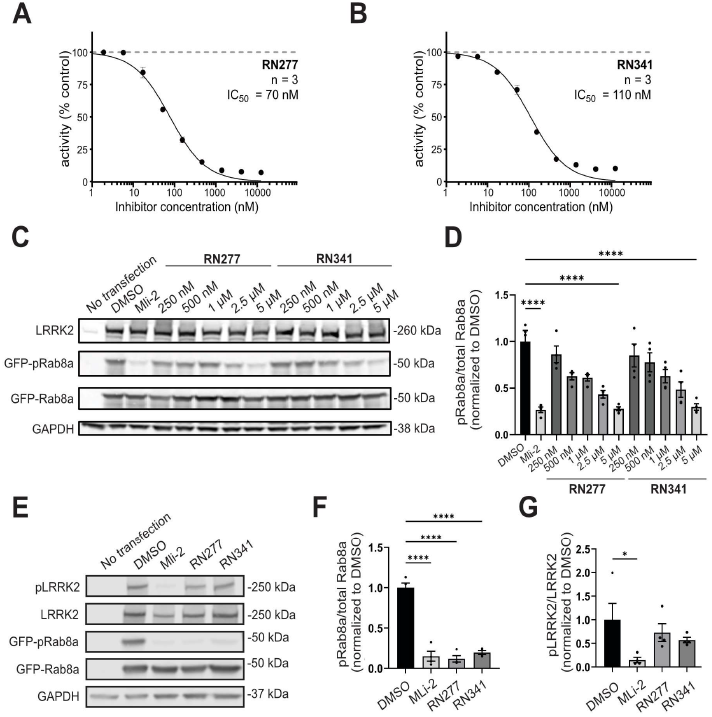
Inhibition of LRRK2’s phosphorylation of Rab8a in vitro and in cellulo. **(A)** and **(B)** Dose response curve of RN277 (**30**) and RN341 (**32**) inhibiting LRRK2^RCKW^-mediated phosphorylation of Rab8a. Activity was calculated as the percentage (%) of phosphorylated Rab8a vs. non-phosphorylated Rab8a detected in the presence of different concentrations of RN277/RN341; **(C)** Western blots from 293T cells transiently co-transfected with LRRK2 (full-length) and GFP-Rab8a for 48h prior to treatment with a dilution series of RN277 (**30**) and RN341 (**32**) for 4h. DMSO and MLi-2 (500 nM) treatment for 4h were used as negative and positive controls, respectively. Lysed cells were immunoblotted for LRRK2, total GFP-Rab8a, phospho-Rab8a (pT72) and GAPDH as a loading control; **(D)** Quantification from four independent western blots showing the ratio of GFP-pRab8a to total GFP-Rab8a upon treatment with RN277 (**30**) and RN341 (**32**) at the indicated concentrations. Statistical analysis was performed using one-way ANOVA analysis with Tukey’s multiple comparison of means. ****p<0.0001, error bars ± s.e.m. **(E)** Western blots from 293T cells transiently co-transfected with LRRK2 (full-length) and GFP-Rab8a, treated with DMSO (control), 500 nM MLi-2, 5 µM RN277 or 5 µM RN341 for 4h, 48h post-transfection. Lysed cells were immunoblotted for LRRK2, phospho-LRRK2 (pS935), total GFP-Rab8a, phospho-Rab8a (pT72) and GAPDH as a loading control; **(F)** Quantification from four independent western blots showing the ratio of GFP-pRab8a to total GFP-Rab8a upon treatment with the indicated inhibitors (as in **D**). Statistical analysis was performed using a one-way ANOVA analysis with Tukey’s multiple comparison of means. ****p<0.0001, error bars ± s.e.m.; **(G)** Quantification from four independent western blots showing the ratio of pLRRK2 to total LRRK2 upon treatment with the indicated inhibitors. Statistical analysis was performed using a one-way ANOVA analysis with Tukey’s multiple comparison of means. *p 0.0469, error bars ± s.e.m. Note that data from one replicate shown in Fig. 4D forms part of the dataset presented in F/G.

Next, we determined whether RN277 (**30**) and RN341 inhibited LRRK2 kinase-mediated phosphorylation of Rab8a in cells. To do this, we co-transfected 293T cells with full-length, untagged LRRK2 and GFP-Rab8a. Transfected cells were treated with increasing concentrations of RN277 (**30**) and RN341 (**32**), and the level of Rab8a phosphorylation was assessed by western blotting with a phospho-specific Rab8a antibody. Toxicity did not appear to be an issue with either of these compounds, at any of the concentrations used.

Both compounds reduced phosphorylation of Rab8a in a dose-dependent manner (**Figure 4 C, D**). At 5 µM, both RN277 (**30**) and RN341 (**32**) reduced the level of phosphorylated Rab8a to almost background levels, as seen following treatment with the well-characterized type-I inhibitor MLi-2 (**Figure 4 C-F**). In addition, we tested the effect of RN277 and RN341 (**32**) on phosphorylation of the LRRK2 biomarker site S935 using a phospho-specific LRRK2 S935 antibody. The ability to inhibit phosphorylation of LRRK2 substrate Rabs (such as Rab8a) without significantly affecting the phosphorylation of LRRK2 biomarker sites is a known feature of type-II LRRK2 kinase inhibitors (**31**). In contrast to treatment with 500 nM MLi-2, which led to almost complete dephosphorylation of LRRK2 S935, neither RN277 (**30**) nor RN341 (**32**) significantly reduced the phosphorylation level at this site, identifying this site as a specific marker for type-I LRRK2 inhibition (**Figure 4 E, G**).

#### Type-II inhibitors rescue LRRK2 inhibited kinesin motility *in vitro*

Previous work shows that LRRK2^RCKW^ can associate with microtubules (*19, 23*–*25*) and block dynein and kinesin motor protein movement *in vitro* (**19**). As demonstrated with non-specific inhibitors in these studies, type-II inhibitors, which stabilize the open kinase conformation, rescue motor protein motility, while type-I inhibitors that stabilize the closed kinase form, do not (**19**). Given our structural evidence that RN277 (**30**) stabilized the open kinase conformation of LRRK2 (**Figure 3 B, C**), we hypothesized that our developed selective type-II compounds would prevent LRRK2 filamentation on microtubules and therefore relieve LRRK2^RCKW^ dependent inhibition of kinesin motility in single-molecule *in vitro* motility assays (**Figure 5 A**). As previously described (*19, 25*), LRRK2^RCKW^ significantly impedes kinesin motility (**Figure 5 A-E**). The LRRK2-specific type-I inhibitor MLi-2, which stabilizes the closed kinase conformation (**18**), did not rescue this effect (**Figure 5 B, C**). In contrast, RN277 (**30**), RN341 (**32**) and the commercially available broad-spectrum type-II kinase inhibitor Ponatinib rescued kinesin motility from the inhibition observed in the presence of LRRK2^RCKW^ (**Figure 5 D, E**). Thus, our data showed that RN277 (**30**) and RN341 (**32**) displayed a key hallmark of type-II kinase inhibitors, namely an ability to rescue kinesin motility in the presence of LRRK2^RCKW^, likely due to stabilization of the open kinase conformation that prevents LRRK2 filamentation on microtubules.

**Fig. 5.**
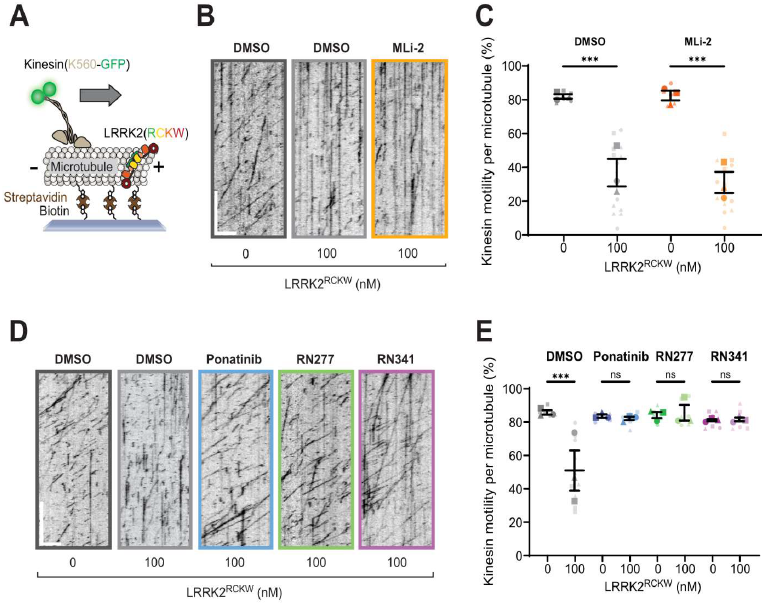
LRRK2-specific type-II inhibitors RN277 and RN341 rescue kinesin motility in the presence of LRRK2^RCKW^. **(A)** Schematic of the single-molecule *in vitro* motility assay; **(B)** Example kymographs from single-molecule motility assays showing kinesin motility with DMSO or the type-I inhibitor MLi-2 (5 µM) in the presence or absence of LRRK2^RCKW^. Scale bars 5 µm (x) and 30 s (y); **(C)** Quantification of the percentage (mean ± s.e.m) of motile events per microtubule as a function of LRRK2^RCKW^ concentration in the absence (DMSO) or presence of MLi-2 (5 µM). Three technical replicates were collected per condition, with data points represented as circles, triangles and squares corresponding to single data points (microtubules) within each replicate. Statistical analysis was performed using the mean of each technical replicate; DMSO condition ***p 0.0007, MLi-2 condition ***p 0.0003, One-way ANOVA with Sidaks multiple comparison test within drug only; **(D)** Example kymographs from single-molecule motility assays showing kinesin motility with DMSO or the type-II inhibitors Ponatinib, RN277 and RN341 (5 µM) in the presence or absence of LRRK2^RCKW^. Scale bars 5 µm (x) and 30 s (y); **(E)** Quantification of the percentage (mean ± s.e.m) of motile events per microtubule as a function of LRRK2^RCKW^ concentration in the absence (DMSO) or presence of type-II inhibitors Ponatinib, RN277 and RN341 (5 µM). Three technical replicates were collected per condition, with data points represented as circles, triangles and squares corresponding to single data points (microtubules) within each replicate. Statistical analysis was performed using the mean of each technical replicate; ***p 0.0003, One-way ANOVA with Sidaks multiple comparison test within drug only.

## Discussion

Here, we presented the development and characterization of the first selective type-II LRRK2 kinase inhibitors RN277 and RN341. The success of our structure-directed approach combining parental LRRK2-specific type-I and broad-spectrum type-II inhibitors demonstrated a general rational design strategy that can be further exploited to develop new type-II inhibitors based on type-I inhibitors with favorable pharmacological properties. Due to its interaction with the allosteric and often diverse DFG-out pocket, the type-II binding mode has been initially described as a strategy for the development of more selective inhibitors. However, kinome wide selectivity studies showed that type-II DFG-out pockets are accessible in most kinases and that type-II inhibitors are often more promiscuous than type-I compounds (*20, 37*). Despite the excellent selectivity of available type-I inhibitors, the development of selective type-II inhibitors inspired by the hinge binding moieties of type-I inhibitors was a formidable medicinal chemistry effort. However, we believe that the PD field will benefit from our lead compounds by enabling new potential avenues for research and drug development that have, until now, been impeded by the many off-targets and the toxicity of commercially available type-II LRRK2 inhibitors.

Both RN277 and RN341 showed excellent potency and selectivity. Both compounds bound to LRRK2, increasing the thermostability of LRRK2^KW^ in DSF assays (**Table 3**), and inhibited the phosphorylation of Rab8a by LRRK2^RCKW^ *in vitro* with 70 nM and 110 nM IC_50_ values respectively (**Figure 4**). Both compounds also inhibited LRRK2 phosphorylation of Rab8a in cells (**Figure 4**). In two kinome selectivity screens of 103 and 350 kinases, only four major off-targets were identified for RN341 (STK10, MAPK14, JNK2 and TTK) with cellular activity in the micromolar EC_50_ region (**Figure 3**). This represents a drastically improved selectivity profile compared to the broad-spectrum LRRK2 type-II inhibitors currently available.

Both RN277 and RN341 displayed the key phenotypic and structural hallmarks of LRRK2 type-II kinase inhibitors. Both compounds preserve phosphorylation at the biomarker S935 site on LRRK2 (**Figure 4**), a phenomenon that was previously described for other broad spectrum LRRK2 type-II inhibitors. This phenotype demonstrates a key difference in the downstream effects of type-I and type-II LRRK2 inhibition, with type-I LRRK2 inhibitors inducing dephosphorylation at these sites (**31**). Although the underlying mechanism for these differences is not known, it is hypothesized that the conformational changes in LRRK2 induced by type-I and type-II inhibitors modulate the accessibility of LRRK2 biomarker sites to upstream phosphatases and/or kinases. Our new LRRK2-specific type-II inhibitors represent key tools that could be used to assess the conformation-specific accessibility and association of LRRK2 with these upstream modifiers. Additionally, both RN277 and RN341 rescue the motility of kinesin on microtubules in the presence of LRRK2^RCKW^, an effect that is specific to type-II inhibitors due to their promoting the open kinase conformation (*19, 25*) (**Figure 5**). An open question in the field is whether the increased size and number of cytosolic lamellar bodies in the lungs of non-human primates following repeated dosing with type-I LRRK2 inhibitors (**32**) could result from LRRK2-induced blockage of microtubule-based intracellular transport. The future use and characterization of RN277 and RN341 should help in answering this critical question.

We also present a 3 Å resolution cryo-EM structure of LRRK2^RCKW^ bound to RN277 (**Figure 2**). Given the structural and chemical similarity of RN277 and RN341, we anticipate that they will bind LRRK2 in a largely comparable manner. In the LRRK2^RCKW^:RN277 structure, we show that RN277 displays the typical binding mode of a type-II inhibitor, including the positioning of the LRRK2 catalytic triad in the “DYG-out” position. However, we also observed conformational differences compared to the cryo-EM structure of LRRK2^RCKW^ bound to the broad-spectrum type-II inhibitor GZD-824 (**18**). This makes the new chemotype of RN277 and RN341 interesting for future investigations into the conformation-dependent regulation of LRRK2 and for the design of additional LRRK2-targeting inhibitory compounds.

The LRRK2-selective type-II inhibitors RN277 and RN341 represent a new set of tool compounds, which will open new avenues for therapeutic development for PD and that harbor significant research potential for uncovering the broader impact of using conformation-specific LRRK2 inhibitors *in vivo* and in cells.

## Supporting information

Supplementary_all

## Acknowledgements

We would like to thank Al Garofalo for providing assay platforms in the early stages of the project. For the help with enzymatic activity assays, we thank Benedict-Tilman Berger. We appreciate Leoni Fritsche for supporting the synthesis of three derivatives biologically tested in this study during an internship under the supervision of NDR.

We also thank the UC San Diego Cryo-EM Facility, and the UC San Diego Physics Computing Facility for IT support.

This paper was typeset with the bioRxiv word template by @Chrelli: (www.github.com/chrelli/bioRxiv-word-template).

## Author contributions

Writing and editing: NDR, KJS, MSM, AEL, SLR-P and SK Conceptualization: AEL, SLR-P and SK

Synthesis: NDR

Investigation and analysis: NDR, KJS, MSM, VD, AK, MS, LE and DC Supervision: SM, TH, AEL, SLR-P and SK

Funding acquisition: AEL, SLR-P and SK

## Materials and Methods

### Compound synthesis and analysis

The synthesis schemes (Supporting Information **Scheme 1-6**) and procedures including all intermediate analytics as well as the NMR spectra, HPLC/MS analysis and HRMS data of all hybrid type-II inhibitors is shown in the supporting information. All biologically tested compounds exceeded 95 % purity in HPLC UV-spectra, showed clean NMR spectra and the mass error of the high-resolution mass spectrum was below 5 ppm. (The synthesis and structural characterization of the LRRK2-selective compounds RN277 and RN341 is available at https://dx.doi.org/10.17504/protocols.io.36wgqnxzxgk5/v1).

### Protein expression and purification from insect cells

The LRRK2^KW^ (RRID: Addgene_228879) or LRRK2^RCKW^ (RRID: Addgene_226784) expression constructs (Supporting Information **Table S4**) encoding a TEV cleavable N-terminal His_6_-Z-tag, were expressed in SF9 insect cells (RRID:CVCL_0549) after baculoviral transfection as described previously (**18**). In brief, the virus was generated according to Bac-to-Bac expression system protocols (Invitrogen). For protein expression, cells were infected at a density of 2×10^6^ cells/mL with recombinant virus. Infected cells were cultured in serum-free Insect-XPRESS Medium (Lonza) at 27 °C shaking with 90 rpm for 66 h. After harvesting by centrifugation, the cell pellet was washed with PBS before sonication in lysis buffer (containing 20 mM HEPES [pH7.4], 500 mM NaCl, 20 mM imidazole, 0.5 mM TCEP, 5 % glycerol, 20 µM GDP, 5 mM MgCl_2_). The lysate was clarified by ultra-centrifugation for 1 h at 100000 g at 4°C and loaded onto pre-equilibrated Ni-NTA Sepharose (Quiagen). After stringent wash with lysis buffer, the protein was eluted in lysis buffer supplemented with 300 mM imidazole. After reducing the NaCl concentration to 250 mM the eluate was loaded onto a SP sepharose column (Cytiva) and eluted by applying a salt gradient ranging from 250 mM to 2.5 M. The fractions containing the protein of interest were pooled and N-terminal tags were removed with TEV protease incubation overnight. Cleaved tag, contaminating proteins and uncleaved protein were removed by running a combination of SP sepharose Ni-NTA column. Finally, the protein was subjected to size-exclusion chromatography using a S200 16/200 column (GE Healthcare) in 20 mM HEPES (pH7.4), 800 mM NaCl, 0.5 mM TCEP, 0.5 % glycerol, 20 µM GDP, 2.5 mM MgCl_2_ coupled to an AKTA Xpress system. (For a step-by-step protocol see: dx.doi.org/10.17504/protocols.io.14egn314pl5d/v1).

### Protein expression and purification from bacteria

The Rab8A substrate, encoding a TEV cleavable N-terminal His_6_-tag (RRID: Addgene_228880, Supporting Information **Table S4**) was expressed in *E. coli* Rosetta cells (Novagen, 70954, https://www.emdmilli-pore.com/US/en/product/RosettaDE3-Competent-Cells-No-vagen,EMD_BIO-70954). Cells were grown shaking at 37 °C until OD_600_=0.8 was reached. After reducing temperature to 18 °C protein expression was induced by adding IPTG and cells were incubated overnight shaking at 180 rpm. Cells were harvested by centrifugation. Ni-NTA purification was essentially carried out as described above. The eluate from Ni-NTA was dialyzed straight away over night while the N-terminal tags were removed by adding TEV protease. Contaminating protein, the cleaved tag as well as non-cleaved protein were removed by Ni-NTA before the protein was concentrated and subjected to size-exclusion chromatography as described above. (For step-by-step protocol see: dx.doi.org/10.17504/protocols.io.261ge514yg47/v1).

The GFP-tagged human kinesin construct (K560-GFP; RRID:Addgene_15219) (Supporting Information **Table S4**) was purified from BL21-CodonPlus (DE3)-RIPL (Agilent 230280, https://www.agilent.com/store/en_US/Prod-230280/230280) *E*.*coli* as previously described (**38**). In brief, transformed *E*.*coli* cultures were grown at 37 °C to OD_600_ of 0.6-0.8 before induction with 0.75 mM IPTG for 16 h at 18 °C. After harvesting by centrifugation, *E. coli* pellets were resuspended in lysis buffer (50 mM Tris [pH7.5], 250 mM sodium chloride, 1 mM magnesium chloride, 20 mM imidazole [pH 7.4]) supplemented with 0.5 mM Mg-ATP, 10 mM beta-mercaptoethanol, 1 mM Pefabloc, complete EDTA-free protease inhibitor cocktail tablet (Roche) and 1 mL of 50 mg/mL lysozyme) and lysed by sonication on ice. Lysates were clarified by centrifugation at 92,600 x g for 30 min at 4 °C and incubated with Ni-NTA agarose beads (Qiagen) for 1 h at 4 °C. Beads and lysate were then transferred to a gravity flow column and washed repeatedly in lysis buffer supplemented with 0.5 mM Mg-ATP, 10 mM beta-mercaptoethanol, and 1 mM Pefabloc. After washing, protein was eluted in elution buffer (50 mM Tris [pH7.5], 250 mM sodium chloride, 1 mM magnesium chloride, 250 mM imidazole [pH7.4]) supplemented with 0.1 mM Mg-ATP, and 10 mM beta-mercaptoethanol. The eluate was transferred to a PD-10 column (GE Healthcare) for desalting and buffer exchanged into BRB80 (80 mM K+PIPES [pH6.8], 2 mM magnesium chloride, 1 mM EGTA) supplemented with 0.1 mM Mg-ATP, 0.1 mM DTT and 10 % sucrose. (For step-by-step protocol see: dx.doi.org/10.17504/protocols.io.4r3l2qr1xl1y/v1).

### Compound screening by differential scanning fluorometry (DSF)

To characterize binding between small molecules and LRRK2^KW^, induced thermal stabilization was measured by differential scanning fluorimetry (DSF) according to previously established protocols (**39**). In brief, LRRK2^KW^ was diluted to 4 µM in 20 mM HEPES [pH7.4], 150 mM NaCl, 0.5 mM TCEP, 0.5 % glycerol, 20 µM GDP, 2.5 mM MgCl_2_ and SYPRO Orange was added (1:1000, Millipore Sigma). 10 µM inhibitor to be tested was added to a final sample volume of 20 µL per well of a 96 well plate (white, Starlab). Fluorescence was monitored with excitation and emission filters set at 465 and 590 nm during gradual heating (1 K/min) from 25 °C to 95 °C in a MX3005P real-time PCR instrument (Stratagene). Data was evaluated using MXPro3005 software (Stratagene, https://www.agilent.com/cs/library/usermanuals/public/70225.pdf, RRID:SCR_016375). The fluorescence was plotted against the temperature and the melting point (T_M_) was determined as the minimum of the first derivative relative to the control. (For step-by-step protocol see: dx.doi.org/10.17504/protocols.io.kxygx3y6kg8j/v1).

### Biochemical LRRK2 activity assay

The biochemical activity assay was based on the phosphorylation reaction between LRRK2^RCKW^ and the SOX-based substrate peptide (AQT0615, Assay Quant Technologies). A dilution series between 15 µM and 0.4 nM of eleven concentrations of the compounds was pipetted into white 384-well plates (Greiner 781207) as duplicates with an ECHO acoustic dispenser (Labcyte). Two wells per compound were used as 0 % (without protein and compound) and 100 % (without compound) control. The purified LRRK2^RCKW^ was diluted to 22 nM with the reaction buffer (50 mM HEPES buffer [pH7.5], 10 mM MgCl_2_, 1 % glycerol, 1 mM DTT, 0.2 mg/mL BSA,

0.01 % Tween20, 5 µM AQT0615) and 10 µL were added in each well with a multichannel pipette (Eppendorf). The 0 % control wells were filled with 10 µL of pure reaction buffer. With the ECHO 5 nL 100 mM ATP was pippeted into each well, the plates were centrifuged at 1500 g_n_ for 2 min and the reaction was incubated at room temperature for 4 h. The fluorescence of the SOX-peptide after 360 nm excitation was measured at 487 nm emission with a PHERAstar plate reader (BMG Labtech). The IC_50_ values were calculated from a non-linear regression of the log[inhibitor] vs. the normalized response with GraphPad Prism 8 (https://www.graphpad.com/, RRID:SCR_002798). (For step-by-step protocol see: dx.doi.org/10.17504/protocols.io.81wgbz8qngpk/v2).

### Differential scanning fluorometry (DSF) 100 kinase selectivity screen

Recombinant protein kinase domains with a concentration of 2 µM were mixed with a 20 µM compound solution in DMSO, 20 mM HEPES [pH7.5], and 500 mM NaCl. SYPRO Orange (5000×, Invitrogen) was added as a fluorescence probe (1 µL per mL). Subsequently, temperature-dependent protein unfolding profiles were measured using the QuantStudio™ 5 realtime PCR machine (Thermo Fisher, Waltham, MA, USA). Excitation and emission filters were set to 465 and 590 nm. The temperature was raised with a step rate of 3 °C per minute. Data points were analyzed with the internal software (Protein Thermal Shift™ Software v1.4, https://www.thermofisher.com/order/catalog/product/4466038) using the Boltzmann equation to determine the inflection point of the transition curve (**40**). Differences in melting temperature are given as ΔTm values in °C. Measurements were performed in duplicates. (For step-by-step protocol see: https://www.eubopen.org/sites/www.eubopen.org/files/attachments/2021/DSF-inhibitorScreening_v1.pdf).

### Co-Crystallization with CLK3

CLK3 kinase domain (residues 127–484) with a TEV-cleavable N-terminal His6-tag (RRID: Addgene_38831; Supporting Information **Table S4**) was transformed into BL21(DE3) cells. Bacteria were grown in Terrific Broth medium (Thermo Fisher) containing 50 mg/mL kanamycin. Protein expression was induced at an OD600 = 2 using 0.5 mM IPTG at 18 °C for 12 h. Cells were lysed by sonication in lysis buffer containing 50 mM HEPES [pH7.5], 500 mM NaCl, 25 mM imidazole, 5 % glycerol, 50 mM L-glutamic acid, 50 mM L-arginine, and 0.5 mM TCEP. After centrifugation, the supernatant was loaded onto a Nickel-Sepharose column (Thermo Fisher) equilibrated with 30 mL lysis buffer. The column was washed with 60 mL lysis buffer. Protein was eluted by an imidazole step gradient (50, 100, 200, 300 mM). CLK3 was dialysed against SEC buffer (50 mM HEPES [pH7.5], 500 mM NaCl, 5 % glycerol, 50 mM L-glutamic acid, 50 mM L-arginine, and 0.5 mM TCEP) overnight and TEV protease was added to remove tag. After a reverse Ni-rebinding column to remove TEV protease and uncleaved protein, the protein was further purified by size exclusion chromatography using a S75 16/60 HiLoad column (GE Healthcare). Pure protein fractions were pooled together and concentrated to ∼11 mg/mL.

CLK3 (∼11 mg/mL) was co-crystallized with RN129 at 4 °C using the sitting-drop vapor diffusion method by mixing protein and well solutions in 2:1, 1:1, and 1:2 ratios (drop size 200 nL). The compound was incubated prior to crystallization with the protein for approx. 30 min on ice (final concentration ∼500 µM). The reservoir solution contained 15 % PEG 6k and 0.1 M Bicine buffer pH8. Ethylene glycol (25 %) was used as cryoprotectant for crystals before flash freezing in liquid nitrogen. Diffraction data were collected at beamline X06SA (Swiss Light Source) at a wavelength of 1.0 Å at 100 K (https://doi.org/10.5281/zenodo.13934795). Data were processed using XDS (version Jan. 10 2022, http://xds.mpimf-heidel-berg.mpg.de/, RRID:SCR_015652) (**41**) and scaled with aimless (version 0.8.2, http://www.ccp4.ac.uk/html/aimless.html, RRID:SCR_015747) (**42**). The PDB structure with the accession code 6YU1 (**43**) was used as an initial search MR model using the program MOLREP (version 11.0, https://www.ccp4.ac.uk/html/molrep.html) (**44**). The final model was built manually using Coot (version 0.8.9, http://www2.mrc-lmb.cam.ac.uk/personal/pemsley/coot/, RRID:SCR_014222) (**45**) and refined with REFMAC5 (http://www.ccp4.ac.uk/html/refmac5/description.html, RRID:SCR_014225) (**46**). Data collection and refinement statistics are summarized in **Table S5** (Supporting Information). Dictionary files for ligands were generated using the Grade Web Server (http://grade.glob-alphasing.org). (Protocol available at https://www.protocols.io/pri-vate/E313820E8AD611EFB8100A58A9FEAC02).

### Cryo-EM

Purified LRRK2^RCKW^ was exchanged into 20 mM HEPES [pH7.4], 150 mM NaCl, 0.5 mM TCEP, 5 % glycerol, 2.5 mM MgCl2 and 20 μM GDP. LRRK2^RCKW^ was incubated with E11-DARPin (RRID: Addgene_226784) in a 1:1.2 molar ratio and RN277 (30 μM) for 10 min at room temperature and 15 min at 4 °C. Complex was diluted to a final concentration of 4.5 μM in the same buffer before plunge freezing. 3 µL of LRRK2^RCKW^:RN277:E11 DARPin complex were applied to a glow-discharged UltraAuFoil Holey Gold 200 mesh R2/2 grid (Quantifoil, Q250AR2A) and incubated in a FEI Vitrobot IV chamber at 4 °C and 95 % humidity for 20 sec. The excess liquid was blotted for 4 sec using filter paper 595 at blot force 5 and vitrified by plunging into liquid ethane cooled down to liquid nitrogen temperature. (Protocol available at dx.doi.org/10.17504/protocols.io.yxmvm35n9l3p/v1).

Cryo-EM data for LRRK2^RCKW^:RN277:E11 DARPin was collected on a Titan Krios G3 (Thermo Fisher Scientific) operated at 300 keV, equipped with a Falcon 4 direct electron detector (Thermo Fisher Scientific) and a Gatan BioContinuum energy filter. Images were collected at defocus values varying between -1.0 to -2.0 μm at a nominal magnification of 130,000x in EF-TEM mode (0.935 Å calibrated pixel size) using a 20-eV slit width in the energy filter and a cumulative electron exposure of ∼55 electrons/Å2. Data was collected automatically using EPU software (Thermo Fisher Scientific, https://www.thermofisher.com/us/en/home/electron-micros-copy/products/software-em-3d-vis/epu-software.html). (Protocol available at dx.doi.org/10.17504/protocols.io.14egn3m5ql5d/v1).

13,972 movies were collected for LRRK2^RCKW^:RN277:E11 DARPin and preprocessed using Patch MotionCor and Path CTF estimation to align and estimate CTF respectively, both available in CryosPARC (**47**) (version 4.3, https://cryosparc.com/, RRID:SCR_016501). Micrographs with a CTF fit worse than 3.5 Å were excluded for further processing. Particles were picked using a Topaz (**48**) model (version 0.2.5, https://cb.csail.mit.edu/cb/topaz/) previously trained for the dataset. Several rounds of reference-free 2D classification yielded a stack of 412,439 particles, which were further extracted with a 400-pixel box using CryoSPARC (**47**) (Supporting Information **Figure S1**). Subsequently, an ab initio and heterogeneous refinement jobs were run to remove bad particles. Particles from two output maps were used as an input for NU-Refinement (C1 symmetry). After 3D Variability and local refinement, we obtained maps at 3.35 and 3.65 Å, respectively. In all maps, FSC and local resolution estimations were performed using the routines implemented in CryoSPARC (**47**).

LRRK2^RCKW^ model was built using the highest-resolution local maps obtained for the complex. Available PDB 6VP7 was used as a starting point. Protein model was split into domains, docked into the corresponding cryo-EM maps using UCSF ChimeraX (**49**) (version 1.7, https://www.cgl.ucsf.edu/chimerax/, RRID:SCR_015872) and merged. For DARPin E11, we generated an initial model using ColabFold (**50**) (https://github.com/sokrypton/ColabFold, RRID:SCR_025453). To obtain an RN277 model, electronic Ligand Building and Optimization Workbench (elBOW) software available in Phenix (**51**) (version 1.21, https://www.phenix-online.org/, RRID:SCR_014224) was run using Simplified Molecular Input Line Entry System (SMILES) notation of molecule as an input. RN277 inhibitor model was fitted in the map density and incorporated in the model. A combination of manual inspection of amino acids in Coot (*45, 52*) (version 0.8.9, http://www2.mrc-lmb.cam.ac.uk/personal/pemsley/coot/, RRID:SCR_014222) and refinement of model into their maps in Phenix (**51**) was used to generate the final model. Data collection and refinement statistics are summarized in **Table S3** (Supporting Information). All structure-related figures were prepared using ChimeraX (**49**). (Protocol available at dx.doi.org/10.17504/protocols.io.81wgbx5m1lpk/v1).

### Kinome-wide selectivity profile

Compound RN341 was tested at two concentrations of 1 μM and 10 μM against a panel of 350 wild-type kinases in the ^33^PanQinase^™^ assay. The assay was performed by Reaction Biology (https://www.reactionbiology.com/services/kinase-assays/kinase-panel-screening/).

### nanoBRET off-target validation

The assay was performed as described previously (**53**). In brief, full-length kinases as listed in **Table S6** (Supporting Information) were obtained as plasmids cloned in frame with terminal NanoLuc-fusion (kind gift from Promega; Supporting Information **Table S6**). Plasmids were transfected into HEK293T cells (ATCC CRL-3216, RRID: CVCL_0063) using FuGENE 4K (Promega, E5911), and proteins were allowed to express for 20 h. Serially diluted inhibitor and corresponding tracer as listed in **Table S6** (Supporting Information) at the Tracer KD concentration taken from TracerDB (tracerdb.org) (**54**) were pipetted into white 384-well plates (Greiner 781207) using an ECHO acoustic dispenser (Labcyte). The corresponding protein-transfected cells were added and reseeded at a density of 2 × 10^5^ cells/mL after trypsinization and resuspending in Opti-MEM without phenol red (Life Technologies). The system was allowed to equilibrate for 3 h at 37 °C/5 % CO2 prior to bioluminescence resonance energy transfer (BRET) measurements. To measure BRET, NanoBRET NanoGlo Substrate + extracellular NanoLuc Inhibitor (Promega, N2540) was added as per the manufacturer’s protocol, and filtered luminescence was measured on a PHERAstar plate reader (BMG Labtech) equipped with a luminescence filter pair (450 nm BP filter (donor) and 610 nm LP filter (acceptor). Competitive displacement data were then graphed using GraphPad Prism 9 software (http://www.graphpad.com/, RRID:SCR_002798) using a normalized 3-parameter curve fit with the following equation: Y = 100/(1 + 10(X – log IC50)). (For step-by-step protocol see: https://www.eubopen.org/sites/www.eubopen.org/files/attachments/2021/NanoBRET-IC50_v1_0.pdf).

### Mass spectroscopy (MS)-based activity assay

To determine IC50 values for selected inhibitors activity assay was performed using LRRK2^RCKW^ and Rab8A substrate (Supporting Information Table **S4**). 20 nM LRRK2^RCKW^ was incubated with 5 µM substrate and a concentration series of the respective inhibitor in 20 mM HEPES [pH7.4], 100 mM NaCl, 0.5 mM TCEP, 0.5 % glycerol, 20 µM GDP, 2.5 mM MgCl2. The reaction was started by adding 1 mM ATP and incubated at room temperature for 4 h. The phosphorylation reaction was stopped by addition of MS buffer (0.1 % formic acid in water). 5 µL of reaction mix was injected into an Agilent 6230 Electrospray Ionization Time-of-Flight mass spectrometer coupled to a 1260 Infinity liquid chromatography unit (0.4 mL/min flow rate using a solvent gradient of water to acetonitrile with 0.1 % formic acid). Data was acquired using the MassHunter LC/MS Data Acquisition software and analysed using the BioConfirm vB.08.00 tool (both Agilent Technology, https://www.agilent.com/en/product/software-informatics/mass-spectrometry-software, RRID: SCR_016657).

Peak intensities of unphosphorylated and phosphorylated Rab8A were quantified, and the relative kinase activated calculated as ratio. To determine the IC50, a non-linear regression with variable slope was fitted to the datapoints with GraphPad Prism (http://www.graphpad.com/, RRID:SCR_002798). (For step-by-step protocol see: dx.doi.org/10.17504/protocols.io.6qpvr385ovmk/v2).

### Cell treatments and Western blot for Rab8a phosphorylation level in cells

293T cells (American Type Culture Collection CRL-3216, RRID: CVCL_0063) were transfected with 1000 ng of wild-type full-length LRRK2 (pcDNA5-LRRK2, RRID: Addgene_229019) and 500 ng GFP-Rab8A (Addgene, RRID: Addgene_49543) using PEI (Polyethylenimine, Polysciences) (Supporting Information **Table S4**). After 48 h, transfected cells were treated with either DMSO, 500 nM MLi-2 (Tocris), or various concentrations of RN277 or RN341 for 4 h at 37 °C and 5 % CO2. Following treatments, cells were washed three times with ice cold PBS. Cells were then lysed in 300 µL ice cold RIPA lysis buffer (50 mM Tris [pH8.0], 150 mM NaCl, 1 % Triton X-100, 0.1 % SDS, 0.5 % sodium deoxycholate, 1 mM DTT) in the presence of complete mini EASYpack protease inhibitor cocktail and the PhosSTOP EASYpack phosphatase inhibitor cocktail (Roche). Resuspended and lysed cells were clarified by centrifugation at 13,000 x g for 15 mins at 4ºC. 1XNuPAGETM LDS Sample Buffer and 1XNuPAGETM Sample Reducing Agent (Invitrogen) were added to the clarified supernatants and the samples were boiled at 95 ºC for 5 min and spun down at 13,000 x g for 10 mins. For western blotting, 15 µL of sample were loaded into a Nu-PAGE^TM^ 4-12 % gradient Bis-Tris gel (Invitrogen) and run at 120 V for 75 mins before transfer to PVDF transfer membrane (0.45 µm pore size, Millipore) for 240 mins at 80 V at 4ºC. The membrane was blocked in 5 % milk in 1X TBS (20 mM Tris [pH8.0], 150 mM NaCl) for 1 h shaking at room temperature. Primary antibodies, monoclonal rabbit anti-LRRK2 (Abcam ab133474, RRID: AB_2713963; 1:1000), monoclonal rabbit anti-LRRK2 phospho-S935 (Abcam ab133450, RRID: AB_2732035; 1:1000; run with identical conditions on a separate gel), monoclonal rabbit anti-Rab8a phospho-T72 (Abcam ab230260, RRID: AB_2814988; 1:1000), monoclonal rabbit anti-GAPDH (Cell Signalling Technology 2118, RRID: AB_561053; 1:1000) and monoclonal mouse anti-GFP (Santa Cruz Biotechnology sc-9996, RRID: AB_627695 1:1000) were diluted in 5 % milk in TBS and incubated with membrane for 16 h at 4 °C. Following incubation, membranes were washed three times with 1X TBS-T (20 mM Tris [pH8.0], 150 mM NaCl , 0.1% Tween 20) before incubation with IRDye 680RD goat anti-rabbit IgG (LI-COR 926-68071, RRID: AB_10956166; 1:10000) and IRDye 800CW goat anti-mouse IgG (LI-COR 926-32210, RRID: AB_621842; 1:10000) secondary antibodies in 5 % milk in 1X TBS for 1h at room temperature. Membranes were then washed three times in 1X TBS-T before imaging on Li-COR Odyssey CLx controlled by Imaging Studio software (https://licor.com/bio/odyssey-clx/?lang=en, RRID:SCR_015795). Image processing and quantification was performed in Image J version 2.14.0 (NIH; https://imagej.net/, RRID:SCR_003070). The signal intensity of each band was assessed using the Analyze>Gels tool. The band intensity for phosphorylated LRRK2 was measured and compared to total LRRK2 levels. Band intensity for phosphorylated Rab8a was measured and compared against total GFP-Rab8a (using anti-GFP antibody). Comparative statistical analysis of phosphorylation levels were analyzed using a one-way ANOVA and corrected using Tukey’s multiple comparison test in GraphPad Prism 10 (http://www.graphpad.com/, RRID:SCR_002798). (For current step-by-step protocol for LRRK transfection and western blotting see dx.doi.org/10.17504/protocols.io.kxygx9bozg8j/v1).

### TIRF microscopy

Imaging for single molecule motility assays was performed with an inverted microscope (Nikon, Ti-E Eclipse) equipped with a 100×1.49 N.A. oil immersion objective (Nikon, Plano Apo) and a MLC400B laser launch (Agilent), with 405 nm, 488 nm, 561 nm and 640 nm laser lines. The excitation and emission paths were filtered using appropriate single bandpass filter cubes (Chroma). The emitted signals were detected with an electron multiplying CCD camera (Andor Technology, iXon Ultra 888) and the stage *xy* position was controlled by a ProScan linear motor stage controller (Prior). Illumination and image acquisition was controlled by NIS Elements Advanced Research software (Nikon; https://www.microscope.healthcare.nikon.com/products/software/nis-elements/nis-elements-advanced-research, RRID:SCR_014329).

### Single-molecule motility assays

Single-molecule TIRF motility experiments were performed in flow chambers using the microscopy set-up described above. Flow chambers were made by adhering 11/2 thickness cover glass (Corning) to a SuperFrost Plus microscope slide (Electron Microscopy Sciences) with double sided tape. Prior to flow chamber assembly, cover glass was washed in 1 M HCl at 60 °C for 1 h and sonicated in 100 % ethanol for 10 min to reduce non-specific binding. Taxol-stabilized microtubules composed of ∼10 % biotin-tubulin (Cytoskeleton) for attachment to streptavidin-coated cover glass and ∼10 % Alexa405 (Thermo Fisher Scientific)-tubulin for visualization were prepared and motility assays were performed as previously described (**19**). In brief, flow chambers were functionalized by incubation with 1 mg/mL biotin-BSA (Sigma) for 3 min, followed by 0.5 mg/mL Strep-davidin for a further 3 min. Taxol-stabilized microtubules were diluted 1:100 in motility assay buffer (30 mM HEPES [pH7.4], 50 mM potassium acetate, 2 mM magnesium acetate, 1 mM EGTA, 10 % glycerol, 1 mM DTT and 20 μM Taxol) and incubated with the flow chamber for 3 min. Following incubation, the flow chambers with adhered microtubules were washed three times with LRRK2 buffer (20 mM HEPES [pH7.4], 150 mM NaCl, 1 mM TCEP, 5 % glycerol, 2.5 mM MgCl_2_ and 20 μM GDP) before incubation for 5 min with either LRRK2 buffer with DMSO or specified kinase inhibitors (0 nM LRRK2^RCKW^) or LRRK2 buffer containing LRRK2^RCKW^ and DMSO or kinase inhibitors as indicated. In all cases, DMSO or the indicated kinase inhibitors were incubated with LRRK2 buffer in the presence or absence of LRRK2^RCKW^ for 10 mins before addition and incubation within the flow chambers. Flow chambers were then washed with motility assay buffer supplemented with 1 mg/mL casein and K560-GFP added in the final imaging buffer (motility assay buffer supplemented with an oxygen scavenger system of 45 μg mL^−1^ glucose catalase (Sigma-Aldrich), and 1.15 mg mL^−1^ glucose oxidase (Sigma-Aldrich), 0.4 % glucose, 71.5 mM β-mercaptoethanol and 1 mM ATP). The final concentration of K560-GFP in the flow chamber was 1.9 nM. K560–GFP was imaged every 500 ms for 2 min with 20 % laser (488) power. (Protocol available at dx.doi.org/10.17504/protocols.io.5qpvok4dzl4o/v1).

### TIRF motility data analysis

Data was blinded prior to analysis. Kymographs were generated from motility movies and quantified for percent motility using ImageJ V.2.14.0 as previously described (*19, 55*). Processive, diffusive and non-motile events were manually annotated for calculation of the percentage of processive events per microtubule. Processive events were defined as runs that moved unidirectionally and did not exhibit directional changes. Diffusive events were those with at least one bidirectional change greater than 600 nm in each direction and non-motile events were defined as those that did not exhibit movement. Single molecule movements with multiple behaviors were counted as multiple events. For percentage motility per microtubule measurements, processive events were divided by total events (non-motile, diffuse and processive) per kymograph.

For statistical analysis, brightness and contrast were adjusted in ImageJ for all motility movies and kymographs. Statistical analyses were performed in GraphPad Prism 10 (http://www.graphpad.com/, RRID:SCR_002798). Specific statistical analysis descriptors, n values and p values can be found in the corresponding figure legends. All TIRF experiments were analyzed from three independent technical replicates.

## Data and materials availability

All data needed to evaluate the conclusions in the paper are present in the paper and/or the Supplementary Materials.

In addition, the cryo-EM maps have been deposited in the Electron Microscopy Data Bank under the following accession codes: EMD-47004 (RoC-COR-A focused refinement map; https://www.ebi.ac.uk/emdb/EMD-47004), EMD-47006 (consensus map; https://www.ebi.ac.uk/emdb/EMD-47006) and EMD-47025 (COR-B-Kinase-WD40-E11 focused refinement map; https://www.ebi.ac.uk/emdb/EMD-47025). The model has been deposited in the Protein Data Bank with the accession code 9DMI (https://www.rcsb.org/structure/9DMI).

The CLK3-RN129 crystal structure has been deposited in the Protein Data Bank with the accession code 9EZ3 (https://www.rcsb.org/structure/9EZ3) and collected dataset deposited at https://doi.org/10.5281/zenodo.13934795.

Primary data for tables and figures is available at https://doi.org/10.5281/zenodo.13765594.

