## Supplementary_all for "Type-II kinase inhibitors that target Parkinson’s Disease-associated LRRK2"


Nicolai D. Raig *et al.*


**This PDF file includes:**

Figs. S1
Tables S1 to S6
Schemes S1 to S6
Synthesis Instructions
NMR spectra
HPLC/MS
HRMS

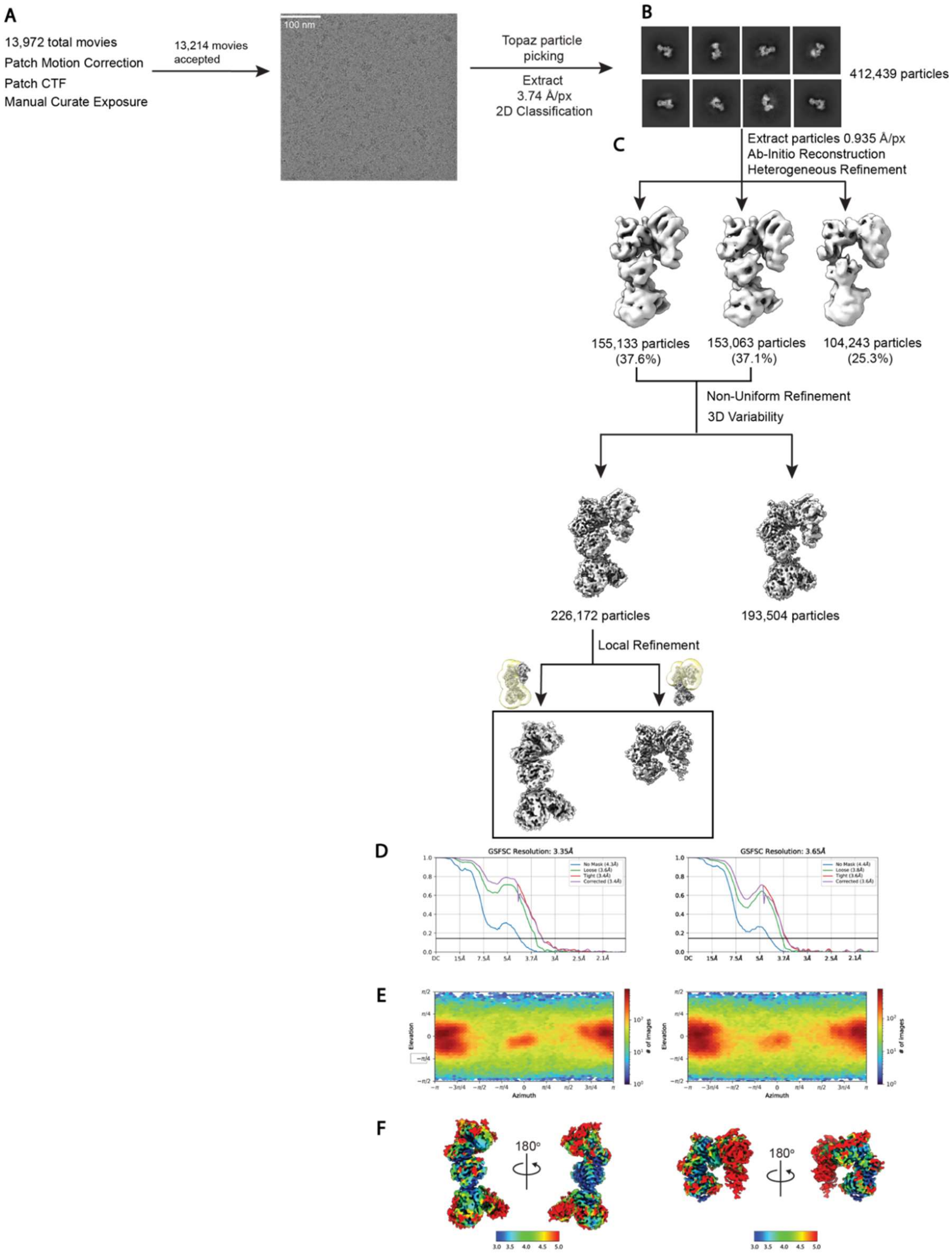


**Fig. S1. Cryo-EM workflow for LRRK2$^{RCKW}$:RN277:E11 DARPin complex dataset. A**. Representative micrograph. **B**. 2D averages for monomer particles dataset. **C**. Data processing strategy for the dataset. **D-E**. FSC plots and Euler angle respectively. **F**. Local resolution maps.

**Table S1. 31 Type-II inhibitors screened against $LRRK2^{KW}$ via DSF assay.**

| Inhibitor | dTm LRRK2(KW) |
|---|---|
| ALW-II-41-27 | -0.5 |
| AMG 900 | 4.5 |
| Apatinib | -1 |
| AST-487 | 10 |
| Axitinib | 7.5 |
| AZ 628 | -0.5 |
| BIRB-796 | 0 |
| BMS 777607 | 6.8 |
| Cabozantinib | 7.5 |
| Foretinib | 11.5 |
| Golvatinib | 0.5 |
| GSK2850163A | 0.5 |
| Imatinib | 0 |
| KI20227 | 3 |
| Linifanib | 5.8 |
| Masitinib | 0 |
| MCP-007 | -0.5 |
| Motesanib | -0.5 |
| Nilotinib | -2.5 |
| Nintedanib | 5 |
| NVP-BHG712 | -0.5 |
| OSI-930 | 0 |
| Pexidartinib | 6.8 |
| PF-4618433 | 11 |
| Quizartinib | 3.5 |
| Rabusertinib | 4 |
| Regorafinib | 5.5 |
| Sorafinib | 7.5 |
| Tivozanib (AV-951) | 11.5 |
| ZM336372 | -1 |
| ZM447439 | 6.5 |

**Table S2. Compound structures, thermal shift data and IC$_{50}$ values of MLi-2 inspired type II hybrid compounds.** The selectivity was calculated as the number of kinases showing a thermal shift greater than 5 K divided by the number of kinases screened.

| Compound | | 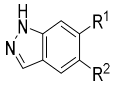 | ΔTm [K] | IC$_{50}$ [nM] | S (5K) |
|---|---|---|---|---|---|
| **47** | $R^1$ = H | $R^2$ = 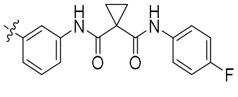 | 3.0 | > 15 µM | n.a. |
| **48** | $R^1$ = H | $R^2$ = 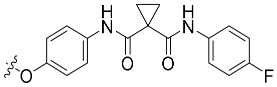 | 1.0 | > 15 µM | n.a. |
| **49** | $R^1$ = H | $R^2$ = 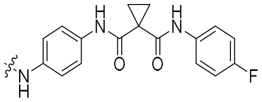 | 2.0 | > 15 µM | n.a. |
| **50** | $R^1$ = H | $R^2$ = 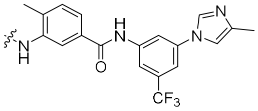 | 1.5 | > 15 µM | n.a. |
| **51** | $R^1$ = 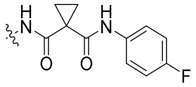 | $R^2$ = H | 1.0 | > 15 µM | n.a. |
| **52** | $R^1$ = 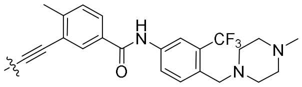 | $R^2$ = H | 10.0 | 897 ± 120 | 0.27 |
| **53** | $R^1$ = 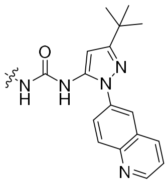 | $R^2$ = H | 2.0 | > 15 µM | n.a. |
| **54** | $R^1$ = 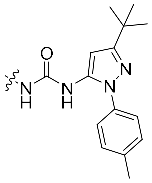 | $R^2$ = H | 2.0 | > 15 µM | n.a. |

**Table S3. Cryo-EM data collection, refinement, and validation statistics.**

| | $LRRK2^{RCKW}$:RN277:E11 DARPin Consensus refinement (EMDB-47006) (PDB 9DMI) | $LRRK2^{RCKW}$:RN277:E11 DARPin Focused refinement COR-B-Kinase-WD40 (EMDB-47025) | $LRRK2^{RCKW}$:RN277:E11 DARPin Focused refinement ROC-COR-A (EMDB-47004) |
|---|---|---|---|
| **Data collection and processing** | | | |
| Magnification | 130000 | 130000 | 130000 |
| Voltage (kV) | 300 | 300 | 300 |
| Electron exposure (e–/Å$^2$) | 55 | 55 | 55 |
| Defocus range (μm) | -1.0 to -3.0 | -1.0 to -3.0 | -1.0 to -3.0 |
| Pixel size (Å) | 0.935 | 0.935 | 0.935 |
| Symmetry imposed | C1 | C1 | C1 |
| Initial particle images (no.) | 412,439 | 412,439 | 412,439 |
| Final particle images (no.) | 226,172 | 226,172 | 226,172 |
| Map resolution (Å) | 3.4 | 3.35 | 3.65 |
| FSC threshold | 0.143 | 0.143 | 0.143 |
| Map resolution range (Å) | 3-5 | 3-5 | 3-5 |
| **Refinement** | | | |
| Initial model used | 6VP7 | -- | -- |
| (PDB code) | | -- | -- |
| Model resolution | -- | -- | -- |
| (Å) | -- | -- | -- |
| FSC threshold | | | |
| Model composition | | | |
| Non-hydrogen | 7954 | -- | -- |
| atoms | 1047 | -- | -- |
| Protein residues | 0 | -- | -- |
| Nucleotide | 1 | -- | -- |
| Ligands | | | |
| *B* factors (Å$^2$) | | | |
| Protein | 103.35 | -- | -- |
| Nucleotide | -- | -- | -- |
| Ligand | 47.77 | -- | -- |
| R.m.s. deviations | | | |
| Bond lengths (Å) | 0.007 | -- | -- |
| Bond angles (°) | 0.938 | -- | -- |
| Validation | | | |
| MolProbity score | 2.24 | -- | -- |

| | | | |
|---|---|---|---|
| Clashscore | 19.70 | -- | -- |
| Ramachandran plot | | | |
| Favored (%) | 92.92 | -- | -- |
| Allowed (%) | 6.98 | -- | -- |
| Disallowed (%) | 0.10 | -- | -- |

**Table S4. Expression constructs used in this study.**

| Construct | Description | Source/Identifier |
|---|---|---|
| **pET-28a(+)-Rab8A** | Rab8A D6 to K176 | RRID: Addgene_228880 |
| **pFastBac1-LRRK2$^{RCKW}$** | LRRK2 R1327 to E2527 | RRID: Addgene_226784 |
| **pFastBac1-LRRK2$^{KW}$** | LRRK2 R1847 to E2527 | RRID: Addgene_228879 |
| **pET17b-K560** | Human Kinesin residues 1-560 with C-terminal GFP | Gift from the Vale lab. RRID:Addgene_15219 |
| **pcDNA5-LRRK2** | LRRK2 full-length, untagged | RRID:Addgene_229019 |
| **EGFP-Rab8a** | | RRID: Addgene_49543 |
| **pQE30- DARPin E11** | N-terminal 8× His tag and a C-terminal 3x-FLAG tag | RRID: Addgene_226784 |
| **pNIC28-Bsa4-CLK3** | N-terminal 6x His tag and TEV cutting site | RRID: Addgene_38831 |

**Table S5. Data collection and Refinement Statistics CLK3 co-crystal structure.**

| **Data collection** | **CLK3-RN129** |
|---|---|
| Beamline | X06SA/PXI SLS |
| Wavelength (Å) | 1.00000 |
| Space group | $P4_1$ |
| Cell dimensions | |
| *a, b, c (Å)* | 74.59, 74.59, 97.04 |
| α, β, γ (°) | 90.00, 90.00, 90.00 |
| Resolution (Å)* | 48.52-2.70 (2.83-2.70) |
| unique observations* | 14673 (1937) |
| $R_{meas}$* | 0.10 (1.47) |
| Completeness (%)* | 100.0 (100.0) |
| Multiplicity* | 14.0 (14.7) |
| mean I/σI* | 17.4 (2.3) |
| CC1/2* | 0.99 (0.76) |
| **Refinement** | |
| $R_{work}$ / $R_{free}$ | 20.4 /26.3 |
| No. of atoms | 2643 |
| Rms deviations | |
| Bond lengths (Å) | 0.006 |
| Bond angles (°) | 1.355 |
| Ramachandran outlier (%) | 0.0 |
| **Protein Data Bank entry** | 9EZ3 |

*Values for the highest resolution shell are shown in parentheses

**Table S6. NanoBRET constructs listed with corresponding tracer and recommended tracer concentration.**

| Target | Tracer (tracerDB ID) | Tracer concentration [nM] | PROMEGA plasmid catalog no. | URL |
|---|---|---|---|---|
| JNK2 | K10 (T000008) | 130 | NV1711 | https://www.promega.com/products/cell-signaling/kinase-target-engagement/nanoluc-mapk9-fusion-vector/?catNum=NV1711 |
| CDKL1 | K10 (T000008) | 250 | NV2881 | https://www.promega.com/products/cell-signaling/kinase-target-engagement/nanoluc-cdkl1-fusion-vector/?catNum=NV2881 |
| STK10 | K10 (T000008) | 500 | NV4261 | https://www.promega.com/products/cell-signaling/kinase-target-engagement/nanoluc-stk10-fusion-vector/?catNum=NV4261 |
| TTK | K9 (T000017) | 660 | NV2191 | https://www.promega.com/products/cell-signaling/kinase-target-engagement/ttk-nanoluc-fusion-vector/?catNum=NV2191 |
| DYRK2 | K10 (T000008) | 1000 | NV3041 | https://www.promega.com/products/cell-signaling/kinase-target-engagement/dyrk2-nanoluc-fusion-vector/?catNum=NV3041 |
| STK17B | K10 (T000008) | 1000 | NV4271 | https://www.promega.com/products/cell-signaling/kinase-target-engagement/stk17b-nanoluc-fusion-vector/?catNum=NV4271 |
| CLK1 | K10 (T000008) | 250 | NV1131 | https://www.promega.com/products/cell-signaling/kinase-target-engagement/nanoluc-clk1-fusion-vector/?catNum=NV1131 |
| SLK | K10 (T000008) | 1000 | NV2051 | https://www.promega.com/products/cell-signaling/kinase-target-engagement/nanoluc-slk-fusion-vector/?catNum=NV2051 |
| MAPK14 | K10 (T000008) | 500 | NV1661 | https://www.promega.com/products/cell-signaling/kinase-target-engagement/mapk14-nanoluc-fusion-vector/?catNum=NV1661 |
| JNK3 | K10 (T000008) | 250 | NV1481 | https://www.promega.com/products/cell-signaling/kinase-target- |

| | | | | |
|---|---|---|---|---|
| | | | | engagement/jnk3-nanoluc-fusion-vector/?catNum=NV1481 |
| DDR2 | K4 (T000037) | 63 | NV1201 | https://www.promega.com/products/cell-signaling/kinase-target-engagement/ddr2-nanoluc-fusion-vector/?catNum=NV1201 |

**Data S1. (separate file)**

The tabular data of all figures and tables can be found in the excel file **Data S1** and at doi.org/10.5281/zenodo.13765594.

**Scheme S1:** Synthesis of PF-360 and GZD-824 inspired hybrid type II inhibitor 1.[a]

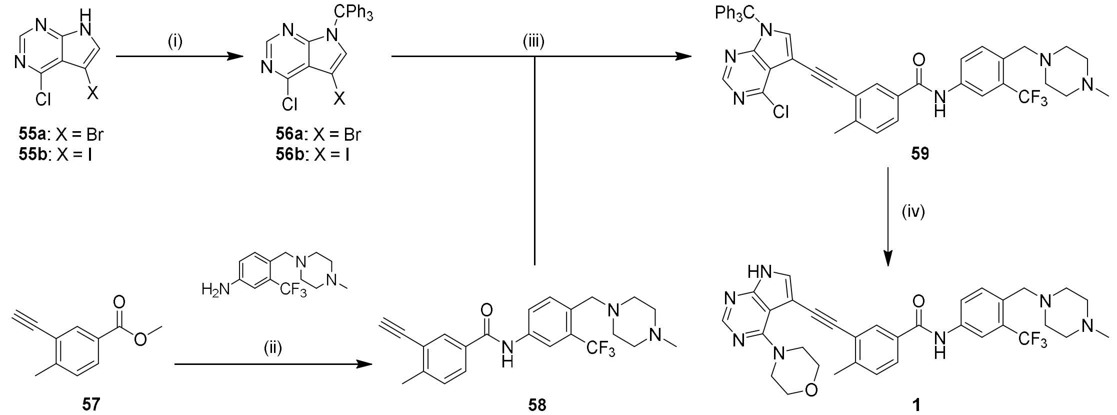


[a]Reagents and conditions: (i) CPh3Cl, NaH, THF, RT, ON; (ii) tBuOK, THF, -20 °C – RT, ON; (iii) [PdCl2(PPh3)2], CuI, DIPEA, DMF, 80 °C, ON; (iv)(a) morpholine, DIPEA, DMF, 120°C, ON (b) TFA, RT, ON.

**Scheme S2:** Synthesis of PF-360 and Foretinib inspired hybrid type II inhibitors 2-8.[a]

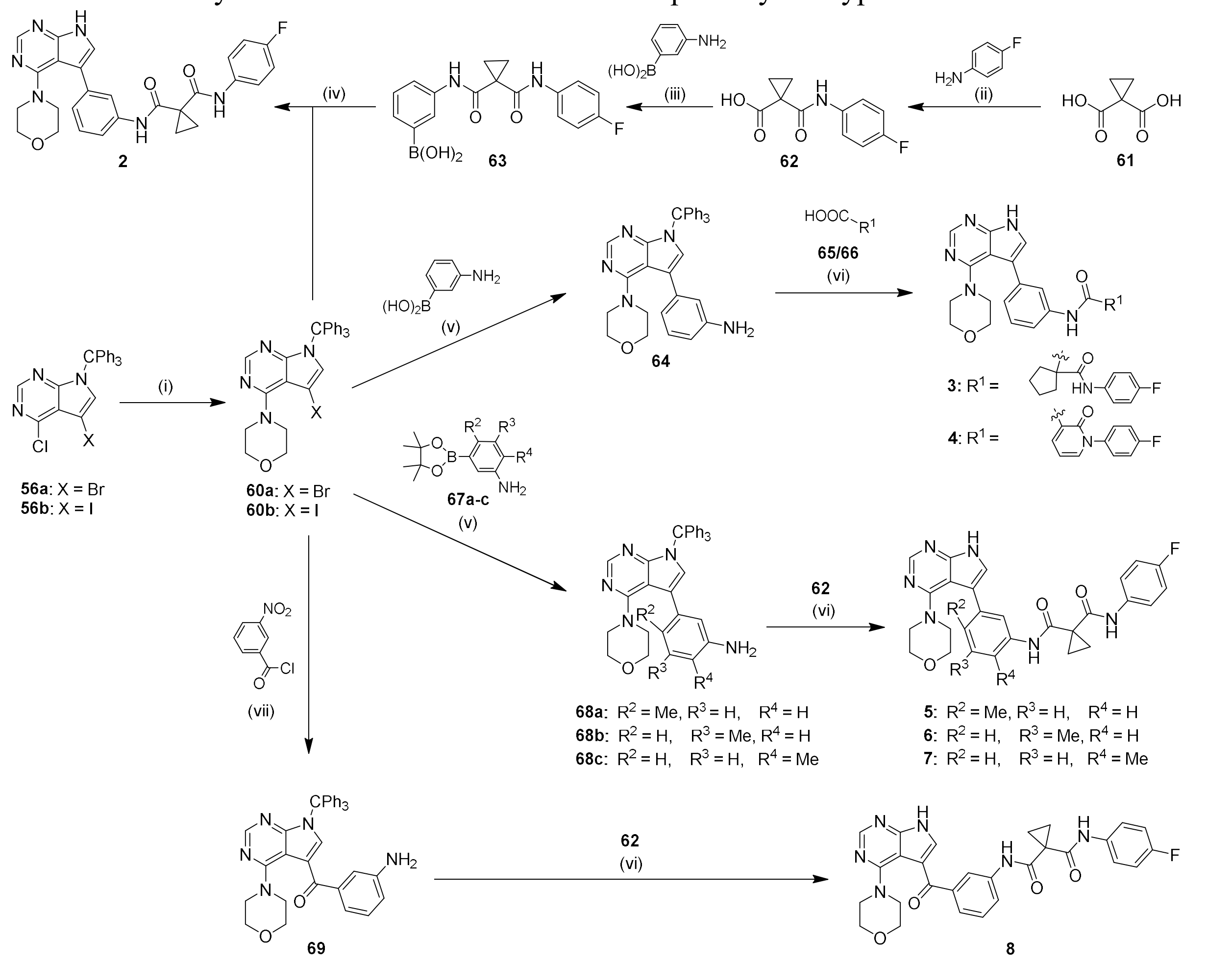

[a]Reagents and conditions: (i) morpholine, K2CO3, DMF, RT, ON; (ii) SOCl2, TEA, THF, 0 °C - RT, ON; (iii) HATU, DIPEA, DMF, RT, ON; (iv)(a) XPhos Pd Gen. 2, XPhos, K3PO4, dioxane/H2O, 80 °C, 4 h (b) TFA, RT, ON; (v) XPhos Pd Gen. 2, XPhos, K3PO4, dioxane/H2O, 80 °C, 4 h; (vi)(a) PyAOP or HATU, DIPEA, DMF, RT, ON (b) TFA, RT, ON; (vii)(a) n-BuLi, THF, -78 °C, 2 h (b) Fe, NH4Cl, MeOH/H2O, 60 °C, ON.

**Scheme S3:** Synthesis of 2 derived hybrid type II inhibitors 9-27.[a]

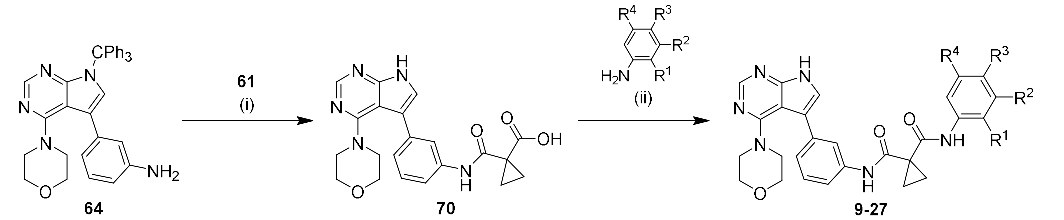


[a]Reagents and conditions: (i)(a) SOCl2, THF, 60 °C, 2 h (b) TFA, RT, ON; (ii) PyAOP, DIPEA, DMF, RT, ON.

**Scheme S4:** Synthesis of Rebastinib inspired hybrid type II inhibitors 28,29 and 31.[a]

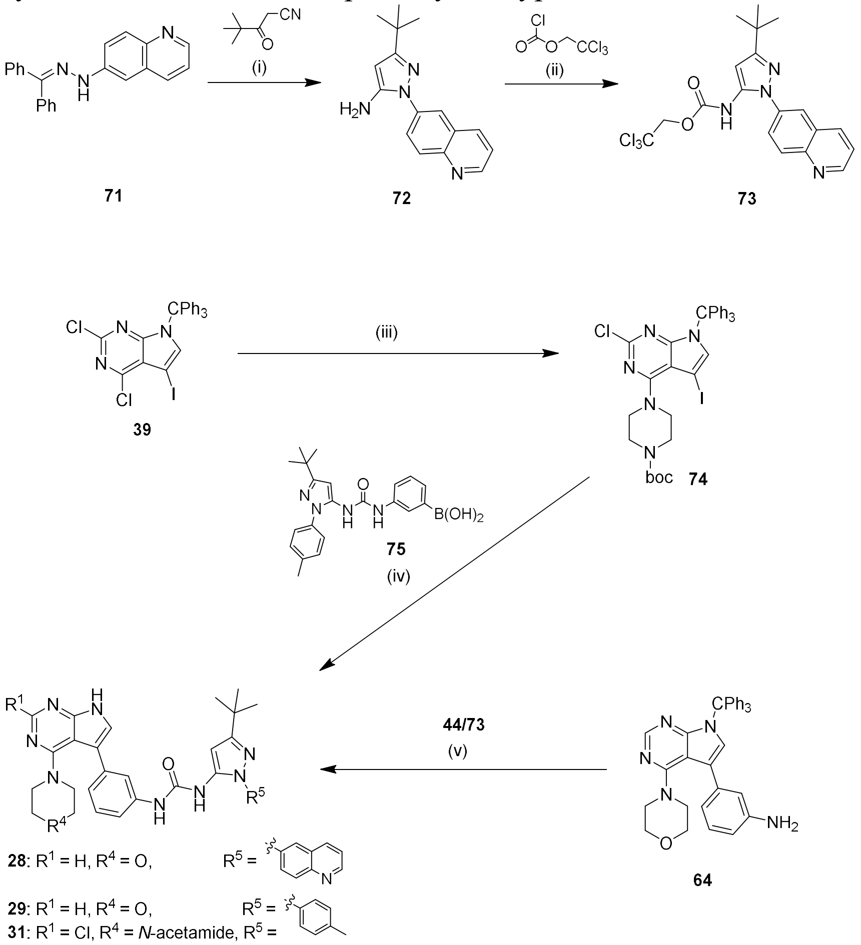


[a]Reagents and conditions: (i) HCl, EtOH/$H_2O$, reflux, ON; (ii) pyridine, DMAP, DCM, -10 °C, 2 h; (iii) morpholine or 1-boc piperazine, $K_2CO_3$, DMF, RT, ON; (iv)(a) XPhos Pd Gen. 2, XPhos, $K_3PO_4$, dioxane/water, 80 °C, 4 h (b) TFA, RT, ON (c) acetyl chloride, TEA, DCM/THF, RT, 3 h; (v)(a) DIPEA, DMF or DMSO, 60 °C, ON (b) TFA, RT, ON.

**Scheme S5:** Synthesis of the type I inhibitors 33 and 34 and the hybrid type II inhibitors 35-37.[a]

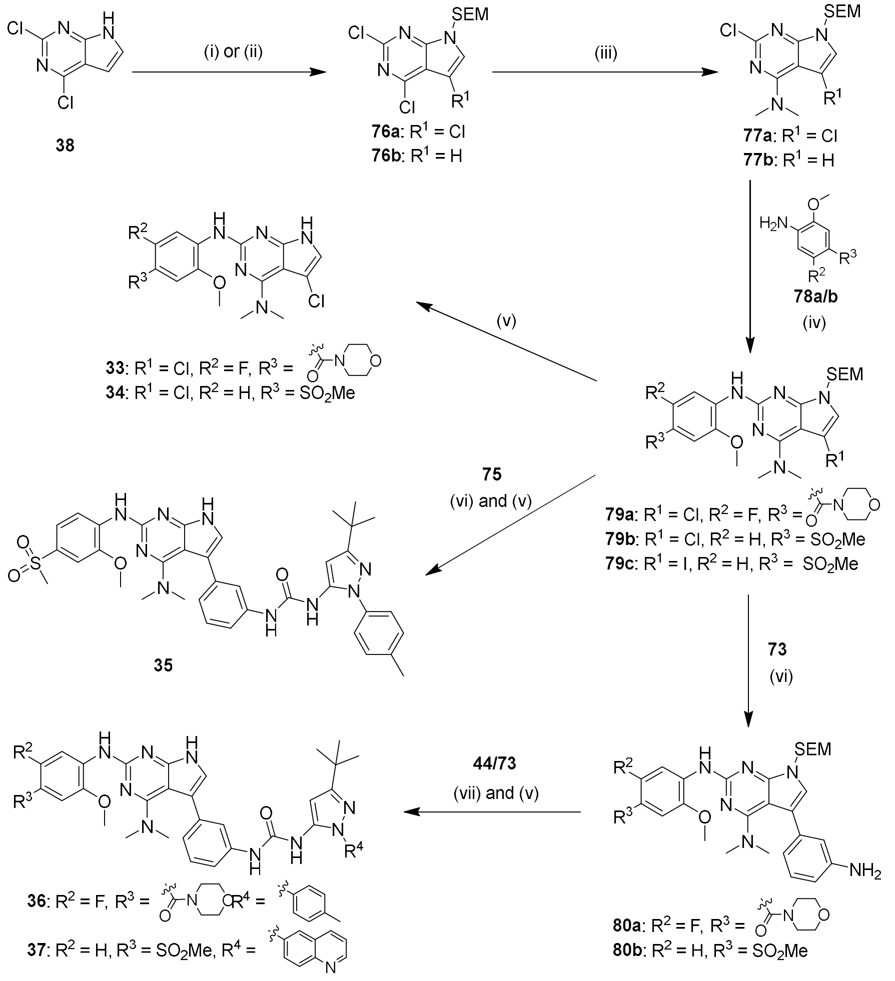


[a]Reagents and conditions: (i)(a) NCS, ACN, 70 °C, ON (b) SEM-Cl and NaH, DMF, 0 °C – RT, 2 h; (ii) SEM-Cl and NaH, DMF, 0 °C – RT, 2 h; (iii) dimethyl amine and DIPEA, EtOH, RT, ON; (iv)[(a) NIS, DCM, RT, 5 h] (b) XPhos Pd Gen. 2, Xphos, K3PO4, dioxane, 60 °C, 1 h; (v)(a) TFA, RT, 1 h (b) aq. NH3 25 %, 60 °C, 2 h; (vi) (a) XPhos Pd Gen. 2, XPhos, K3PO4, dioxane/water, 80 °C, 4 h; (vii) DIPEA, DMSO, 60 -70 °C, ON.

**Scheme S6:** Synthesis of MLi-2 inspired hybrid type II inhibitors 47-54.[a]

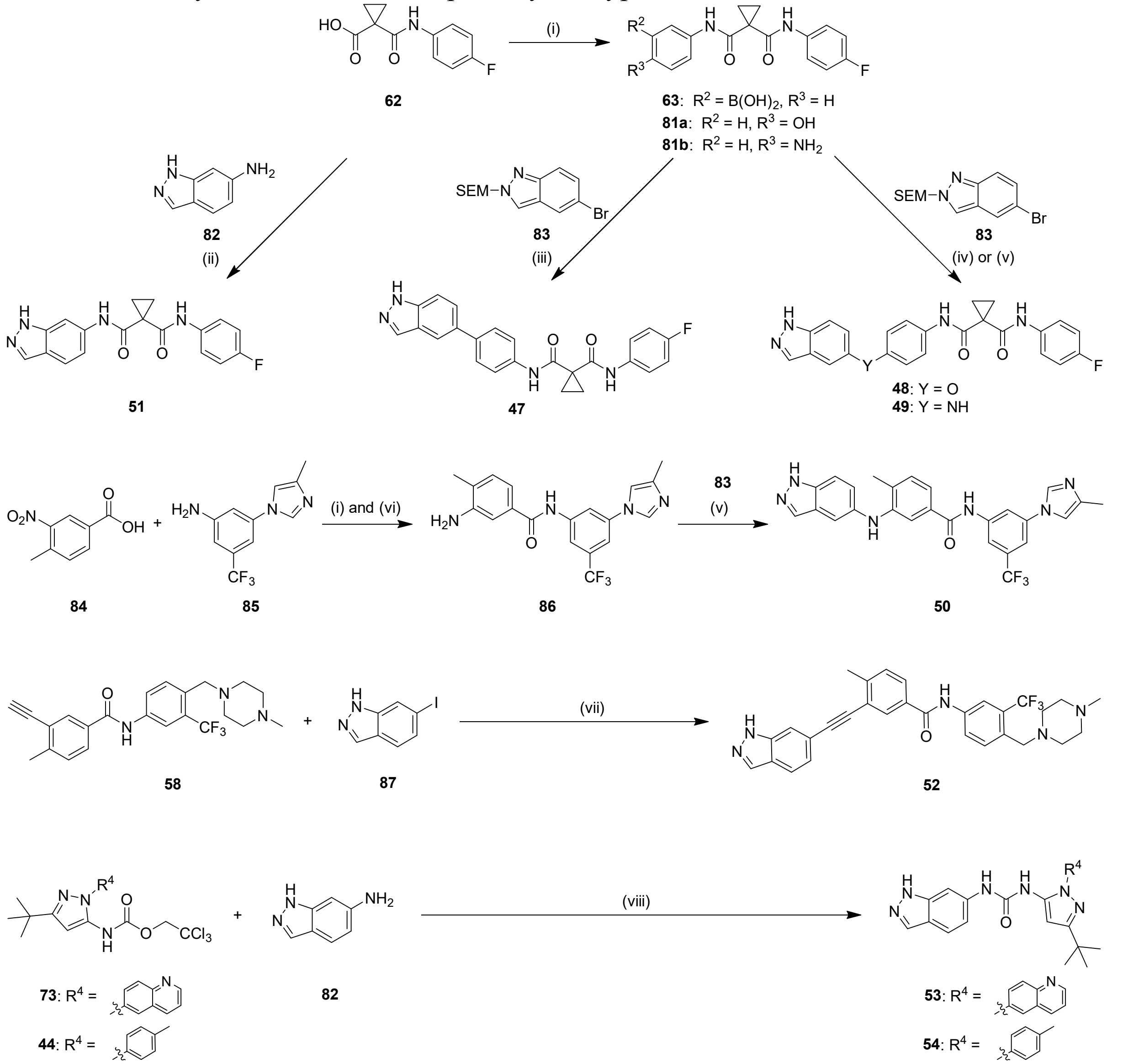


[a]Reagents and Conditions: (i) EDC or HATU, DIPEA, DMF, RT, ON; (ii) T3P, pyridine, ACN, RT, 2 h; (iii) XPhos Pd Gen. 2, XPhos, K3PO4, dioxane/H2O 4:1, 80 °C, 4 h; (iv) tBuBrettPhos Pd Gen.3, tBuBrettPhos, K3PO4, toluene/DME, 100 °C, ON; (v) BrettPhos Pd Gen. 4, BrettPhos, Cs2CO3, toluene/tBuOH, 110 °C, ON; (vi)(a) PyAOP, DIPEA, DMF, RT, ON (b) Pd/C, MeOH, RT, ON; (vii) [Pd(PPh3), CuI, DIPEA, DMF, 80 °C, ON (viii) DIPEA, DMSO, 70 °C, 2-16 h.

### Synthesis Instructions

#### Analytic of small molecules

All synthesized compounds were characterized by mass spectrometry (MS) with electron spray ionization (ESI) and $^{1}H$ NMR. All inhibitors, which were biologically tested, have been additionally characterized by $^{13}C$ NMR, high resolution mass spectrometry (HRMS) and the purity was measured by high performance liquid chromatography (HPLC). The MS spectrograms were recorded with a ThermoFisher Surveyor MSQ or with an Agilent LC/MSD (G6125B). The NMR spectra were measured with a DPX250, an AV400, an AV400HD and an AV500HD from Bruker. The experiments were processed with the deuterated solvents DMSO-$d_6$ or $CDCl_3$ and the resulting spectra were referenced on the solvent signals (DMSO-$d_6$: $^{1}H$-NMR 2.50 ppm, $^{13}C$ NMR 39.52 ppm; $CDCl_3$: $^{1}H$-NMR 7.26 ppm, $^{13}C$ NMR 77.16 ppm). HRMS was measured on a ThermoScientific MALDI LTQ Orbitrap XL or a Bruker micOTOF QII. The HPLC analysis was performed with an Agilent 1260 Infinity II setup containing a flexible pump (G7104C), a multisampler (G7167A), a column compartment (G7116A), a DAD HS detector (G7117C; 254 nm, 280 nm, 310 nm) and the LC/MSD (G6125B, ESI pos. 100-1000). The Poroshell 120 EC-C18 (Agilent, 3 x 150 mm, 2.7 µm) reversed phase column was used with the eluents 0.1 % formic acid in water (A) and 0.1 % formic acid in acetonitrile (B). Two different gradient methods were used with a flowrate of 0.6 mL/min: Method 1: 0 min, 5 % B – 2 min, 80 % B – 5 min, 95 % B – 7 min, 95 % B. Method 2: 0 min, 5 % B – 0.4 min 5 % B – 8 min, 100 % B – 10 min, 100 % B. The UV-detection was performed at 320 nm (150 nm bandwidth).

#### General synthesis and purification methods

The starting materials, reagents and solvents were purchased from common vendors and have been used without further purification. In all procedures with anhydrous solvents the synthesis was carried out under argon atmosphere. The microwave reactions were performed in a CEM Explorer SP48 microwave. Liquid-liquid extractions were always repeated three times, before the organic layers were combined, dried over $MgSO_4$, filtrated and concentrated in vacuo. The flash chromatography was conducted with a puriFlash XS420 device with an UV-VIS detector (200-400 nm) from Interchim. The pre-packed columns from Interchim PF-SIHP for normal phase (NP) flash chromatography and the PF-C18HP for reversed phase (RP) flash chromatography, both with a particle size of 30 µm, were used. The NP flash chromatography was performed either with *n*-hexane and ethyl acetate (EA) and a gradient from 100 % *n*-hexane to 100 % EA or with dichloromethane (DCM) and methanol (MeOH) and a gradient from 100 % DCM to 10 % MeOH. The RP flash chromatography was conducted with acetonitrile (ACN) and water as eluent and a gradient from 95 % water to 100 % acetonitrile. The preparative HPLC was performed with an Agilent 1260 Infinity II setup containing a preparative binary pump (G7161A), an auto sampler (G7157A), a multi wavelength detector (G7165A; 254 nm, 280 nm) and a preparative scale fraction collector (G1364E). The Eclipse XDB-C18 column 120 EC-C18 (Agilent, 21.2 x 250 mm, 7 µm) was used with the eluents 0.1 % trifluoroacetic acid in water (A) and 0.1 % trifluoroacetic acid in acetonitrile (B). A gradient method was used with a flow-rate of 21.0 mL/min: 0 min, 5 % B – 3 min, 5 % B – 28 min, 98 % B – 30 min, 98 % B.

#### General Procedures

When a general procedure was applied, the amount of all reagents was recalculated according to the quantity of starting material used.

**General Procedure A – Suzuki coupling**

The boronic acid or pinacol ester (1 eq.), the aryl halide (1 eq), potassium phosphate (3 eq.), XPhos (0.05 eq.) and XPhos Pd G2 (0.05 eq.) were suspended in dioxane and water (4:1). The reaction was heated to 80 °C and stirred for 4 h.

**General procedure B – aromatic nucleophilic substitution**

The aryl chloride (1 eq.), the amine (1.2 eq.) and potassium carbonate (3 eq.) were solved in anhydrous DMF. The reaction stirred at room temperature overnight. Water was added and the precipitated solid was filtered and washed with water and little amounts of cold ethanol.

**General procedure C – trityl deprotection**

The starting material was solved in TFA and stirred overnight. The reaction mixture was poured in 4 M potassium carbonate solution and extracted with EA.

**General procedure D – amide coupling**

The carboxylic acid (1 eq.), the amine (1 eq.) and PyAOP (1.2 eq.) were solved in anhydrous DMF, before DIPEA (3 eq.) was added. The reaction was stirred at room temperature overnight.

**General procedure E – SEM deprotection**

The starting material was solved in 2 mL TFA and stirred at room temperature for 1 h. The solvent was removed in vacuo, before 10 mL aq. $NH_3$ (25 %) was added. The reaction was heated to 60 °C and stirred for 2 h. Water was added and the reaction mixture was extracted with EA.

**4-methyl-*N*-(4-((4-methylpiperazin-1-yl)methyl)-3-(trifluoromethyl)phenyl)-3-((4-morpholino-7*H*-pyrrolo[2,3-*d*]pyrimidin-5-yl)ethynyl)benzamide (1)**

200 mg **59** (247 µmol) was solved in 20 mL anhydrous DMF. 129 µL DIPEA (741 µmol) and 32 µL morpholine (371 µmol) was added. The reaction was heated to 120 °C and stirred overnight. The solvent was removed and the crude intermediate was purified via flash chromatography (DCM/MeOH). The resulting light brown solid was treated according to **general procedure C**. The crude product was purified via flash chromatography (DCM/MeOH). 67 mg **1** (108 µmol, 43 % yield) was obtained as a yellow solid.

**MS (ESI+)**: m/z = 618.25 $[M+H]^+$

**$^1$H NMR** (500 MHz, DMSO-*d*$_6$): δ = 12.43 (s, 1H), 10.56 (s, 1H), 8.32 (s, 1H), 8.23 (d, *J* = 2.1 Hz, 1H), 8.12 – 8.08 (m, 2H), 7.91 (dd, *J* = 8.0, 1.8 Hz, 1H), 7.87 (s, 1H), 7.71 (d, *J* = 8.6 Hz, 1H), 7.51 (d, *J* = 8.2 Hz, 1H), 3.82 – 3.76 (m, 5H), 3.75 – 3.70 (m, 4H), 3.66 (s, 2H), 3.41 – 3.24 (m, 4H), 3.22 – 2.99 (m, 4H), 2.73 (s, 3H), 2.54 (s, 3H) ppm.

**$^{13}$C NMR** (101 MHz, DMSO-*d*$_6$): δ = 164.81, 158.83, 157.93 (d, TFA), 152.25, 151.22, 143.41, 138.52, 132.08, 131.47, 131.02, 130.32, 129.92, 129.79, 127.54 (t), 125.35, 123.51, 123.17, 122.93, 117.36 (q, TFA), 103.67, 94.36, 89.92, 88.26, 66.02, 56.63, 53.00, 49.75, 48.93, 42.53, 20.46, 13.47 ppm.

**$^{19}$F NMR** (470 MHz, DMSO-*d*$_6$): δ = -57.94 ($CF_3$, 3 F), -73.50 (TFA, 3 F) ppm.

**HRMS**: $[C_{33}H_{34}F_3N_7O_2+H]^+$ calculated: m/z = 618.27988, found: m/z = 618.27967

**HPLC**: $t_R$ = 5.812 min (Method 1), ≥ 95 % purity

**$N$-(4-fluorophenyl)-$N$-(3-(4-morpholino-7$H$-pyrrolo[2,3-$d$]pyrimidin-5-yl)phenyl)cyclopropane-1,1-dicarboxamide (2)**

The **general procedure A** was conducted with the boronic acid **63** (50 mg, 146 µmol) and the aryl halide **60a** (85 mg, 161 µmol). The solvent was removed and the intermediate product was treated according to the **general procedure C**. The obtained crude product was purified via RP flash chromatography. 15 mg of **2** (30.0 µmol, 20 % yield) was obtained as a white solid.

**MS (ESI+):** m/z = 501.20 $[M+H]^+$

**$^1$H NMR** (400 MHz, DMSO-$d_6$): δ = 12.11 (s, 1H), 10.21 (s, 1H), 10.01 (s, 1H), 8.34 (s, 1H), 7.80 (t, $J$ = 1.8 Hz, 1H), 7.66 – 7.60 (m, 2H), 7.60 – 7.55 (m, 1H), 7.43 (d, $J$ = 1.3 Hz, 1H), 7.37 (t, $J$ = 7.9 Hz, 1H), 7.27 – 7.22 (m, 1H), 7.18 – 7.11 (m, 2H), 3.47 – 3.40 (m, 4H), 3.18 (d, $J$ = 4.3 Hz, 4H), 1.54 – 1.46 (m, 4H) ppm.

**$^{13}$C NMR** (101 MHz, DMSO-$d_6$): δ = 168.49, 168.14, 159.69, 153.09, 150.21, 138.84, 135.70, 135.06, 128.67, 123.11, 122.53, 122.46, 122.33, 120.12, 118.11, 115.81, 115.12, 114.90, 102.33, 65.40, 49.32, 31.33, 15.56 ppm.

**HRMS**: $[C_{27}H_{25}FN_6O_3+H]^+$ calculated: m/z = 501.20449, found: m/z = 501.20350

**HPLC**: $t_R$ = 6.533 min (Method 2), ≥ 95 % purity

**$N$-(4-fluorophenyl)-$N$-(3-(4-morpholino-7H-pyrrolo[2,3-$d$]pyrimidin-5-yl)phenyl)cyclopentane-1,1-dicarboxamide (3)**

The **general procedure D** was conducted with the carboxylic acid **65** (77 mg, 307 µmol) and the amine **64** (150 mg, 279 µmol). Water was added, the formed precipitate was filtrated and was treated according to the **general procedure C**. The obtained crude product was purified via flash chromatography (DCM/MeOH). The resulting solid was washed with ACN. 10 mg of **3** (67.8 µmol, 7 % yield) was obtained as a white solid.

**MS (ESI+)**: m/z = 529.20 $[M+H]^+$

**$^1$H NMR** (400 MHz, DMSO-$d_6$): δ = 12.09 (d, $J$ = 1.9 Hz, 1H), 9.53 (s, $J$ = 5.0 Hz, 1H), 9.52 (s, 1H), 8.34 (s, 1H), 7.84 (s, 1H), 7.70 – 7.64 (m, 2H), 7.62 (d, $J$ = 8.9 Hz, 1H), 7.42 (d, $J$ = 2.5 Hz, 1H), 7.34 (t, $J$ = 7.9 Hz, 1H), 7.22 (d, $J$ = 7.7 Hz, 1H), 7.15 – 7.08 (m, 2H), 3.43 – 3.35 (m, 4H), 3.19 – 3.10 (m, 4H), 2.39 – 2.25 (m, 4H), 1.68 – 1.62 (m, 4H) ppm.

**$^{13}$C NMR** (101 MHz, DMSO-$d_6$): δ = 170.55, 170.52, 159.67, 159.34, 156.96, 153.06, 150.18, 139.23, 135.59, 135.48, 128.58, 122.74, 122.34, 122.26, 120.40, 118.05, 115.89, 115.00, 114.78, 102.28, 65.35, 63.78, 49.31, 34.05, 24.48 ppm.

**HRMS**: $[C_{29}H_{29}FN_6O_3+H]^+$ calculated: m/z = 529.23579, found: m/z = 529.23717

**HPLC**: $t_R$ = 6.844 min (Method 2), ≥ 95 % purity

**1-(4-fluorophenyl)-$N$-(3-(4-morpholino-7$H$-pyrrolo[2,3-$d$]pyrimidin-5-yl)phenyl)-2-oxo-1,2-dihydropyridine-3-carboxamide (4)**

The **general procedure D** was conducted with the carboxylic acid **66** (72 mg, 307 µmol) and the amine **64** (150 mg, 279 µmol). Water was added, the formed precipitate was filtrated and was treated according to the **general procedure C**. The obtained crude product was purified via flash chromatography (DCM/MeOH). 91 mg of **4** (178 µmol, 64 % yield) was obtained as a white solid.

**MS (ESI+)**: m/z = 511.15 $[M+H]^+$

**$^1$H NMR** (400 MHz, DMSO-$d_6$): δ = 12.22 (s, 1H), 12.07 (s, 1H), 8.59 (dd, $J$ = 7.3, 2.1 Hz, 1H), 8.37 (s, 1H), 8.11 (dd, $J$ = 6.6, 2.1 Hz, 1H), 7.91 (s, 1H), 7.64 – 7.58 (m, 3H), 7.53 (d, $J$ = 2.5 Hz, 1H), 7.46 – 7.39 (m, 3H), 7.28 (d, $J$ = 7.7 Hz, 1H), 6.72 (t, $J$ = 6.9 Hz, 1H), 3.49 – 3.40 (m, 4H), 3.22 – 3.16 (m, 4H) ppm.

$^{13}$C NMR (101 MHz, DMSO-$d_6$): δ = 163.07, 161.89, 161.21, 160.63, 159.38, 152.72, 149.67, 144.72, 143.92, 138.43, 136.33, 136.30, 136.04, 129.33, 129.24, 123.31, 122.82, 120.52, 119.35, 117.35, 116.17, 115.94, 115.78, 106.99, 102.30, 65.32, 49.44 ppm.
**HRMS**: $[C_{28}H_{23}FN_6O_3 +Na]^+$ calculated: m/z = 533.17079, found: m/z = 533.17312
**HPLC**: $t_R$ = 6.401 min (Method 2), ≥ 95 % purity

***N*-(4-fluorophenyl)-*N*-(4-methyl-3-(4-morpholino-7H-pyrrolo[2,3-*d*]pyrimidin-5-yl)phenyl)cyclopropane-1,1-dicarboxamide (5)**

The **general procedure D** was conducted with the carboxylic acid **62** (41 mg, 185 µmol) and the amine **68a** (85 mg, 154 µmol). Water was added, the formed precipitate was filtrated and was treated according to the **general procedure C**. The obtained crude product was purified via RP flash chromatography. 10 mg of **5** (19 µmol, 12 % yield) was obtained as a white solid.
**MS (ESI+)**: m/z = 515.20 $[M+H]^+$
$^{1}$**H NMR** (400 MHz, DMSO-$d_6$): δ = 12.04 (s, 1H), 10.04 (s, 1H), 10.03 (s, 1H), 8.31 (s, 1H), 7.66 – 7.57 (m, 2H), 7.53 (dd, *J* = 8.2, 2.0 Hz, 1H), 7.49 (d, *J* = 1.8 Hz, 1H), 7.28 (d, *J* = 2.4 Hz, 1H), 7.23 (d, *J* = 8.2 Hz, 1H), 7.13 (t, *J* = 8.9 Hz, 2H), 3.25 (s, 4H), 3.10 (s, 4H), 2.18 (s, 3H), 1.46 (s, 4H) ppm.
$^{13}$**C NMR** (101 MHz, DMSO-$d_6$): δ = 168.37, 168.07, 159.42, 152.45, 150.19, 136.52, 135.43, 135.07, 131.54, 130.00, 122.64, 122.38, 122.32, 122.18, 115.08, 114.90, 114.15, 103.86, 65.49, 49.11, 31.22, 19.51, 15.49 ppm.
**HRMS**: $[C_{28}H_{27}FN_6O_3+H]^+$ calculated: m/z = 515.22014, found: m/z = 515.21943
**HPLC**: $t_R$ = 3.706 min (Method 1), ≥ 95 % purity

***N*-(4-fluorophenyl)-*N*-(3-methyl-5-(4-morpholino-7*H*-pyrrolo[2,3-*d*]pyrimidin-5-yl)phenyl)cyclopropane-1,1-dicarboxamide (6)**

The **general procedure D** was conducted with the carboxylic acid **62** (136 mg, 609 µmol) and the amine **68b** (280 mg, 508 µmol). Water was added, the formed precipitate was filtrated and was treated according to the **general procedure C**. The obtained crude product was purified via RP flash chromatography. 18 mg of **6** (35.0 µmol, 7 % yield) was obtained as a white solid.
**MS (ESI+)**: m/z = 515.20 $[M+H]^+$
$^{1}$**H NMR** (400 MHz, DMSO-$d_6$): δ = 12.09 (d, *J* = 1.9 Hz, 1H), 10.16 (s, 1H), 10.00 (s, 1H), 8.34 (s, 1H), 7.68 – 7.57 (m, 3H), 7.42 (d, *J* = 2.5 Hz, 2H), 7.14 (t, *J* = 8.9 Hz, 2H), 7.08 (s, 1H), 3.49 – 3.41 (m, 4H), 3.18 (d, *J* = 4.1 Hz, 4H), 2.35 (s, 3H), 1.54 – 1.44 (m, 4H) ppm.
$^{13}$**C NMR** (101 MHz, DMSO-$d_6$): δ = 168.64, 168.12, 159.75, 159.54, 157.15, 153.07, 150.18, 138.77, 137.66, 135.50, 135.04, 135.01, 124.06, 122.61, 122.53, 122.28, 118.74, 117.21, 115.92, 115.14, 114.92, 102.39, 65.49, 49.36, 31.21, 21.29, 15.67 ppm.
**HRMS**: $[C_{28}H_{27}FN_6O_3 +H]^+$ calculated: m/z = 515.22014, found: m/z = 515.21960
**HPLC**: $t_R$ = 3.817 min (Method 1), ≥ 95 % purity

***N*-(4-fluorophenyl)-*N*-(2-methyl-5-(4-morpholino-7*H*-pyrrolo[2,3-*d*]pyrimidin-5-yl)phenyl)cyclopropane-1,1-dicarboxamide (7)**

The **general procedure D** was conducted with the carboxylic acid **62** (194 mg, 870 µmol) and the amine **68c** (400 mg, 725 µmol). Water was added, the formed precipitate was filtrated and was treated according to the **general procedure C**. The obtained crude product was purified via RP flash chromatography. 62 mg of **7** (120 µmol, 16 % yield) was obtained as a white solid.
**MS (ESI+)**: m/z = 515.20 $[M+H]^+$

**$^{1}$H NMR** (400 MHz, DMSO-$d_6$): δ = 12.08 (s, 1H), 10.19 (s, 1H), 10.11 (s, 1H), 8.33 (s, 1H), 7.83 (s, 1H), 7.66 – 7.58 (m, 2H), 7.41 (s, 1H), 7.30 – 7.23 (m, 2H), 7.16 (t, *J* = 8.9 Hz, 2H), 3.48 – 3.41 (m, 6H), 3.24 – 3.14 (m, 4H), 2.26 (s, 4H), 1.61 – 1.53 (m, 4H) ppm.
**$^{13}$C NMR** (101 MHz, DMSO-$d_6$): δ = 169.41, 168.31, 159.72, 157.50, 153.04, 150.15, 136.24, 134.71, 134.69, 133.33, 130.28, 128.60, 124.30, 123.96, 122.83, 122.77, 122.12, 115.63, 115.20, 115.03, 102.40, 65.41, 49.31, 29.79, 17.41, 16.47 ppm.
**HRMS**: $[C_{28}H_{27}FN_6O_3+H]^+$ calculated: m/z = 515.22014, found: m/z = 515.21933
**HPLC**: $t_R$ = 3.781 min (Method 1), ≥ 95 % purity

***N*-(4-fluorophenyl)-*N*-(3-(4-morpholino-7*H*-pyrrolo[2,3-*d*]pyrimidine-5-carbonyl)phenyl)cyclopropane-1,1-dicarboxamide (8)**

The **general procedure D** was conducted with the carboxylic acid **11** (35.5 mg, 159 µmol) and the amine **69** (60 mg, 106 µmol). Water was added, the formed precipitate was filtrated and was treated according to the **general procedure C**. The obtained crude product was purified via RP flash chromatography. 12 mg of **8** (22.7 µmol, 21 % yield) was obtained as a white solid.
**MS (ESI+)**: m/z = 529.15 $[M+H]^+$
**$^{1}$H NMR** (400 MHz, DMSO-$d_6$): δ = 12.62 (s, 1H), 10.24 (s, 1H), 10.04 (s, 1H), 8.35 (s, 1H), 8.19 (t, *J* = 1.8 Hz, 1H), 7.90 (ddd, *J* = 8.1, 1.9, 1.0 Hz, 1H), 7.72 (s, 1H), 7.66 – 7.59 (m, 2H), 7.59 – 7.55 (m, 1H), 7.48 (t, *J* = 7.9 Hz, 1H), 7.18 – 7.10 (m, 2H), 3.56 – 3.48 (m, 4H), 3.42 – 3.37 (m, 4H), 1.45 (s, 4H) ppm.
**$^{13}$C NMR** (101 MHz, DMSO-$d_6$): δ = 188.36, 168.34, 167.95, 158.77, 154.05, 151.64, 139.00, 138.48, 135.23, 135.21, 132.34, 128.69, 124.60, 124.19, 122.32, 122.24, 121.55, 116.10, 115.12, 114.90, 101.49, 65.77, 48.09, 31.70, 15.33 ppm.
**HRMS**: $[C_{28}H_{25}FN_6O_4+H]^+$ calculated: m/z = 529.19941, found: m/z = 529.19875
**HPLC**: $t_R$ = 6.579 min (Method 1), ≥ 95 % purity

***N*-(1-methyl-1*H*-pyrazol-4-yl)-*N*-(3-(4-morpholino-7*H*-pyrrolo[2,3-*d*]pyrimidin-5-yl)phenyl)cyclopropane-1,1-dicarboxamide (9)**

**9** was synthesized according to the **general procedure D** from 46 mg **70** (113 µmol) and 13 mg 1-methyl-1*H*-pyrazol-4-amine (135 µmol). The solvent was removed and the crude product was purified via RP flash chromatography. 41 mg of **9** (84 µmol, 74 % yield) was obtained as a white solid.
**MS (ESI+)**: m/z = 487.15 $[M+H]^+$
**$^{1}$H NMR** (400 MHz, DMSO-$d_6$): δ = 12.12 (d, *J* = 1.7 Hz, 1H), 10.55 (s, 1H), 9.88 (s, 1H), 8.35 (s, 1H), 7.89 (s, 1H), 7.79 (s, 1H), 7.58 (dd, *J* = 8.1, 0.9 Hz, 1H), 7.49 (s, 1H), 7.45 (d, *J* = 2.5 Hz, 1H), 7.37 (t, *J* = 7.9 Hz, 1H), 7.25 (d, *J* = 7.7 Hz, 1H), 3.78 (s, 3H), 3.48 – 3.41 (m, 4H), 3.21 – 3.14 (m, 6H), 1.54 – 1.38 (m, 4H) ppm.
**$^{13}$C NMR** (101 MHz, DMSO-$d_6$): δ = 167.97, 167.57, 159.74, 153.11, 150.22, 138.80, 135.76, 130.28, 128.77, 123.03, 122.41, 121.83, 121.31, 119.82, 117.78, 115.79, 102.37, 65.42, 49.37, 48.61, 38.65, 30.24, 15.89 ppm.
**HRMS**: $[C_{25}H_{26}N_8O_3+H]^+$ calculated: m/z = 487.22006, found: m/z = 487.21978
**HPLC**: $t_R$ = 3.167 min (Method 1), ≥ 95 % purity

***N*-(2-methyl-1,2,3,4-tetrahydroisoquinolin-6-yl)-*N*-(3-(4-morpholino-7*H*-pyrrolo[2,3-*d*]pyrimidin-5-yl)phenyl)cyclopropane-1,1-dicarboxamide (10)**

**10** was synthesized according to the **general procedure D** from 100 mg **70** (245 µmol) and 48 mg 2-methyl-1,2,3,4-tetrahydroisoquinolin-6-amine (295 µmol). The solvent was removed, and the residue was purified via flash chromatography (DCM/MeOH). 8 mg of **10** (14.5 µmol, 6 % yield) was obtained as a white solid.
**MS (ESI+)**: m/z = 552.25 $[M+H]^+$
**$^1$H NMR** (400 MHz, DMSO-*d*$_6$): δ = 12.12 (s, 1H), 10.21 (s, 1H), 9.90 (s, 1H), 8.35 (s, 1H), 7.80 (s, 1H), 7.57 (d, *J* = 8.9 Hz, 1H), 7.45 (d, *J* = 2.4 Hz, 1H), 7.41 – 7.31 (m, 3H), 7.25 (d, *J* = 7.8 Hz, 1H), 6.97 (d, *J* = 8.3 Hz, 1H), 3.44 (d, *J* = 5.8 Hz, 6H), 3.19 (d, *J* = 4.3 Hz, 4H), 2.78 (t, *J* = 5.7 Hz, 2H), 2.59 (t, *J* = 5.8 Hz, 2H), 2.34 (s, 3H), 1.55 – 1.45 (m, 4H) ppm.
**$^{13}$C NMR** (101 MHz, DMSO-*d*$_6$): δ = 168.42, 159.69, 153.09, 150.20, 138.75, 136.52, 135.70, 133.70, 130.12, 128.67, 126.19, 123.14, 122.35, 120.36, 120.18, 118.23, 118.16, 115.78, 102.33, 65.41, 56.95, 52.24, 49.33, 45.60, 31.07, 28.74, 15.71 ppm.
**HRMS**: $[C_{31}H_{33}N_7O_3+H]^+$ calculated: m/z = 552.27176, found: m/z = 552.27080
**HPLC**: $t_R$ = 3.028 min (Method 1), ≥ 95 % purity

***N*-(2,3-dihydro-1*H*-inden-4-yl)-*N*-(3-(4-morpholino-7*H*-pyrrolo[2,3-*d*]pyrimidin-5-yl)phenyl)cyclopropane-1,1-dicarboxamide (11)**

**11** was synthesized according to the **general procedure D** from 60 mg **70** (147 µmol) and 35.5 mg 2,3-dihydro-1*H*-inden-4-amine (177 µmol). The solvent was removed, and the residue was purified via RP flash chromatography. 55 mg of **11** (105 µmol, 71 % yield) was obtained as a white solid.
**MS (ESI+)**: m/z = 523.15 $[M+H]^+$
**$^1$H NMR** (400 MHz, DMSO-*d*$_6$): δ = 12.13 (d, *J* = 2.1 Hz, 1H), 10.19 (s, 1H), 10.10 (s, 1H), 8.35 (s, 1H), 7.77 (s, 1H), 7.54 (d, *J* = 8.4 Hz, 1H), 7.48 (d, *J* = 7.9 Hz, 1H), 7.46 (d, *J* = 2.5 Hz, 1H), 7.40 (t, *J* = 7.9 Hz, 1H), 7.28 (d, *J* = 7.7 Hz, 1H), 7.10 (t, *J* = 7.7 Hz, 1H), 7.02 (d, *J* = 7.3 Hz, 1H), 3.47 – 3.40 (m, 4H), 3.22 – 3.15 (m, 4H), 2.87 (t, *J* = 7.4 Hz, 2H), 2.80 (t, *J* = 7.4 Hz, 2H), 1.98 (p, *J* = 7.3 Hz, 2H), 1.59 (s, 4H) ppm.
**$^{13}$C NMR** (101 MHz, DMSO-*d*$_6$): δ = 169.52, 168.29, 159.70, 153.10, 150.22, 144.66, 138.32, 135.98, 135.76, 133.96, 128.75, 126.55, 123.50, 122.42, 120.66, 120.13, 118.66, 115.70, 102.36, 65.43, 49.34, 32.68, 30.04, 29.68, 24.39, 16.64 ppm.
**HRMS**: $[C_{30}H_{30}N_6O_3+H]^+$ calculated: m/z = 523.24522, found: m/z = 523.24424
**HPLC**: $t_R$ = 4.040 min (Method 1), ≥ 95 % purity

***N*-(3-(4-morpholino-7*H*-pyrrolo[2,3-*d*]pyrimidin-5-yl)phenyl)-*N*-phenylcyclopropane-1,1-dicarboxamide (12)**

**12** was synthesized according to the **general procedure D** from 100 mg **70** (245 µmol) and 25 mg aniline (270 µmol). The solvent was removed and the residue was purified via RP flash chromatography. The resulting pale yellow solid was washed with 5 mL ACN 27 mg of **12** (56 µmol, 23 % yield) was obtained as a white solid.
**MS (ESI+)**: m/z = 483.20 $[M+H]^+$
**$^1$H NMR** (400 MHz, DMSO-*d*$_6$): δ = 12.11 (d, *J* = 2.1 Hz, 1H), 10.17 (s, 1H), 10.00 (s, 1H), 8.34 (s, 1H), 7.80 (t, *J* = 1.6 Hz, 1H), 7.61 (d, *J* = 7.6 Hz, 2H), 7.57 (d, *J* = 8.1 Hz, 1H), 7.44 (d, *J* = 2.5 Hz, 1H), 7.37 (t, *J* = 7.9 Hz, 1H), 7.30 (t, *J* = 7.9 Hz, 2H), 7.25 (d, *J* = 7.7 Hz, 1H), 7.07 (t, *J* = 7.4 Hz, 1H), 3.49 – 3.38 (m, 4H), 3.22 – 3.11 (m, 4H), 1.54 – 1.46 (m, 4H) ppm.

$^{13}$**C NMR** (101 MHz, DMSO-*d*$_6$): δ = 168.46, 168.35, 159.69, 153.08, 150.20, 138.79, 138.67, 135.69, 128.66, 128.47, 123.66, 123.16, 122.34, 120.56, 120.19, 118.19, 115.80, 102.33, 65.40, 49.32, 31.32, 15.63 ppm.
**HRMS**: $[C_{27}H_{26}N_6O_3+H]^+$ calculated: m/z = 483.21392, found: m/z = 483.21370
**HPLC**: $t_R$ = 3.675 min (Method 1), ≥ 95 % purity

***N*-(4-chlorophenyl)-*N*-(3-(4-morpholino-7H-pyrrolo[2,3-*d*]pyrimidin-5-yl)phenyl)cyclopropane-1,1-dicarboxamide (13)**

**13** was synthesized according to the **general procedure D** from 100 mg **70** (245 µmol) and 34.4 mg 4-chloroaniline (270 µmol). The solvent was removed and the residue was purified via RP flash chromatography. The resulting pale yellow solid was washed with 5 mL ACN. 21 mg of **13** (40.6 µmol, 17 % yield) was obtained as a white solid.
**MS (ESI+)**: m/z = 517.20 $[M+H]^+$
$^1$**H NMR** (400 MHz, DMSO-*d*$_6$): δ = 12.11 (d, *J* = 1.8 Hz, 1H), 10.13 (s, 1H), 10.12 (s, 1H), 8.34 (s, 1H), 7.79 (s, 1H), 7.69 – 7.63 (m, 2H), 7.58 (d, *J* = 8.8 Hz, 1H), 7.43 (d, *J* = 2.5 Hz, 1H), 7.40 – 7.33 (m, 3H), 7.24 (d, *J* = 7.7 Hz, 1H), 3.46 – 3.38 (m, 4H), 3.22 – 3.12 (m, 4H), 1.53 – 1.44 (m, 4H) ppm.
$^{13}$**C NMR** (101 MHz, DMSO-*d*$_6$): δ = 168.95, 168.55, 160.14, 153.55, 150.67, 139.32, 138.24, 136.15, 129.12, 128.81, 127.65, 123.58, 122.78, 122.48, 120.64, 118.63, 116.27, 102.78, 65.87, 49.78, 32.09, 15.99 ppm.
**HRMS**: $[C_{27}H_{25}ClN_6O_3+H]^+$ calculated: m/z = 517.17494, found: m/z = 517.17472
**HPLC**: $t_R$ = 3.923 min (Method 1), ≥ 95 % purity

***N*-(3-(4-morpholino-7*H*-pyrrolo[2,3-*d*]pyrimidin-5-yl)phenyl)-*N*-(p-tolyl)cyclopropane-1,1-dicarboxamide (14)**

**14** was synthesized according to the **general procedure D** from 100 mg **70** (245 µmol) and 31.6 mg p-toluidine (295 µmol). Water was added, the formed precipitate was filtrated and washed with 5 mL DCM. 60 mg of **14** (121 µmol, 49 % yield) was obtained as a white solid.
**MS (ESI+)**: m/z = 497.30 $[M+H]^+$
$^1$**H NMR** (400 MHz, DMSO-*d*$_6$): δ = 10.23 (s, 1H), 9.90 (s, 1H), 8.35 (s, 1H), 7.80 (s, 1H), 7.56 (d, *J* = 8.0 Hz, 1H), 7.49 (d, *J* = 8.4 Hz, 2H), 7.44 (d, *J* = 2.4 Hz, 1H), 7.37 (t, *J* = 7.9 Hz, 1H), 7.25 (d, *J* = 7.6 Hz, 1H), 7.10 (d, *J* = 8.3 Hz, 2H), 3.48 – 3.39 (m, 4H), 3.22 – 3.12 (m, 4H), 2.25 (s, 3H), 1.56 – 1.44 (m, 4H) ppm.
$^{13}$**C NMR** (101 MHz, DMSO-*d*$_6$): δ = 168.41, 159.70, 153.09, 150.21, 138.77, 136.08, 135.71, 132.67, 128.87, 128.68, 123.16, 122.36, 120.66, 120.15, 118.15, 115.80, 102.35, 65.41, 49.34, 31.10, 20.46, 15.69 ppm.
**HRMS**: $[C_{28}H_{28}N_6O_3+H]^+$ calculated: m/z = 497.22957, found: m/z = 497.22931
**HPLC**: $t_R$ = 3.803 min (Method 1), ≥ 95 % purity

***N*-(4-ethylphenyl)-*N*-(3-(4-morpholino-7*H*-pyrrolo[2,3-*d*]pyrimidin-5-yl)phenyl)cyclopropane-1,1-dicarboxamide (15)**

**15** was synthesized according to the **general procedure D** from 100 mg **70** (245 µmol) and 31.6 mg 4-ethylaniline (295 µmol). Water was added, the formed precipitate was filtrated and washed with 5 mL ACN. 99 mg of **15** (194 µmol, 79 % yield) was obtained as a white solid.
**MS (ESI+)**: m/z = 511.25 $[M+H]^+$

$^{1}$**H NMR** (400 MHz, DMSO-*d$_6$*): δ = 12.18 (s, 1H), 10.30 (s, 1H), 10.01 (s, 1H), 8.34 (s, 1H), 7.82 (s, 1H), 7.58 (d, *J* = 8.0 Hz, 1H), 7.52 (d, *J* = 8.4 Hz, 2H), 7.43 (d, *J* = 2.4 Hz, 1H), 7.36 (t, *J* = 7.9 Hz, 1H), 7.24 (d, *J* = 7.6 Hz, 1H), 7.13 (d, *J* = 8.4 Hz, 2H), 3.43 (s, 4H), 3.17 (s, 4H), 2.59 – 2.53 (m, 2H), 1.58 – 1.42 (m, 4H), 1.14 (t, *J* = 7.6 Hz, 3H) ppm.
$^{13}$**C NMR** (101 MHz, DMSO-*d$_6$*): δ = 168.46, 159.70, 153.05, 150.18, 139.11, 138.80, 136.32, 135.69, 128.65, 127.65, 123.15, 122.38, 120.72, 120.12, 118.14, 115.77, 102.35, 65.41, 49.34, 31.19, 27.61, 15.73, 15.68 ppm.
**HRMS**: $[C_{29}H_{30}N_6O_3+H]^+$ calculated: m/z = 511.24522, found: m/z = 511.24492
**HPLC**: $t_R$ = 3.994 min (Method 1), ≥ 95 % purity

##### *N*-(4-methoxyphenyl)-*N*-(3-(4-morpholino-7*H*-pyrrolo[2,3-*d*]pyrimidin-5-yl)phenyl)cyclopropane-1,1-dicarboxamide (16)

**16** was synthesized according to the **general procedure D** from 100 mg **70** (245 µmol) and 36 mg 4-methoxyaniline (295 µmol). Water was added, the formed precipitate was filtrated and washed with 5 mL ACN. 92 mg of **16** (179 µmol, 73 % yield) was obtained as a white solid.
**MS (ESI+)**: m/z = 513.25 $[M+H]^+$
$^{1}$**H NMR** (400 MHz, DMSO-*d$_6$*): δ = 12.19 (d, *J* = 1.9 Hz, 1H), 10.41 (s, 1H), 9.92 (s, 1H), 8.34 (s, 1H), 7.82 (s, 1H), 7.58 (d, *J* = 8.4 Hz, 1H), 7.52 (d, *J* = 9.0 Hz, 2H), 7.43 (d, *J* = 2.5 Hz, 1H), 7.36 (t, *J* = 7.9 Hz, 1H), 7.24 (d, *J* = 7.8 Hz, 1H), 6.87 (d, *J* = 9.1 Hz, 2H), 3.71 (s, 3H), 3.47 – 3.40 (m, 4H), 3.20 – 3.13 (m, 4H), 1.58 – 1.42 (m, 4H) ppm.
$^{13}$**C NMR** (101 MHz, DMSO-*d$_6$*): δ = 168.46, 168.38, 159.70, 155.59, 153.04, 150.17, 138.82, 135.69, 131.63, 128.67, 123.08, 122.38, 120.01, 118.02, 115.75, 113.56, 102.35, 65.40, 55.17, 49.34, 31.01, 15.68 ppm.
**HRMS**: $[C_{28}H_{28}N_6O_4+H]^+$ calculated: m/z = 513.22448, found: m/z = 513.22418
**HPLC**: $t_R$ = 3.630 min (Method 1), ≥ 95 % purity

##### *N*-(3,4-difluorophenyl)-*N*-(3-(4-morpholino-7*H*-pyrrolo[2,3-*d*]pyrimidin-5-yl)phenyl)cyclopropane-1,1-dicarboxamide (17)

**17** was synthesized according to the **general procedure D** from 80 mg **70** (196 µmol) and 30.4 mg 3,4-difluoroaniline (236 µmol). Water was added, the formed precipitate was filtrated and washed with 5 mL ACN. 92 mg of **17** (177 µmol, 90 % yield) was obtained as a white solid.
**MS (ESI+)**: m/z = 519.15 $[M+H]^+$
$^{1}$**H NMR** (400 MHz, DMSO-*d$_6$*): δ = 12.11 (s, 1H), 10.20 (s, 1H), 10.12 (s, 1H), 8.34 (s, 1H), 7.87 – 7.74 (m, 2H), 7.59 (d, *J* = 8.4 Hz, 1H), 7.43 (d, *J* = 2.4 Hz, 1H), 7.40 – 7.32 (m, 3H), 7.24 (d, *J* = 7.6 Hz, 1H), 3.48 – 3.37 (m, 4H), 3.25 – 3.11 (m, 4H), 1.48 (s, 4H) ppm.
$^{13}$**C NMR** (101 MHz, DMSO-*d$_6$*): δ = 168.53, 167.89, 159.67, 153.08, 150.20, 138.89, 135.68, 128.65, 123.10, 122.31, 120.17, 118.15, 117.18, 117.00, 116.71, 116.68, 116.66, 116.62, 115.80, 109.57, 109.36, 102.30, 65.39, 49.31, 31.74, 15.43 ppm.
**HRMS**: $[C_{27}H_{24}F_2N_6O_3+H]^+$ calculated: m/z = 519.19507, found: m/z = 519.19475
**HPLC**: $t_R$ = 3.837 min (Method 1), ≥ 95 % purity

##### *N*-(3-chloro-4-fluorophenyl)-*N*-(3-(4-morpholino-7*H*-pyrrolo[2,3-*d*]pyrimidin-5-yl)phenyl)cyclopropane-1,1-dicarboxamide (18)

**18** was synthesized according to the **general procedure D** from 80 mg **70** (196 µmol) and 34.3 mg 3-chloro-4-fluoroaniline (236 µmol). Water was added, the formed precipitate was filtrated and washed with 5 mL DCM. 73 mg of **18** (136 µmol, 69 % yield) was obtained as a pale yellow solid.

**MS (ESI+)**: m/z = 535.15 $[M+H]^+$

**$^{1}$H NMR** (400 MHz, DMSO-*$d_6$*): δ = 12.11 (s, 1H), 10.16 (s, 1H), 10.14 (s, 1H), 8.34 (s, 1H), 7.96 (dd, *J* = 6.9, 2.5 Hz, 1H), 7.78 (s, 1H), 7.62 – 7.52 (m, 2H), 7.43 (d, *J* = 2.5 Hz, 1H), 7.40 – 7.33 (m, 2H), 7.24 (d, *J* = 7.7 Hz, 1H), 3.49 – 3.37 (m, 4H), 3.23 – 3.12 (m, 4H), 1.53 – 1.43 (m, 4H) ppm.

**$^{13}$C NMR** (101 MHz, DMSO-*$d_6$*): δ = 168.58, 167.83, 159.66, 154.49, 153.08, 152.08, 150.19, 138.90, 136.09, 136.06, 135.68, 128.65, 123.08, 122.32, 122.04, 120.77, 120.70, 120.15, 118.89, 118.71, 118.13, 116.70, 116.48, 115.81, 102.30, 65.39, 49.31, 31.71, 15.45 ppm.

**HRMS**: $[C_{27}H_{24}ClFN_6O_3+H]^+$ calculated: m/z = 535.16552, found: m/z = 535.16501

**HPLC**: $t_R$ = 3.963 min (Method 1), ≥ 95 % purity

##### *N*-(3-bromo-4-fluorophenyl)-*N*-(3-(4-morpholino-7*H*-pyrrolo[2,3-*d*]pyrimidin-5-yl)phenyl)cyclopropane-1,1-dicarboxamide (19)

**19** was synthesized according to the **general procedure D** from 40 mg **70** (98 µmol) and 22.5 mg 3-bromo-4-fluorobenzenaminium chloride (118 µmol). The solvent was removed and the residue was purified via preparative HPLC. The resulting TFA salt was solved in EA and washed with 1M aq. $NaHCO_3$ solution. 20 mg of **19** (34.5 µmol, 35 % yield) was obtained as a white solid.

**MS (ESI+)**: m/z = 579.10 $[M+H]^+$

**$^{1}$H NMR** (400 MHz, DMSO-*$d_6$*): δ = 12.39 (s, 1H), 10.16 (s, 1H), 10.14 (s, 1H), 8.40 (s, 1H), 7.78 (s, 1H), 7.59 (d, *J* = 8.0 Hz, 2H), 7.49 (d, *J* = 1.9 Hz, 1H), 7.38 (t, *J* = 7.9 Hz, 1H), 7.33 (t, *J* = 8.8 Hz, 1H), 7.23 (d, *J* = 7.6 Hz, 1H), 3.49 – 3.38 (m, *J* = 4.1 Hz, 4H), 3.30 – 3.19 (m, 4H), 1.48 (d, *J* = 4.7 Hz, 4H) ppm.

**$^{13}$C NMR** (101 MHz, DMSO-*$d_6$*): δ = 168.58, 167.88, 158.41, 158.06, 155.57, 153.17, 151.52, 148.28, 138.99, 136.32, 136.29, 135.29, 128.82, 124.86, 123.12, 122.87, 121.46, 121.39, 120.23, 118.40, 116.55, 116.49, 116.26, 107.34, 107.12, 101.86, 65.34, 49.44, 31.71, 15.47 ppm.

**HRMS**: $[C_{27}H_{24}BrFN_6O_3+H]^+$ calculated: m/z = 579.11501, found: m/z = 579.11507

**HPLC**: $t_R$ = 3.966 min (Method 1), ≥ 95 % purity

##### *N*-(4-fluoro-3-methylphenyl)-*N*-(3-(4-morpholino-7*H*-pyrrolo[2,3-*d*]pyrimidin-5-yl)phenyl)cyclopropane-1,1-dicarboxamide (20)

**20** was synthesized according to the **general procedure D** from 80 mg **70** (196 µmol) and 29.5 mg 4-fluoro-3-methylaniline (236 µmol). Water was added, the formed precipitate was filtrated and washed with 5 mL DCM. 81 mg of **20** (157 µmol, 80 % yield) was obtained as a white solid.

**MS (ESI+)**: m/z = 515.15 $[M+H]^+$

**$^{1}$H NMR** (400 MHz, DMSO-*$d_6$*): δ = 12.11 (s, 1H), 10.21 (s, 1H), 9.95 (s, 1H), 8.35 (s, 1H), 7.79 (s, 1H), 7.57 (d, *J* = 8.5 Hz, 1H), 7.52 (dd, *J* = 7.0, 2.2 Hz, 1H), 7.47 – 7.41 (m, 2H), 7.37 (t, *J* = 7.9 Hz, 1H), 7.25 (d, *J* = 7.6 Hz, 1H), 7.07 (t, *J* = 9.2 Hz, 1H), 3.48 – 3.39 (m, 4H), 3.22 – 3.12 (m, 4H), 2.20 (d, *J* = 1.1 Hz, 3H), 1.55 – 1.44 (m, 4H) ppm.

**$^{13}$C NMR** (101 MHz, DMSO-*$d_6$*): δ = 168.47, 168.25, 159.68, 158.10, 155.72, 153.09, 150.20, 138.81, 135.70, 134.66, 128.68, 124.01, 123.83, 123.77, 123.72, 123.12, 122.35, 120.16, 119.91, 119.83, 118.14, 115.81, 114.76, 114.53, 102.33, 65.41, 49.33, 31.17, 15.65, 14.31, 14.28 ppm.

**HRMS**: $[C_{28}H_{27}FN_6O_3+H]^+$ calculated: m/z = 515.22014, found: m/z = 515.21878

**HPLC**: $t_R$ = 3.859 min (Method 1), ≥ 95 % purity

**_N_-(4-fluoro-3-methoxyphenyl)-_N_-(3-(4-morpholino-7_H_-pyrrolo[2,3-_d_]pyrimidin-5-yl)phenyl)cyclopropane-1,1-dicarboxamide (21)**

**21** was synthesized according to the **general procedure D** from 80 mg **70** (196 µmol) and 33.3 mg 4-fluoro-3-methoxyaniline (236 µmol). Water was added, the formed precipitate was filtrated and recrystallized from DCM. 74 mg of **21** (139 µmol, 71 % yield) was obtained as a white solid.

**MS (ESI+)**: m/z = 531.25 $[M+H]^+$

**$^1$H NMR** (400 MHz, DMSO-$d_6$): δ = 12.11 (s, 1H), 10.16 (s, 1H), 10.01 (s, 1H), 8.34 (s, 1H), 7.79 (s, 1H), 7.55 (dd, *J* = 34.4, 7.0 Hz, 2H), 7.43 (s, 1H), 7.36 (d, *J* = 7.2 Hz, 1H), 7.30 – 7.03 (m, 3H), 3.79 (s, 3H), 3.42 (s, 4H), 3.17 (s, 4H), 1.49 (s, 4H) ppm.

**$^{13}$C NMR** (101 MHz, DMSO-$d_6$): δ = 168.39, 168.17, 159.68, 153.09, 150.20, 146.65, 146.54, 138.85, 135.68, 135.56, 135.53, 128.66, 123.08, 122.33, 120.18, 118.15, 115.80, 115.43, 115.24, 112.37, 112.30, 106.57, 102.32, 65.40, 55.78, 49.32, 31.48, 15.51 ppm.

**HRMS**: $[C_{28}H_{27}FN_6O_4+H]^+$ calculated: m/z = 531.21506, found: m/z = 531.21420

**HPLC**: $t_R$ = 3.716 min (Method 1), ≥ 95 % purity

**_N_-(3-(tert-butyl)phenyl)-_N_-(3-(4-morpholino-7_H_-pyrrolo[2,3-_d_]pyrimidin-5-yl)phenyl)cyclopropane-1,1-dicarboxamide (22)**

**22** was synthesized according to the **general procedure D** from 60 mg **70** (147 µmol) and 24.2 mg 3-(*tert*-butyl)aniline (162 µmol). Water was added, the formed precipitate was filtrated and washed with 5 mL ACN. 33 mg of **22** (61.3 µmol, 41 % yield) was obtained as a white solid.

**MS (ESI+)**: m/z = 539.25 $[M+H]^+$

**$^1$H NMR** (400 MHz, DMSO-$d_6$): δ = 12.11 (d, *J* = 2.1 Hz, 1H), 10.19 (s, 1H), 9.92 (s, 1H), 8.34 (s, 1H), 7.79 (t, *J* = 1.6 Hz, 1H), 7.62 (t, *J* = 1.9 Hz, 1H), 7.58 (d, *J* = 8.1 Hz, 1H), 7.49 (d, *J* = 8.0 Hz, 1H), 7.44 (d, *J* = 2.5 Hz, 1H), 7.37 (t, *J* = 7.9 Hz, 1H), 7.27 – 7.19 (m, 2H), 7.10 (ddd, *J* = 7.8, 1.6, 0.9 Hz, 1H), 3.46 – 3.40 (m, 4H), 3.20 – 3.14 (m, 4H), 1.54 – 1.45 (m, 4H), 1.26 (s, 9H) ppm.

**$^{13}$C NMR** (101 MHz, DMSO-$d_6$): δ = 168.44, 168.37, 159.71, 153.09, 151.03, 150.21, 138.82, 138.45, 135.70, 128.69, 128.09, 123.13, 122.35, 120.61, 120.20, 118.18, 117.83, 117.53, 115.81, 102.35, 65.41, 49.34, 34.43, 31.36, 31.11, 15.59 ppm.

**HRMS**: $[C_{31}H_{34}N_6O_3+H]^+$ calculated: m/z = 539.27652, found: m/z = 539.27631

**HPLC**: $t_R$ = 4.296 min (Method 1), ≥ 95 % purity

**_N_-(2,4-difluorophenyl)-_N_-(3-(4-morpholino-7_H_-pyrrolo[2,3-_d_]pyrimidin-5-yl)phenyl)cyclopropane-1,1-dicarboxamide (23)**

**23** was synthesized according to the **general procedure D** from 50 mg **70** (123 µmol) and 19 mg 2,4-difluoroaniline (147 µmol). The solvent was removed and the residue was purified via RP flash chromatography. 18 mg of **23** (34.7 µmol, 28 % yield) was obtained as a white solid.

**MS (ESI+)**: m/z = 519.15 $[M+H]^+$

**$^1$H NMR** (400 MHz, DMSO-$d_6$): δ = 12.37 (s, 1H), 10.32 (s, 1H), 10.18 (s, 1H), 8.40 (s, 1H), 7.87 – 7.75 (m, 2H), 7.51 (t, *J* = 6.9 Hz, 2H), 7.40 (t, *J* = 7.8 Hz, 1H), 7.37 – 7.29 (m, 1H), 7.26 (d, *J* = 7.3 Hz, 1H), 7.08 (t, *J* = 7.6 Hz, 1H), 3.49 – 3.38 (m, 4H), 3.29 – 3.19 (m, 4H), 1.67 – 1.54 (m, 4H) ppm.

**$^{13}$C NMR** (101 MHz, DMSO-$d_6$): δ = 169.07, 158.54, 151.75, 148.54, 138.45, 135.40, 128.85, 126.46, 126.36, 123.47, 122.87, 122.33, 122.22, 120.57, 118.75, 116.35, 111.21, 111.18, 110.99, 110.96, 104.35, 104.11, 104.09, 103.84, 101.94, 65.33, 49.42, 29.73, 16.83 ppm.

**HRMS**: $[C_{27}H_{24}F_2N_6O_3+H]^+$ calculated: m/z = 519.19507, found: m/z = 519.19471

**HPLC**: $t_R$ = 3.763 min (Method 1), ≥ 95 % purity

**_N_-(4-fluoro-2-methylphenyl)-_N_-(3-(4-morpholino-7_H_-pyrrolo[2,3-_d_]pyrimidin-5-yl)phenyl)cyclopropane-1,1-dicarboxamide (24)**

**24** was synthesized according to the **general procedure D** from 38 mg **70** (93.3 µmol) and 14.0 mg 4-fluoro-2-methylaniline (112 µmol). The solvent was removed and the residue was purified via RP flash chromatography. The resulting pale yellow solid was washed with 5 mL ACN. 24 mg of **24** (46.6 µmol, 50 % yield) was obtained as a white solid.
**MS (ESI+)**: m/z = 515.15 $[M+H]^+$
**$^1$H NMR** (400 MHz, DMSO-$d_6$): δ = 12.12 (s, 1H), 10.45 (s, 1H), 9.76 (s, 1H), 8.35 (s, 1H), 7.79 (s, 1H), 7.54 (d, *J* = 7.8 Hz, 1H), 7.50 – 7.42 (m, 2H), 7.38 (t, *J* = 7.9 Hz, 1H), 7.26 (d, *J* = 7.5 Hz, 1H), 7.09 (d, *J* = 8.4 Hz, 1H), 7.01 (t, *J* = 7.3 Hz, 1H), 3.43 (s, 4H), 3.17 (s, 4H), 2.20 (s, 3H), 1.57 (s, 4H) ppm.
**$^{13}$C NMR** (101 MHz, DMSO-$d_6$): δ = 169.24, 168.71, 160.73, 159.70, 158.33, 153.09, 150.21, 138.54, 135.77, 135.13, 135.05, 132.29, 132.27, 128.75, 127.27, 127.19, 123.32, 122.40, 120.17, 118.21, 116.61, 116.39, 115.74, 112.58, 112.36, 102.36, 65.41, 49.34, 29.78, 17.62, 16.44 ppm.
**HRMS**: $[C_{28}H_{27}FN_6O_3+H]^+$ calculated: m/z = 515.22014, found: m/z = 515.21978
**HPLC**: $t_R$ = 3.761 min (Method 1), ≥ 95 % purity

**_N_-(5-bromo-2,4-difluorophenyl)-_N_-(3-(4-morpholino-7_H_-pyrrolo[2,3-_d_]pyrimidin-5-yl)phenyl)cyclopropane-1,1-dicarboxamide (25)**

**25** was synthesized according to the **general procedure D** from 60 mg **70** (147 µmol) and 36.8 mg 5-bromo-2,4-difluoroaniline (177 µmol). The solvent was removed and the residue was purified via preparative HPLC. The resulting TFA salt was solved in EA and washed with 1M aq. $NaHCO_3$ solution. 20 mg of **25** (33.5 µmol, 22 % yield) was obtained as a white solid.
**MS (ESI+)**: m/z = 597.10 $[M+H]^+$
**$^1$H NMR** (400 MHz, DMSO-$d_6$): δ = 12.12 (s, 1H), 10.65 (s, 1H), 10.13 (s, 1H), 8.35 (s, 1H), 8.28 (s, 1H), 7.81 (s, 1H), 7.61 – 7.50 (m, 2H), 7.44 (s, 1H), 7.38 (t, *J* = 7.9 Hz, 1H), 7.26 (d, *J* = 7.4 Hz, 1H), 3.50 – 3.38 (m, 4H), 3.22 – 3.10 (m, 4H), 1.69 – 1.44 (m, 4H) ppm.
**$^{13}$C NMR** (101 MHz, DMSO-$d_6$): δ = 169.42, 169.24, 159.71, 154.48, 154.39, 153.10, 152.50, 152.41, 150.22, 138.55, 135.74, 128.70, 127.85, 123.30, 122.37, 120.50, 118.45, 115.78, 105.23, 102.36, 102.17, 102.00, 65.40, 49.33, 29.80, 17.12 ppm.
**HRMS**: $[C_{27}H_{23}BrF_2N_6O_3+H]^+$ calculated: m/z = 597.10558, found: m/z = 597.10531
**HPLC**: $t_R$ = 4.167 min (Method 1), ≥ 95 % purity

**_N_-(3-bromo-2,4-difluorophenyl)-_N_-(3-(4-morpholino-7_H_-pyrrolo[2,3-_d_]pyrimidin-5-yl)phenyl)cyclopropane-1,1-dicarboxamide (26)**

**26** was synthesized according to the **general procedure D** from 60 mg **70** (147 µmol) and 36.8 mg 3-bromo-2,4-difluoroaniline (177 µmol). The solvent was removed and the residue was purified via RP flash chromatography. 32 mg of **26** (53.6 µmol, 36 % yield) was obtained as a white solid.
**MS (ESI+)**: m/z = 599.05 $[M+H]^+$
**$^1$H NMR** (400 MHz, DMSO-$d_6$): δ = 12.12 (s, 1H), 10.41 (s, 1H), 10.16 (s, 1H), 8.35 (s, 1H), 7.84 – 7.75 (m, 2H), 7.54 (d, *J* = 8.6 Hz, 1H), 7.45 (d, *J* = 2.4 Hz, 1H), 7.39 (t, *J* = 7.9 Hz, 1H), 7.31 – 7.24 (m, 2H), 3.49 – 3.40 (m, 4H), 3.23 – 3.14 (m, 4H), 1.65 – 1.54 (m, 4H) ppm.
**$^{13}$C NMR** (101 MHz, DMSO-$d_6$): δ = 169.22, 168.76, 159.70, 157.21, 157.18, 154.79, 154.75, 153.31, 153.27, 153.09, 150.85, 150.81, 150.21, 138.44, 135.75, 128.74, 125.39, 125.30, 123.46,

123.36, 123.32, 123.23, 123.19, 122.42, 120.45, 118.48, 115.74, 111.74, 111.71, 111.52, 111.48, 102.35, 97.36, 97.13, 97.11, 96.88, 65.42, 49.34, 30.00, 16.79 ppm.
**HRMS**: $[C_{27}H_{23}BrF_2N_6O_3+H]^+$ calculated: m/z = 597.10558, found: m/z = 597.10548
**HPLC**: $t_R$ = 4.045 min (Method 1), ≥ 95 % purity

##### *N*-(3-bromo-5-fluorophenyl)-*N*-(3-(4-morpholino-7*H*-pyrrolo[2,3-*d*]pyrimidin-5-yl)phenyl)cyclopropane-1,1-dicarboxamide (27)

**27** was synthesized according to the **general procedure D** from 60 mg **70** (147 µmol) and 33.6 mg 3-bromo-5-fluoroaniline (177 µmol). The solvent was removed and the residue was purified via RP flash chromatography. 21 mg of **27** (36.2 µmol, 24 % yield) was obtained as a white solid.
**MS (ESI+)**: m/z = 581.05 $[M+H]^+$
**$^1$H NMR** (400 MHz, DMSO-*d6*): δ = 12.11 (d, *J* = 2.0 Hz, 1H), 10.32 (s, 1H), 10.07 (s, 1H), 8.34 (s, 1H), 7.77 (d, *J* = 1.1 Hz, 2H), 7.63 – 7.55 (m, 2H), 7.42 (d, *J* = 2.5 Hz, 1H), 7.37 (t, *J* = 7.9 Hz, 1H), 7.27 – 7.20 (m, 2H), 3.47 – 3.39 (m, 4H), 3.21 – 3.12 (m, 4H), 1.47 (s, 4H) ppm.
**$^{13}$C NMR** (101 MHz, DMSO-*d6*): δ = 168.73, 167.64, 163.20, 160.76, 159.66, 153.08, 150.20, 141.82, 141.69, 138.92, 135.67, 128.64, 123.10, 122.29, 121.55, 121.43, 120.22, 118.72, 118.69, 118.17, 115.81, 113.39, 113.14, 106.20, 105.94, 102.29, 65.39, 49.31, 32.14, 15.39 ppm.
**HRMS**: $[C_{27}H_{24}BrFN_6O_3+H]^+$ calculated: m/z = 579.11501, found: m/z = 579.11442
**HPLC**: $t_R$ = 4.162 min (Method 1), ≥ 95 % purity

##### 1-(3-(*tert*-butyl)-1-(quinolin-6-yl)-1*H*-pyrazol-5-yl)-3-(3-(4-morpholino-7*H*-pyrrolo[2,3-*d*]pyrimidin-5-yl)phenyl)urea (28)

60 mg **64** (112 µmol) and 54.2 mg **73** (123 µmol) were solved in 4 mL anhydrous DMF, before 58 µL DIPEA (335 µmol) was added. The reaction was heated to 60 °C and stirred overnight. The solvent was removed and the crude intermediate was treated according to the **general procedure C**. The crude product was purified via RP flash chromatography. 20 mg of **28** (34.0 µmol, 30 % yield) was obtained as a pale yellow solid.
**MS (ESI+)**: m/z = 588.25 $[M+H]^+$
**$^1$H NMR** (400 MHz, DMSO-*d6*): δ = 12.10 (d, *J* = 2.1 Hz, 1H), 9.10 (s, 1H), 8.95 (dd, *J* = 4.2, 1.6 Hz, 1H), 8.60 (s, 1H), 8.46 (d, *J* = 7.1 Hz, 1H), 8.34 (s, 1H), 8.19 – 8.14 (m, 2H), 7.97 (dd, *J* = 9.1, 2.3 Hz, 1H), 7.64 – 7.57 (m, 2H), 7.42 (d, *J* = 2.5 Hz, 1H), 7.36 – 7.28 (m, 2H), 7.16 – 7.11 (m, 1H), 6.46 (s, 1H), 3.47 – 3.40 (m, 4H), 3.21 – 3.11 (m, 4H), 1.32 (s, 9H) ppm.
**$^{13}$C NMR** (101 MHz, DMSO-*d6*): δ = 161.40, 159.66, 153.05, 151.54, 150.99, 150.18, 146.36, 139.43, 137.77, 136.33, 136.20, 135.97, 130.16, 128.97, 127.96, 126.50, 122.31, 122.18, 122.06, 121.74, 117.91, 115.89, 115.86, 102.34, 95.50, 65.40, 49.34, 32.13, 30.17 ppm.
**HRMS**: $[C_{33}H_{33}N_9O_2+H]^+$ calculated: m/z = 588.28300, found: m/z = 588.28173
**HPLC**: $t_R$ = 6.351 min (Method 1), ≥ 95 % purity

##### 1-(3-(*tert*-butyl)-1-(*p*-tolyl)-1*H*-pyrazol-5-yl)-3-(3-(4-morpholino-7*H*-pyrrolo[2,3-*d*]pyrimidin-5-yl)phenyl)urea (29)

**29** was synthesized according to the procedure of **28** from 199 mg **64** (371 µmol) and 150 mg **44** (371 µmol). 50 mg of **29** (90.8 µmol, 24 % yield) was obtained as a white solid.
**MS (ESI+)**: m/z = 551.30 $[M+H]^+$
**$^1$H NMR** (500 MHz, DMSO-*d6*): δ = 12.34 (s, 1H), 9.28 (s, 1H), 8.50 (s, 1H), 8.39 (s, 1H), 7.61 (s, 1H), 7.48 (d, *J* = 2.1 Hz, 1H), 7.41 (d, *J* = 8.3 Hz, 2H), 7.35 – 7.29 (m, 4H), 7.15 – 7.09 (m, 1H), 6.35 (s, 1H), 3.51 – 3.43 (m, 4H), 3.27 – 3.21 (m, 4H), 2.36 (s, 3H), 1.27 (s, 9H) ppm.

$^{13}$**C NMR** (101 MHz, DMSO-*d6*): δ = 160.49, 151.83, 151.73, 139.65, 137.05, 136.69, 136.09, 135.64, 129.64, 129.05, 124.21, 122.75, 121.66, 117.85, 116.56, 116.02, 101.99, 95.44, 65.35, 49.45, 40.43, 32.00, 30.20, 20.58 ppm.
**HRMS**: $[C_{31}H_{34}N_8O_2+H]^+$ calculated: m/z = 551.28775, found: m/z = 551.28696
**HPLC**: $t_R$ = 4.004 min (Method 1), ≥ 95 % purity

**1-(3-(*tert*-butyl)-1-(*p*-tolyl)-1*H*-pyrazol-5-yl)-3-(3-(2-chloro-4-morpholino-7*H*-pyrrolo[2,3-*d*]pyrimidin-5-yl)phenyl)urea (30)**

**30** was synthesized according to the procedure of **28** from 150 mg **45a** (262 µmol) and 159 mg **44** (393 µmol). 56 mg of **30** (95.7 µmol, 36 % yield) was obtained as a white solid.
**MS (ESI+)**: m/z = 585.25 $[M+H]^+$
$^{1}$**H NMR** (500 MHz, DMSO-*d6*): δ = 12.27 (d, *J* = 2.3 Hz, 1H), 9.11 (s, 1H), 8.35 (s, 1H), 7.58 (t, *J* = 1.7 Hz, 1H), 7.43 (d, *J* = 2.5 Hz, 1H), 7.42 – 7.38 (m, 2H), 7.36 – 7.28 (m, 4H), 7.09 (dt, *J* = 7.2, 1.5 Hz, 1H), 6.36 (s, 1H), 3.48 – 3.41 (m, 4H), 3.24 – 3.19 (m, 4H), 2.37 (s, 3H), 1.27 (s, 9H) ppm.
$^{13}$**C NMR** (101 MHz, DMSO-*d6*): δ = 160.52, 160.20, 153.87, 151.53, 151.15, 139.63, 137.08, 136.78, 136.05, 135.63, 129.67, 129.11, 124.32, 122.66, 121.63, 117.78, 116.41, 116.10, 100.72, 95.05, 65.28, 49.12, 32.00, 30.20, 20.58 ppm.
**HRMS**: $[C_{31}H_{33}ClN_8O_2+H]^+$ calculated: m/z = 585.24878, found: m/z = 585.24852
**HPLC**: $t_R$ = 4.899 min (Method 1), ≥ 95 % purity

**1-(3-(4-(4-acetylpiperazin-1-yl)-2-chloro-7*H*-pyrrolo[2,3-*d*]pyrimidin-5-yl)phenyl)-3-(3-(*tert*-butyl)-1-(*p*-tolyl)-1*H*-pyrazol-5-yl)urea (31)**

The **general procedure A** was conducted with the aryl halide 468 mg **74** (663 µmol) and the boronic acid 248 mg **75** (632 µmol). Water was added and the reaction mixture was extracted with DCM. The residue was treated according to **general procedure C**. The crude intermediate was purified via RP flash chromatography. The obtained white solid (50 mg, 85.6 µmol) was solved in 6 mL anhydrous THF and DCM (1:1), before 14.3 µL TEA (103 µmol) and 7.3 µL acetyl chloride (103 µmol) were added. The reaction was stirred at room temperature for 3 h. The solvent was removed and the crude product was purified via RP flash chromatography. 31 mg of **31** (49.5 µmol, 8 % yield) was obtained as a white solid.
**MS (ESI+)**: m/z = 626.35 $[M+H]^+$
$^{1}$**H NMR** (500 MHz, DMSO-*d6*): δ = 12.28 (s, 1H), 9.11 (s, 1H), 8.35 (s, 1H), 7.61 (s, 1H), 7.51 – 7.22 (m, 9H), 7.10 (s, 1H), 6.35 (s, 1H), 3.29 (s, 4H), 3.18 (s, 3H), 2.36 (s, 4H), 1.93 (s, 3H), 1.27 (s, 9H) ppm.
$^{13}$**C NMR** (101 MHz, DMSO-*d6*): δ = 168.18, 160.52, 160.05, 153.90, 151.56, 151.10, 139.65, 137.09, 136.79, 136.04, 135.61, 129.66, 129.15, 124.35, 122.73, 121.78, 117.82, 116.43, 116.18, 100.83, 95.00, 48.69, 44.74, 32.00, 30.20, 21.09, 20.57 ppm.
**HRMS**: $[C_{33}H_{36}ClN_9O_2+H]^+$ calculated: m/z = 626.27533, found: m/z = 626.2729
**HPLC**: $t_R$ = 4.409 min (Method 1), ≥ 95 % purity

**1-(3-(*tert*-butyl)-1-(*p*-tolyl)-1H-pyrazol-5-yl)-3-(3-(2-(ethylthio)-4-(methylamino)-7*H*-pyrrolo[2,3-*d*]pyrimidin-5-yl)phenyl)urea (32)**

**32** was synthesized according to the procedure of **28** from 122 mg **46** (225 µmol) and 146 mg **44** (360 µmol). 25 mg of **32** (45.1 µmol, 15 % yield) was obtained as a white solid.
**MS (ESI+)**: m/z = 555.25 $[M+H]^+$

$^{1}$**H NMR** (500 MHz, DMSO-*d6*): δ = 11.65 (d, *J* = 1.9 Hz, 1H), 9.11 (s, 1H), 8.40 (s, 1H), 7.50 (t, *J* = 1.6 Hz, 1H), 7.40 (d, *J* = 8.3 Hz, 2H), 7.33 (dd, *J* = 8.0, 2.3 Hz, 3H), 7.25 (dd, *J* = 8.1, 1.0 Hz, 1H), 7.09 – 7.05 (m, 2H), 6.38 (s, 1H), 5.78 (q, *J* = 4.5 Hz, 1H), 3.10 (q, *J* = 7.3 Hz, 2H), 2.92 (d, *J* = 4.7 Hz, 3H), 2.36 (s, 3H), 1.34 (t, *J* = 7.3 Hz, 3H), 1.27 (s, 9H) ppm.
$^{13}$**C NMR** (101 MHz, DMSO-*d6*): δ = 162.19, 160.51, 156.25, 151.85, 151.65, 139.64, 137.14, 136.81, 136.07, 135.65, 129.67, 129.56, 124.41, 122.12, 118.72, 118.54, 116.66, 115.82, 97.25, 94.88, 31.99, 30.21, 27.59, 24.20, 20.57, 15.20 ppm.
**HRMS**: $[C_{30}H_{34}N_8OS+H]^+$ calculated: m/z = 555.26491, found: m/z = 555.2635
**HPLC**: $t_R$ = 5.146 min (Method 1), ≥ 95 % purity

**(4-((5-chloro-4-(dimethylamino)-7*H*-pyrrolo[2,3-*d*]pyrimidin-2-yl)amino)-2-fluoro-5-methoxyphenyl)(morpholino)methanone (33)**

**33** was synthesized according to the **general procedure E** from 60 mg **79a** (104 µmol). The crude product was purified via RP flash chromatography and preparative HPLC. 18 mg of **33** (26.6 µmol, 25 % yield) was obtained as a light green twofold TFA salt.
**MS (ESI+)**: m/z = 449.10 $[M+H]^+$
$^{1}$**H NMR** (500 MHz, DMSO-*d6*): δ = 11.91 (s, 1H), 8.52 (d, *J* = 12.4 Hz, 1H), 7.88 (s, 1H), 7.23 (d, *J* = 2.5 Hz, 1H), 7.00 (d, *J* = 6.2 Hz, 1H), 3.92 (s, 3H), 3.64 (s, 4H), 3.56 (s, 2H), 3.31 (s, 2H), 3.24 (s, 6H) ppm.
$^{13}$**C NMR** (101 MHz, DMSO-*d6*): δ = 164.22, 158.24 (q, TFA), 152.67, 150.80, 143.58, 131.71, 131.62, 118.85, 116.63, 114.32, 113.87, 113.71, 109.82, 109.77, 104.49, 104.25, 102.29, 96.38, 66.30, 66.01, 56.54, 47.21, 42.06, 41.41 ppm.
**HRMS**: $[C_{20}H_{22}ClFN_6O_3+H]^+$ calculated: m/z = 449.14987, found: m/z = 449.14936
**HPLC**: $t_R$ = 4.252 min (Method 1), ≥ 95 % purity

**5-chloro-N2-(2-methoxy-4-(methylsulfonyl)phenyl)-N4,N4-dimethyl-7*H*-pyrrolo[2,3-*d*]pyrimidine-2,4-diamine (34)**

**34** was synthesized according to the **general procedure E** from 100 mg **79b** (190 µmol). The crude product was recrystallized from ACN. 55 mg of **34** (124 µmol, 65 % yield) was obtained as a grey solid.
**MS (ESI+)**: m/z = 396.10 $[M+H]^+$
$^{1}$**H NMR** (500 MHz, DMSO-*d6*): δ = 11.74 (s, 1H), 8.80 (d, *J* = 8.6 Hz, 1H), 7.68 (s, 1H), 7.50 (dd, *J* = 8.6, 1.9 Hz, 1H), 7.44 (d, *J* = 2.0 Hz, 1H), 7.19 (d, *J* = 2.6 Hz, 1H), 4.00 (s, 3H), 3.21 (s, 6H), 3.19 (s, 3H) ppm.
$^{13}$**C NMR** (101 MHz, DMSO-*d6*): δ = 158.90, 153.30, 151.97, 146.56, 134.54, 131.34, 120.31, 118.58, 115.87, 108.28, 101.92, 96.78, 56.36, 43.99, 41.22 ppm.
**HRMS**: $[C_{16}H_{18}ClN_5O_3S+H]^+$ calculated: m/z = 396.08916, found: m/z = 396.08899
**HPLC**: $t_R$ = 4.260 min (Method 1), ≥ 95 % purity

**1-(3-(*tert*-butyl)-1-(*p*-tolyl)-1*H*-pyrazol-5-yl)-3-(3-(4-(dimethylamino)-2-((2-methoxy-4-(methylsulfonyl)phenyl)amino)-7*H*-pyrrolo[2,3-*d*]pyrimidin-5-yl)phenyl)urea (35)**

The **general procedure A** was conducted with the aryl halide **79c** (75 mg, 121 µmol) and the boronic acid **75** (50 mg, 128 µmol). The solvent was removed and the residue was treated according to the **general procedure E**. The crude product was purified via RP flash chromatography and preparative HPLC. 21 mg of **35** (22.4 µmol, 18 % yield) was obtained as a yellow twofold TFA salt.

**MS (ESI+)**: m/z = 708.30 $[M+H]^+$

**$^1$H NMR** (500 MHz, DMSO-$d_6$): δ = 12.35 (s, 1H), 9.14 (s, 1H), 8.80 (d, *J* = 8.5 Hz, 1H), 8.49 (s, 2H), 7.85 (s, 1H), 7.54 (dd, *J* = 8.6, 2.0 Hz, 1H), 7.50 (d, *J* = 2.0 Hz, 1H), 7.48 – 7.44 (m, 1H), 7.43 – 7.39 (m, 2H), 7.36 – 7.28 (m, 4H), 7.04 (d, *J* = 1.7 Hz, 1H), 6.38 (s, 1H), 4.03 (s, 3H), 3.42 (s, 6H), 3.21 (s, 3H), 2.38 (s, 3H), 1.28 (s, 9H) ppm.

**$^{13}$C NMR** (101 MHz, DMSO-$d_6$): δ = 160.53, 158.24 (q, TFA), 151.64, 147.32, 139.88, 137.10, 136.80, 136.05, 133.34, 132.74, 132.62, 131.83, 129.68, 129.30, 124.36, 120.16, 118.76, 117.43, 117.21, 117.12, 114.79, 114.43, 108.67, 100.51, 98.60, 95.22, 56.47, 43.90, 32.02, 30.23, 20.60 ppm.

**HRMS**: $[C_{37}H_{41}N9O_4S+H]^+$ calculated: m/z = 708.30750, found: m/z = 708.30706

**HPLC**: $t_R$ = 4.501 min (Method 1), ≥ 95 % purity

**1-(3-(*tert*-butyl)-1-(*p*-tolyl)-1*H*-pyrazol-5-yl)-3-(3-(4-(dimethylamino)-2-((5-fluoro-2-methoxy-4-(morpholine-4-carbonyl)phenyl)amino)-7*H*-pyrrolo[2,3-*d*]pyrimidin-5-yl)phenyl)urea (36)**

The **general procedure E** was conducted with **80a** (62.5 mg, 98.3 µmol). The crude intermediate was recrystallized from ACN and water. The obtained grey solid was resolved in 5 mL anhydrous DMSO, before 64 mg **44** (158 µmol) and 51.7 µL DIPEA (297 µmol) were added. The reaction was heated to 60 °C and stirred overnight. The solvent was removed and the crude product was purified via RP flash chromatography. 43 mg of **36** (56.5 µmol, 57 % yield) was obtained as an off-white solid.

**MS (ESI+)**: m/z = 761.35 $[M+H]^+$

**$^1$H NMR** (500 MHz, DMSO-$d_6$): δ = 11.71 (s, 1H), 9.09 (s, 1H), 8.63 (d, *J* = 12.6 Hz, 1H), 8.37 (s, 1H), 7.51 (t, *J* = 1.6 Hz, 1H), 7.42 – 7.38 (m, 2H), 7.37 – 7.31 (m, 3H), 7.31 – 7.24 (m, 2H), 7.10 (d, *J* = 2.3 Hz, 1H), 7.05 (dt, *J* = 7.1, 1.5 Hz, 1H), 6.99 (d, *J* = 6.3 Hz, 1H), 6.36 (s, 1H), 3.93 (s, 3H), 3.65 (s, 4H), 3.57 (s, 4H), 2.80 (s, 6H), 2.37 (s, 3H), 1.27 (s, 9H) ppm.

**$^{13}$C NMR** (101 MHz, DMSO-$d_6$): δ = 164.34, 160.50, 152.82, 151.52, 150.95, 143.23, 139.47, 137.19, 137.07, 136.78, 136.49, 136.27, 136.05, 129.68, 129.34, 128.79, 124.40, 123.44, 121.92, 119.30, 117.66, 117.12, 115.80, 109.66, 109.62, 96.92, 94.83, 73.69, 66.31, 66.03, 56.51, 47.24, 42.08, 40.69, 31.99, 30.20, 30.13, 20.58 ppm.

**HRMS**: $[C_{41}H_{45}FN_{10}O_4+H]^+$ calculated: m/z = 761.3682, found: m/z = 761.3660

**HPLC**: $t_R$ = 4.412 min (Method 1), ≥ 95 % purity

**1-(3-(*tert*-butyl)-1-(quinolin-6-yl)-1*H*-pyrazol-5-yl)-3-(3-(4-(dimethylamino)-2-((2-methoxy-4-(methylsulfonyl)phenyl)amino)-7*H*-pyrrolo[2,3-*d*]pyrimidin-5-yl)phenyl)urea (37)**

120 mg of **80b** (206 µmol) and 182 mg **73** (412 µmol) were solved in 10 mL anhydrous DMSO, before 108 µL DIPEA (618 µmol) was added. The reaction was heated to 70 °C and stirred overnight. The solvent was removed and the residue was treated according to **general procedure E**. The crude product was purified via preparative HPLC. 27 mg of **37** (27.8 µmol, 13 % yield) was obtained as a green twofold TFA salt.

**MS (ESI+)**: m/z = 745.35 $[M+H]^+$

**$^1$H NMR** (500 MHz, DMSO-$d_6$): δ = 12.34 (s, 1H), 9.12 (s, 1H), 9.01 (dd, *J* = 4.2, 1.4 Hz, 1H), 8.79 (d, *J* = 8.6 Hz, 1H), 8.75 (s, 1H), 8.57 (d, *J* = 8.0 Hz, 1H), 8.22 (d, *J* = 2.2 Hz, 1H), 8.20 (d, *J* = 9.0 Hz, 1H), 8.02 (dd, *J* = 9.0, 2.2 Hz, 1H), 7.84 (s, 1H), 7.68 (dd, *J* = 8.3, 4.3 Hz, 1H), 7.55

(dd, $J$ = 8.6, 1.7 Hz, 1H), 7.51 (d, $J$ = 1.5 Hz, 1H), 7.46 (d, $J$ = 7.4 Hz, 1H), 7.35 – 7.28 (m, 2H), 7.05 (s, 1H), 6.48 (s, 1H), 4.04 (s, 3H), 3.42 (s, 6H), 3.22 (s, 3H), 1.33 (s, 9H) ppm.
**$^{13}$C NMR** (101 MHz, DMSO-$d_6$): δ = 161.69, 158.36 (q, TFA), 151.83, 150.56, 147.51, 145.39, 139.96, 137.94, 137.63, 136.67, 131.96, 129.79, 129.48, 129.45, 128.24, 127.12, 122.43, 122.19, 120.31, 119.03, 117.70, 117.09, 114.77, 114.67, 108.88, 100.74, 98.71, 96.07, 56.65, 44.05, 32.32, 30.33 ppm.
**HRMS**: $[C_{39}H_{40}N_{10}O_4S+H]^+$ calculated: m/z = 745.30275, found: m/z = 745.30151
**HPLC**: $t_R$ = 4.031 min (Method 1), ≥ 95 % purity

**2,4-dichloro-5-iodo-7-trityl-7*H*-pyrrolo[2,3-*d*]pyrimidine (40)**

10.0 g 2,4-dichloro-7*H*-pyrrolo[2,3-*d*]pyrimidine (53.2 mmol) was solved in 50 mL anhydrous DCM and 15.6 g NIS (69.1 mmol) was added in portions. The reaction was stirred at room temperature overnight. The precipitate was filtrated and washed with 20 mL DCM. The white intermediate product (**39**) was resolved in 50 mL chloroform. 10.7 mL TEA (76.5 mmol) was added, before 11.7 g (chloromethanetriyl)tribenzene (42.1 mmol) was added in portions. The reaction was stirred at room temperature overnight. The solvent was removed and the residue was suspended in MeOH. The solid was filtrated and washed with 20 mL MeOH. 20.8 g of **40** (37.4 mmol, 98 % yield) was obtained as a white solid.
**MS (ESI+):** m/z = 243.05 $[C_{25}H_{16}Cl_2IN_3\text{-}C_6HCl_2IN_3^-]^+$
**$^1$H NMR** (250 MHz, $CDCl_3$): δ = 7.43 (s, 1H), 7.34 – 7.27 (m, 9H), 7.19 – 7.10 (m, 6H) ppm.

**4-(2-chloro-5-iodo-7-trityl-7*H*-pyrrolo[2,3-*d*]pyrimidin-4-yl)morpholine (41a)**

**41a** synthesized according to the **general procedure B** from 5.00 g **40** (8.99 mmol) and 1.55 mL morpholine (18.0 mmol). 5.38 g of **41a** (8.86 mmol, 98 % yield) was obtained as a white solid.
**MS (ESI+):** m/z = 243.05 $[CPh_3\text{-}M^-]^+$
**$^1$H NMR** (250 MHz, $CDCl_3$): δ = 7.25 – 7.16 (m, 9H), 7.14 (d, $J$ = 0.5 Hz, 1H), 7.13 – 7.04 (m, 6H), 3.90 – 3.77 (m, 4H), 3.58 – 3.45 (m, 4H) ppm.

**2-chloro-5-iodo-*N*-methyl-7-trityl-7*H*-pyrrolo[2,3-*d*]pyrimidin-4-amine (41b)**

**41b** synthesized according to the **general procedure B** from 5.00 g **40** (8.99 mmol) and 8.99 mL methylamine (2 M inTHF, 18.0 mmol). 4.89 g of **41b** (8.88 mmol, 99 % yield) was obtained as a white solid.
**MS (ESI+):** m/z = 308.9 $[M\text{-}CPh_3^+ + 2H^+]^+$
**$^1$H NMR** (250 MHz, $CDCl_3$): δ = 7.22 – 7.13 (m, 9H), 7.10 – 7.02 (m, 6H), 6.88 (s, 1H), 6.05 (q, $J$ = 4.3 Hz, 1H), 3.03 (d, $J$ = 4.9 Hz, 3H) ppm.

**3-(*tert*-butyl)-1-(*p*-tolyl)-1*H*-pyrazol-5-amine (43)**

5.00 g *p*-tolylhydrazine (40.9 mmol) and 5.12 g 4,4-dimethyl-3-oxopentanenitrile (40.9 mmol) were solved in 50 mL anhydrous methanol. The reaction was heated to 80 °C and stirred overnight. After quant. conversion, the solvent was removed in vacuo. The crude product **43** was obtained as a pale yellow solid and was used for the synthesis of **44** without further purification.
**MS (ESI+)**: m/z = 230.15 $[M+H]^+$
**$^1$H NMR** (250 MHz, DMSO-$d_6$): δ = 7.55 – 7.32 (m, 1H), 5.71 (s, 1H), 2.40 (s, 1H), 1.32 (s, 2H), 1.09 (s, 1H) ppm.

**2,2,2-trichloroethyl (3-(*tert*-butyl)-1-(*p*-tolyl)-1*H*-pyrazol-5-yl)carbamate (44)**

A solution of 9.39 g **43** (41 mmol) in 100 mL EA and water (2:1), was cooled to 0 °C and 4.09 g sodium hydroxide (102 mmol) was added, followed by 6.76 mL trichloroethylchloroformate (49.1 mmol). The reaction mixture was warmed to room temperature and stirred for 1 hour. The layers were separated and the organic layer was washed with brine. 16.5 g of **44** (40.8 mmol,99 % yield) was obtained as a pale brown solid.

**MS (ESI+)**: m/z = 404.05 $[M+H]^+$

**$^1$H NMR** (500 MHz, DMSO-$d_6$): δ = 9.90 (s, 1H), 7.37 (d, *J* = 8.4 Hz, 2H), 7.26 (d, *J* = 8.3 Hz, 2H), 6.27 (s, 1H), 4.85 (s, 2H), 2.34 (s, 3H), 1.28 (s, 9H) ppm.

**3-(2-chloro-4-morpholino-7-trityl-7*H*-pyrrolo[2,3-*d*]pyrimidin-5-yl)aniline (45a)**

**45a** was synthesized according to the **general procedure A** from 5.50 g **41a** (9.06 mmol) and 1.37 g (3-aminophenyl)boronic acid (9.97 mmol). Water was added and the reaction mixture was extracted with DCM. The crude orange product was recrystallized from EtOH. 5.02 g of **45a** (8.77 mmol, 97 % yield) was obtained as a white solid

**MS (ESI+):** m/z = 572.20 $[M+H]^+$

**$^1$H NMR** (250 MHz, $CDCl_3$): δ = 7.25 – 7.02 (m, 16H), 6.96 (s, 1H), 6.69 (d, *J* = 7.5 Hz, 1H), 6.61 (s, 1H), 6.52 (d, *J* = 7.8 Hz, 1H), 3.50 – 3.40 (m, 4H), 3.28 – 3.19 (m, 4H) ppm.

**5-(3-aminophenyl)-2-chloro-*N*-methyl-7-trityl-7*H*-pyrrolo[2,3-*d*]pyrimidin-4-amine (45b)**

**45b** was synthesized according to the **general procedure A** from 3.80 g **41b** (6.90 mmol) and 1.13 g (3-aminophenyl)boronic acid (8.28 mmol). Water was added and the precipitate was filtrated and recrystallized from EtOH. 3.51 g of **45b** (6.80 mmol, 99 % yield) was obtained as a white solid.

**MS (ESI+):** m/z = 516.15 $[M+H]^+$

**$^1$H NMR** (250 MHz, DMSO-$d_6$): δ = 7.39 – 7.03 (m, 15H), 6.84 – 6.64 (m, 2H), 6.63 – 6.45 (m, 3H), 5.88 (q, *J* = 4.3 Hz, 1H), 5.24 (s, 2H), 2.86 (d, *J* = 4.7 Hz, 3H) ppm.

**5-(3-aminophenyl)-2-(ethylthio)-*N*-methyl-7-trityl-7*H*-pyrrolo[2,3-*d*]pyrimidin-4-amine (46)**

300 mg **45b** (581 µmol) and 489 mg sodium ethyl thiolate (5.81 mmol) were suspended in 3 mL anhydrous DMF. The reaction was heated to 90 °C in the microwave and stirred for 1 h. Water was added and the precipitate was filtrated and washed with 20 mL EtOH. 250 mg of **46** (461 µmol, 79 % yield) was obtained as a white solid.

**MS (ESI+):** m/z = 542.20 $[M+H]^+$

**$^1$H NMR** (250 MHz, $CDCl_3$): δ = 9.35 – 9.06 (m, 16H), 8.78 – 8.49 (m, 4H), 7.10 (q, *J* = 4.4 Hz, 1H), 5.73 (s, 2H), 4.86 (d, *J* = 4.8 Hz, 3H), 4.30 (q, *J* = 7.2 Hz, 2H), 2.78 (t, *J* = 7.2 Hz, 3H) ppm.

***N*-(3-(2*H*-indazol-5-yl)phenyl)-*N*-(4-fluorophenyl)cyclopropane-1,1-dicarboxamide (47)**

The **general procedure A** was conducted with the boronic acid **63** (80 mg, 234 µmol) and the aryl halide **83** (84.2 mg, 257 µmol). The solvent was removed and the residue was solved in 8 mL anhydrous THF, before 2.34 mL tetrabutylammoniumfluorid (1 M in THF, 2.34 mmol) was added. The mixture was heated to 80 °C and stirred overnight. Water was added and the reaction mixture was extracted with EA. The crude product was purified via flash chromatography (DCM/MeOH). 60 mg of **47** (145 µmol, 62 % yield) was obtained as a white solid.

**MS (ESI+)**: m/z = 415.15 $[M+H]^+$

**$^{1}$H NMR** (400 MHz, DMSO-*d$_6$*): δ = 13.12 (s, 1H), 10.14 (s, 1H), 10.05 (s, 1H), 8.13 (d, *J* = 1.2 Hz, 1H), 8.00 (s, 1H), 7.98 (s, 1H), 7.69 – 7.56 (m, 5H), 7.45 – 7.34 (m, 2H), 7.19 – 7.08 (m, 2H), 1.49 (d, *J* = 2.4 Hz, 4H) ppm.
**$^{13}$C NMR** (101 MHz, DMSO-*d$_6$*): δ = 168.24, 168.20, 159.45, 157.07, 141.11, 139.36, 135.17, 135.14, 133.99, 132.61, 129.06, 125.49, 123.51, 122.46, 122.39, 122.01, 121.93, 118.91, 118.74, 118.13, 115.12, 114.90, 110.56, 31.50, 15.47 ppm.
**HRMS**: $[C_{24}H_{19}FN_4O_2+H]^+$ calculated: m/z = 415.15648, found: m/z = 415.15641
**HPLC**: $t_R$ = 7.692 min (Method 1), ≥ 95 % purity

***N*-(4-((2*H*-indazol-5-yl)oxy)phenyl)-*N*-(4-fluorophenyl)cyclopropane-1,1-dicarboxamide (48)**

80 mg **83** (244 µmol), 92 mg **81a** (293 µmol), 156 mg potassium phosphate (733 µmol), 6 mg *t*BuBrettPhos (12 µmol) and 10 mg *t*BuBrettPhos Pd G3 (12 µmol) were suspended in toluene and dimethoxyethane (4:1). The reaction was heated to 100 °C and stirred overnight. The reaction mixture was filtrated through Celite®, the solvent was removed and the residue was solved in 8 mL anhydrous THF, before 2.44 mL tetrabutylammoniumfluorid (1 M in THF, 2.44 mmol) was added. The mixture was heated to 80 °C and stirred overnight. Water was added and the reaction mixture was extracted with EA. The crude product was purified via flash chromatography (DCM/MeOH) and RP flash chromatography. 21 mg of **48** (48.8 µmol, 20 % yield) was obtained as a white solid.
**MS (ESI+)**: m/z = 431.05 $[M+H]^+$
**$^{1}$H NMR** (400 MHz, DMSO-*d$_6$*): δ = 13.08 (s, 1H), 10.07 (s, 1H), 10.00 (s, 1H), 8.00 (s, 1H), 7.66 – 7.53 (m, *J* = 22.7, 12.3, 6.9 Hz, 5H), 7.29 (d, *J* = 2.0 Hz, 1H), 7.18 – 7.07 (m, 3H), 6.97 – 6.90 (m, 2H), 1.45 (s, 4H) ppm.
**$^{13}$C NMR** (101 MHz, DMSO-*d$_6$*): δ = 168.24, 168.01, 159.45, 157.06, 153.78, 150.57, 136.94, 135.14, 133.95, 133.33, 123.18, 122.40, 122.32, 122.27, 119.77, 117.99, 115.14, 114.92, 111.53, 108.46, 31.27, 15.44 ppm.
**HRMS**: $[C_{24}H_{19}FN_4O_3+H]^+$ calculated: m/z = 431.15140, found: m/z = 431.15118
**HPLC**: $t_R$ = 4.313 min (Method 1), ≥ 95 % purity

***N*-(4-((2*H*-indazol-5-yl)amino)phenyl)-*N*-(4-fluorophenyl)cyclopropane-1,1-dicarboxamide (49)**

100 mg **83** (306 µmol), 115 mg **81a** (367 µmol), 299 mg caesium carbonate (917 µmol), 8 mg BrettPhos (15 µmol) and 14 mg BrettPhos Pd G4 (15 µmol) were suspended in toluene and *tert*-butanol (4:1). The reaction was heated to 110 °C and stirred overnight. The reaction mixture was filtrated through Celite®, the solvent was removed and the residue was solved in 8 mL anhydrous THF, before 1.83 mL tetrabutylammoniumfluorid (1 M in THF, 1.83 mmol) was added. The mixture was heated to 80 °C and stirred overnight. Water was added and the reaction mixture was extracted with EA. The crude product was purified via flash chromatography (DCM/MeOH) and RP flash chromatography. 42 mg of **49** (97.8 µmol, 32 % yield) was obtained as a white solid.
**MS (ESI+)**: m/z = 430.15 $[M+H]^+$
**$^{1}$H NMR** (400 MHz, DMSO-*d$_6$*): δ = 12.84 (s, 1H), 10.16 (s, 1H), 9.79 (s, 1H), 7.89 (s, 2H), 7.68 – 7.58 (m, 2H), 7.43 (dd, *J* = 13.7, 8.8 Hz, 3H), 7.35 (d, *J* = 1.1 Hz, 1H), 7.19 – 7.08 (m, 3H), 6.96 (d, *J* = 8.9 Hz, 2H), 1.45 (d, *J* = 2.6 Hz, 4H) ppm.
**$^{13}$C NMR** (101 MHz, DMSO-*d$_6$*): δ = 168.45, 167.85, 159.45, 157.07, 141.35, 136.85, 136.08, 135.13, 132.57, 130.27, 123.48, 122.37, 122.30, 122.25, 121.06, 115.58, 115.17, 114.95, 110.79, 106.14, 30.88, 15.55 ppm.

**HRMS**: $[C_{24}H_{20}FN_5O_2+H]^+$ calculated: m/z = 430.16738, found: m/z = 430.16542
**HPLC**: $t_R$ = 7.161 min (Method 2), ≥ 95 % purity

**3-((2*H*-indazol-5-yl)amino)-4-methyl-*N*-(3-(4-methyl-1*H*-imidazol-1-yl)-5-(trifluoromethyl)phenyl)benzamide (50)**

**50** was synthesized according to the procedure of **49** from 80 mg **86** (214 µmol). 17 mg of **50** (34.7 µmol, 18 % yield) was obtained as a white solid.
**MS (ESI+)**: m/z = 491.15 $[M+H]^+$
**$^1$H NMR** (400 MHz, DMSO-*d6*): δ = 12.94 (s, 1H), 10.49 (s, 1H), 8.26 (s, 1H), 8.17 (s, 1H), 8.10 (s, 1H), 7.95 (t, *J* = 2.4 Hz, 1H), 7.68 (s, 1H), 7.57 (d, *J* = 1.5 Hz, 1H), 7.51 (d, *J* = 8.8 Hz, 1H), 7.47 – 7.42 (m, 2H), 7.37 (s, 1H), 7.35 – 7.30 (m, 2H), 7.20 (dd, *J* = 8.9, 2.0 Hz, 1H), 2.33 (s, 3H), 2.16 (d, *J* = 0.4 Hz, 3H) ppm.
**$^{13}$C NMR** (101 MHz, DMSO-*d6*): δ = 166.15, 144.33, 141.40, 138.87, 137.85, 136.66, 136.49, 134.93, 132.70, 132.38, 130.92, 130.76, 130.60, 130.54, 124.99, 123.48, 122.28, 118.34, 114.82, 114.46, 114.18, 111.36, 110.86, 109.24, 18.12, 13.54 ppm.
**HRMS**: $[C_{26}H_{21}F_3N_6O+H]^+$ calculated: m/z = 491.18017, found: m/z = 491.18039
**HPLC**: $t_R$ = 6.360 min (Method 1), ≥ 95 % purity

***N*-(4-fluorophenyl)-*N*-(1*H*-indazol-6-yl)cyclopropane-1,1-dicarboxamide (51)**

80 mg **62** (358 µmol), 434 µL pyridine (5.38 mmol), 426 µL propanephosphonic acid anhydride (T3P, 50 wt. % in EA, 717 µmol) and 53 mg 1*H*-indazol-6-amine (394 µmol) were solved in 3 mL ACN and stirred for 2 h at room temperature. Water was added and the resulting solid was filtrated and purified via RP flash chromatography. 61 mg of **51** (179 µmol, 50 % yield) was obtained as a white solid.
**MS (ESI+)**: m/z = 339.05 $[M+H]^+$
**$^1$H NMR** (500 MHz, DMSO-*d6*): δ = 12.91 (s, 1H), 10.21 (s, 1H), 9.99 (s, 1H), 8.09 (s, 1H), 7.96 (s, 1H), 7.71 – 7.55 (m, 3H), 7.25 – 7.07 (m, 3H), 1.49 (s, 4H) ppm.
**$^{13}$C NMR** (101 MHz, DMSO-*d6*): δ = 168.37, 168.15, 159.24, 157.33, 140.21, 136.97, 135.13, 135.11, 133.25, 122.53, 122.47, 120.27, 119.32, 115.09, 114.91, 100.27, 31.58, 15.48 ppm.
**HRMS**: $[C_{18}H_{15}FN_4O_2+H]^+$ calculated: m/z = 339.12518, found: m/z = 339.1244
**HPLC**: $t_R$ = 3.927 min (Method 1), ≥ 95 % purity

**3-((1*H*-indazol-6-yl)ethynyl)-4-methyl-*N*-(4-((4-methylpiperazin-1-yl)methyl)-3-(trifluoromethyl)phenyl)benzamide (52)**

100 mg **58** (241 µmol) and 65 mg 6-iodo-1*H*-indazole (265 µmol) was solved in 3 mL anhydrous DMF. 16.9 mg bis(triphenylphosphine)palladium(II)-chloride (24 µmol), 4.6 mg copper(I) iodide (24 µmol) and 69 µL DIPEA (722 mmol) were added. The reaction was heated to 80 °C and stirred overnight. The reaction mixture was filtrated through Celite®, the solvent was removed in vacuo and the residue was purified via RP flash chromatography. 64 mg **52** (120 mmol, 50 % yield) was obtained as a light yellow solid.
**MS (ESI+)**: m/z = 532.20 $[M+H]^+$
**$^1$H NMR** (500 MHz, DMSO-*d6*): δ = 13.26 (s, 1H), 10.55 (s, 1H), 9.35 (s, 1H), 8.25 – 8.14 (m, 3H), 8.12 (d, *J* = 8.2 Hz, 1H), 7.92 (dd, *J* = 7.9, 1.2 Hz, 1H), 7.85 (d, *J* = 8.3 Hz, 1H), 7.79 (s, 1H), 7.71 (d, *J* = 8.5 Hz, 1H), 7.53 (d, *J* = 8.1 Hz, 1H), 7.30 (d, *J* = 8.3 Hz, 1H), 3.67 (s, 2H), 3.46 – 2.99 (m, 8H), 2.78 (s, 3H), 2.59 (s, 3H) ppm.

$^{13}$**C NMR** (101 MHz, DMSO-*d*$_6$): δ = 164.71, 143.77, 138.49, 134.56, 134.48, 132.05, 131.44, 130.93, 130.50, 129.93, 128.07, 127.62, 127.39, 125.32, 123.44, 123.30, 123.14, 122.30, 121.20, 119.31, 117.31 (q, $CF_3$), 113.29, 94.78, 87.12, 56.56, 52.97, 49.64, 42.43, 20.39 ppm.
**HRMS**: $[C_{30}H_{28}F_3N_5O+H]^+$ calculated: m/z = 532.23187, found: m/z = 532.2311
**HPLC**: $t_R$ = 3.546 min (Method 1), ≥ 95 % purity

**1-(3-(*tert*-butyl)-1-(quinolin-6-yl)-1*H*-pyrazol-5-yl)-3-(1*H*-indazol-6-yl)urea (53)**

40 mg 1*H*-indazol-6-amine (300 µmol) and 146 mg **73** (330 µmol) were solved in DMSO, before 86 µL DIPEA (901 µmol) was added. The reaction was heated to 70 °C and stirred overnight. Water was added and the reaction mixture was extracted with EA. The crude product was purified via RP flash chromatography. 70 mg of **53** (165 µmol, 55 % yield) was obtained as a white solid.
**MS (ESI+)**: m/z = 426.20 $[M+H]^+$
$^1$**H NMR** (500 MHz, DMSO-*d*$_6$): δ = 9.15 (s, 1H), 8.96 (dd, *J* = 4.2, 1.7 Hz, 1H), 8.61 (s, 1H), 8.48 (dd, *J* = 8.5, 1.1 Hz, 1H), 8.20 – 8.16 (m, 2H), 7.98 (dd, *J* = 9.1, 2.4 Hz, 1H), 7.93 (s, 2H), 7.63 – 7.59 (m, 2H), 6.85 (dd, *J* = 8.7, 1.7 Hz, 1H), 6.50 (s, 1H), 1.33 (s, 9H) ppm.
$^{13}$**C NMR** (101 MHz, DMSO-*d*$_6$): δ = 161.43, 151.61, 150.99, 146.37, 140.59, 137.78, 137.53, 136.36, 136.22, 133.26, 130.18, 127.97, 126.55, 122.18, 122.08, 120.74, 118.58, 113.58, 97.42, 95.39, 32.14, 30.19 ppm.
**HRMS**: $[C_{24}H_{23}N_7O+H]^+$ calculated: m/z = 426.20368, found: m/z = 426.2032
**HPLC**: $t_R$ = 3.607 min (Method 1), ≥ 95 % purity

**1-(3-(*tert*-butyl)-1-(*p*-tolyl)-1*H*-pyrazol-5-yl)-3-(1*H*-indazol-6-yl)urea (54)**

**54** was synthesized according to the procedure of **53** from 100 mg 1*H*-indazol-6-amine (751 µmol) and 334 mg **44** (826 µmol). 144 mg of **54** (371 µmol, 49 % yield) was obtained as a white solid.
**MS (ESI+)**: m/z = 389.05 $[M+H]^+$
$^1$**H NMR** (500 MHz, DMSO-*d*$_6$): δ = 12.81 (s, 1H), 9.16 (s, 1H), 8.36 (s, 1H), 7.93 (s, 2H), 7.61 (d, *J* = 8.6 Hz, 1H), 7.41 (d, *J* = 8.4 Hz, 2H), 7.34 (d, *J* = 8.3 Hz, 2H), 6.84 (dd, *J* = 8.7, 1.6 Hz, 1H), 6.39 (s, 1H), 2.37 (s, 3H), 1.28 (s, 9H) ppm.
$^{13}$**C NMR** (101 MHz, DMSO-*d*$_6$): δ = 160.52, 151.58, 140.60, 137.62, 137.12, 136.80, 136.05, 133.25, 129.69, 124.38, 120.72, 118.51, 113.51, 97.28, 94.92, 32.01, 30.22, 20.59 ppm.
**HRMS**: $[C_{22}H_{24}N_6O+H]^+$ calculated: m/z = 389.20844, found: m/z = 389.2075
**HPLC**: $t_R$ = 4.258 min (Method 1), ≥ 95 % purity

**5-bromo-4-chloro-7-trityl-7*H*-pyrrolo[2,3-*d*]pyrimidine (56a)**

5.0 g 5-bromo-4-chloro-7*H*-pyrrolo[2,3-*d*]pyrimidine (21.5 mmol) was solved in 50 mL anhydrous chloroform and 6.0 mL triethylamine (43.0 mmol) was added. The solution was cooled to 0 °C, before 7.20 g tritylchloride (25.8 mmol) was added in portions and the reaction was stirred at room temperature overnight. The solvent was removed, the crude product was washed with ethanol and dried in vacuo. 9.48 g **56a** (20.0 mmol, 93 % yield) was obtained as a white solid.
**MS (ESI+):** m/z = 485.05 $[M+H]^+$
$^1$**H NMR** (250 MHz, $CDCl_3$): δ = 8.29 (s, 1H), 7.34 – 7.26 (m, 10H), 7.19 – 7.10 (m, 6H) ppm.

**4-chloro-5-iodo-7-trityl-7*H*-pyrrolo[2,3-*d*]pyrimidine (56b)**

**56b** was synthesized according to the procedure of **56a** from 10 g 4-chloro-5-iodo-7*H*-pyrrolo[2,3-*d*]pyrimidine (35.8 mmol). 17.9 g of **56b** (34.3 mmol, 96 % yield) was obtained as a white solid.

**MS (ESI+):** m/z = 544.16 [M+Na]$^+$
$^1$**H NMR** (250 MHz, DMSO-*d6*): δ = 8.31 (s, 1H), 7.45 (s, 1H), 7.39 – 7.27 (m, 9H), 7.19 – 7.09 (m, 6H) ppm.

**methyl 3-ethynyl-4-methylbenzoate (57)**

1.0 g Methyl-3-Iodo-4-methyl-benzoate (3.62 mmol) was solved in 5 mL anhydrous ACN. 51 mg bis(triphenylphosphine)palladium(II)-chloride (73 µmol), 14 mg copper(I) iodide (73 µmol) and 1.51 mL TEA (10.9 mmol) were added. 0.62 mL ethinyltrimethylsilane (4.35 mmol) was added dropwise. The reaction was heated to 60 °C and stirred overnight. The reaction mixture was filtrated through Celite® and concentrated in vacuo. The residue was solved in 10 mL methanol and 1.5 g potassium carbonate (10.9 mmol) was added. The mixture was stirred for 1 h at room temperature and filtrated through Celite®. The solvent was removed in vacuo and the residue was purified via flash chromatography (*n*-hexane/EA). 473 mg of **57** (2.72 mmol, 75 % yield) was obtained as an orange solid.
**MS (ESI+)**: m/z = 175.25 [M+H]$^+$
$^1$**H NMR** (400 MHz, DMSO-*d6*): δ = 7.94 (d, *J* = 1.8 Hz, 1H), 7.86 (dd, *J* = 8.0, 1.8 Hz, 1H), 7.45 (d, *J* = 8.0 Hz, 1H), 4.50 (s, 1H), 3.84 (s, 3H), 2.45 (s, 3H) ppm.

**3-ethynyl-4-methyl-*N*-(4-((4-methylpiperazin-1-yl)methyl)-3-(trifluoromethyl)phenyl)benzamide (58)**

355 mg **57** (2.04 mmol) and 557 mg 4-((4-methylpiperazin-1-yl)methyl)-3-(trifluoromethyl)aniline (2,04 mmol) were solved in 10 mL anhydrous THF. The solution was cooled to -20 °C, before 343 mg potassium-*tert*-butanolate (3.06 mmol) was added. The reaction was stirred for 1.5 h while warming slowly to room temperature and stirred overnight. The reaction mixture was poured on water and extracted with EA. The residue was purified via flash chromatography (DCM/MeOH). 683 mg **58** (1.64 mmol, 80 % yield) was obtained as a red solid.
**MS (ESI+)**: m/z = 416.34 [M+H]$^+$
$^1$**H NMR** (500 MHz, DMSO-*d6*): δ = 10.48 (s, 1H), 8.19 (d, *J* = 2.1 Hz, 1H), 8.08 (d, *J* = 1.9 Hz, 1H), 8.04 (dd, *J* = 8.5, 1.9 Hz, 1H), 7.90 (dd, *J* = 8.0, 1.9 Hz, 1H), 7.70 (d, *J* = 8.6 Hz, 1H), 7.47 (d, *J* = 8.1 Hz, 1H), 4.51 (s, 1H), 3.56 (s, 2H), 2.46 (s, 3H), 2.43 – 2.25 (m, 8H), 2.16 (s, 3H) ppm.

**3-((4-chloro-7-trityl-7*H*-pyrrolo[2,3-*d*]pyrimidin-5-yl)ethynyl)-4-methyl-*N*-(4-((4-methylpiperazin-1-yl)methyl)-3-(trifluoromethyl)phenyl)benzamide (59)**

1.00 g **58** (2.41 mmol) and 1.38 g **56b** (2.65 mmol) was solved in 3 mL anhydrous DMF. 16.9 mg bis(triphenylphosphine)palladium(II)-chloride (24 µmol), 4.6 mg copper(I) iodide (24 µmol) and 69 µL DIPEA (722 mmol) were added. The reaction was heated to 80 °C and stirred overnight. The reaction mixture was filtrated through Celite®, the solvent was removed in vacuo and the residue was purified via flash chromatography (DCM/MeOH). 664 mg **59** (820 µmol, 34 % yield) was obtained as an orange solid.
**MS (ESI+):** m/z = 809.30 [M+H]$^+$
$^1$**H NMR** (250 MHz, DMSO-*d6*): δ = 10.49 (s, 1H), 8.40 (s, 1H), 8.16 (dd, *J* = 16.3, 1.8 Hz, 2H), 8.04 (dd, *J* = 8.5, 1.8 Hz, 1H), 7.89 (dd, *J* = 8.0, 1.8 Hz, 1H), 7.79 (s, 1H), 7.70 (d, *J* = 8.4 Hz, 1H), 7.49 (d, *J* = 8.2 Hz, 1H), 7.41 – 7.27 (m, 9H), 7.25 – 7.15 (m, 6H), 2.56 (s, 3H), 2.45 – 2.29 (m, 8H), 2.29 – 2.20 (m, 2H), 2.16 (s, 3H) ppm.

**4-(5-bromo-7-trityl-7*H*-pyrrolo[2,3-*d*]pyrimidin-4-yl)morpholine (60a)**

**60a** was synthesized according to the **general procedure B** from 5.0 g **56a** (10.5 mmol) and 1.09 mL morpholine (12.6 mmol). 5.38 g of **60a** (10.2 mmol, 97 % yield) was obtained as a white solid.
**MS (ESI+):** m/z = 525.10 $[M+H]^+$
**$^1$H NMR** (250 MHz, $CDCl_3$): δ = 8.10 (s, 1H), 7.33 – 7.27 (m, 9H), 7.20 – 7.12 (m, 6H), 7.10 (s, 1H), 3.98 – 3.88 (m, 4H), 3.73 – 3.61 (m, 4H) ppm.

**4-(5-iodo-7-trityl-7*H*-pyrrolo[2,3-*d*]pyrimidin-4-yl)morpholine (60b)**

**60b** was synthesized according to the **general procedure B** from 2.8 g **56b** (5.37 mmol) and 0.56 mL morpholine (5.44 mmol). 2.97 g of **60b** (5.18 mmol, 97 % yield) was obtained as a white solid.
**MS (ESI+):** m/z = 573.10 $[M+H]^+$
**$^1$H NMR** (250 MHz, $CDCl_3$): δ = 8.11 (s, 1H), 7.33 – 7.26 (m, 13H), 7.21 (s, 1H), 7.19 – 7.11 (m, 8H), 4.06 – 3.87 (m, 5H), 3.68 – 3.54 (m, 4H) ppm.

**1-((4-fluorophenyl)carbamoyl)cyclopropane-1-carboxylic acid (62)**

2 g cyclopropane-1,1-dicarboxylic acid (15.4 mmol) and 2.25 mL triethyl amine (16.1 mmol) were solved in 40 mL of anhydrous THF. After the solution was cooled to 0 °C 1.92 g thionyl chloride (16.1 mmol) were added dropwise. 4.44 g 4-fluoro aniline (16.9 mmol) solved in 10 mL anhydrous THF was added dropwise to the solution, which was allowed to warm to room temperature (rt) and stirred overnight (on). The pH was adjusted to 9 with 1 M NaOH solution and afterwards the solution was brought to pH 4 with 4 M HCl (aq). The mixture was filtrated, and the filtrate was extracted with ethyl acetate (EA). The organic phase was washed with 1 M NaOH (aq). The watery phase was adjusted to pH 4 with 4 M HCl (aq) and was extracted with EA. 2.39 g of **62** (10.7 mmol, 70 % yield) was obtained as an off-white solid.
**MS (ESI+)**: m/z = 224.0 $[M+H]^+$
**$^1$H NMR** (250 MHz, DMSO-$d_6$): δ = 10.57 (s, 1H), 7.67-7.57 (m, 2H), 7.20-7.07 (m, 2H), 1.42 (s, 4H) ppm.

**(3-(1-((4-fluorophenyl)carbamoyl)cyclopropane-1-carboxamido)phenyl)boronic acid (63)**

348 mg of **62** (1.56 mmol) and 711 mg HATU (1.87 mmol) were dissolved in anhydrous 5 mL DMF, before 543 µL DIPEA (3.12 mmol) and 235 mg (3-aminophenyl)-boronic acid (1.72 mmol) were added. The reaction mixture was stirred over night at room temperature. Water was added and the mixture was extracted with EA. The obtained solid was purified via RP flash chromatography. 391 g of **63** (1.14 mmol, 73 % yield) was obtained as a white solid.
**MS (ESI+)**: m/z = 343.05 $[M+H]^+$
**$^1$H NMR** (250 MHz, DMSO-$d_6$): δ = 10.05 (s, 2H), 8.00 (s, 2H), 7.85 (s, 1H), 7.77 – 7.68 (m, 1H), 7.67 – 7.57 (m, 2H), 7.51 (d, $J$ = 7.3 Hz, 1H), 7.27 (t, $J$ = 7.7 Hz, 1H), 7.20 – 7.07 (m, 2H), 1.48 (s, 4H) ppm.

**3-(4-morpholino-7-trityl-7*H*-pyrrolo[2,3-*d*]pyrimidin-5-yl)aniline (64)**

**64** was synthesized according to the **general procedure A** from the aryl halide **60a** (3.0 g, 5.71 mmol) and (3-aminophenyl)boronic acid (938 mg, 6.85 mmol). Water was added, the precipitated solid was filtrated and washed with water and ethanol. 2.63 g of **64** (4.89 mmol, 86 % yield) was obtained as a white solid.

**MS (ESI+):** m/z = 538.20 $[M+H]^+$
**$^1$H NMR** (250 MHz, DMSO-$d_6$): δ = 7.98 (s, 1H), 7.37 – 7.24 (m, 9H), 7.20 – 7.11 (m, 6H), 7.04 (t, *J* = 7.7 Hz, 1H), 6.96 (s, 1H), 6.62 (s, 1H), 6.57 – 6.45 (m, 2H), 5.16 (s, 2H), 3.52 – 3.44 (m, 4H), 3.23 – 3.14 (m, 4H) ppm.

**1-((4-fluorophenyl)carbamoyl)cyclopentane-1-carboxylic acid (65)**

**65** was synthesized according to the procedure of **62** from 1 g cyclopentane-1,1-dicarboxylic acid (6.32 mmol). The resulting crude product was purified via flash chromatography (DCM/MeOH). 240 mg of **65** (955 µmol, 15 % yield) was obtained as an orange solid.
**MS (ESI+):** m/z = 252.05 $[M+H]^+$
**$^1$H NMR** (250 MHz, DMSO-$d_6$): δ = 12.56 (s, 1H), 9.59 (s, 1H), 7.71 – 7.49 (m, 2H), 7.25 – 7.00 (m, 2H), 2.37 – 1.95 (m, 4H), 1.77 – 1.47 (m, 4H) ppm.

**1-(4-fluorophenyl)-2-oxo-1,2-dihydropyridine-3-carboxylic acid (66)**

1.00 g methyl 2-oxo-2H-pyran-3-carboxylate (6.49 mmol) and 4-fluoroaniline (6.81 mmol) were solved in 10 mL anhydrous DMF. The reaction was cooled to 0 °C and stirred for 5 h. 1.31 g EDC (8.43 mmol) and 198 mg DMAP (1.62 mmol) was added and the reaction was stirred at room temperature overnight. The solvent was removed and the intermediate product was purified via flash chromatography (*n*-hexane/EA). The obtained red solid was solved in THF and water (5:1) and 849 mg lithium hydroxide monohydrate (20.2 mmol) was added. The reaction was stirred at room temperature overnight. 0.5 M $HCl_{aq}$ was added and the reaction mixture was extracted with EA. 955 mg of **66** (4.10 mmol, 63 % yield) was obtained as a light brown solid.
**MS (ESI+):** m/z = 234.00 $[M+H]^+$
**$^1$H NMR** (250 MHz, DMSO-$d_6$): δ = 14.22 (s, 1H), 8.49 (ddd, *J* = 7.2, 2.0, 1.1 Hz, 1H), 8.20 (ddd, *J* = 6.6, 2.1, 1.1 Hz, 1H), 7.73 – 7.54 (m, 2H), 7.52 – 7.32 (m, 2H), 6.91 – 6.70 (m, 1H) ppm.

**4-methyl-3-(4,4,5,5-tetramethyl-1,3,2-dioxoborolan-2-yl)aniline (67a)**

791 mg potassium acetate (8.06 mmol), 611 mg bis(pinacolato)diboron (2.41 mmol) and 89.3 mg Pd(dppf)$Cl_2$ (134 µmol) were suspended in 10 mL anhydrous dioxane. A solution of 500 mg 3-bromo-4-methylaniline (2.69 mmol) in 5 mL anhydrous dioxane was added dropwise. The reaction was heated to 90 °C and stirred overnight. The reaction mixture was filtered through Celite®. Water was added to the filtrate and was extracted with EA. The crude product was purified via flash chromatography (DCM/MeOH). 215 mg of **67a** (922 µmol, 34 % yield) was obtained as brown resin.
**MS (ESI+):** m/z = 234.28 $[M+H]^+$
**$^1$H NMR** (400 MHz, DMSO-$d_6$): δ = 6.92 (d, *J* = 2.6 Hz, 1H), 6.81 (d, *J* = 8.1 Hz, 1H), 6.50 (dd, *J* = 8.0 Hz, *J* = 2.6 Hz, 1H), 4.78 (s, 2H), 2.27 (s, 3H), 1.27 (s, 12H) ppm.

**3-methyl-5-(4,4,5,5-tetramethyl-1,3,2-dioxoborolan-2-yl)aniline (67b)**

1.00 g 3-bromo-5-methylaniline (5.37 mmol), 1.14 g bis(pinacolato)diboron (8.06 mmol), 2.16 g potassium ethylhexanoate (11.8 mmol) and 128 mg XPhos (269 µmol) were solved in 20 mL anhydrous dioxane and heated to 70 °C. 49.2 mg AllylPdCl (270 µmol) was added, and the reaction was stirred overnight. 1M $NaHCO_3$ solution was added and the reaction mixture was extracted with EA. The crude product was purified by flash chromatography (*n*-hexane/EA). 242 mg **67b** (1.08 mmol, 20 % yield) was obtained as pale-yellow solid.
**MS (ESI+):** m/z = 234.27 $[M+H]^+$

**$^1$H NMR** (400 MHz, DMSO-*d*$_6$): δ = 6.74 (d, 1H, *J* = 2.0 Hz), 6.65 (s, 1H) 6.48 (s, 1H), 4.92 (s, 2H), 2.14 (s, 2H), 1.26 (s, 12H) ppm.

**2-methyl-5-(4,4,5,5-tetramethyl-1,3,2-dioxoborolan-2-yl)aniline (67c)**

**67c** was synthesized according to the procedure of **67a** from 1.00 g 5-bromo-2-methylaniline (5.37 mmol). The crude product was purified via flash chromatography (*n*-hexane/EA). 1.04 g of **67c** (4.46 mmol, 83 %yield) was obtained as white solid.
**MS (ESI+):** m/z = 233.90 $[M+H]^+$
**$^1$H NMR** (400 MHz, DMSO-*d*$_6$): δ = 6.98 (d, *J* = 0.8 Hz, 1H), 6.92 (d, 1H, *J* = 7.3 Hz, 1H, *H*-3), 6.79 (dd, *J* = 7.2 Hz, *J* = 1.0 Hz, 1H), 4.76 (s, 2H), 2.05 (s, 3H), 1.26 (s, 12H) ppm.

**4-methyl-3-(4-:Horpholino-7-trityl-7*H*-pyrrolo[2,3-*d*]pyrimidin-5-yl)aniline (68a)**

**68a** was synthesized according to the **general procedure A** from the aryl halide **60b** (202 mg, 353 µmol) and the pinacol ester **67a** (100 mg, 429 µmol). Water was added, the precipitated solid was filtrated and washed with water and ethanol. 145 mg of **68a** (263 µmol, 74 % yield) was obtained as an off-white solid.
**MS (ESI+):** m/z = 552.11 $[M+H]^+$
**$^1$H NMR** (400 MHz, DMSO-*d*$_6$): δ = 7.95 (s, 1H), 7.36 – 7.25 (m, 9H), 7.18 – 7.08 (m, 7H), 6.91 (d, *J* = 8.0 Hz, 1H), 6.78 (s, 1H), 6.49 – 6.44 (m, 2H), 4.91 (s, 2H), 3.47–3.44 (m, 4H), 3.15–3.13 (m, 4H), 1.95 (s, 3H) ppm.

**3-methyl-5-(4-morpholino-7-trityl-7*H*-pyrrolo[2,3-*d*]pyrimidin-5-yl)aniline (68b)**

**68b** was synthesized according to the **general procedure A** from the aryl halide **60b** (297 mg, 519 µmol) and the pinacol ester **67b** (145 mg, 622 µmol). Water was added, the precipitated solid was filtrated and washed with water and ethanol. 283 mg of **68b** (513 µmol, 99 % yield) was obtained as an off-white solid.
**MS (ESI+):** m/z = 552.00 $[M+H]^+$
**$^1$H NMR** (400 MHz, DMSO-*d*6): 7.97 (s, 1H), 7.35 – 7.25 (m, 9H), 7.18 – 7.07 (m, 6H), 6.96 (s, 1H), 6.42 (s, *J* = 16.4 Hz, 1H), 6.38 (s, 1H), 6.31 (s, 1H), 5.06 (s, 2H), 3.47–3.44 (m, 4H), 3.19–3.16 (m, 4H),2.18 (s, 3H) ppm.

**2-methyl-5-(4-morpholino-7-trityl-7*H*-pyrrolo[2,3-*d*]pyrimidin-5-yl)aniline (68c)**

**68c** was synthesized according to the **general procedure A** from the aryl halide **60b** (409 mg, 715 µmol) and the pinacol ester **67c** (200 mg, 858 µmol). Water was added, the precipitated solid was filtrated and washed with water and ethanol. 96.0 mg of **68c** (174 µmol, 24 % yield) was obtained as an off-white solid.
**MS (ESI+):** m/z = 552.06 $[M+H]^+$
**$^1$H NMR** (400 MHz, DMSO-*d*$_6$): δ = 7.97 (s, 1H), 7.34–7.25 (m, 9H), 7.15–7.14 (m, 6H), 6.94 – 6.92 (m, 2H), 6.68 (s, 1H), 6.50 (d, J = 7.5 Hz, 1H), 4.92 (s, 2H), 3.47– 3.45 (m, 4H), 3.19–3.17 (m, 4H), 2.06 (s, 3H) ppm.

**(3-aminophenyl)(4-morpholino-7-trityl-7*H*-pyrrolo[2,3-*d*]pyrimidin-5-yl)methanone (69)**

To a cooled solution of 500 mg **60a** (952 µmol) in 20 mL anhydrous THF, 455 µL *n*BuLi (2.5 M in *n*-hexane, 1.14 mmol) was added at -78 °C. The reaction was stirred for 1 h, before 212 mg 3-nitrobenzoyl chloride (1.14 mmol) was added. The reaction was slowly warmed room temperature and stirred for 1 h. Water was added and the reaction mixture was extracted with EA. The

intermediate was purified via flash chromatography (*n*-hexane/EA). The obtained white solid was suspended in 18 mL methanol together with 136 mg iron powder (2.43 mmol) and 2 mL conc. aq. ammonium chloride solution. The reaction was heated to 70 °C and stirred overnight. The reaction mixture was filtrated through Celite®. Water was added to the filtrate, and it was extracted with EA. 120 mg **69** (212 µmol, 22 % yield) was obtained as a white solid.
**MS (ESI+):** m/z = 566.25 $[M+H]^+$
$^1$**H NMR** (250 MHz, DMSO-*d*6): δ = 7.99 (s, 1H), 7.45 (s, 1H), 7.37 – 6.89 (m, 18H), 6.77 (d, *J* = 8.4 Hz, 1H), 5.38 (s, 2H), 3.56 – 3.46 (m, 4H), 3.45 – 3.36 (m, 4H) ppm.

**1-((3-(4-morpholino-7*H*-pyrrolo[2,3-*d*]pyrimidin-5-yl)phenyl)carbamoyl)cyclopropane-1-carboxylic acid (70)**

726 mg cyclopropane-1,1-dicarboxylic acid (5.58 mmol) was solved in 30 mL anhydrous THF and 405 µL thionyl chloride (5.58 mmol) was added slowly. The solution was heated to 60 °C and stirred for 2 h. The raction mixture was cooled to room temperature, before a suspension of 2.50 g **64** (4.65 mmol) in 20 mL anhydrous THF was added slowly and stirred overnight. The precipitated white solid was filtrated and washed with water. A white solid was obtained. The **general procedure C** was conducted with the intermediate. The watery layer was adjusted to pH3 with 4M aq. HCl and extracted with EA. The crude product was recrystallized from ACN/MeOH. 1.02 g **70** (2.51 mmol, 54 % yield) was obtained as a white solid.
**MS (ESI+):** m/z = 408.10 $[M+H]^+$
$^1$**H NMR** (250 MHz, DMSO-$d_6$): δ = 13.76 (s, 1H), 12.15 (s, 1H), 8.34 (s, 1H), 7.75 (s, 1H), 7.53 (d, *J* = 8.1 Hz, 1H), 7.46 (d, *J* = 1.7 Hz, 1H), 7.33 (t, *J* = 7.7 Hz, 1H), 7.17 (d, *J* = 7.4 Hz, 1H), 3.47 (s, 4H), 3.17 (s, 4H), 1.24 (d, *J* = 14.1 Hz, 4H) ppm.

**quinolin-6-yl trifluoromethanesulfonate (71.0)**

2.00 g quinolin-6-ol (13.8 mmol) and 2.25 mL pyridine (27.9 mmol) were solved in 50 mL anhydrous DCM. The solution was cooled to 0 °C and 16.5 mL of a 1 M solution of sulfonic anhydride (16.5 mmol) in DCM was added dropwise. The cooling bath was removed and the reaction solution was stirred overnight at room temperature. The reaction mixture was washed with water and brine. The crude product was purified by flash chromatography (*n*-hexane/EA 80/20 → 50/50). 3.06 g of **71.0** (11.0 mmol, 80 % yield) was obtained as a colorless oil.
**MS (ESI+):** m/z = 278.08 $[M+H]^+$
$^1$**H NMR** (400 MHz, DMSO-$d_6$): δ = 9.03 (dd, *J* = 4.2, 1.7 Hz, 1H), 8.54 – 8.52 (m, 1H), 8.24 (d, *J* = 2.9 Hz, 1H), 8.21 (d, *J* = 9.3 Hz, 1H), 7.86 (dd, *J* = 9.3, 2.9 Hz, 1H), 7.67 (dd, *J* = 8.4, 4.2 Hz, 1H) ppm.

**6-(2-(diphenylmethylene)hydrazineyl)quinoline (71)**

1.80 g of **71.0** (6.49 mmol), 1.66 g (diphenylmethylene)hydrazine (8.44 mmol), 4.23 g $Cs_2CO_3$ (13.0 mmol), 153 mg XPhos Pd G2 (195 µmol) and 93 mg XPhos (195 µmol) were suspended in 40 mL anhydrous toluene and 8mL *t*-BuOH. The reaction mixture was heated to 90 °C and stirred overnight. The cooled reaction mixture was concentrated *in vacuo* and the residue was rinsed with EA and filtered over Celite®. The crude product was purified via flash chromatography (*n*-hexane/EA 90/10 → 50/50). 1.60 g of **71** (4.95 mmol, 76 % yield) was obtained as a yellow solid.
**MS (ESI+):** m/z = 324.15 $[M-H]^-$

**$^1$H NMR** (400 MHz, DMSO-*d6*): δ = 9.24 (s, 1H), 8.61 (dd, *J* = 4.2, 1.7 Hz, 1H), 8.19 – 8.12 (m, 1H), 7.85 (d, *J* = 9.2 Hz, 1H, *H*-6), 7.76 (dd, *J* = 9.2, 2.4 Hz, 1H), 7.67 – 7.51 (m, 7H), 7.40 – 7.29 (m, 6H) ppm.

**3-(*tert*-butyl)-1-(quinolin-6-yl)-1*H*-pyrazol-5-amine (72)**

1.53 g **71** (4.73 mmol) and 1.78 g 4,4-dimethyl-3-oxopentanenitrile (14.2 mmol) were solved with 7 mL ethanol. 4 mL concentrated HCl (12 M, 48.0 mmol) was added. The mixture was heated to reflux and stirred overnight. The cooled reaction mixture was concentrated *in vacuo* and the residue was washed with diethyl ether. The crude product was dissolved in EA and washed with saturated $Na_2CO_3$ solution. The residue was purified by flash chromatography (*n*-hexane/EA 70/30 → 30/70). 590 mg of **72** (2.22 mmol, 69 % yield) was obtained as an orange oil.
**MS (ESI+):** *m/z* = 267.27 $[M+H]^+$
**$^1$H NMR** (400 MHz, DMSO-*d6*): δ = 8.88 (dd, *J* = 4.2, 1.7 Hz, 1H), 8.40 (dd, *J* = 8.3, 1.7 Hz, 1H), 8.12 (d, *J* = 1.7 Hz, 1H), 8.11 – 8.03 (m, 2H), 7.55 (dd, *J* = 8.3, 4.2 Hz, 1H), 5.46 (s, 1H), 5.41 (s, 2H), 1.25 (s, 9H) ppm.

**2,2,2-trichloroethyl (3-(*tert*-butyl)-1-(quinolin-6-yl)-1*H*-pyrazol-5-yl)carbamate (73)**

1.30 g of **72** (4.87 mmol), 16.0 mg 4-DMAP (132 µmol) and 1.30 mL pyridine (16.1 mmol) were solved in 30 mL anhydrous DCM and cooled to – 10 °C. A solution of 2,2,2-trichloroethyl carbonochloridate (950 µL, 6.91 mmol) in 3 mL DCM was added dropwise to the cooled solution over a period of 20 min. After 2 h of stirring, 20 mL $H_2O$ were added, and the solution was stirred for 10 min. The mixture was washed with brine. The crude product was purified by flash chromatography (*n*-hexane/EA 90/10 → 50/50). 689 mg of **73** (1.56 mmol, 32 % yield) was obtained as a yellowish solid.
**MS (ESI+):** *m/z* = 441.19 $[M+H]^+$
**$^1$H NMR** (250 MHz, (DMSO-*d6*): δ = 10.15 (s, 1H), 8.94 (dd, *J* = 4.2, 1.7 Hz, 1H), 8.43 – 8.38 (m, 1H), 8.17 – 8.03 (m, 2H), 7.90 (dd, *J* = 9.0, 2.4 Hz, 1H), 7.59 (dd, *J* = 8.3, 4.2 Hz, 1H), 6.38 (s, 1H), 4.85 (s, 2H), 1.31 (s, 9H) ppm.

***tert*-butyl 4-(2-chloro-5-iodo-7-trityl-7*H*-pyrrolo[2,3-*d*]pyrimidin-4-yl)piperazine-1-carboxylate (74)**

**74** synthesized according to the **general procedure B** from 5.00 g **39** (8.99 mmol) and 1.76 g *tert*-butyl piperazine-1-carboxylate (9.44 mmol). 6.35 g of **74** (8.99 mmol, 100 % yield) was obtained as a white solid.
**MS (ESI+):** m/z = 436.20 $[M-CPh_3^-]^+$
**$^1$H NMR** (250 MHz, DMSO-*d6*): δ = 7.40 – 7.26 (m, 9H), 7.15 (s, 1H), 7.14 – 7.06 (m, 1H), 3.62 – 3.41 (m, 8H), 1.42 (s, 9H) ppm.

**(3-(3-(3-(*tert*-butyl)-1-(*p*-tolyl)-1*H*-pyrazol-5-yl)ureido)phenyl)boronic acid (75)**

500 mg (3-aminophenyl)boronic acid (3.65 mmol) and 1.48 g **44** (3.65 mmol) were solved in 20 mL anhydrous ACN, before 1.91 mL DIPEA (20.0 mmol) was added. The reaction was heated to 70 °C and stirred 3 h. 1 M aq. HCl was added and the reaction mixture was extracted with EA. The crude product was purified via RP flash chromatography. 888 mg of **75** (2.26 µmol, 62 % yield) was obtained as a pale yellow solid.
**MS (ESI+):** m/z = 393.15 $[M+H]^+$

$^1$**H NMR** (250 MHz, DMSO-*d*$_6$): δ = 8.94 (s, 1H), 8.30 (s, 1H), 7.98 (s, 2H), 7.66 (s, 1H), 7.56 (d, *J* = 8.0 Hz, 1H), 7.45 – 7.30 (m, 5H), 7.23 (t, *J* = 7.7 Hz, 1H), 6.37 (s, 1H), 2.38 (s, 3H), 1.28 (s, 9H) ppm.

**2,4,5-trichloro-7*H*-pyrrolo[2,3-*d*]pyrimidine (76a.0)**

4.00 g 2,4-dichloro-7*H*-pyrrolo[2,3-*d*]pyrimidine (21.3 mmol) and 3.41 g NCS (25.5 mmol) were solved in 40 mL anhydrous ACN. The reaction was heated to 70 °C and stirred overnight. The precipitate was filtrated and washed with 20 mL ACN. 2.66 g of **76a.0** (12.0 mmol, 56 % yield) was obtained as a white solid.
**MS (ESI+):** m/z = 221.90 $[M+H]^+$
$^1$**H NMR** (250 MHz, DMSO-*d*$_6$): δ = 13.05 (s, 1H), 7.92 (dd, *J* = 2.5, 0.9 Hz, 1H) ppm.

**2,4,5-trichloro-7-((2-(trimethylsilyl)ethoxy)methyl)-7*H*-pyrrolo[2,3-*d*]pyrimidine (76a)**

2.60 g **76a.0** (11.7 mmol) was solved in anhydrous DMF and the solution was cooled to 0 °C. 701 mg sodium hydride (60 wt. % in paraffin, 17.5 mmol) was added in portions and the reaction was stirred for 0.5 h. 2.48 mL (2-(chloromethoxy)ethyl)trimethylsilane (14.0 mmol) was added dropwise. The reaction was warmed to room temperature and stirred for 2 h. Water was added and the reaction mixture was extracted with EA. The crude product was purified via flash chromatography (*n*-hexane/EA). 4.03 g of **76a** (11.3 mmol, 97 % yield) was obtained as a white solid.
**MS (ESI+):** m/z = 352.00 $[M+H]^+$
$^1$**H NMR** (250 MHz, DMSO-*d*$_6$): δ = 8.14 (s, 1H), 5.56 (s, 2H), 3.61 – 3.47 (m, 2H), 0.92 – 0.78 (m, 2H), -0.08 (s, 9H) ppm.

**2,4-dichloro-7-((2-(trimethylsilyl)ethoxy)methyl)-7*H*-pyrrolo[2,3-*d*]pyrimidine (76b)**

**76b** was synthesized according to the procedure of **76a** from 2,4-dichloro-7*H*-pyrrolo[2,3-*d*]pyrimidine (1.00 g, 5.32 mmol). 1.55 g **76b** (4.87 mmol, 91 % yield) was obtained as a white solid.
**MS (ESI+):** m/z = 318.00 $[M+H]^+$
$^1$**H NMR** (250 MHz, $CDCl_3$): δ = 7.37 (d, *J* = 3.7 Hz, 1H), 6.66 (d, *J* = 3.7 Hz, 1H), 5.60 (s, 2H), 3.60 – 3.45 (m, 2H), 0.98 – 0.85 (m, 2H), -0.04 (s, 9H) ppm.

**2,5-dichloro-*N,N*-dimethyl-7-((2-(trimethylsilyl)ethoxy)methyl)-7*H*-pyrrolo[2,3-*d*] pyrimidin-4-amine (77a)**

4.12 g **76a** (11.7 mmol) was solved in 20 mL abs. EtOH, before 8.76 mL dimethylamine (2 M in THF, 17.5 mmol) and 4.07 mL DIPEA (23.4 mmol) were added. The reaction was stirred at room temperature overnight. Water was added and the reaction mixture was extracted with EA. The combined organic layers were washed with 1 M aq. HCl, with 1 M aueous $NaHCO_3$ and with brine. 4.20 g of **77a** (11.6 mmol, 99 % yield) was obtained as a red resin.
**MS (ESI+):** m/z = 361.00 $[M+H]^+$
$^1$**H NMR** (250 MHz, DMSO-*d*$_6$): δ = 7.63 (s, *J* = 6.8 Hz, 1H), 5.43 (s, 2H), 3.57 – 3.46 (m, 2H), 3.23 (s, 6H), 0.89 – 0.78 (m, 2H), -0.08 (s, *J* = 3.3 Hz, 9H) ppm.

**2-chloro-*N,N*-dimethyl-7-((2-(trimethylsilyl)ethoxy)methyl)-7*H*-pyrrolo[2,3-*d*] pyrimidin-4-amine (77b)**

**77b** was synthesized according to the procedure of **77a** from 1.69 g **76b** (5.31 mmol). 1.69 g of **77b** (5.18 mmol, 97 % yield) was obtained as a light yellow resin.
**MS (ESI+):** m/z = 327.10 $[M+H]^+$
$^1$**H NMR** (250 MHz, DMSO-$d_6$): δ = 7.30 (dd, *J* = 3.7, 0.6 Hz, 1H), 6.73 (dd, *J* = 3.7, 0.7 Hz, 1H), 5.44 (s, 2H), 3.53 – 3.44 (m, 2H), 3.28 (s, 6H), 0.88 – 0.78 (m, 2H), -0.08 (d, *J* = 0.8 Hz, 9H) ppm.

**(4-amino-2-fluoro-5-methoxyphenyl)(morpholino)methanone (78a)**

4.75 g 2,5-difluoro-4-nitrobenzoic acid (23.4 mmol) was solved in 75 mL methanol, before 8.30 g potassium hydroxide was added. The reaction was stirred at room temperature for 3 h. The solvent was removed in vacuo and the pH was adjusted to pH1 with 1 M aq. HCl. The reaction mixture was extracted with EA. The **general procedure D** was conducted with the obtained yellow solid (carboxylic acid) and 2.41 mL morpholine (amine, 27.9 mmol). The solvent was removed, the residue was resolved in EA and washed with 1 M aq. $NaHCO_3$ and 1 M aq. HCl. The obtained orange resin was solved in 45 mL methanol and 5 mL conc. aq. $NH_4Cl$ and 13.0 g iron powder was added. The reaction was heated to 80 °C and stirred for 1 h. The reaction mixture was filtrated through Celite®, the solvent was removed in vacuo, water was added and the reaction mixture was extracted with EA. The crude product was purified via flash chromatography (DCM/MeOH) and RP flash chromatography. 2.91 g of **78a** (11.4 mmol, 49 % yield) was obtained as a white foam.
**MS (ESI+):** m/z = 255.05 $[M+H]^+$
$^1$**H NMR** (250 MHz, DMSO-$d_6$): δ = 6.72 (d, *J* = 6.3 Hz, 1H), 6.40 (d, *J* = 11.6 Hz, 1H), 5.41 (s, 2H), 3.75 (s, 3H), 3.61 – 3.51 (m, 8H) ppm.

**2-methoxy-4-(methylsulfonyl)aniline (78b)**

5.00 g 4-fluoro-2-methoxy-1-nitrobenzene (29.2 mmol) and 3.28 g 4-fluoro-2-methoxy-1-nitrobenzene (32.1 mmol) were solved in 15 mL anhydrous DMA. The reaction was heated to 80 °C and stirred overnight. Water was added and the precipitate was filtrated and washed with water. The obtained off white solid and 7.12 g iron powder were suspended in 90 mL methanol and 10 mL aq. $NH_4Cl$. The reaction was heated to 60 °C and stirred for 1 h. The reaction mixture was filtrated through Celite®, the solvent was removed in vacuo, water was added and the reaction mixture was extracted with EA. 4.88 g of **78b** (24.3 mmol, 83 % yield) was obtained as a pale violet solid.
**MS (ESI+):** m/z = 202.00 $[M+H]^+$
$^1$**H NMR** (250 MHz, $CDCl_3$): δ = 7.35 (dd, *J* = 8.2, 2.0 Hz, 1H), 7.24 (d, *J* = 1.9 Hz, 1H), 6.72 (d, *J* = 8.2 Hz, 1H), 4.27 (s, 2H), 3.89 (s, 3H), 3.00 (s, 3H) ppm.

**(4-((5-chloro-4-(dimethylamino)-7-((2-(trimethylsilyl)ethoxy)methyl)-7*H*-pyrrolo[2,3-*d*]pyrimidin-2-yl)amino)-2-fluoro-5-methoxyphenyl)(morpholino)methanone (79a)**

2.00 g **77a** (5.53 mmol), 1.55 g **78a** (6.09 mmol), 3.52 g $K_3PO_4$ (16.6 mmol), 132 mg XPhos (277 µmol) and 218 mg XPhos Pd G2 (277 µmol) were suspended in 30 mL anhydrous dioxane. The reaction was heated to 60 °C and stirred for 1 h. Water was added and the reaction mixture was extracted with DCM. The crude product was purified via flash chromatography (*n*-hexane/EA). 1.14 g of **79a** (1.97 mmol, 35 % yield) was obtained as a pale yellow solid.
**MS (ESI+):** m/z = 567.25 $[M+H]^+$

**5-chloro-N2-(2-methoxy-4-(methylsulfonyl)phenyl)-N4,N4-dimethyl-7-((2-(trimethylsilyl)ethoxy)methyl)-7*H*-pyrrolo[2,3-*d*]pyrimidine-2,4-diamine (79b)**

**79b** was synthesized according to the procedure of **79a** from 1.97 g **77a** (5.44 mmol) and 1.20 g **78b** (5.98 mmol). 1.24 g **79b** (2.36 mmol, 43 % yield) was obtained as a pale orange solid.
**MS (ESI+):** m/z = 526.15 $[M+H]^+$
$^1$**H NMR** (250 MHz, DMSO-$d_6$): δ = 8.82 (d, *J* = 8.6 Hz, 1H), 7.71 (s, 1H), 7.52 (dd, *J* = 8.6, 2.0 Hz, 1H), 7.44 (d, *J* = 2.0 Hz, 1H), 7.38 (s, 1H), 5.45 (s, 2H), 4.00 (s, 3H), 3.60 – 3.50 (m, 2H), 3.21 (s, 6H), 3.18 (s, 3H), 0.90 – 0.80 (m, 2H), -0.13 (s, 9H) ppm.

**5-iodo-N2-(2-methoxy-4-(methylsulfonyl)phenyl)-N4,N4-dimethyl-7-((2-(trimethylsilyl)ethoxy)methyl)-7*H*-pyrrolo[2,3-*d*]pyrimidine-2,4-diamine (79c)**

1.50 g **77b** (4.59 mmol), 1.02 g **78b** (5.05 mmol), 2.92 g $K_3PO_4$ (13.8 mmol), 109 mg XPhos (229 µmol) and 180 mg XPhos Pd G2 (229 µmol) were suspended in 30 mL *iso*-butanol. The reaction was heated to 80 °C and stirred for 1 h. Water was added and the reaction mixture was extracted with DCM. The combined organic layers were washed with 1 M aq. HCl and with 1 M aq. $NaHCO_3$. The crude product was purified via flash chromatography (*n*-hexane/EA). 1.14 g of **79c** (1.97 mmol, 35 % yield) was obtained as a pale yellow solid.
**MS (ESI+):** m/z = 618.10 $[M+H]^+$
$^1$**H NMR** (250 MHz, DMSO-$d_6$): δ = 8.89 – 8.82 (m, 1H), 7.67 (s, 1H), 7.52 (dd, *J* = 8.6, 2.0 Hz, 1H), 7.43 (d, *J* = 2.0 Hz, 1H), 6.96 (s, 1H), 5.46 (s, 2H), 4.00 (s, 3H), 3.61 – 3.51 (m, 2H), 3.28 (s, 6H), 3.17 (s, 3H), 0.92 – 0.82 (m, 3H), -0.13 (s, 9H) ppm.

**(4-((5-(3-aminophenyl)-4-(dimethylamino)-7-((2-(trimethylsilyl)ethoxy)methyl)-7*H*-pyrrolo[2,3-*d*]pyrimidin-2-yl)amino)-2-fluoro-5-methoxyphenyl)(morpholino)methanone (80a)**

**80a** was synthesized according to the **general procedure A** from the aryl halide **79a** (540 mg, 932 µmol) and (3-aminophenyl)boronic acid (140 mg, 1.03 mmol). Water was added and the reaction mixture was extracted with EA. The crude product was purified via flash chromatography (DCM/MeOH). 556 mg of **80a** (874 µmol, 93 % yield) was obtained as an orange resin.
**MS (ESI+):** m/z = 636.45 $[M+H]^+$

**5-(3-aminophenyl)-N2-(2-methoxy-4-(methylsulfonyl)phenyl)-N4,N4-dimethyl-7-((2-(trimethylsilyl)ethoxy)methyl)-7*H*-pyrrolo[2,3-*d*]pyrimidine-2,4-diamine (80b)**

**80b** was synthesized according to the **general procedure A** from the aryl halide **79c** (600 mg, 972 µmol) and (3-aminophenyl)boronic acid (146 mg, 1.07 mmol). Water was added and the reaction mixture was extracted with EA. 560 mg of **80b** (961 µmol, 99 % yield) was obtained as a light brown solid.
**MS (ESI+):** m/z = 583.25 $[M+H]^+$

***N*-(4-fluorophenyl)-*N*-(4-hydroxyphenyl)cyclopropane-1,1-dicarboxamide (81a)**

To a solution of 2.50 g **62** (11.2 mmol) and 1.47 g 4-aminophenol (13.4 mmol) in 30 mL anhydrous DMF was added 2.58 g EDC (13.4 mmol). The solution was stirred at room temperature overnight. The solution was added to 100 mL water and extracted with EA. The obtained solid was purified via flash chromatography (*c*-hexane/EA 1:1), affording 2.68 g of **81a** (8.53 mmol, 76 % yield) as a white solid.
**MS (ESI+)**: m/z = 315.14 $[M+H]^+$

$^1$**H NMR** (400 MHz, DMSO-*d$_6$*): δ = 10.15 (s, 1H), 9.71 (s, 1H), 9.20 (s, 1H), 7.61 (dd, *J* = 9.0, 5.1 Hz, 2H), 7.34 (d, *J* = 8.8 Hz, 2H), 7.14 (t, *J* = 8.9 Hz, 2H), 6.71 – 6.66 (m, 2H), 1.44 (d, *J* = 1.6 Hz, 4H) ppm.

**N-(4-aminophenyl)-N-(4-fluorophenyl)cyclopropane-1,1-dicarboxamide (81b)**

**81b** was synthesized according to the procedure of **63** from 1.00 g **62** (4.48 mmol) and 533 mg benzene-1,4-diamine (4.93 mmol). 1.00 g of **81b** (3.19 mmol,71 % yield) was obtained as a white solid.
**MS (ESI+)**: m/z = 314.10 $[M+H]^+$
$^1$**H NMR** (400 MHz, DMSO-*d$_6$*): δ = 10.23 (s, 1H), 9.55 (s, 1H), 7.61 (dd, *J* = 8.7, 4.9 Hz, 2H), 7.30 – 7.02 (m, 4H), 6.49 (d, *J* = 8.6 Hz, 2H), 4.89 (s, 2H), 1.43 (d, *J* = 2.9 Hz, 4H) ppm.

**5-bromo-2-((2-(trimethylsilyl)ethoxy)methyl)-2*H*-indazole (83)**

1.00 g 5-bromo-1*H*-indazole (5.08 mmol) and 1.30 mL (6.09 mmol) *N*-cyclohexyl-*N*-methylcyclohexanamine were solved in 25 mL anhydrous THF. 1.08 mL (2-(chloromethoxy)ethyl)trimethylsilane (6.09 mmol) were added dropwise, before the mixture was stirred at room temperature overnight. Water was added and the reaction mixture was extracted with DCM. The crude product was purified via flash chromatography (*n*-hexane/EA). 970 mg of **83** (2.97 mmol, 58 % yield) was obtained as a yellow oil.
**MS (ESI+)**: m/z = 327.00 $[M+H]^+$
$^1$**H NMR** (250 MHz, DMSO-*d$_6$*): δ = 8.06 (d, *J* = 0.9 Hz, 1H), 7.85 (dd, *J* = 1.8, 0.7 Hz, 1H), 7.62 (dt, *J* = 9.2, 0.9 Hz, 1H), 7.35 (dd, *J* = 9.2, 1.8 Hz, 1H), 5.71 (s, 2H), 3.70 – 3.55 (m, 2H), 1.01 – 0.85 (m, 2H), -0.03 (s, 9H) ppm.

**4-methyl-*N*-(3-(4-methyl-1*H*-imidazol-1-yl)-5-(trifluoromethyl)phenyl)-3-nitrobenzamide (86.0)**

826 mg 4-methyl-3-nitrobenzoic acid (4.56 mmol) and 1.89 g HATU (4.97 mmol) were solved in 20 mL anhydrous DMF and 1.08 mL DIPEA (6.22 mmol) was added. After 5 min of stirring 1.00 g 3-(4-methyl-1H-imidazol-1-yl)-5-(trifluoromethyl)aniline (4.15 mmol) was added. The reaction was stirred at room temperature overnight. The solvent was removed and the residue was purified via RP chromatography. 1.51 g of **86.0** (3.37 mmol, 91 %) was obtained as a yellow solid.
**MS (ESI+)**: m/z = 405.15 $[M+H]^+$
$^1$**H NMR** (250 MHz, DMSO-*d$_6$*): δ = 10.84 (s, 1H), 8.61 (d, *J* = 1.8 Hz, 1H), 8.30 – 8.17 (m, 3H), 8.11 (s, 1H), 7.78 – 7.67 (m, 2H), 7.47 (s, 1H), 2.60 (s, 3H), 2.18 (s, 3H) ppm.

**3-amino-4-methyl-*N*-(3-(4-methyl-1*H*-imidazol-1-yl)-5-(trifluoromethyl)phenyl)benzamide (86)**

1.50 g **86.0** (3.71 mmol) was solved in 15 mL anhydrous methanol. The flask was flushed with argon, before 150 mg palladium on carbon was added. The suspension was stirred under hydrogen atmosphere at room temperature overnight. The reaction mixture was filtrated through Celite® and the solvent was removed. 1.26 g of **86** (3.73 mmol, 90 %) was obtained as a yellowish solid.
**MS (ESI+)**: m/z = 375.10 $[M+H]^+$
$^1$**H NMR** (500 MHz, DMSO-*d$_6$*): δ = 10.44 (s, 1H), 8.28 (s, 1H), 8.18 (d, *J* = 1.4 Hz, 1H), 8.15 (s, 1H), 7.68 (s, 1H), 7.48 – 7.42 (m, 1H), 7.20 (d, *J* = 1.3 Hz, 1H), 7.16 – 7.05 (m, 2H), 5.13 (s, 2H), 2.18 (d, *J* = 0.8 Hz, 3H), 2.13 (s, 3H) ppm.

### NMR spectra

$^{1}H$ and $^{13}C$ NMR spectra of compound **1**

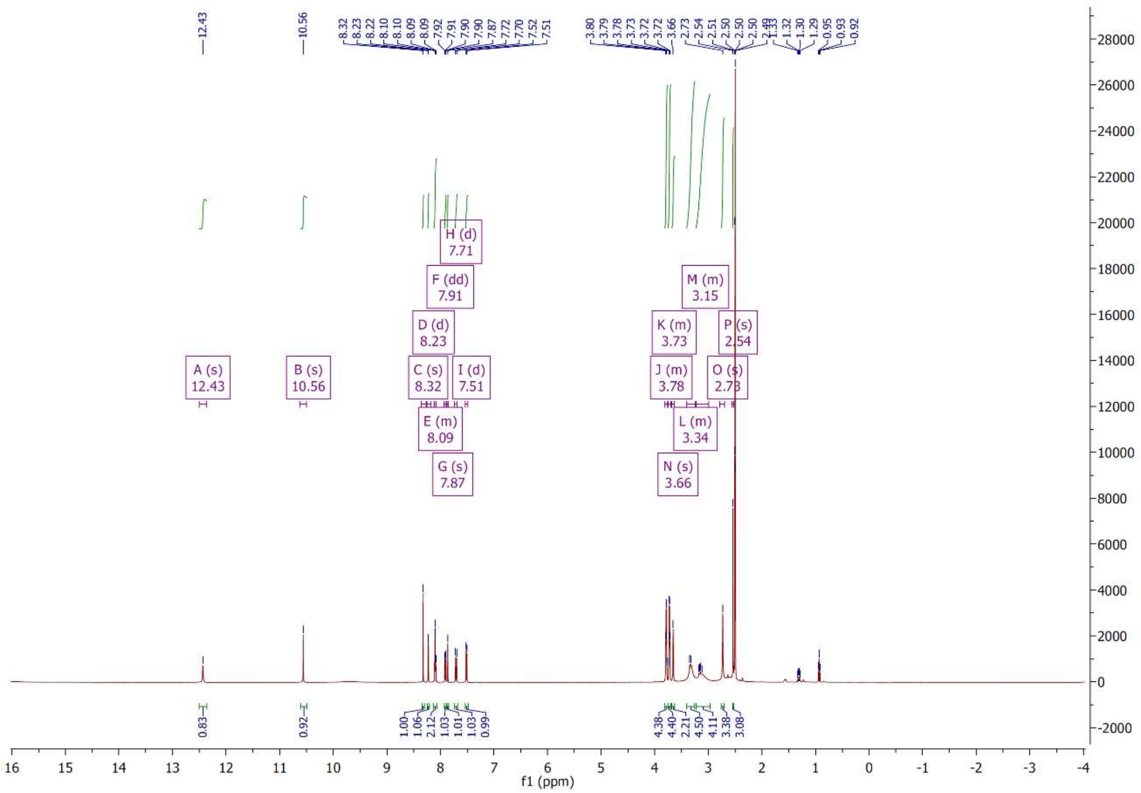

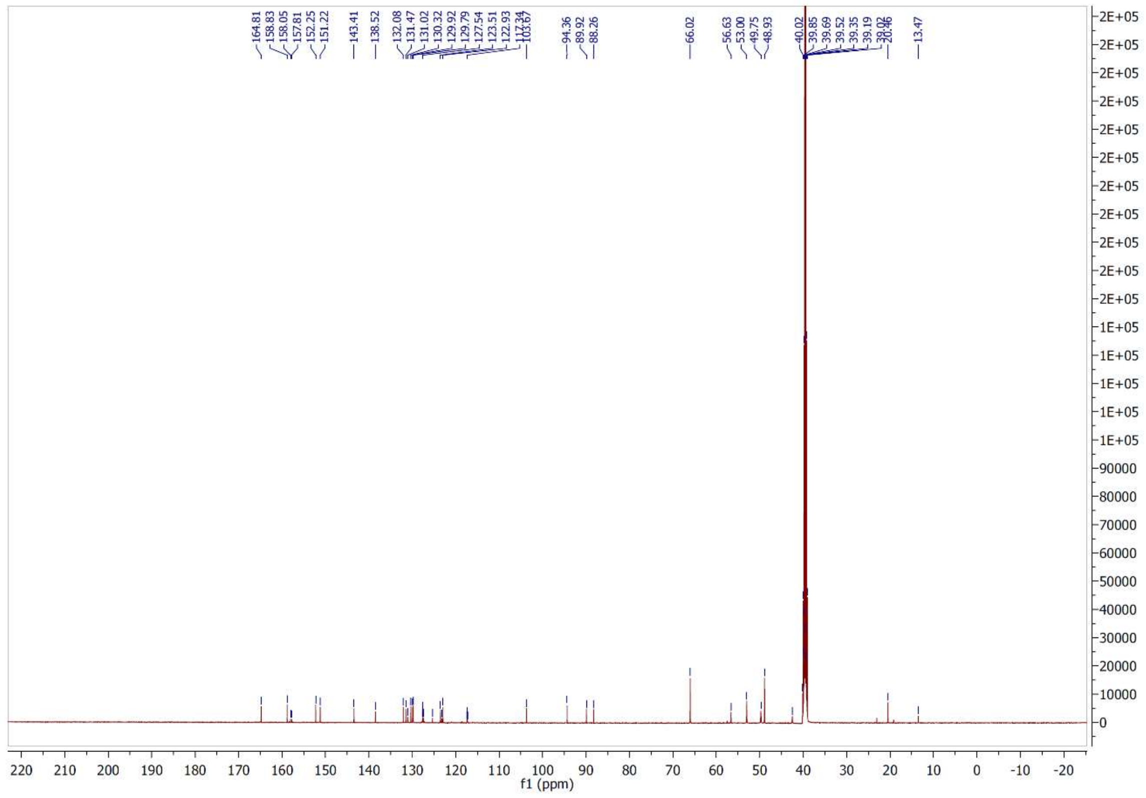

$^{1}H$ and $^{13}C$ NMR spectra of compound **2**

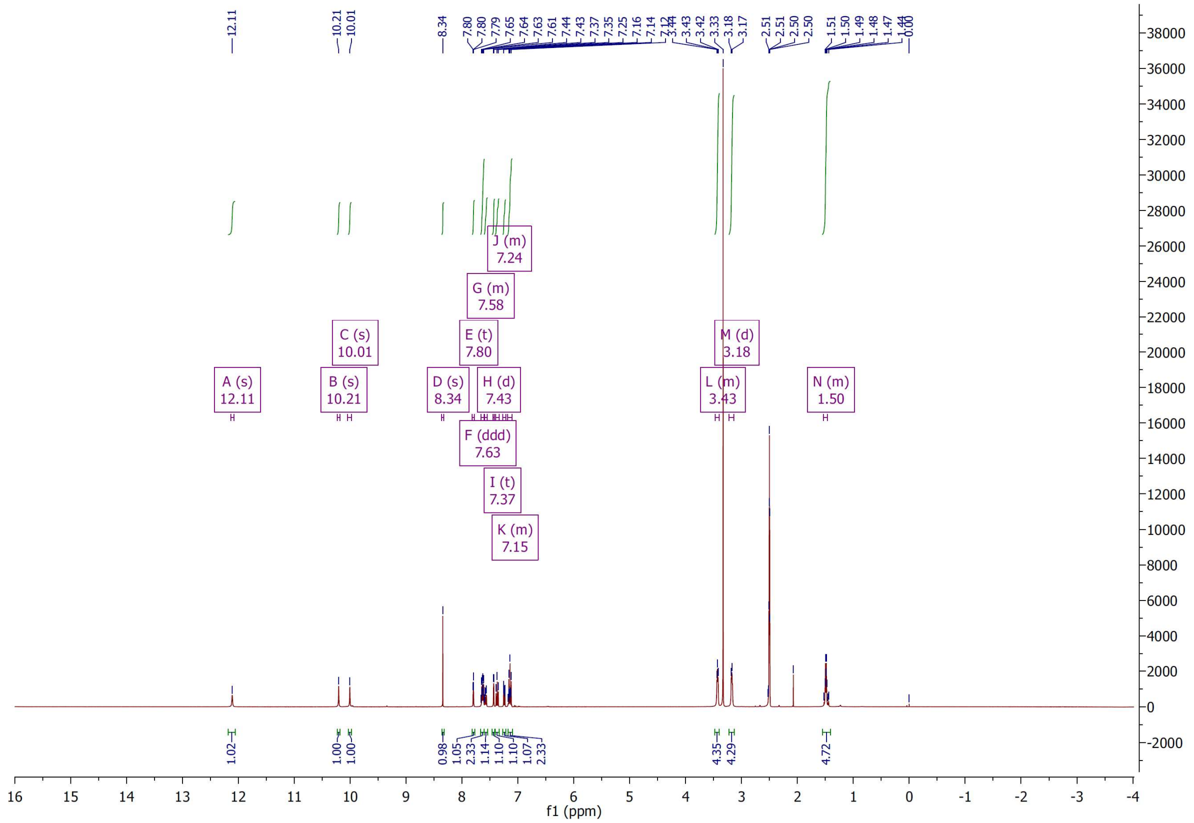

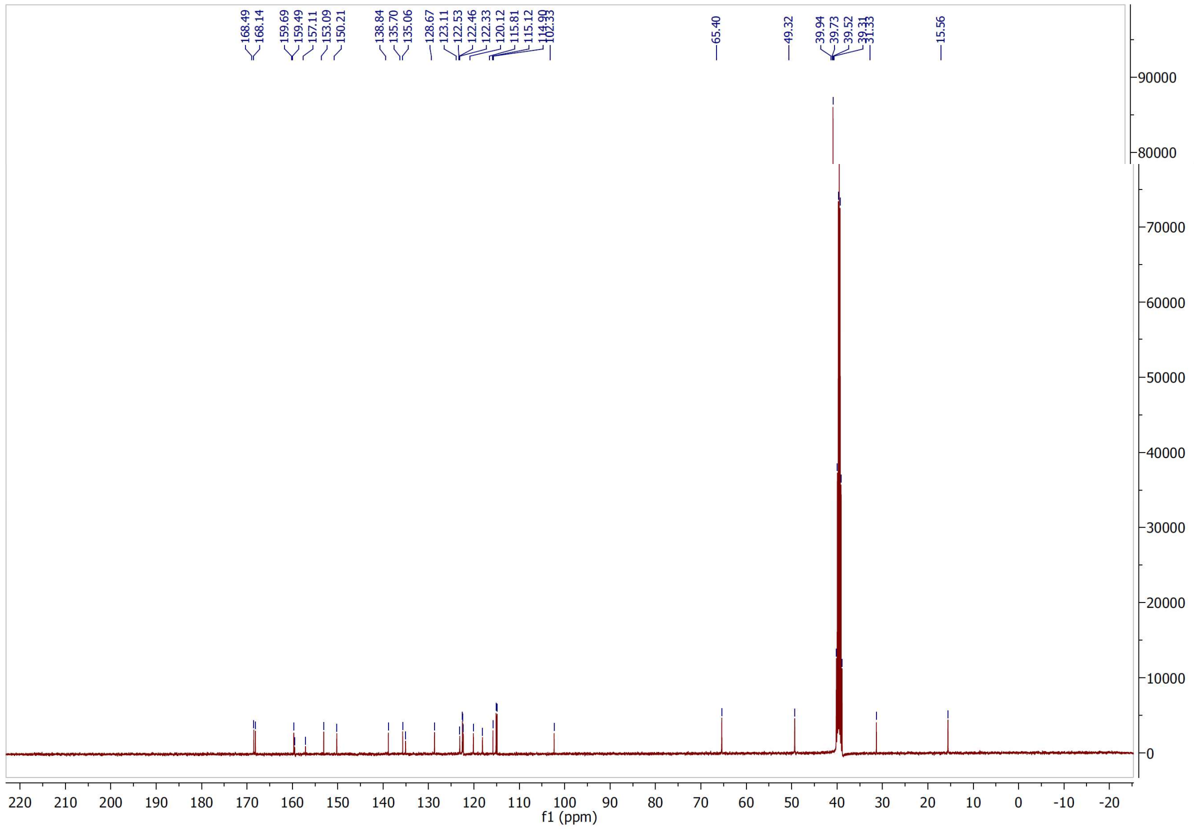

[1]H and [13]C NMR spectra of compound **3**

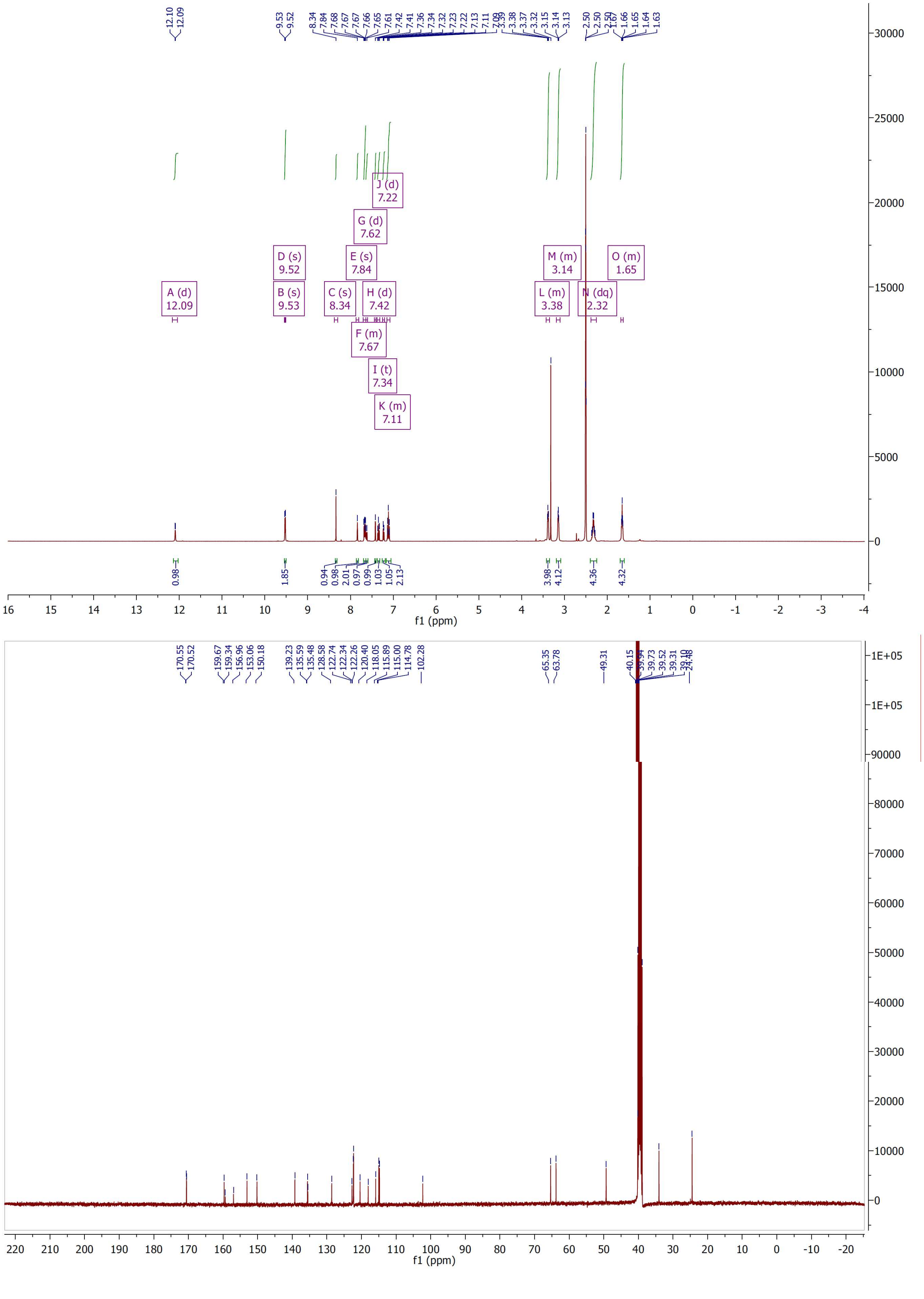

[1]H and [13]C NMR spectra of compound **4**

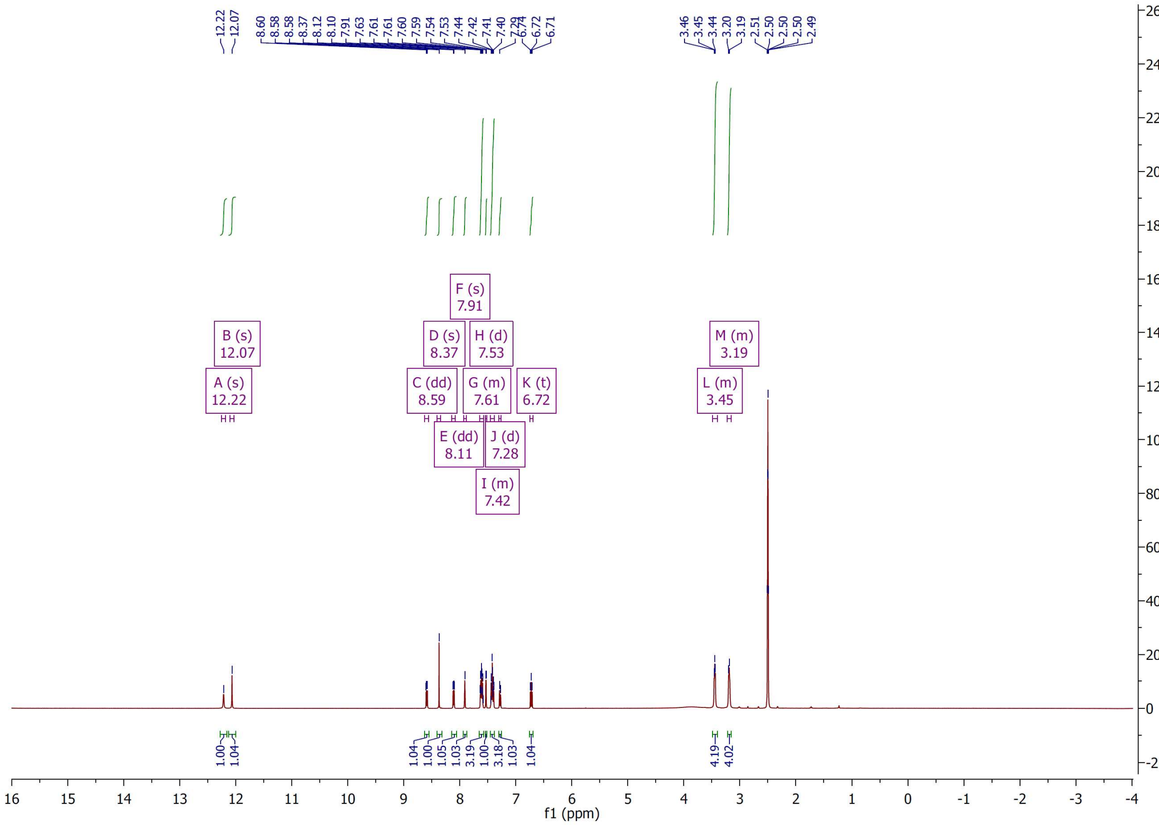


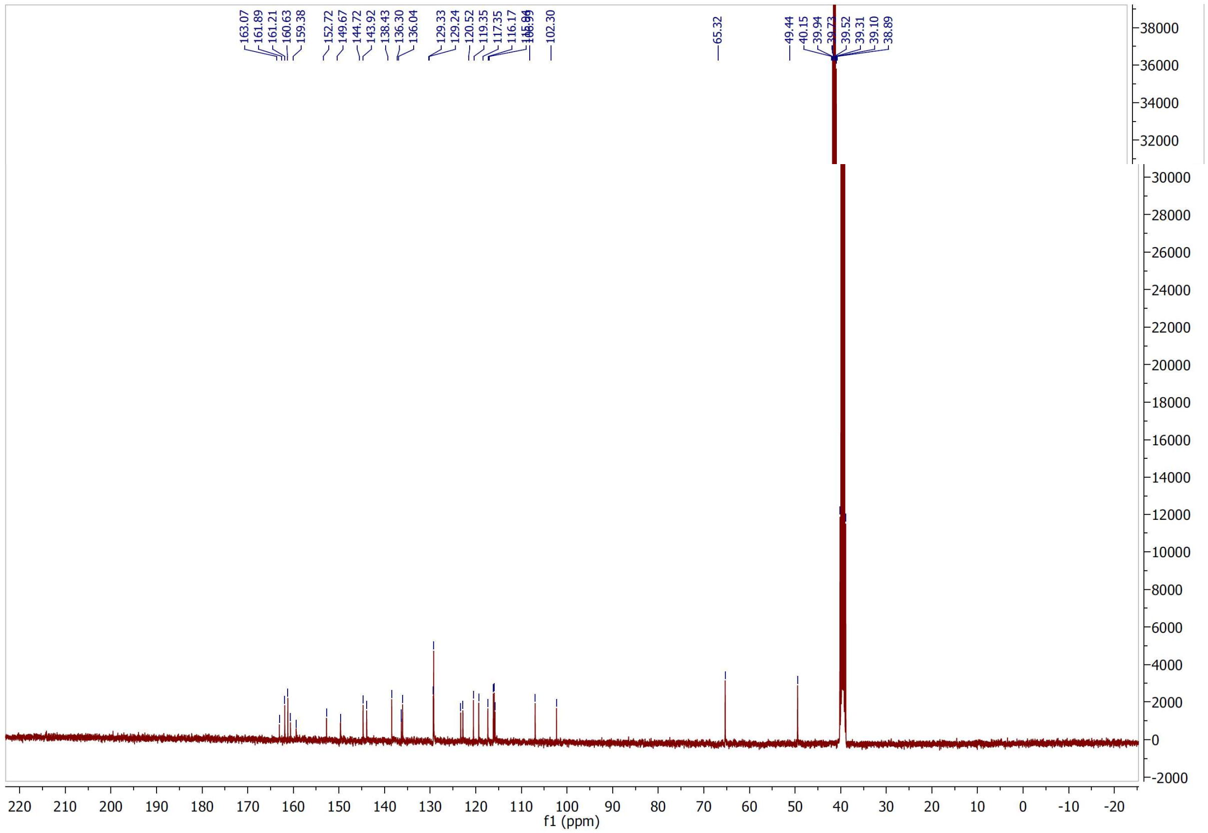

$^{1}H$ and $^{13}C$ NMR spectra of compound **5**

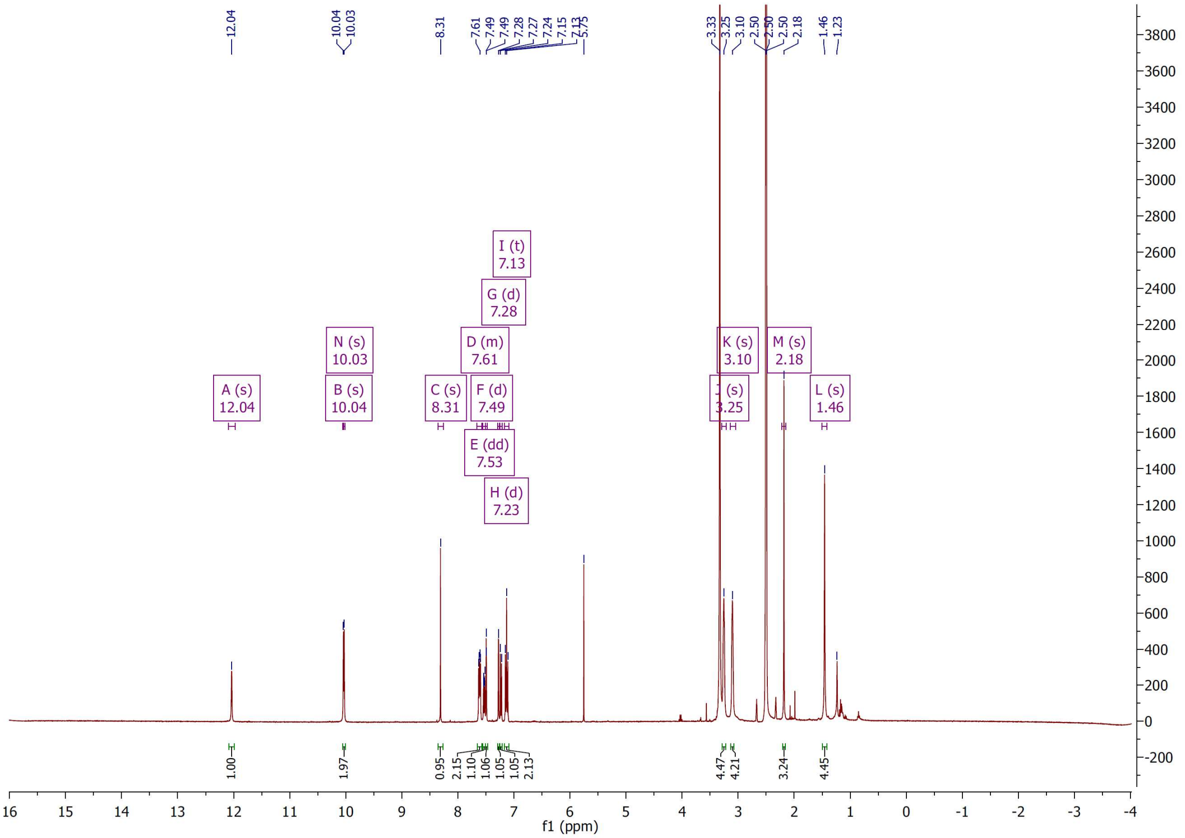

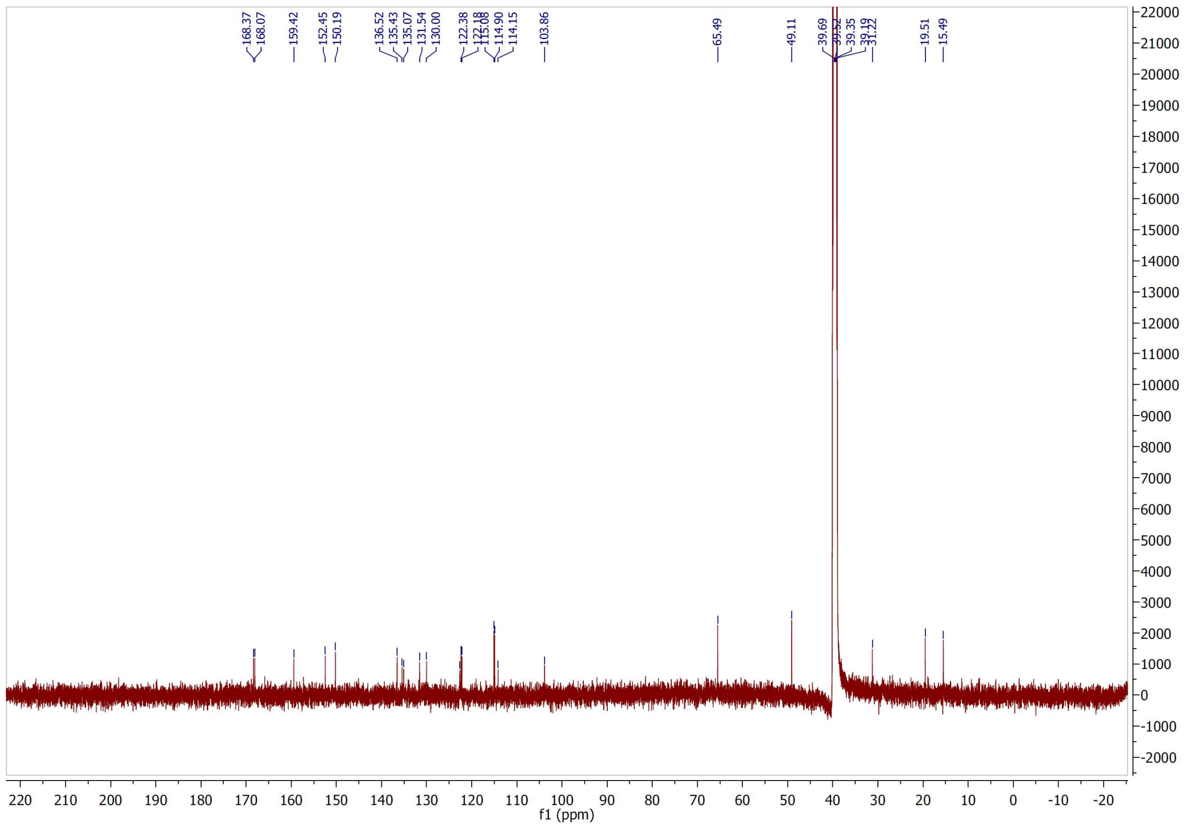

[1]H and [13]C NMR spectra of compound **6**

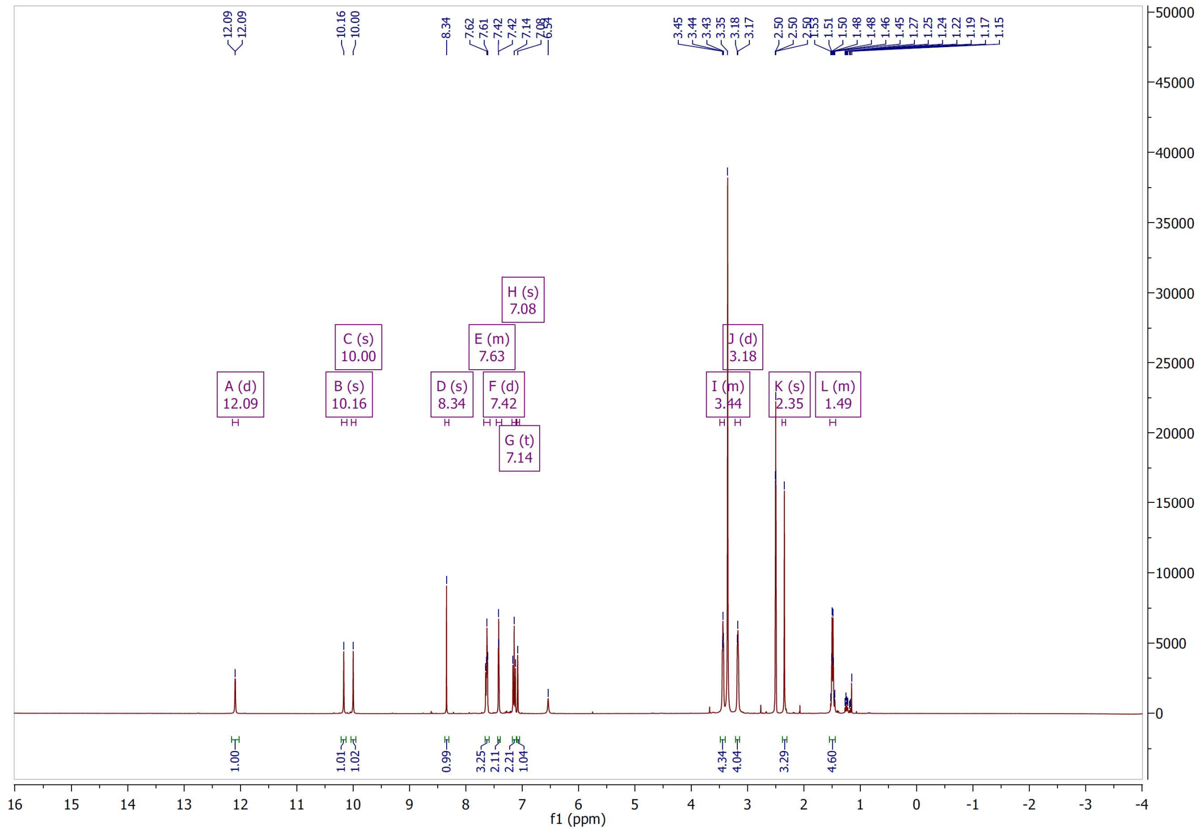

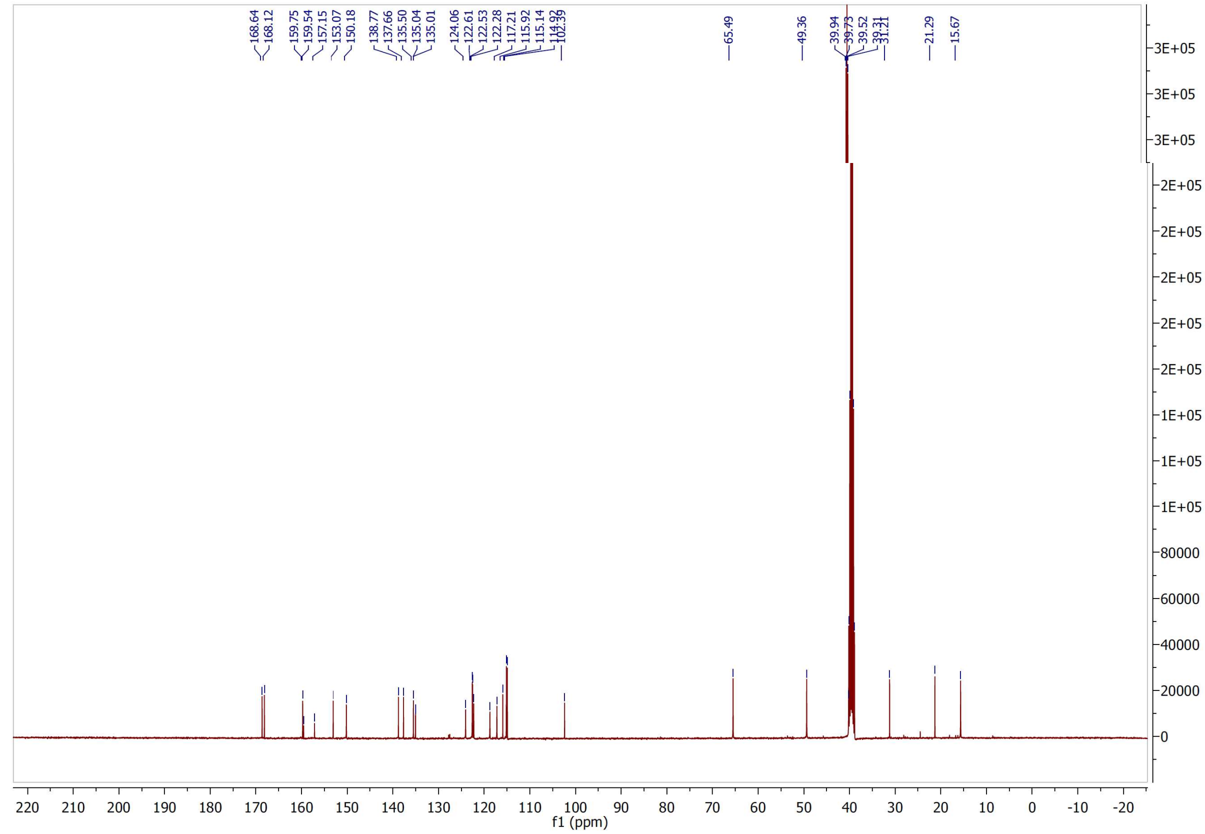

[1]H and [13]C NMR spectra of compound **7**

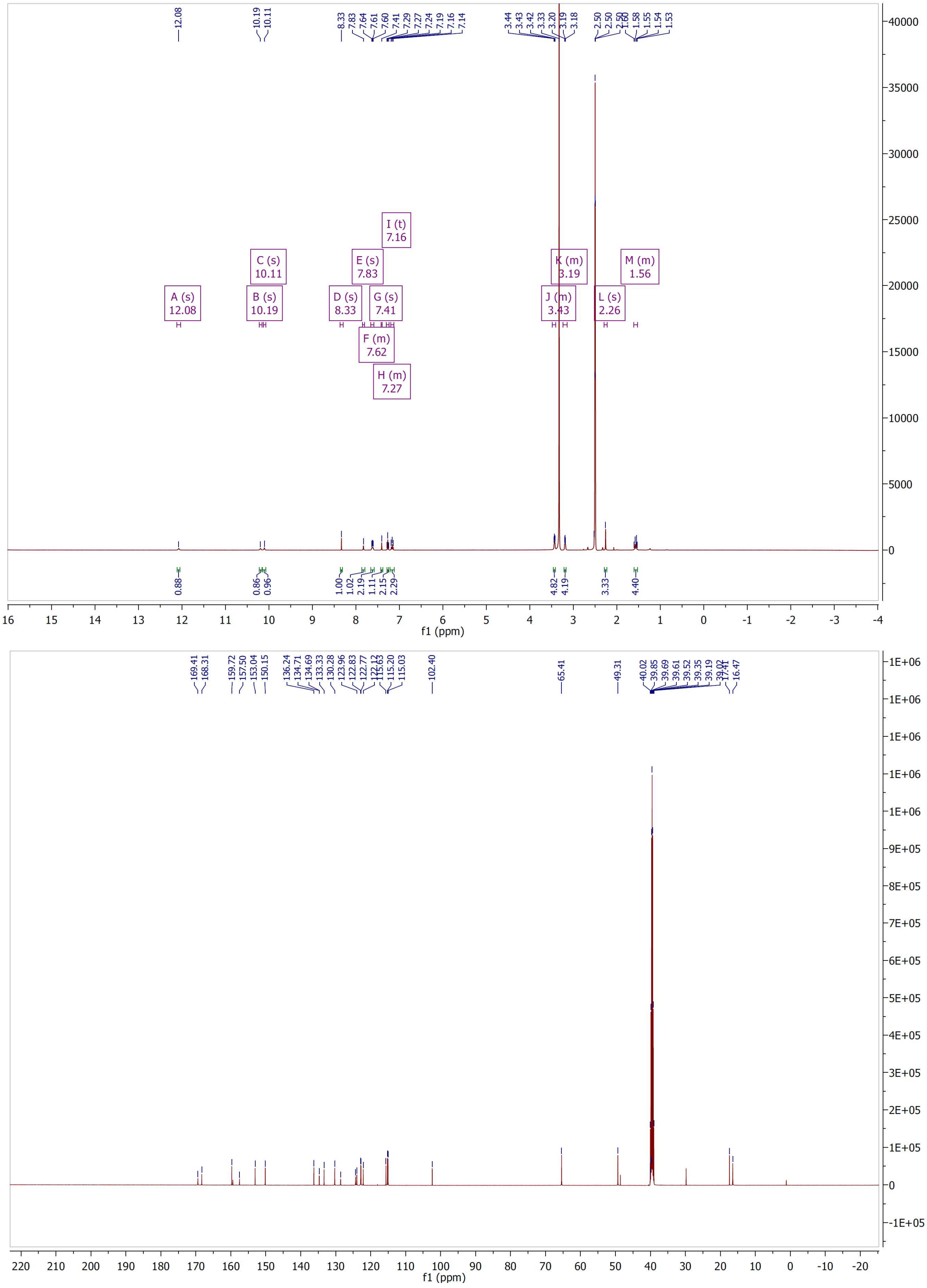

[1]H and [13]C NMR spectra of compound **8**

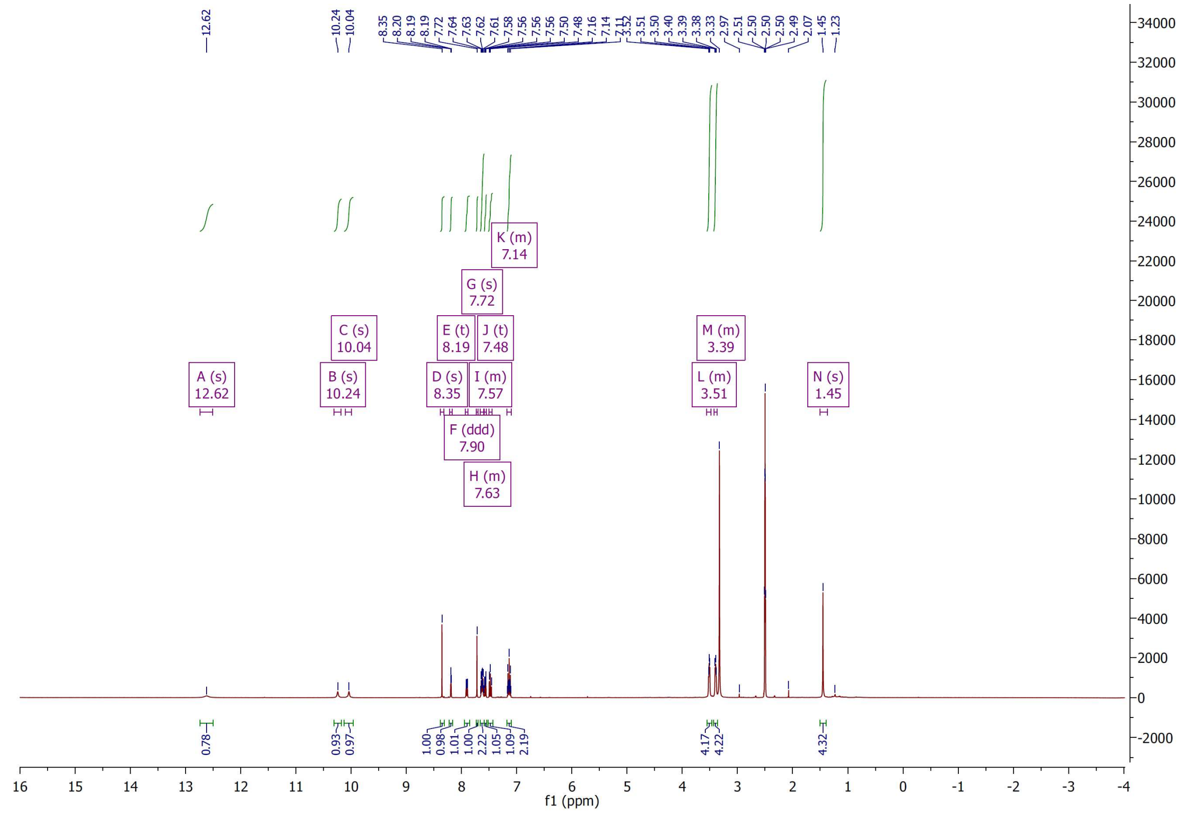

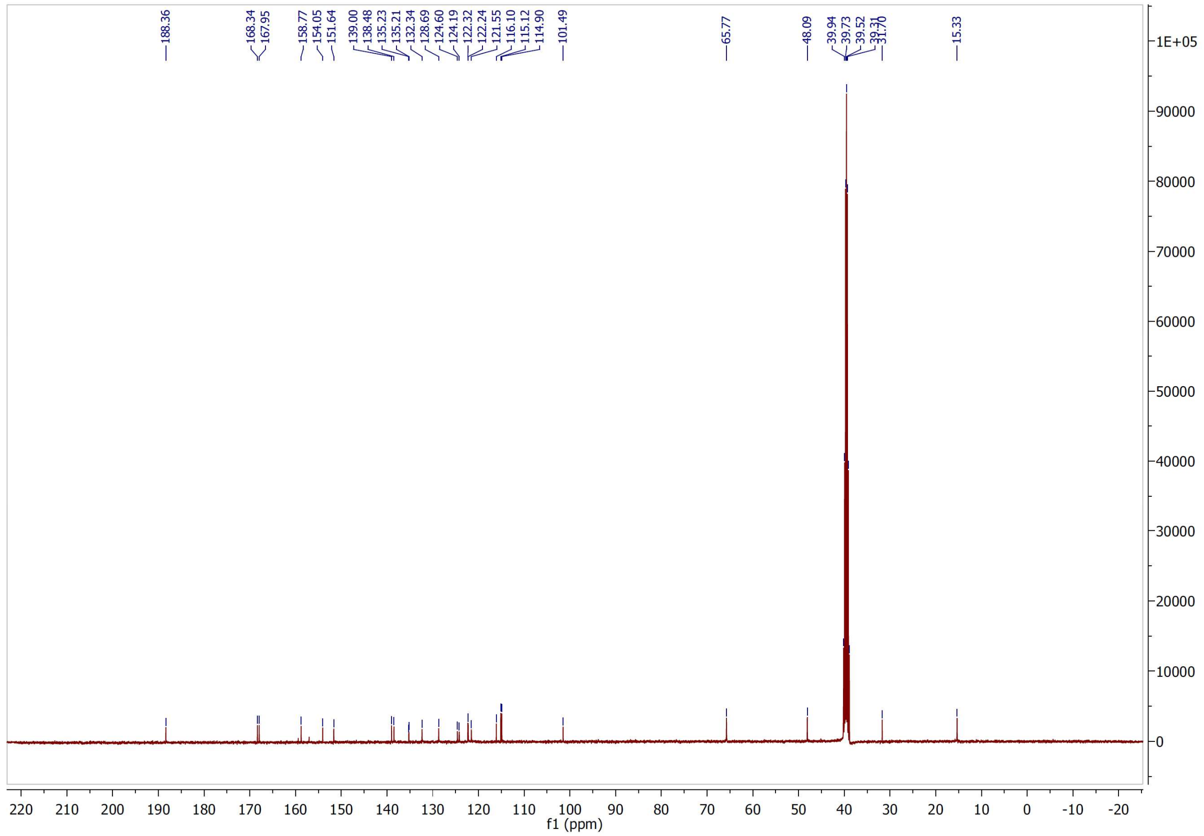

[1]H and [13]C NMR spectra of compound **9**

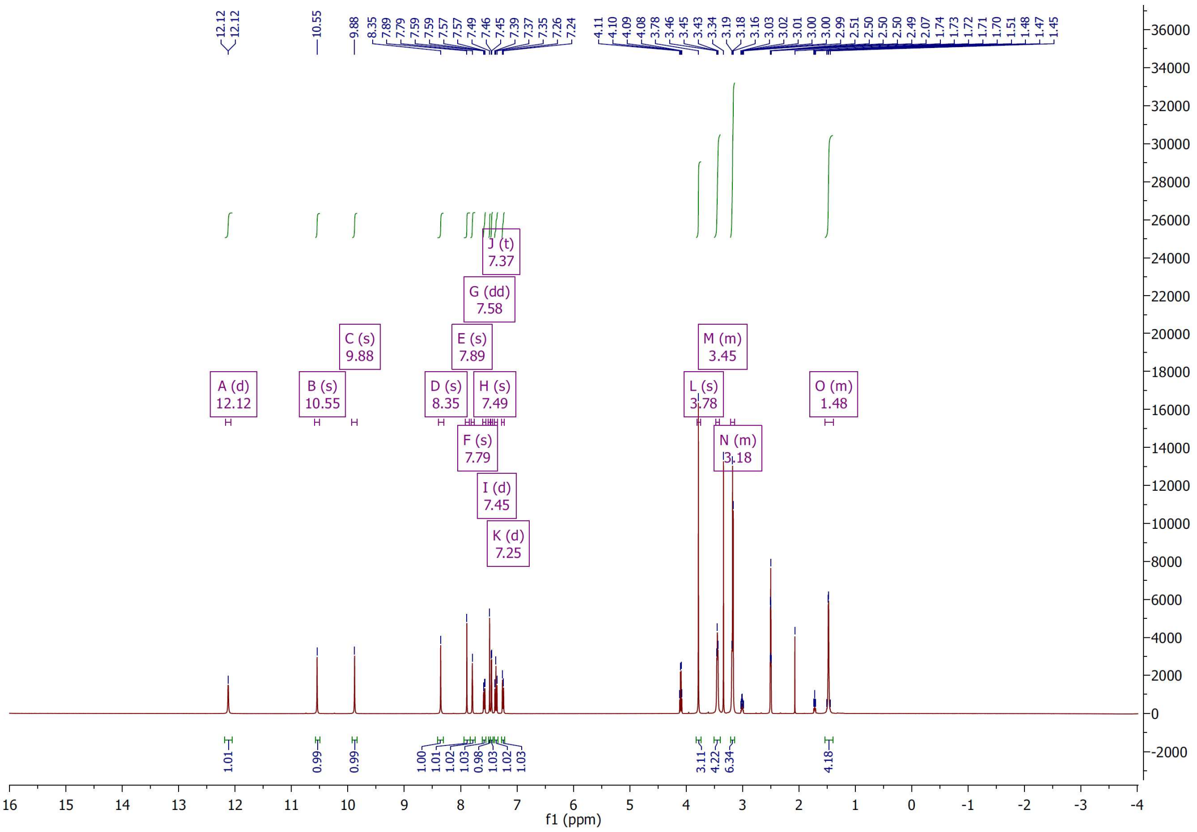

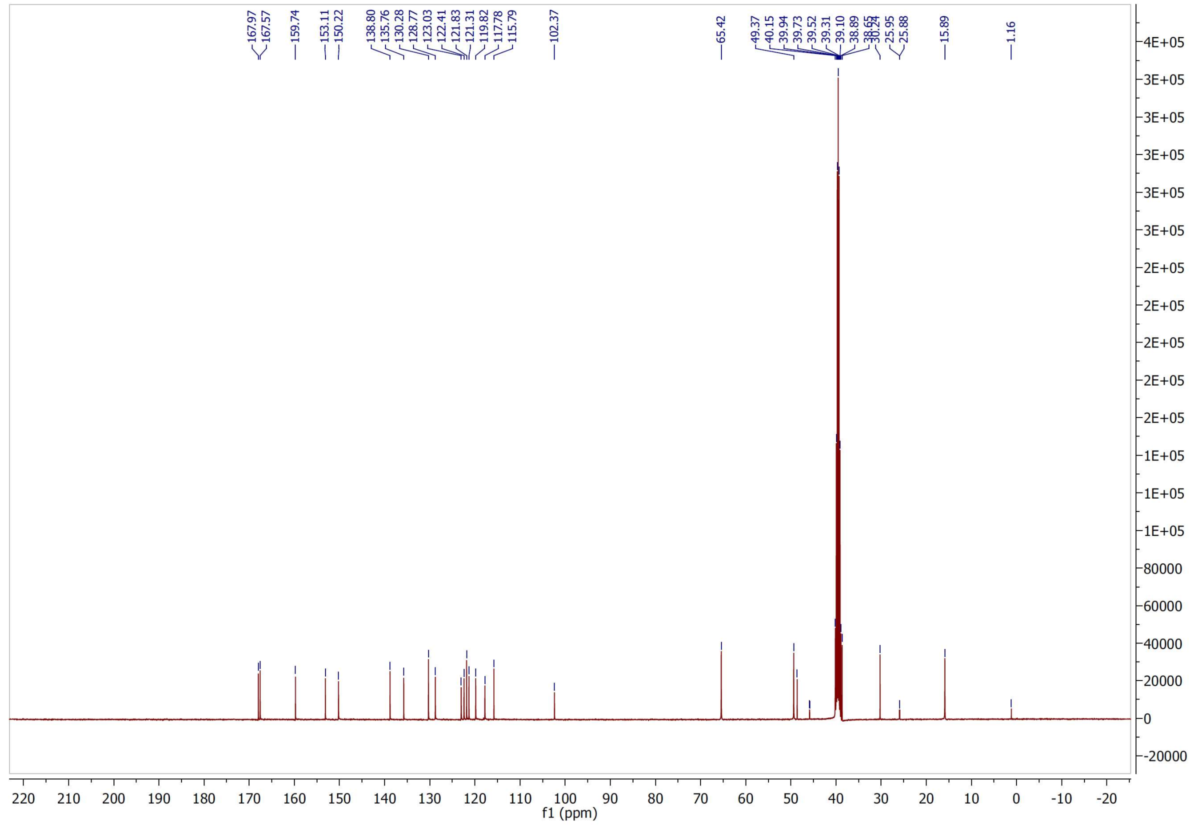

$^{1}H$ and $^{13}C$ NMR spectra of compound **10**

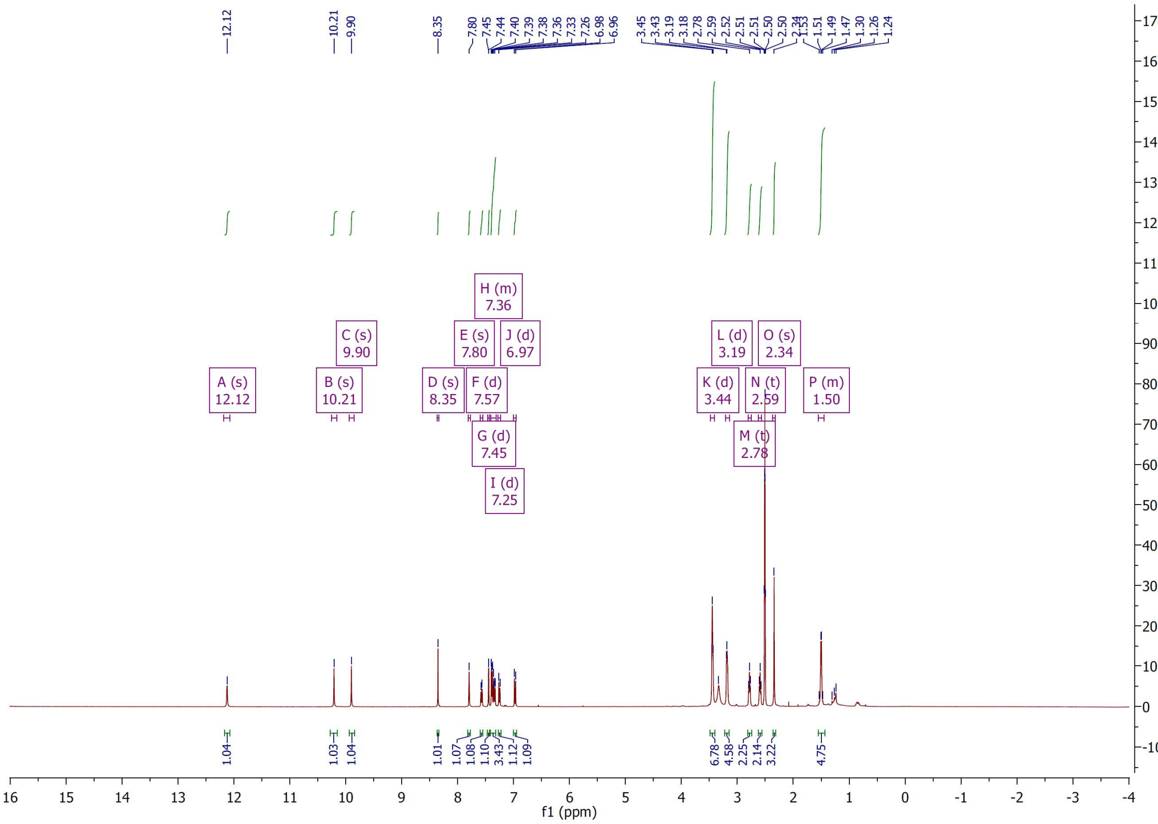


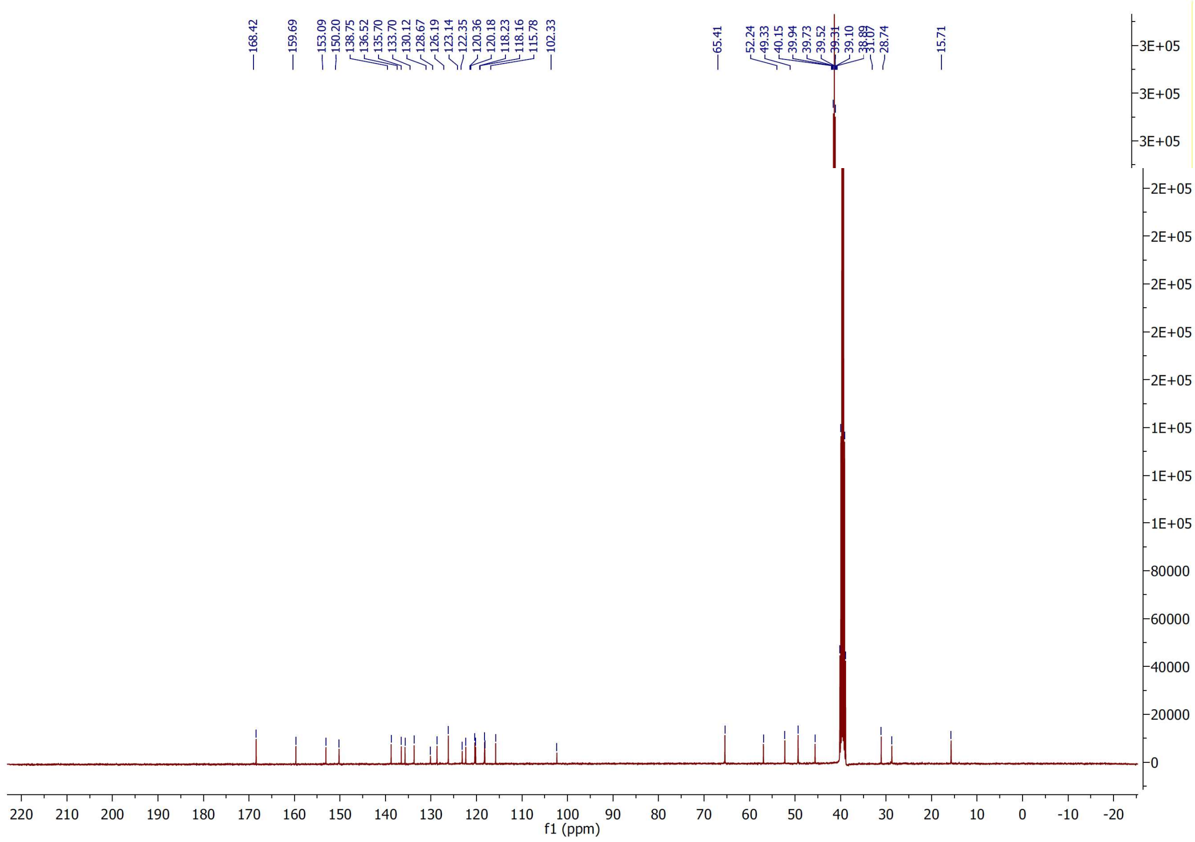

[1]H and [13]C NMR spectra of compound **11**

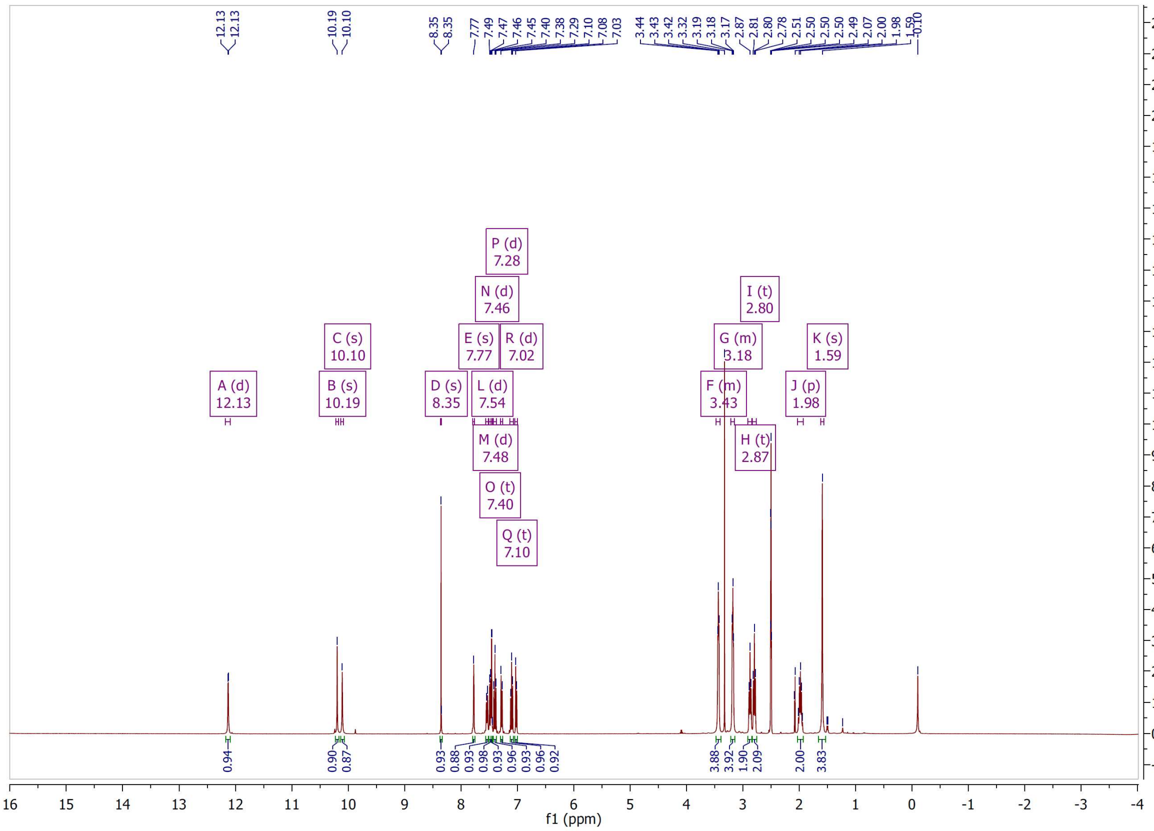


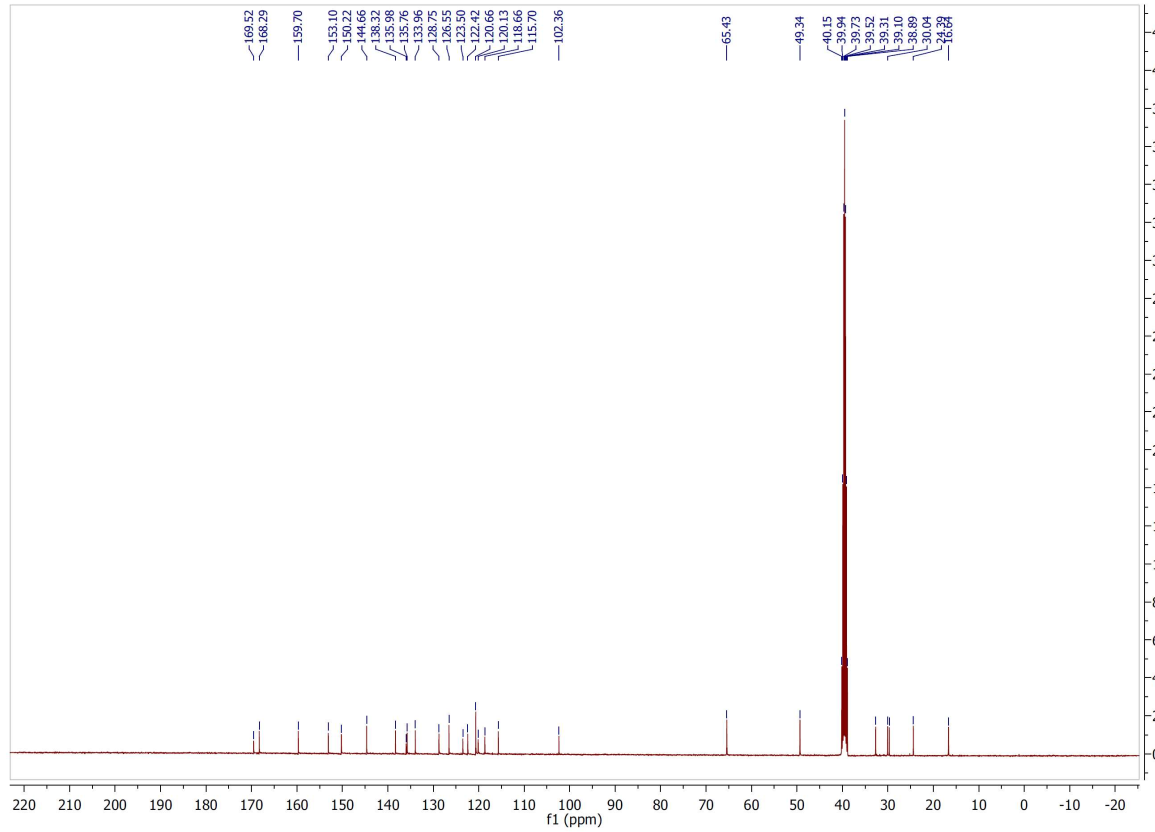

$^{1}H$ and $^{13}C$ NMR spectra of compound **12**

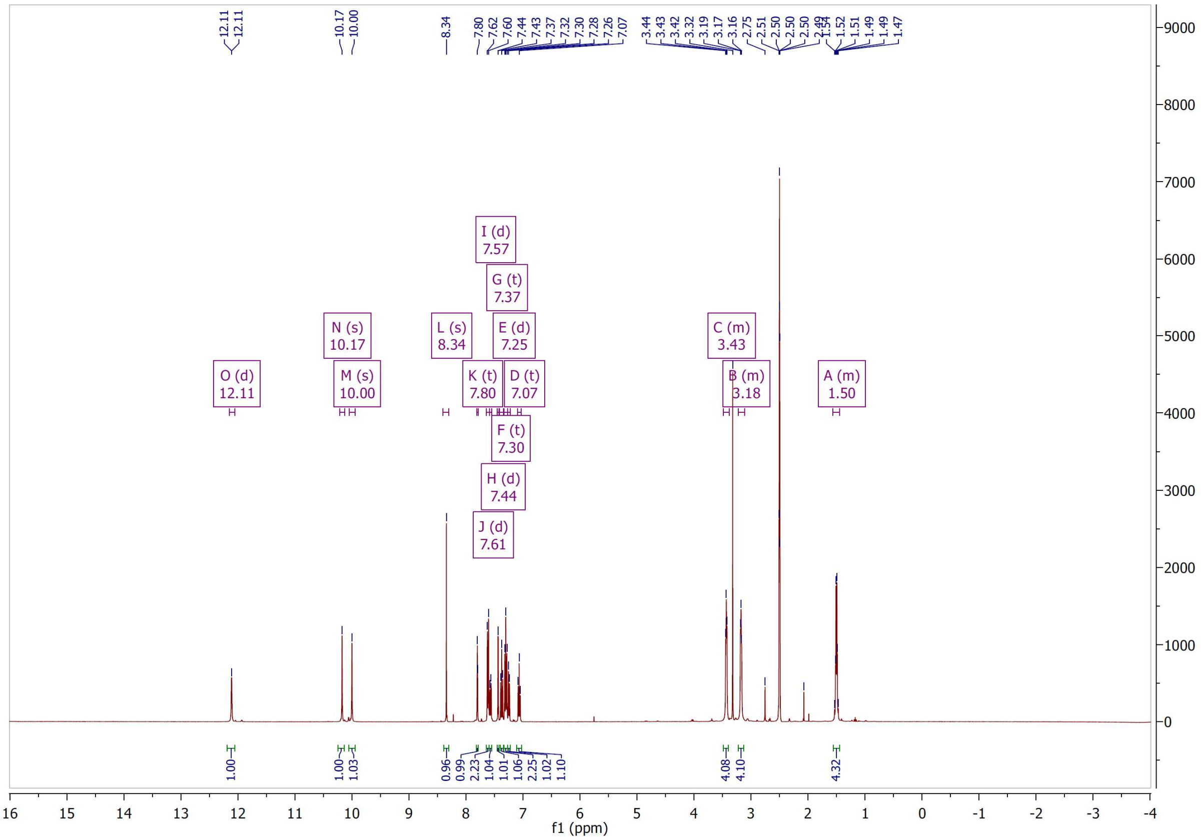


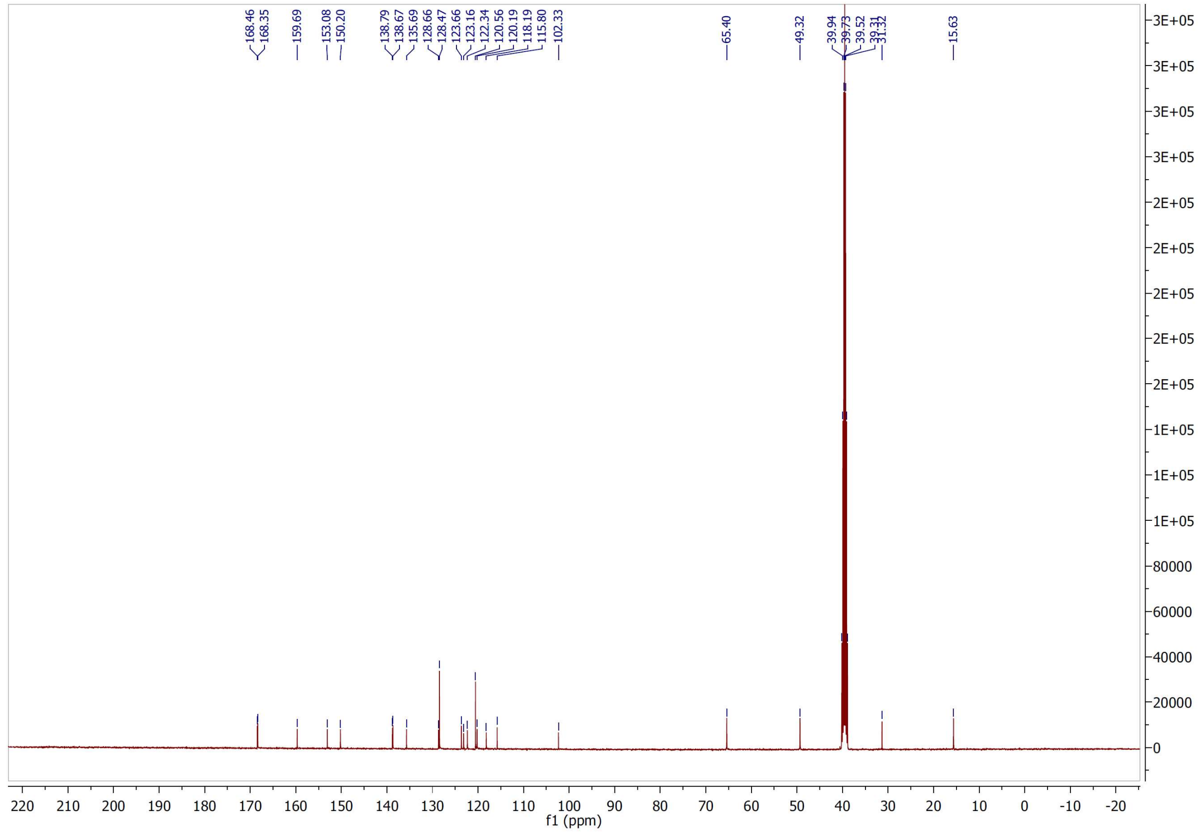

$^{1}H$ and $^{13}C$ NMR spectra of compound **13**

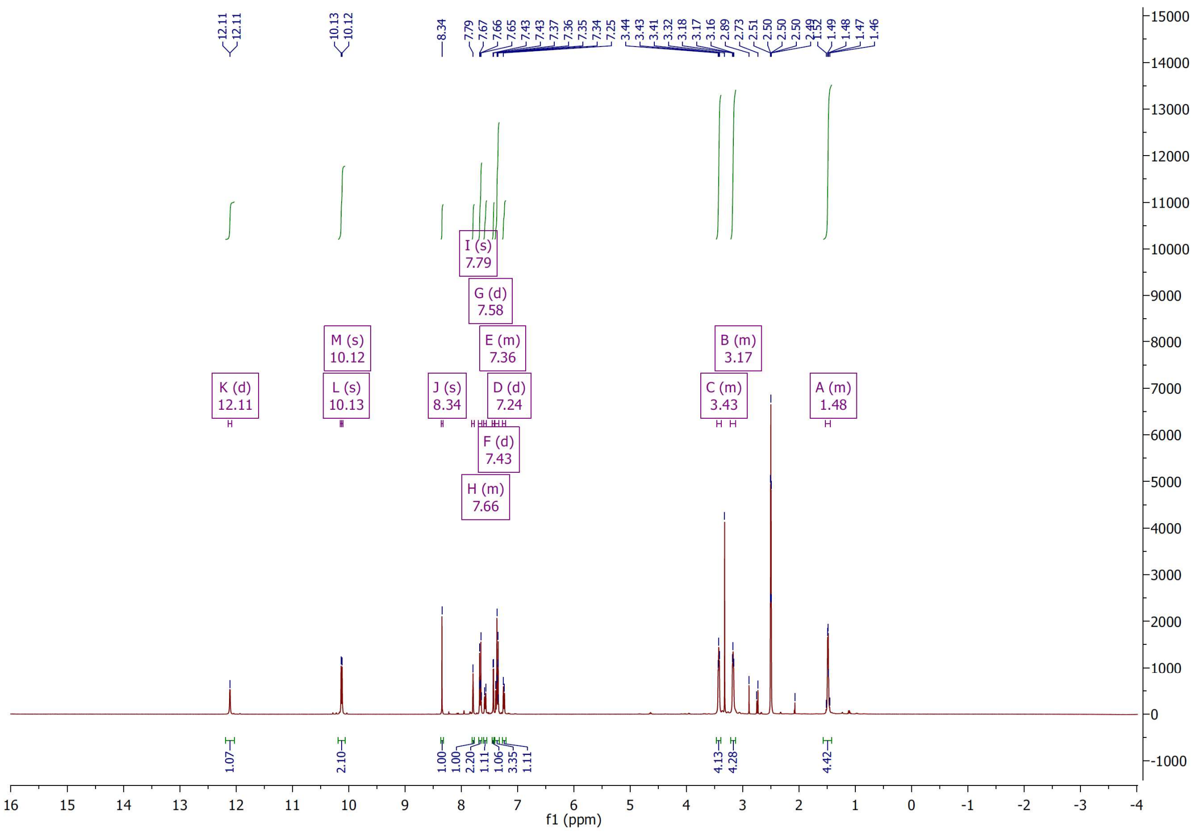


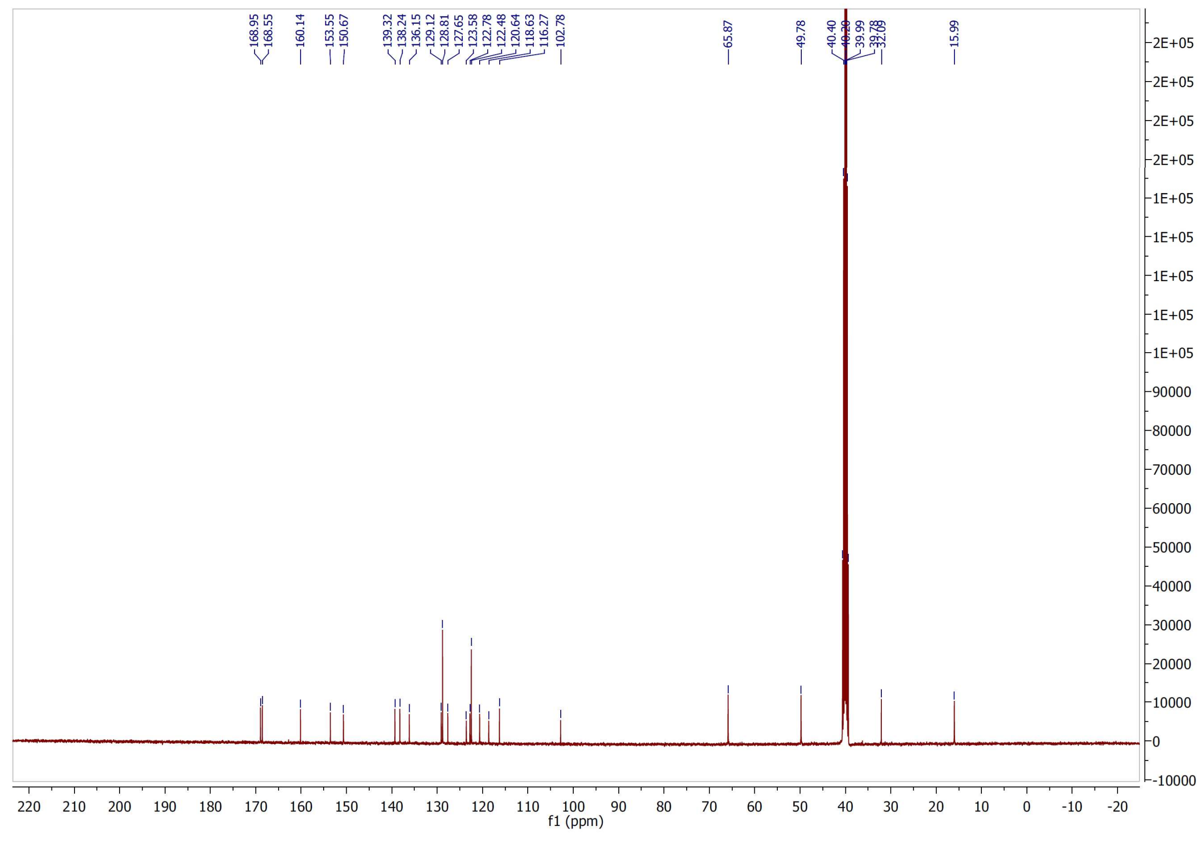

$^{1}H$ and $^{13}C$ NMR spectra of compound **14**

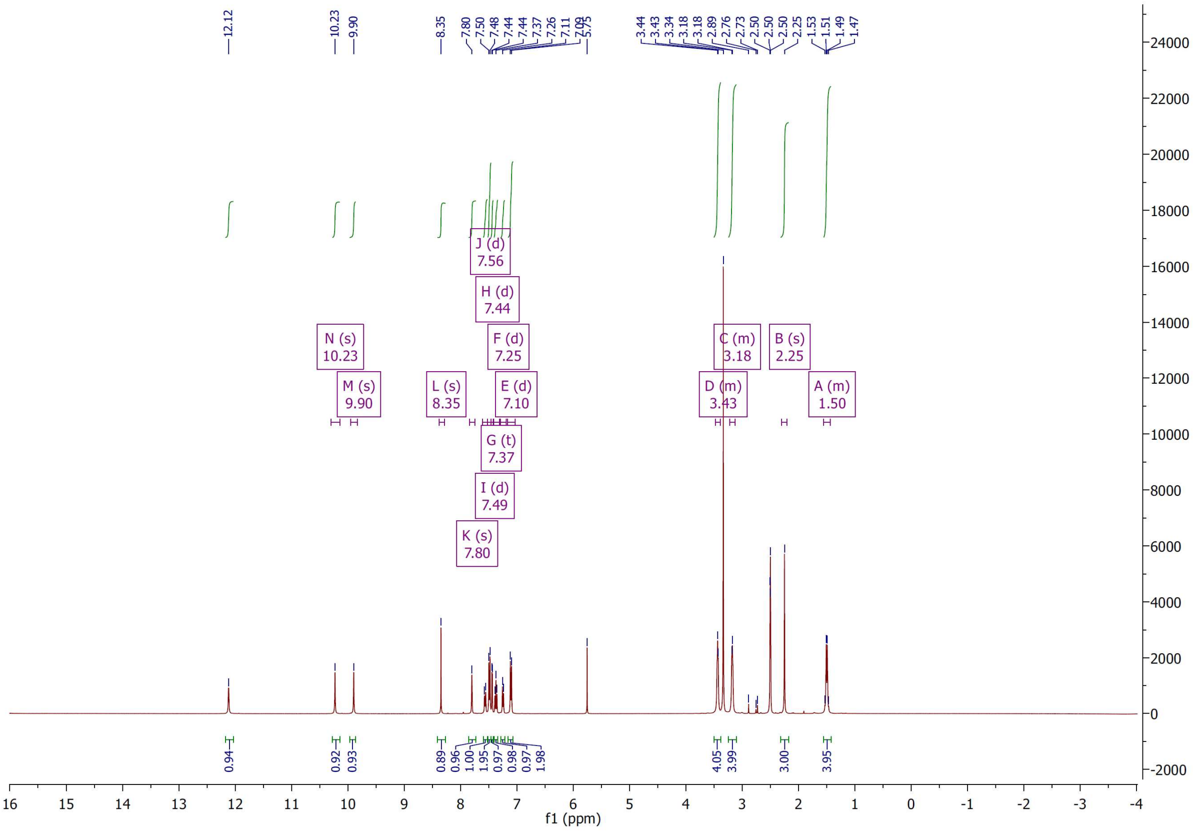


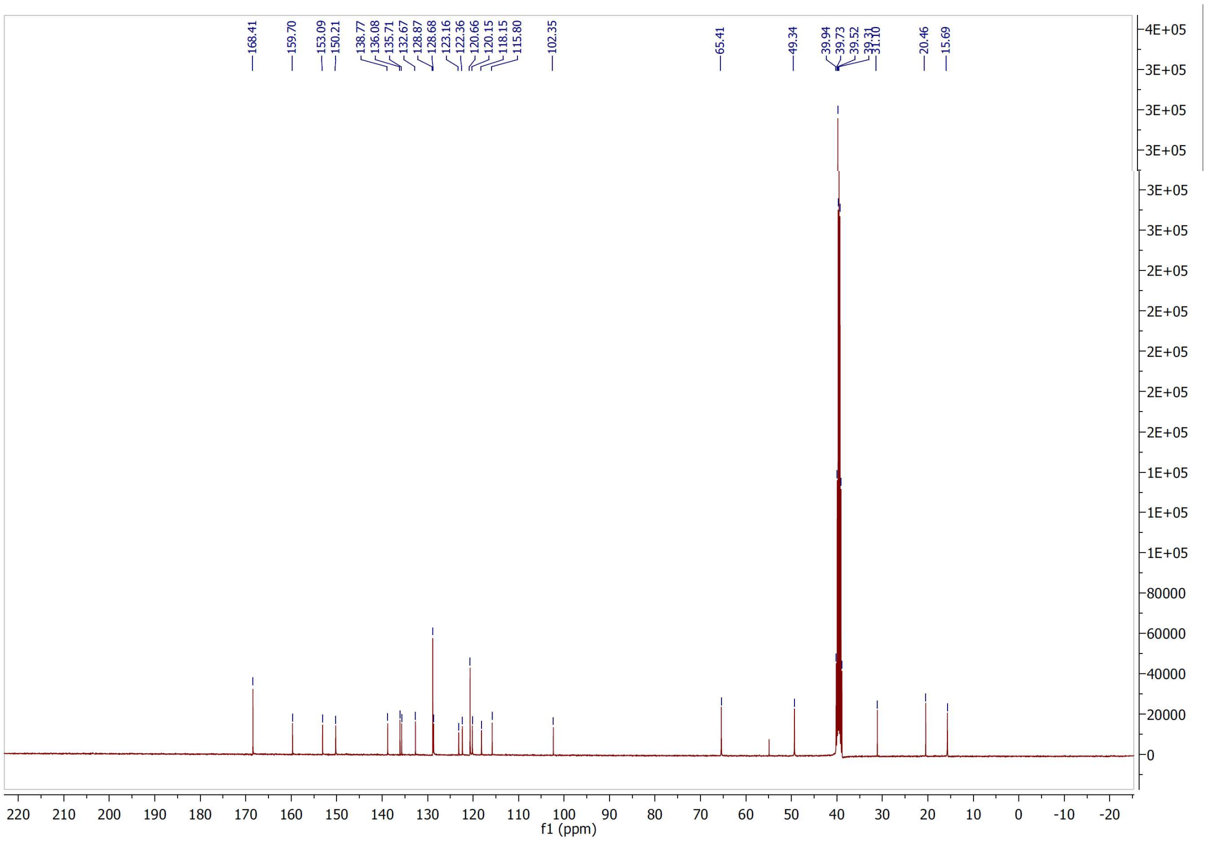

$^{1}H$ and $^{13}C$ NMR spectra of compound **15**

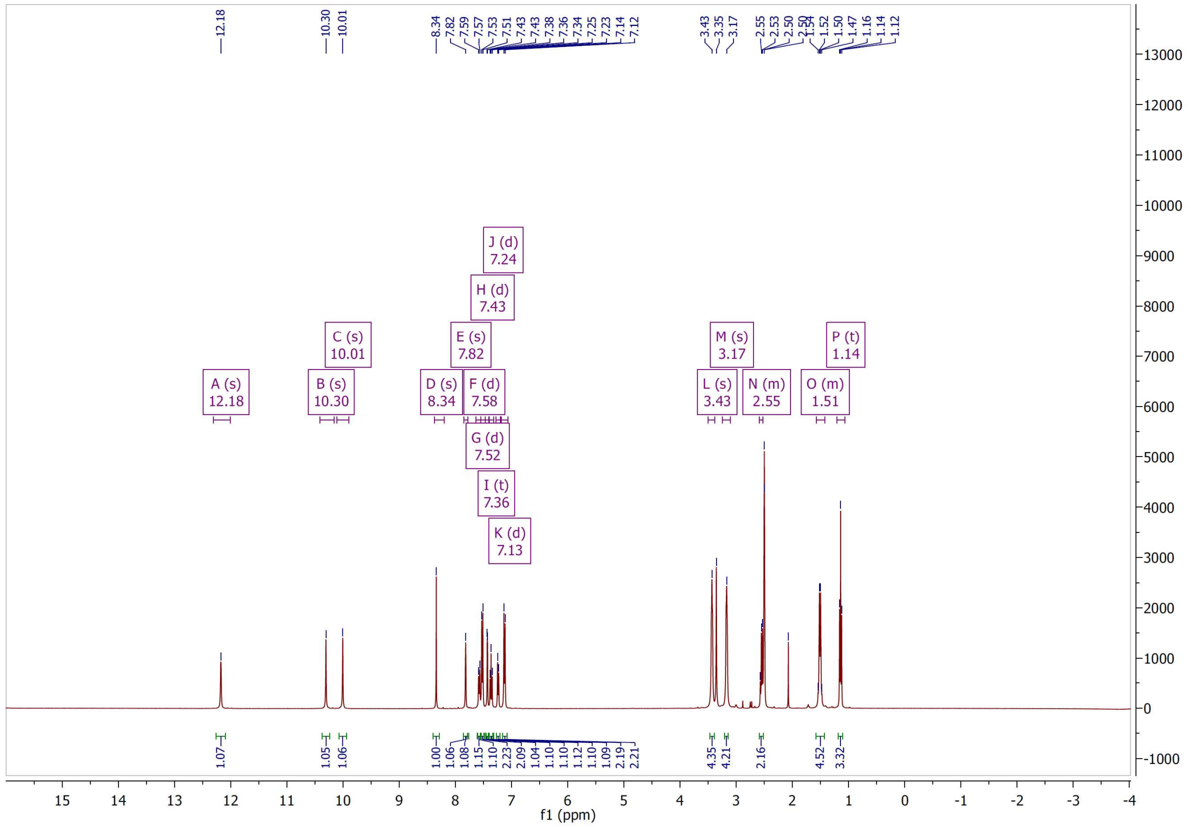

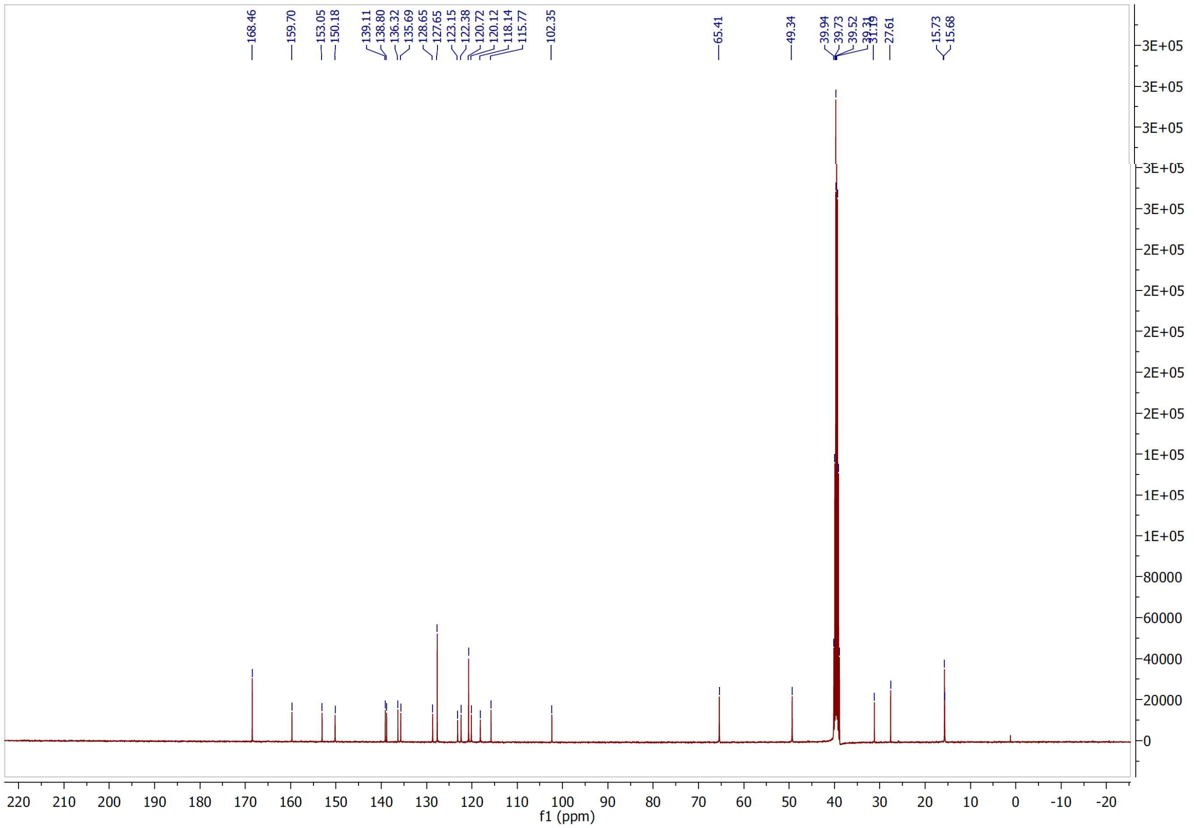

$^{1}H$ and $^{13}C$ NMR spectra of compound **16**

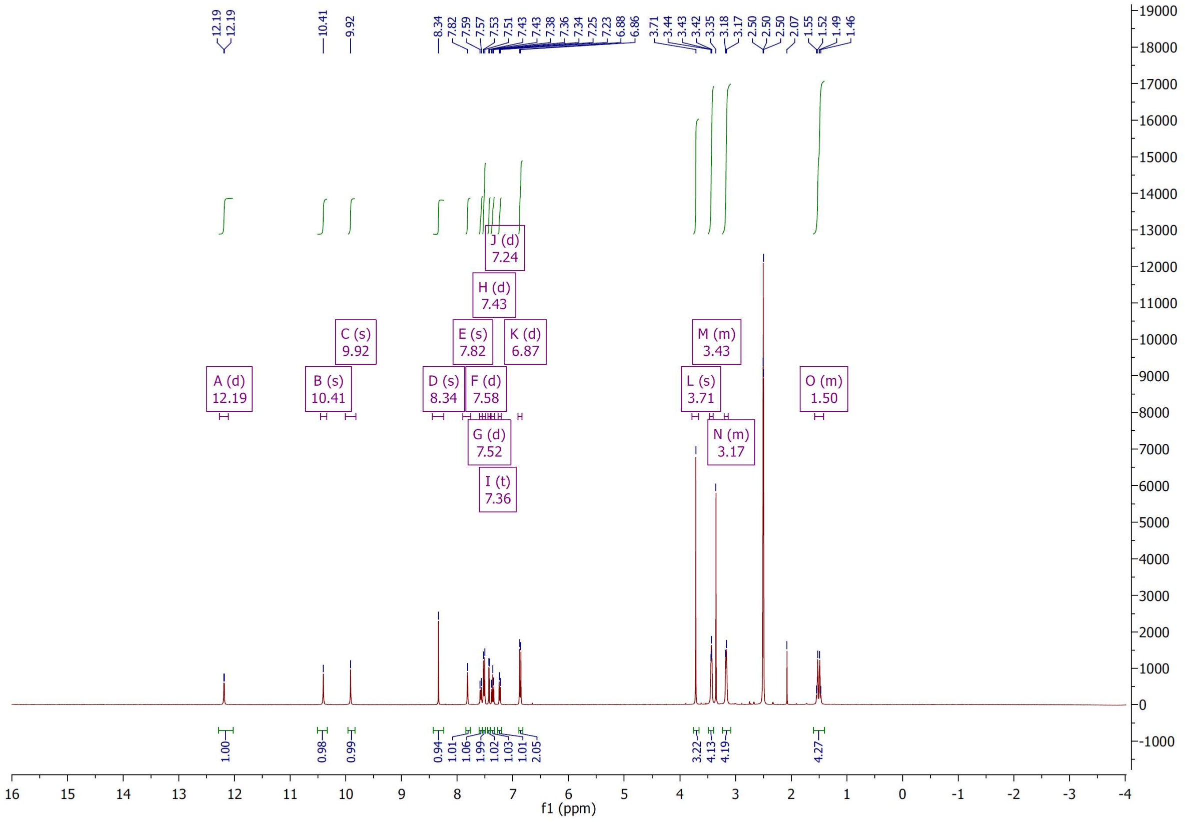


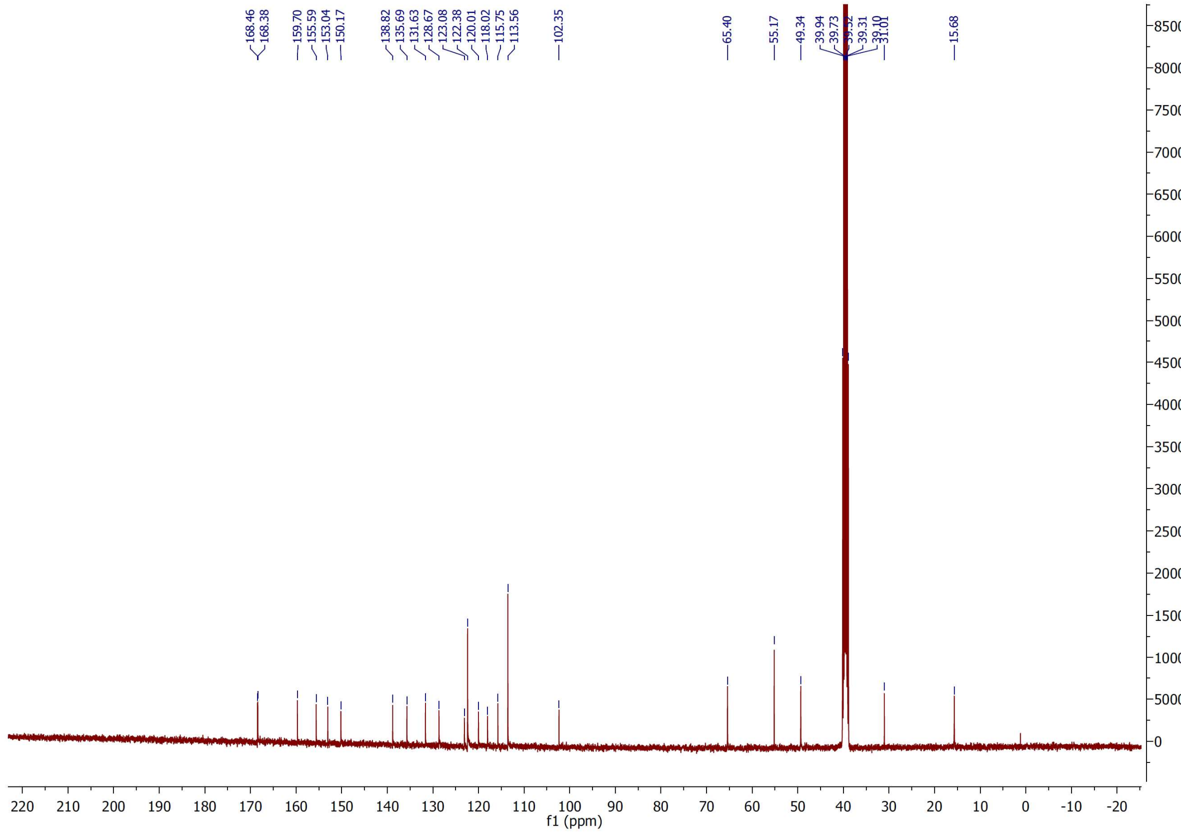

$^{1}H$ and $^{13}C$ NMR spectra of compound **17**

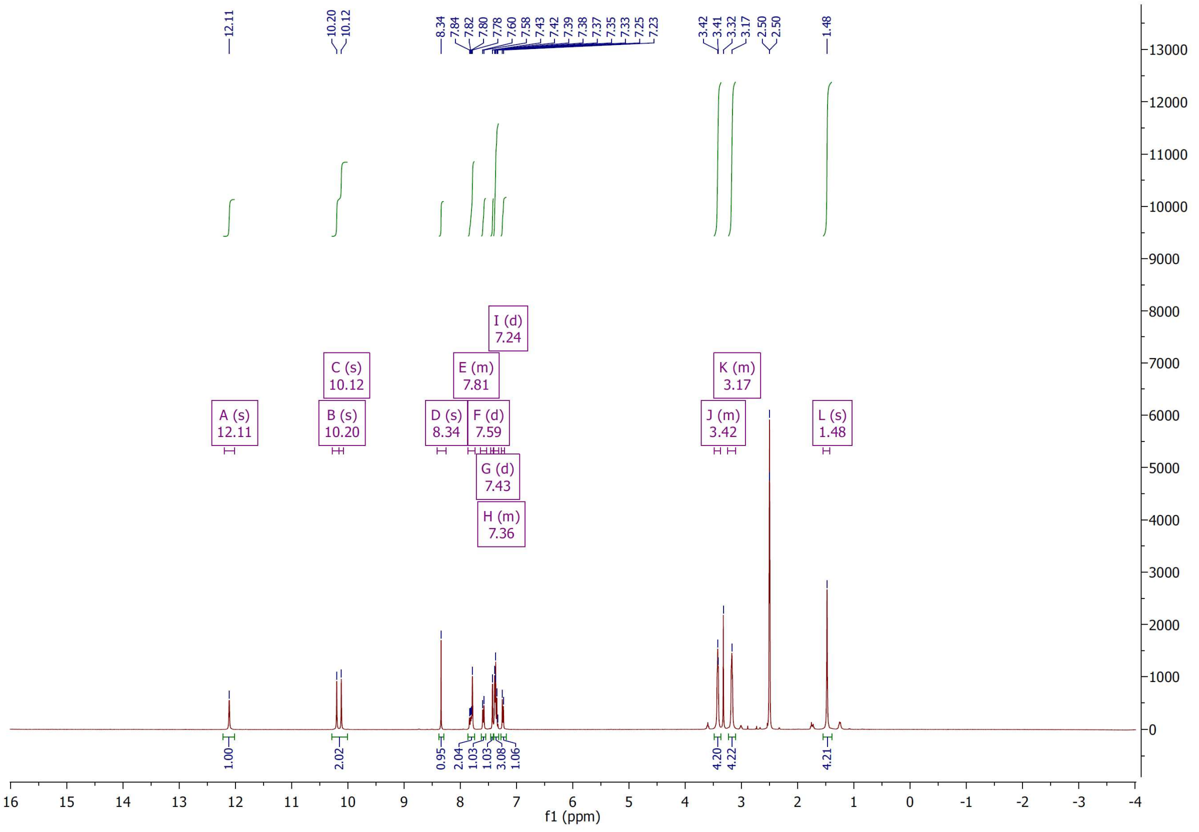


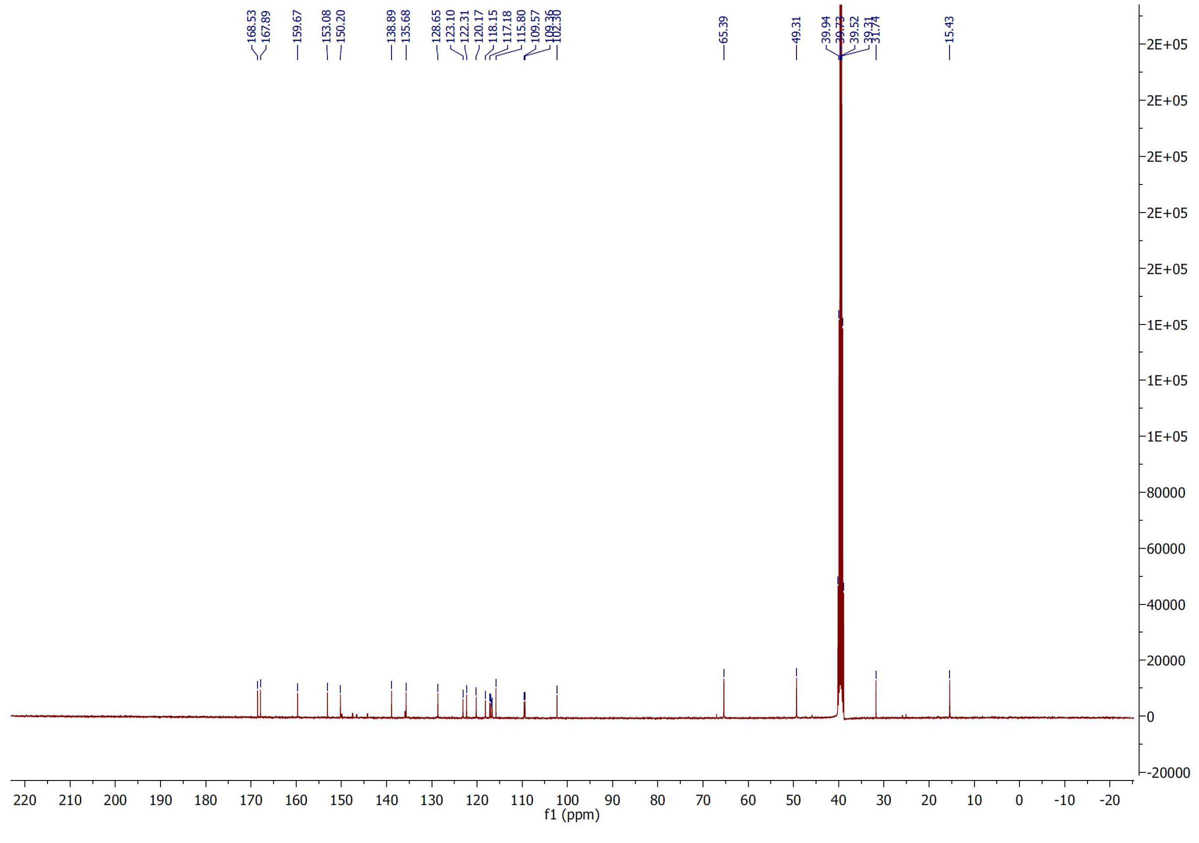

$^{1}H$ and $^{13}C$ NMR spectra of compound **18**

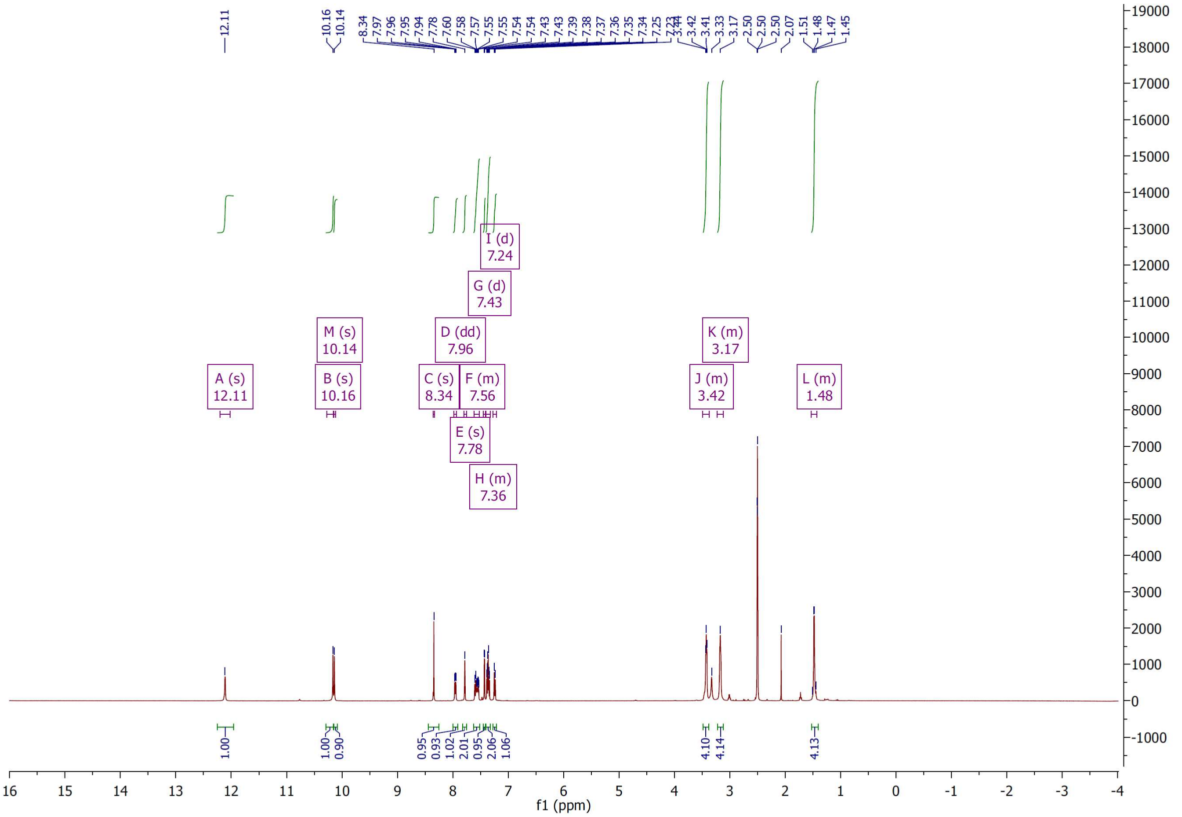

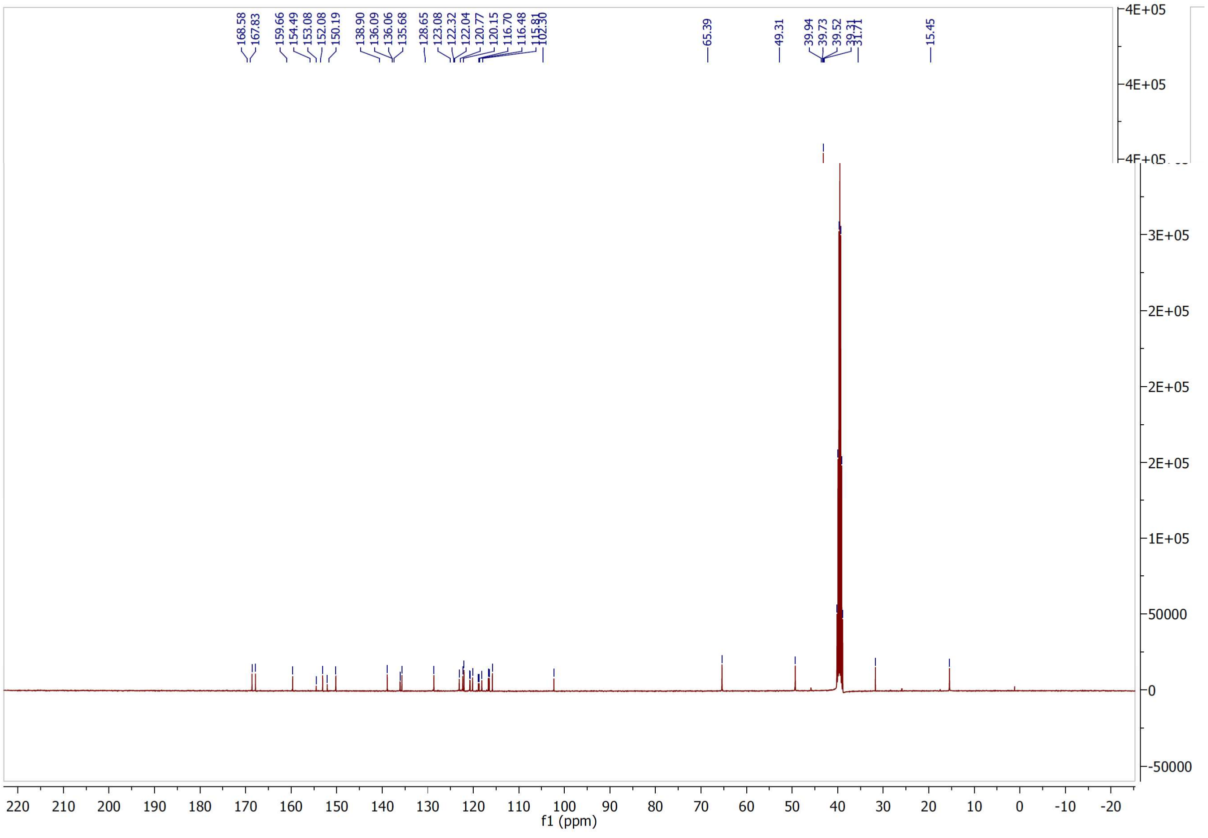

$^{1}H$ and $^{13}C$ NMR spectra of compound **19**

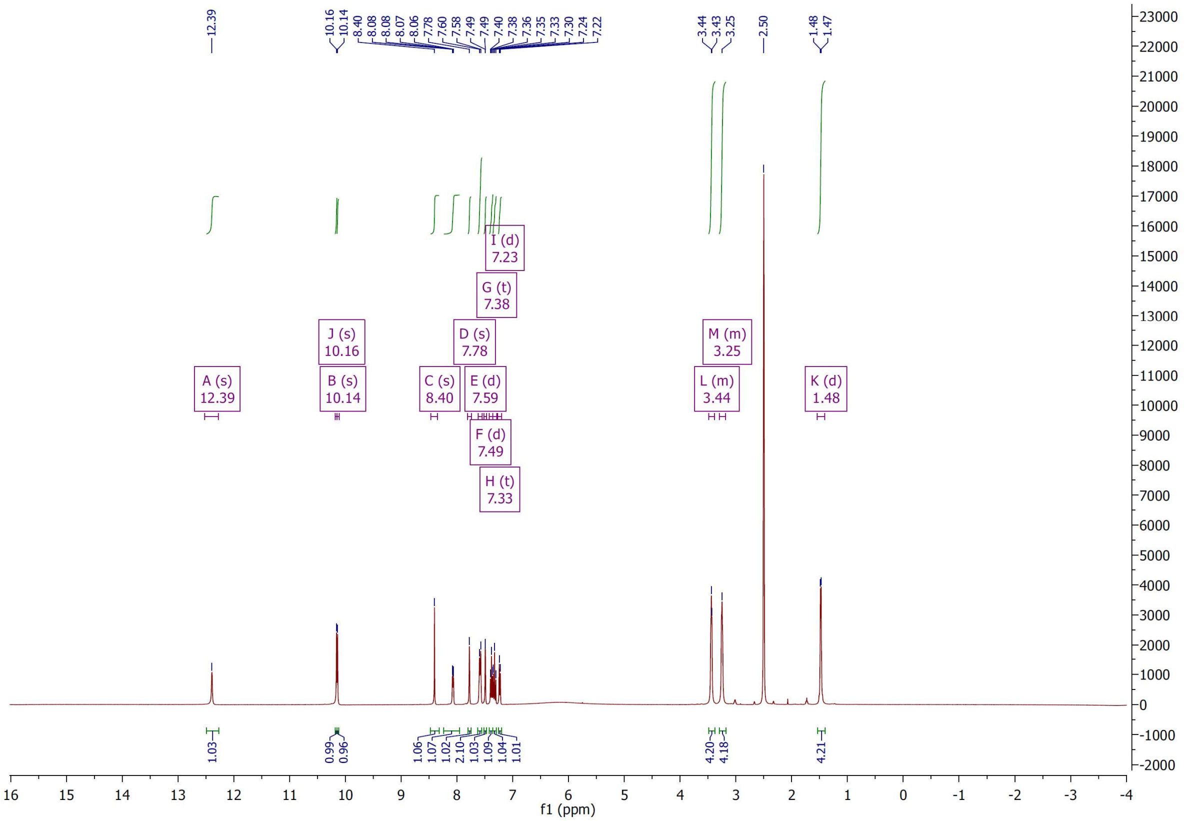

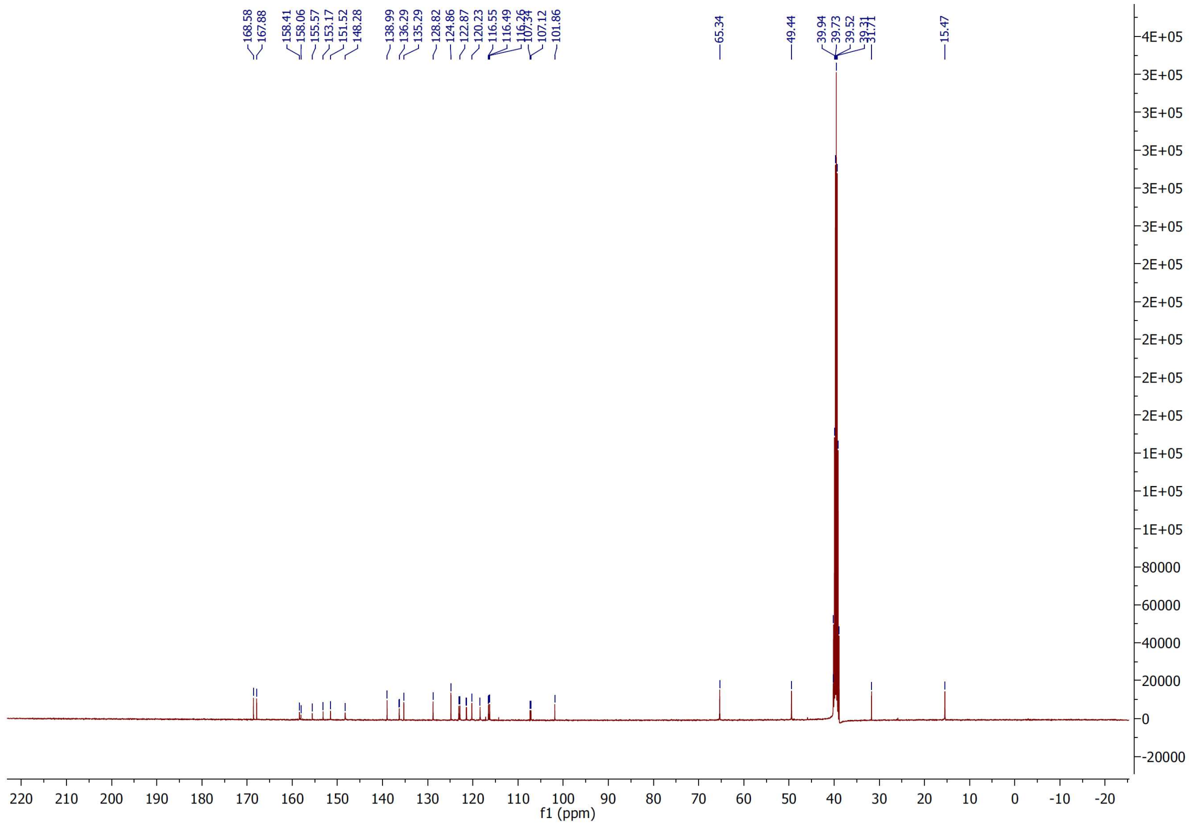

[1]H and [13]C NMR spectra of compound **20**

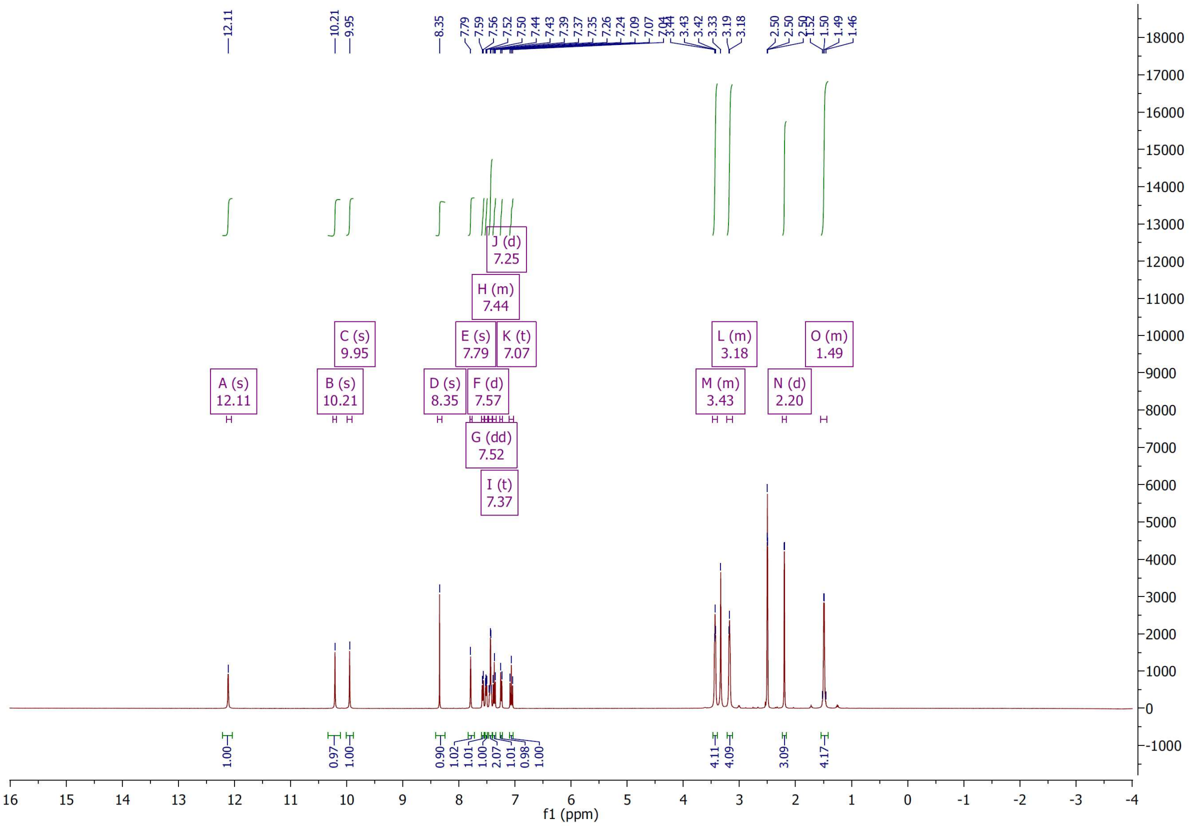

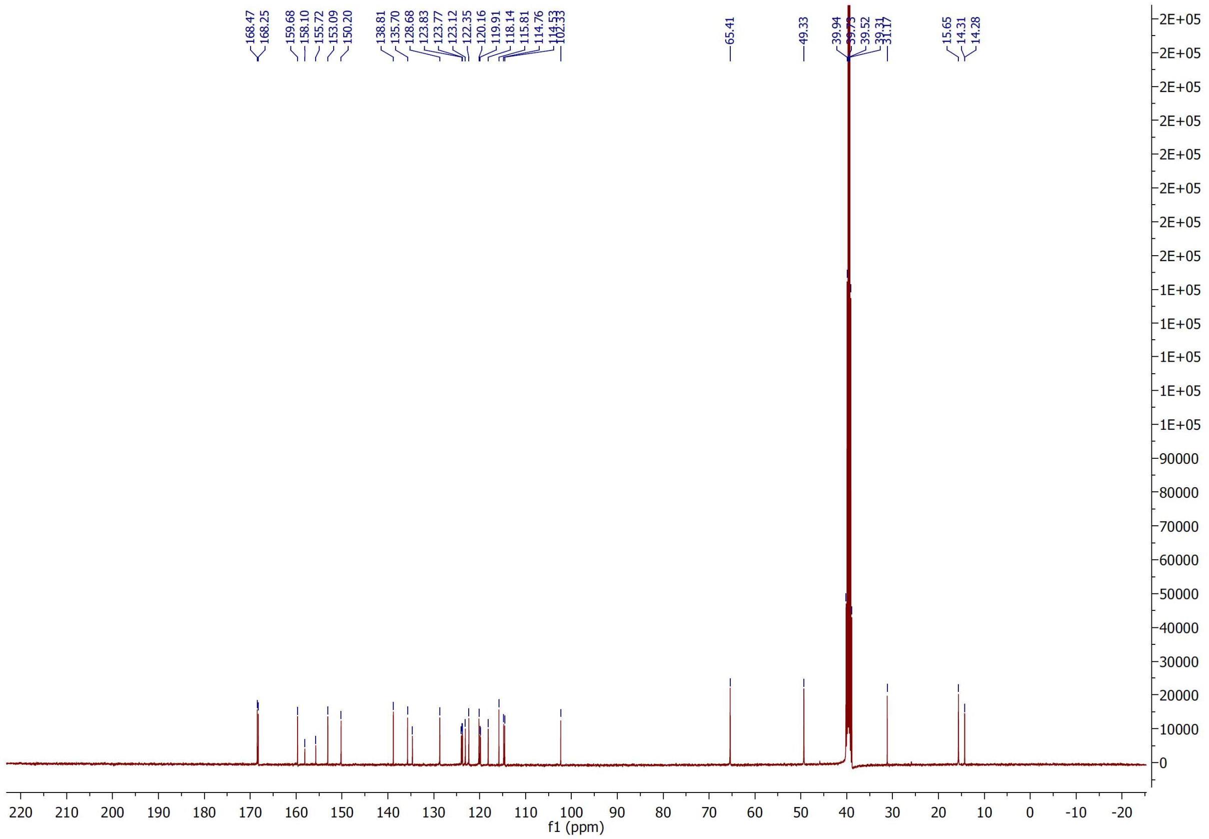

[1]H and [13]C NMR spectra of compound **21**

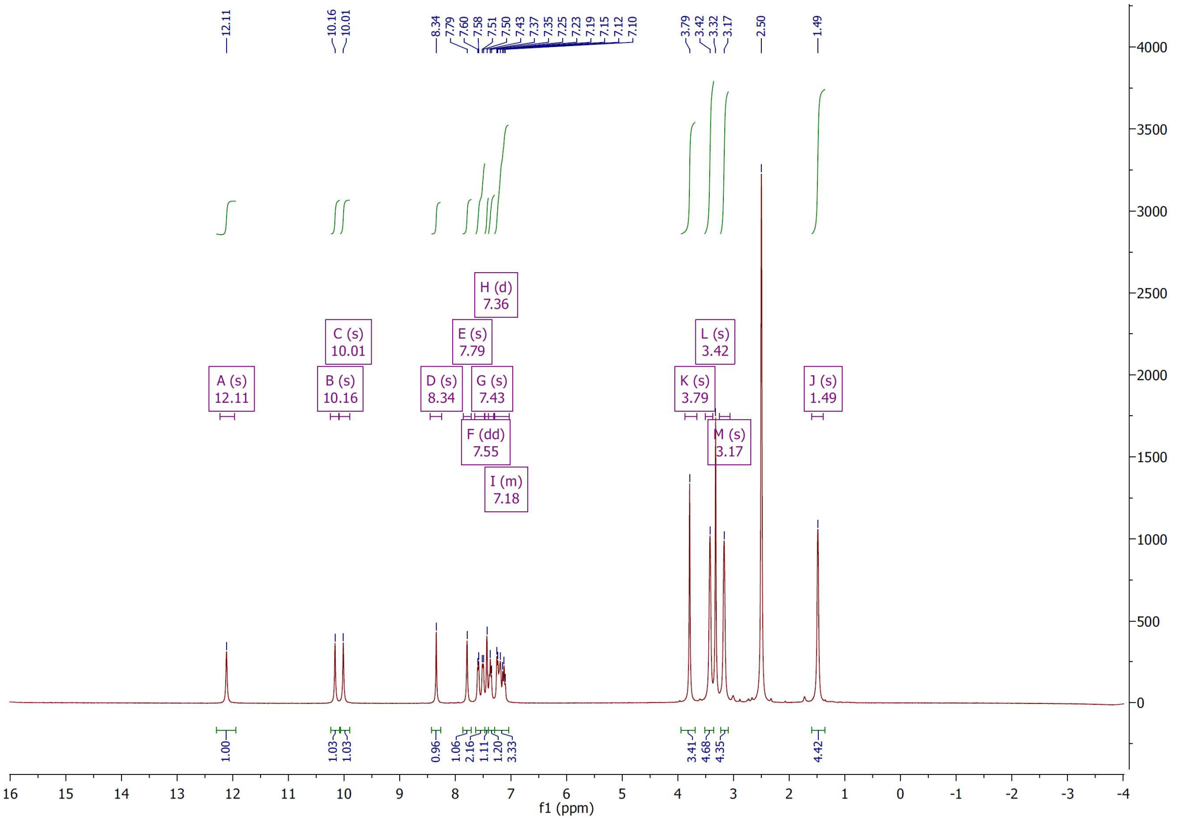

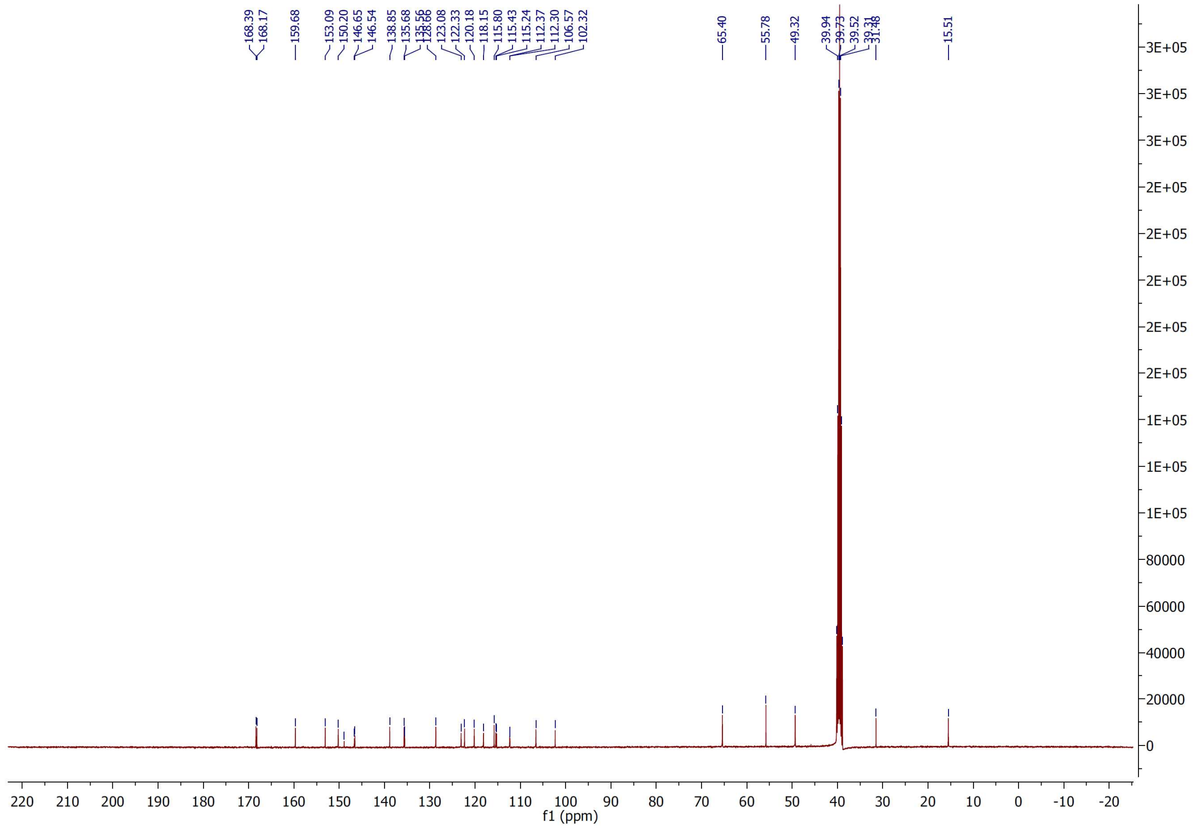

$^{1}H$ and $^{13}C$ NMR spectra of compound **22**

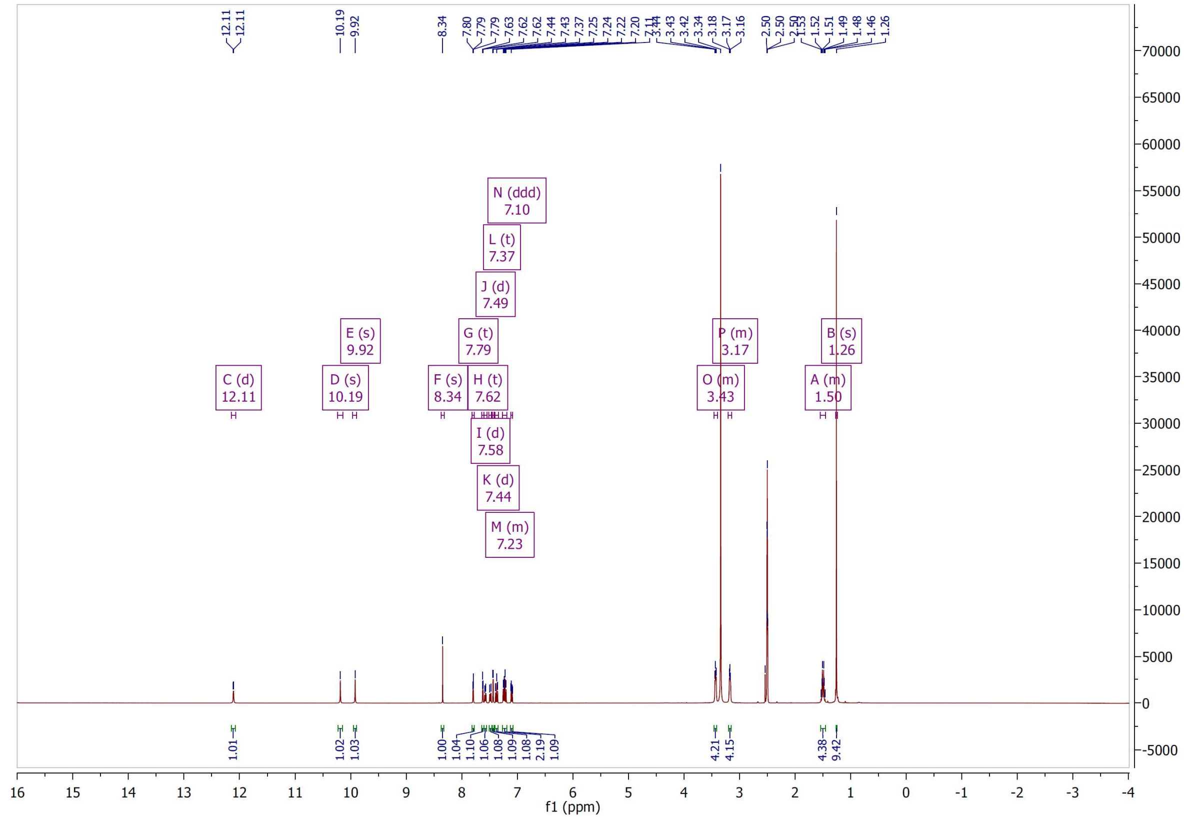

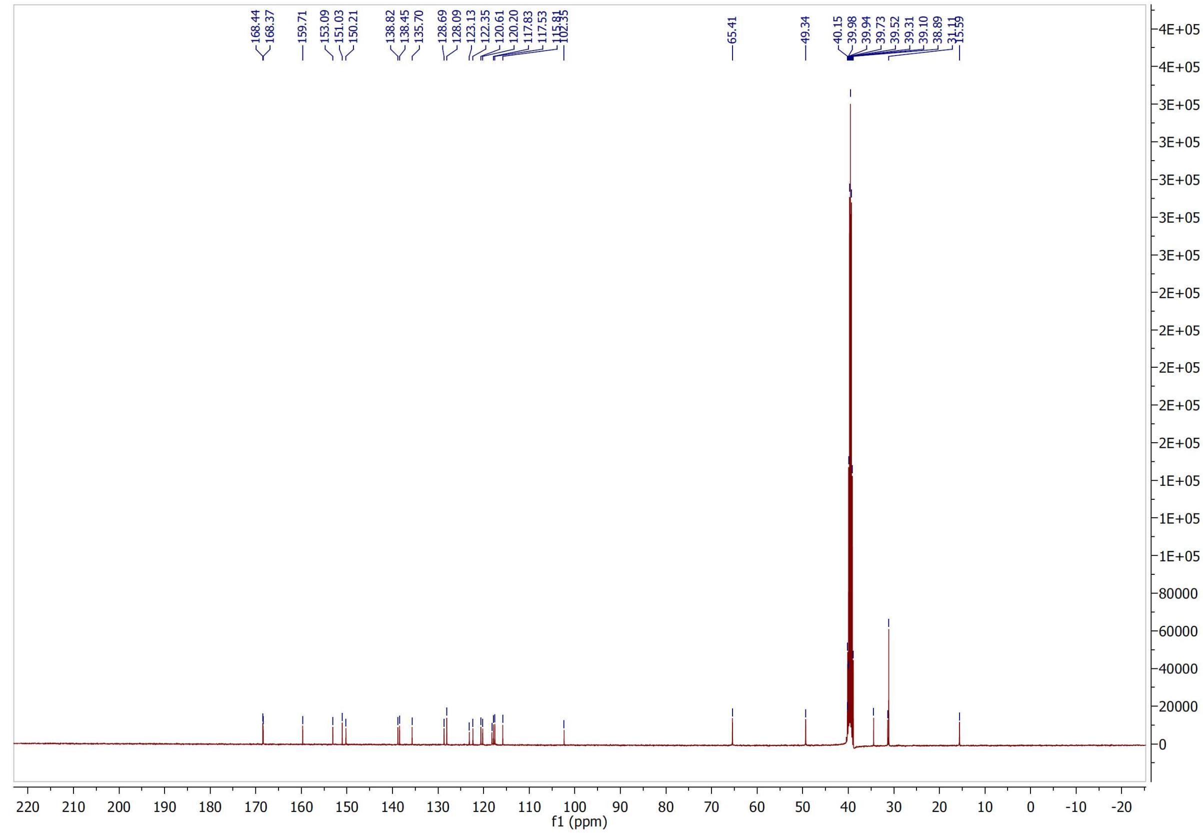

$^{1}H$ and $^{13}C$ NMR spectra of compound **23**

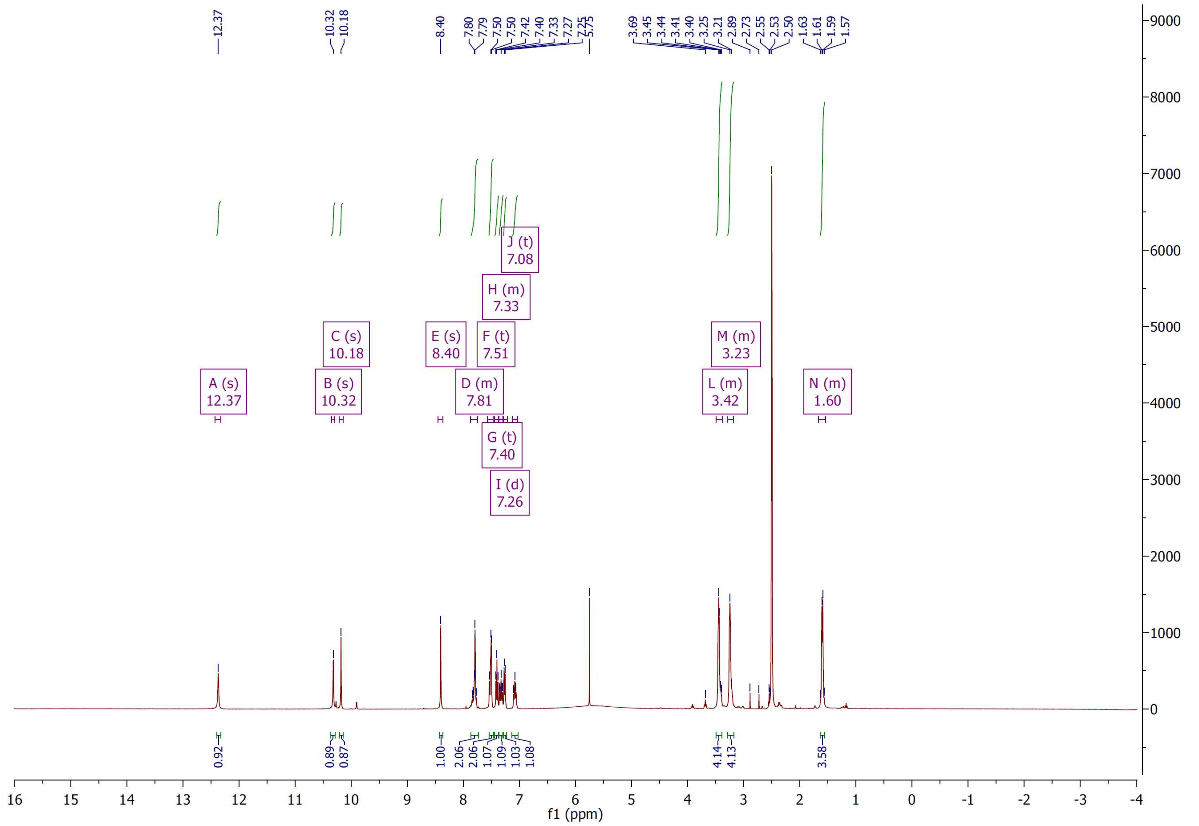

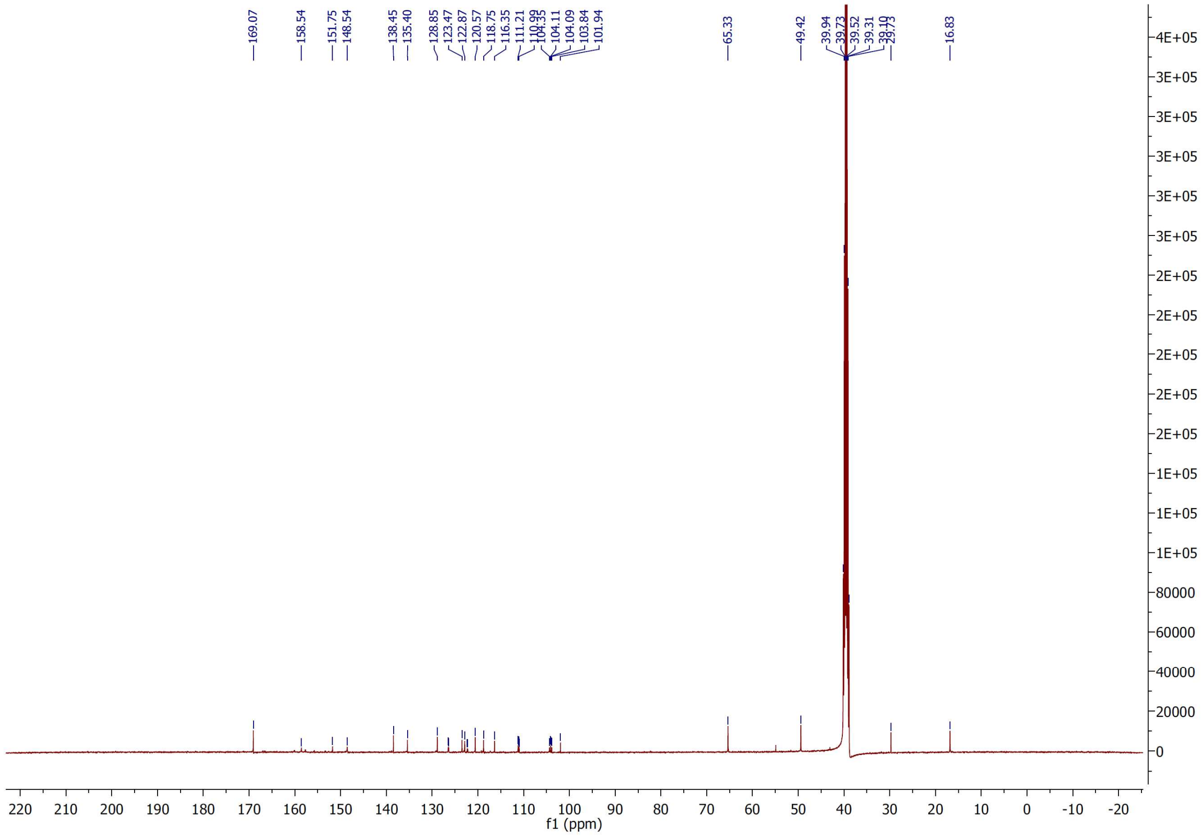

<sup></sup>[1]H and [13]C NMR spectra of compound **24**

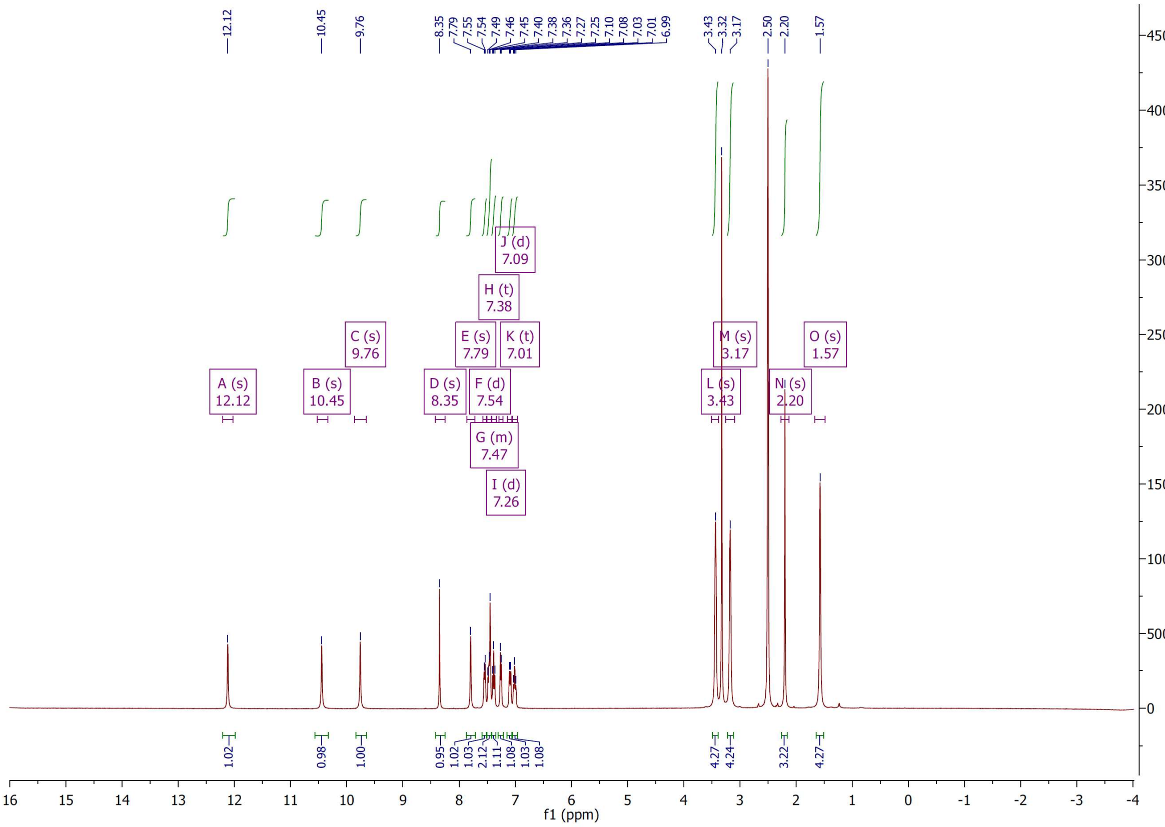

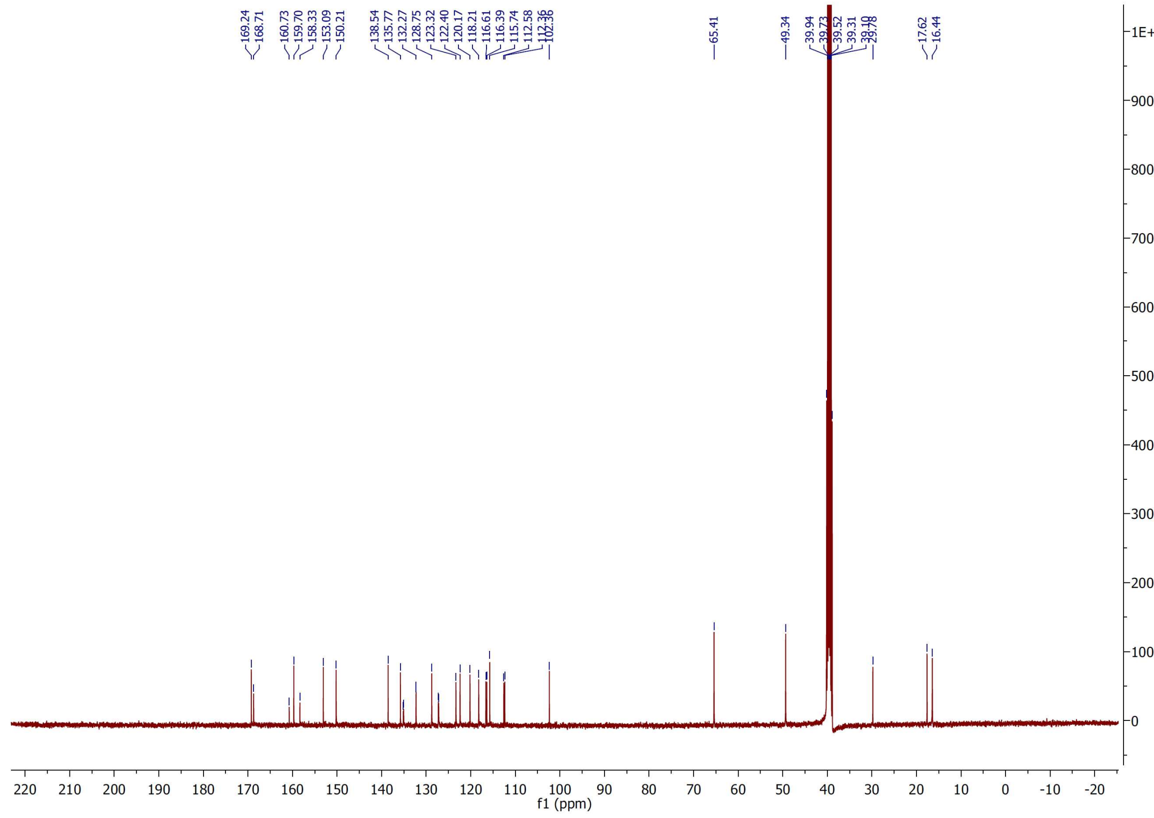

$^1H$ and $^{13}C$ NMR spectra of compound **25**

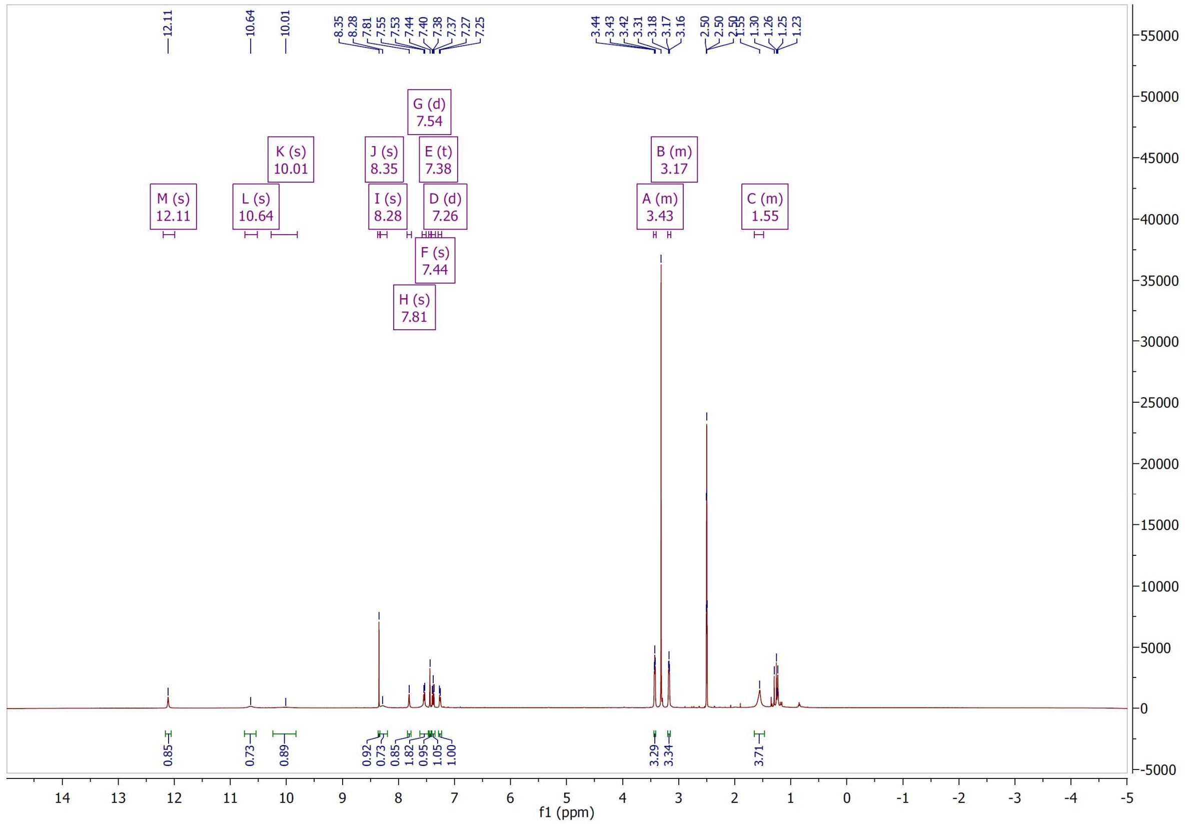


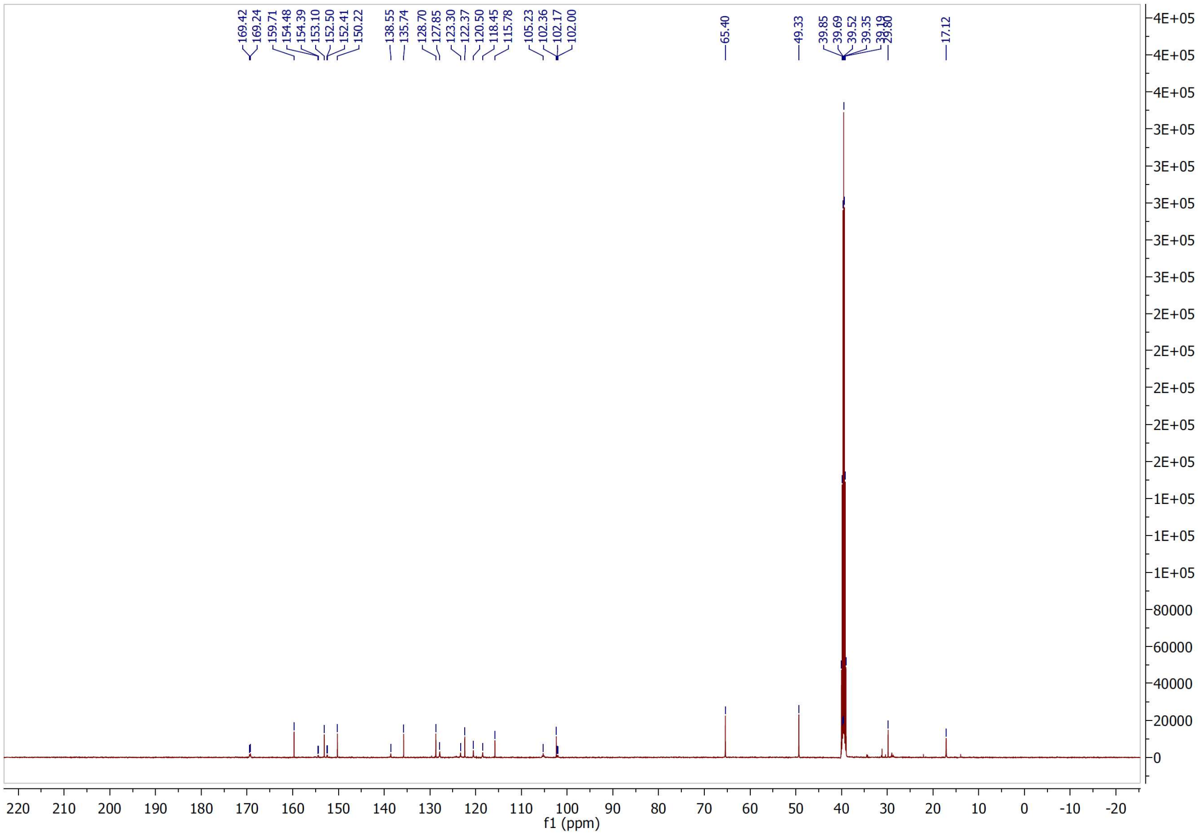

$^{1}H$ and $^{13}C$ NMR spectra of compound **26**

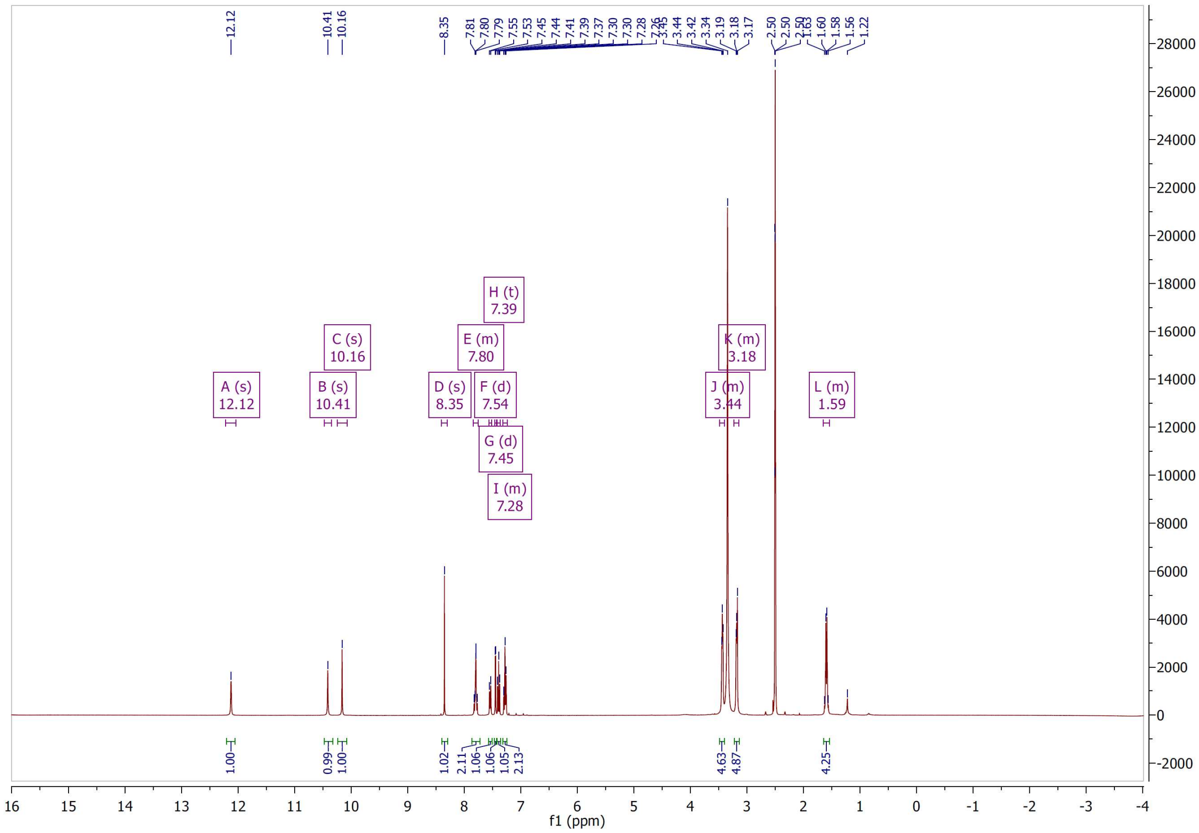

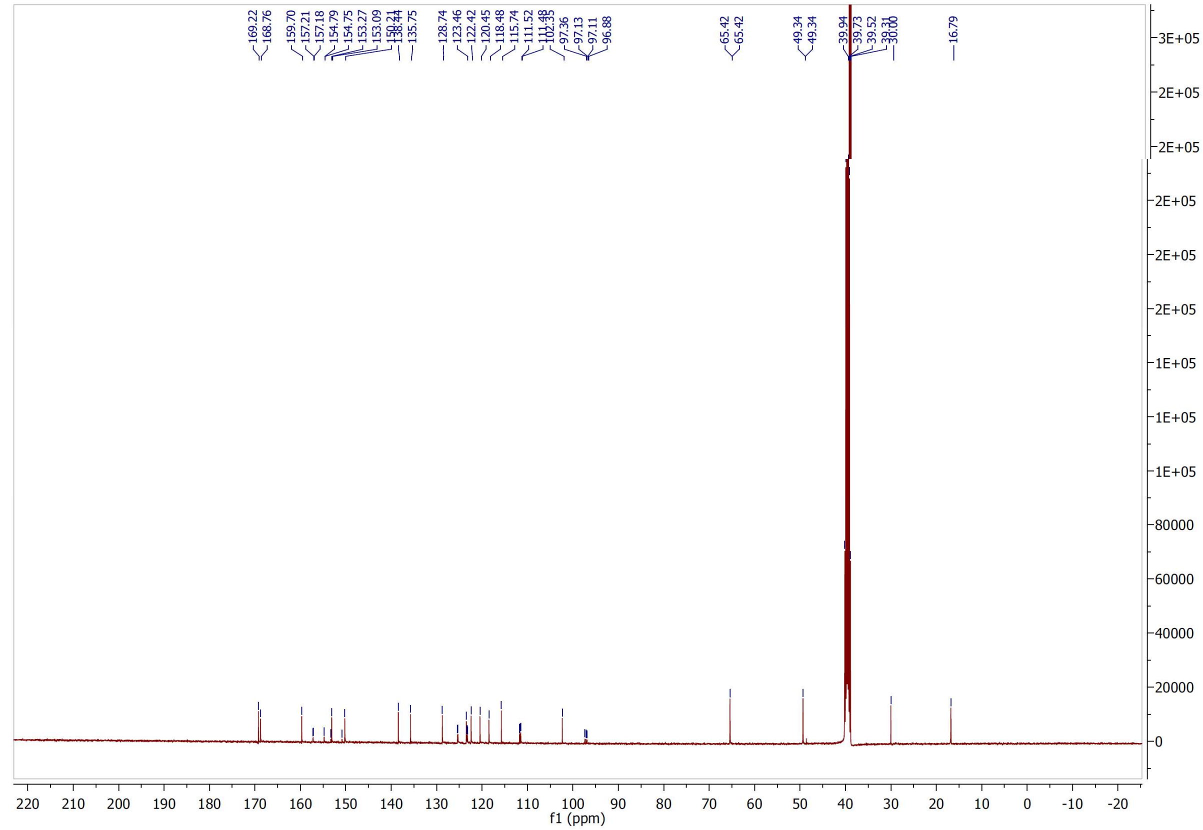

$^{1}H$ and $^{13}C$ NMR spectra of compound **27**

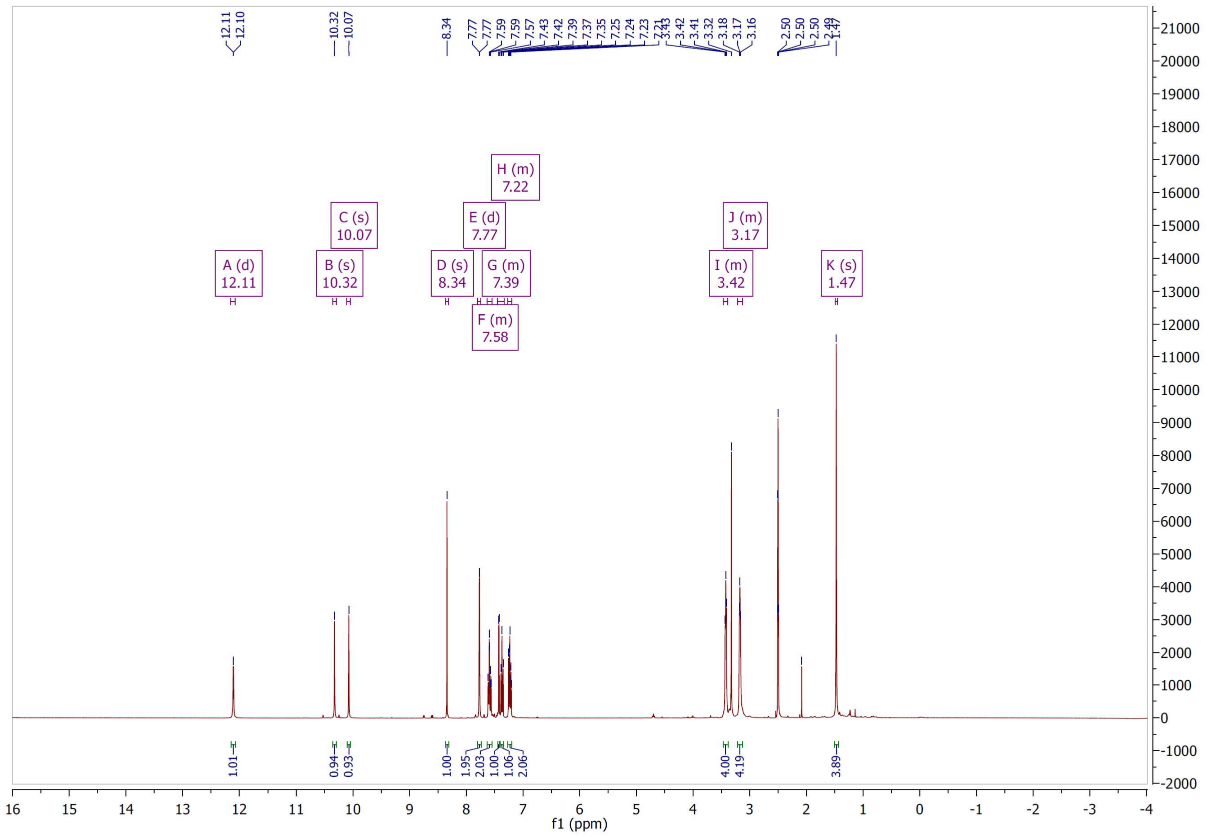

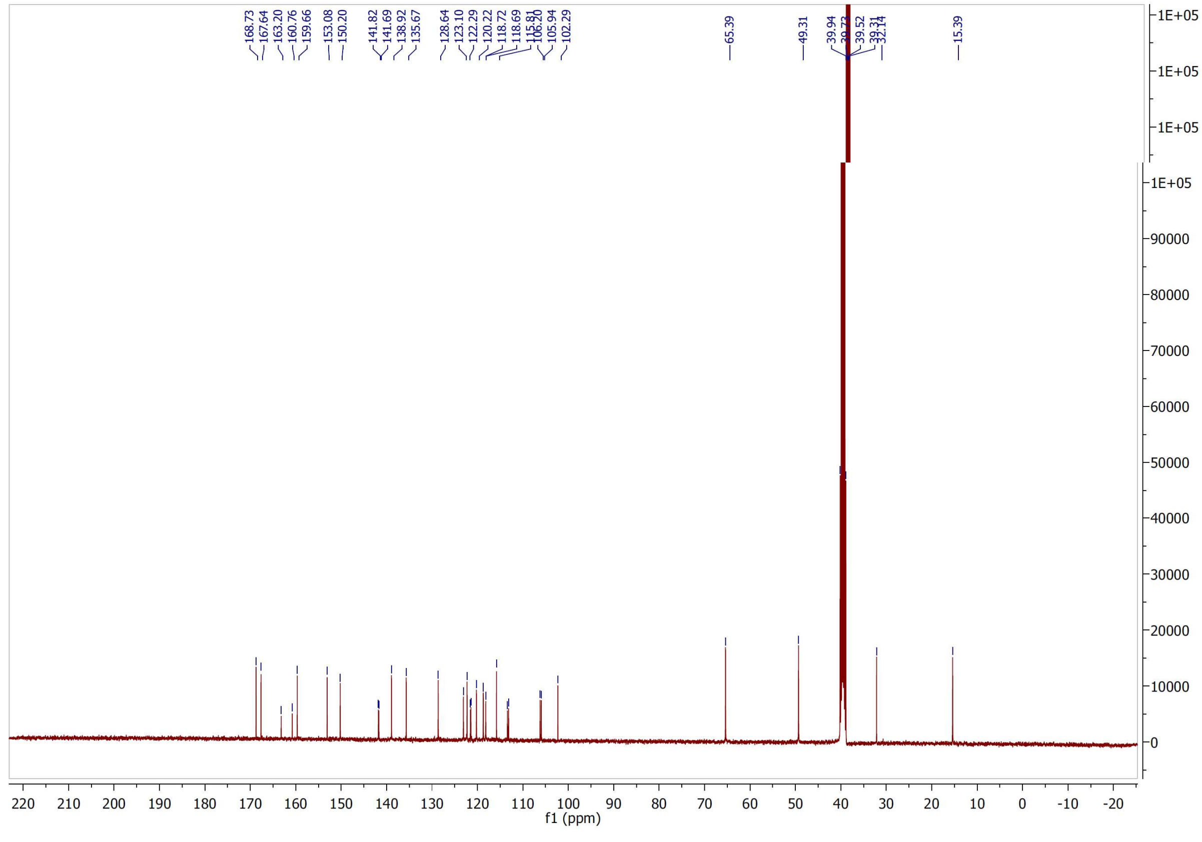

[1]H and [13]C NMR spectra of compound **28**

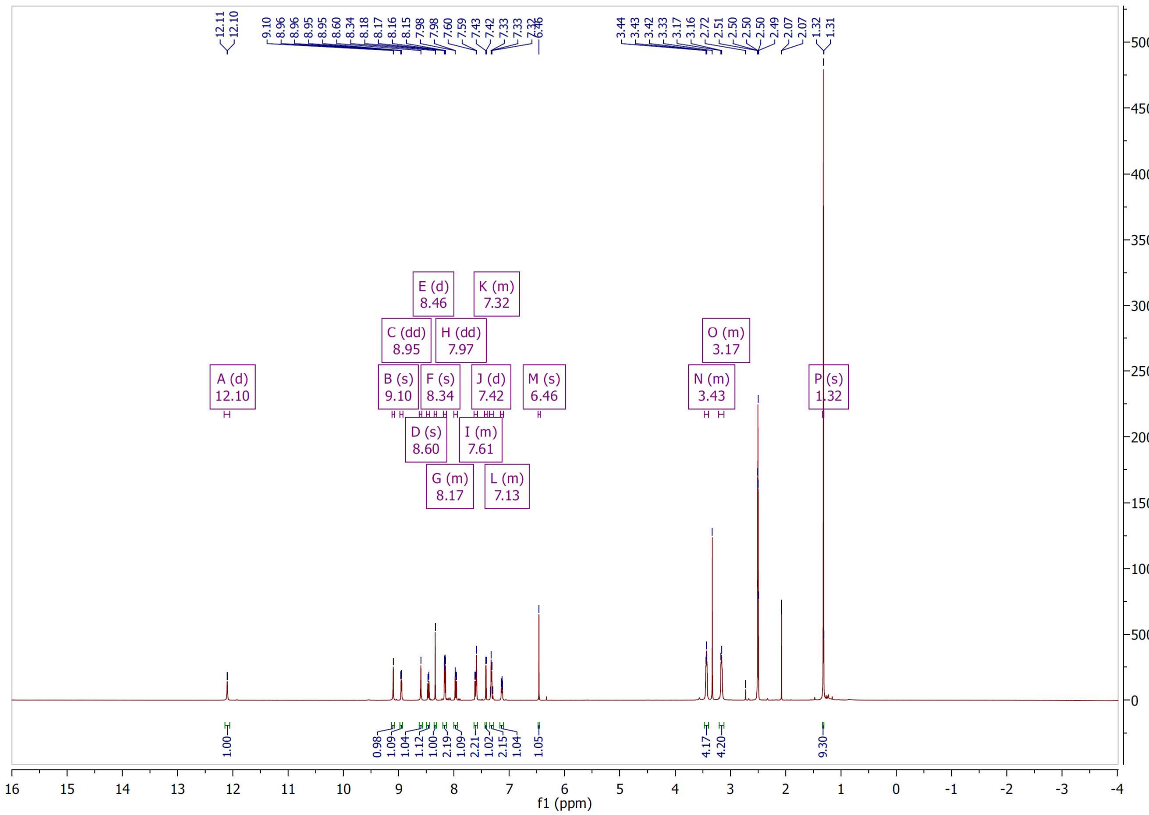

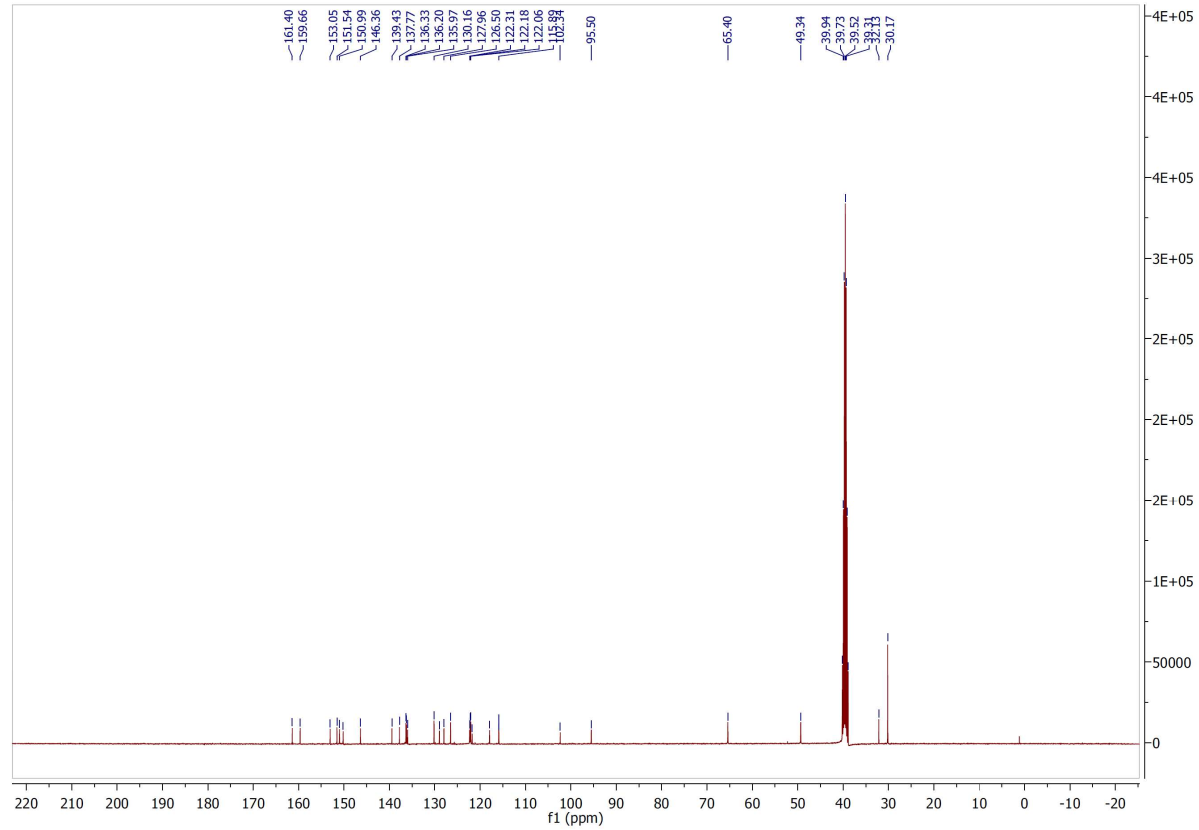

[1]H and [13]C NMR spectra of compound **29**

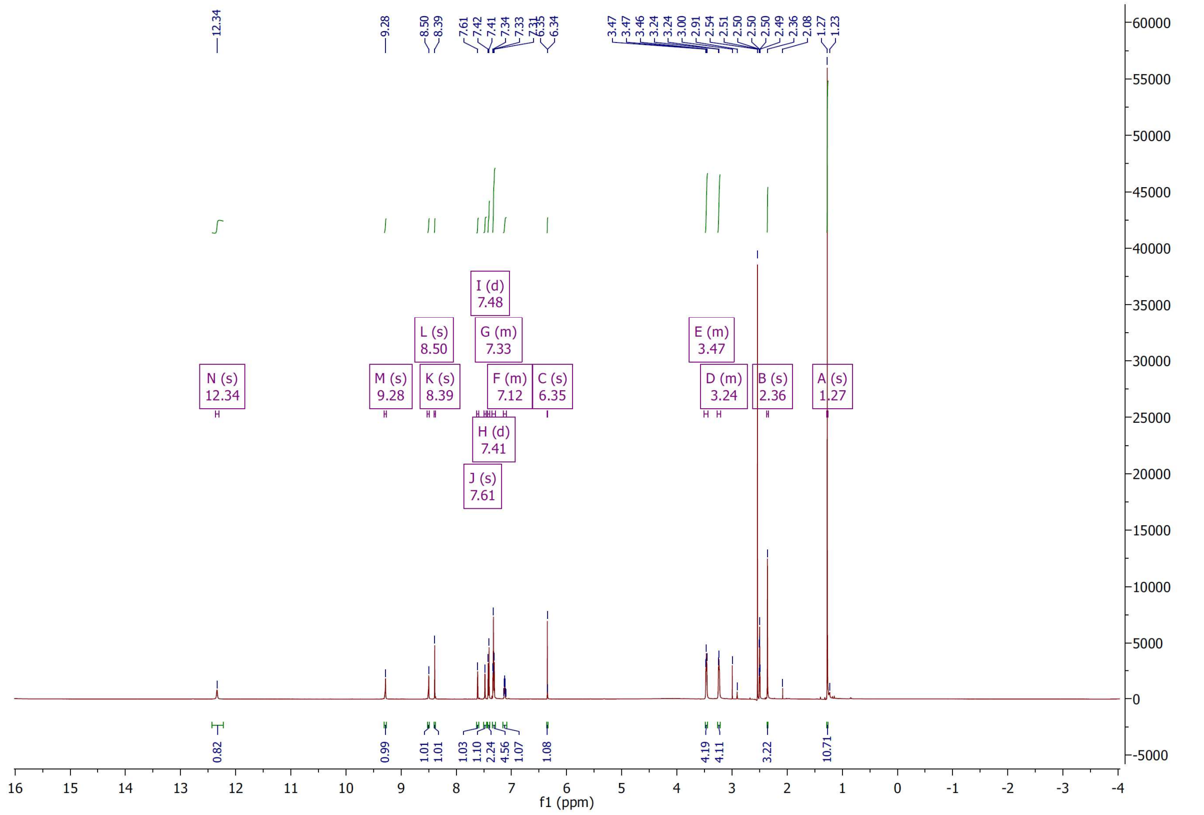

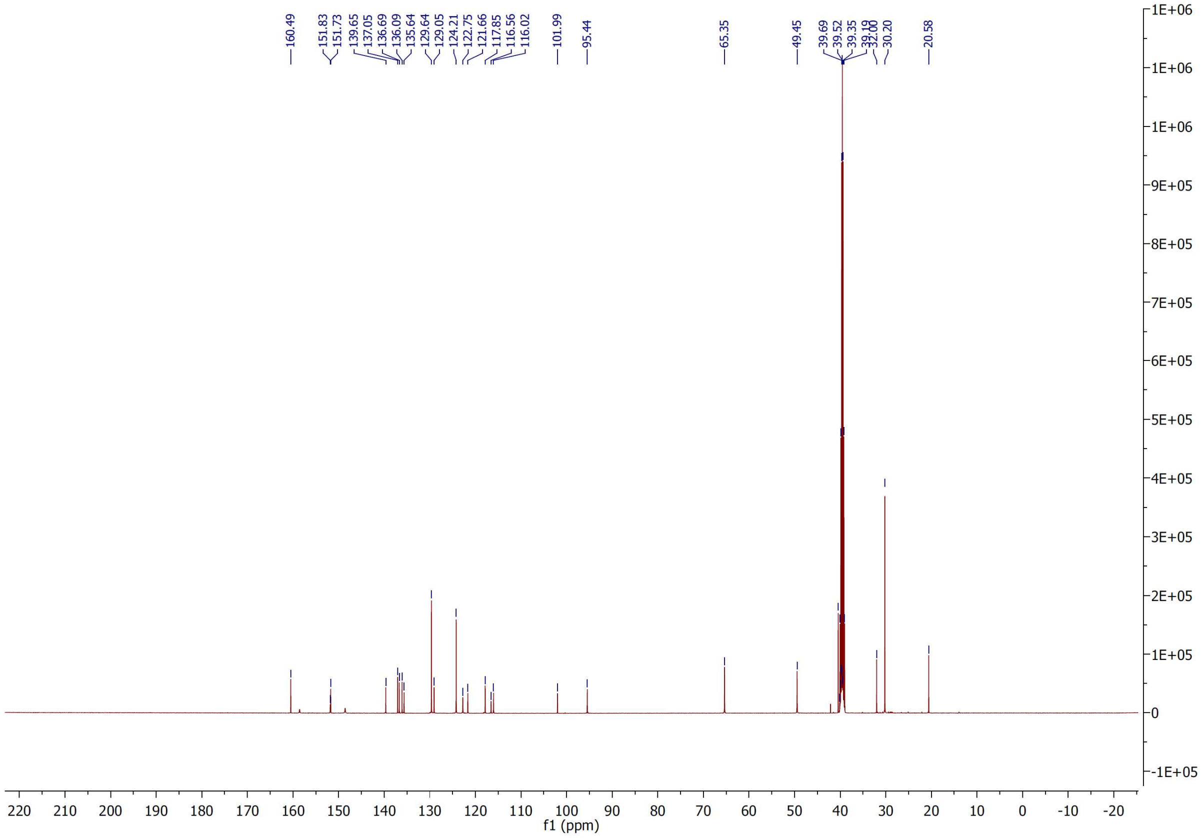

[1]H and [13]C NMR spectra of compound **30**

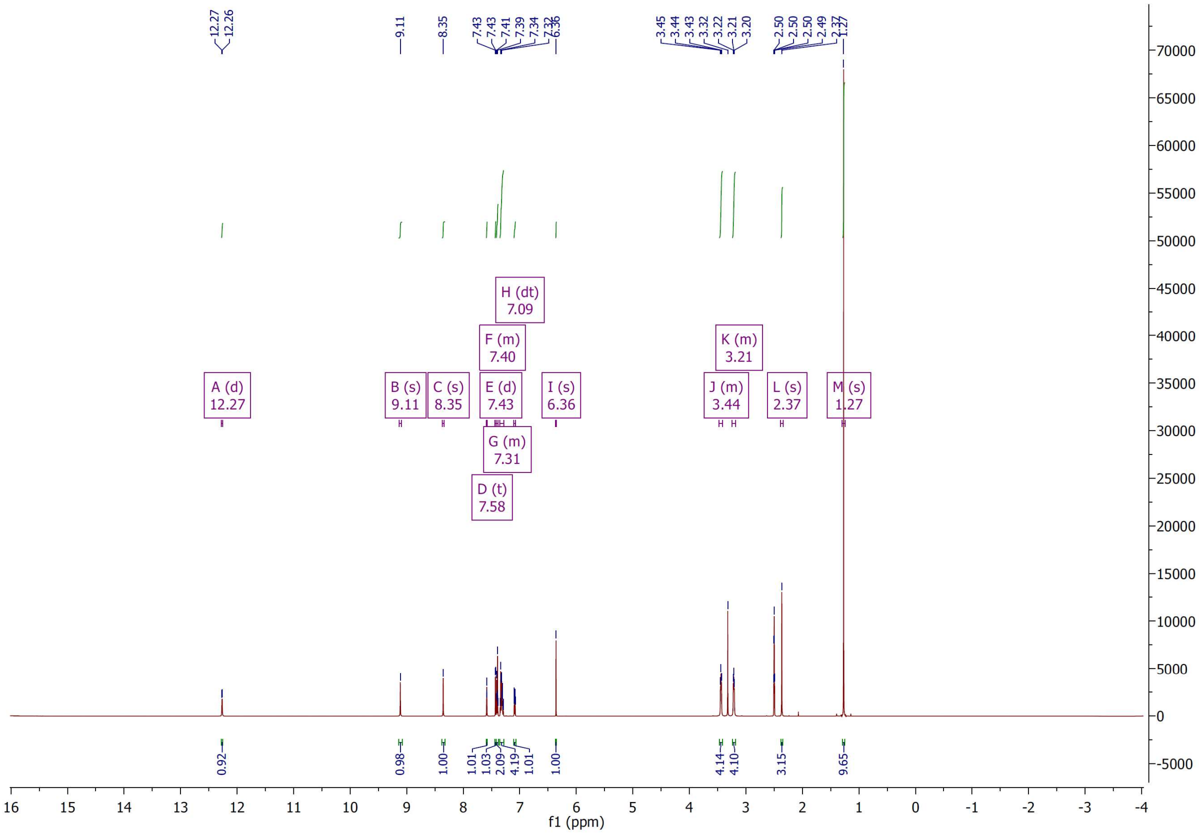

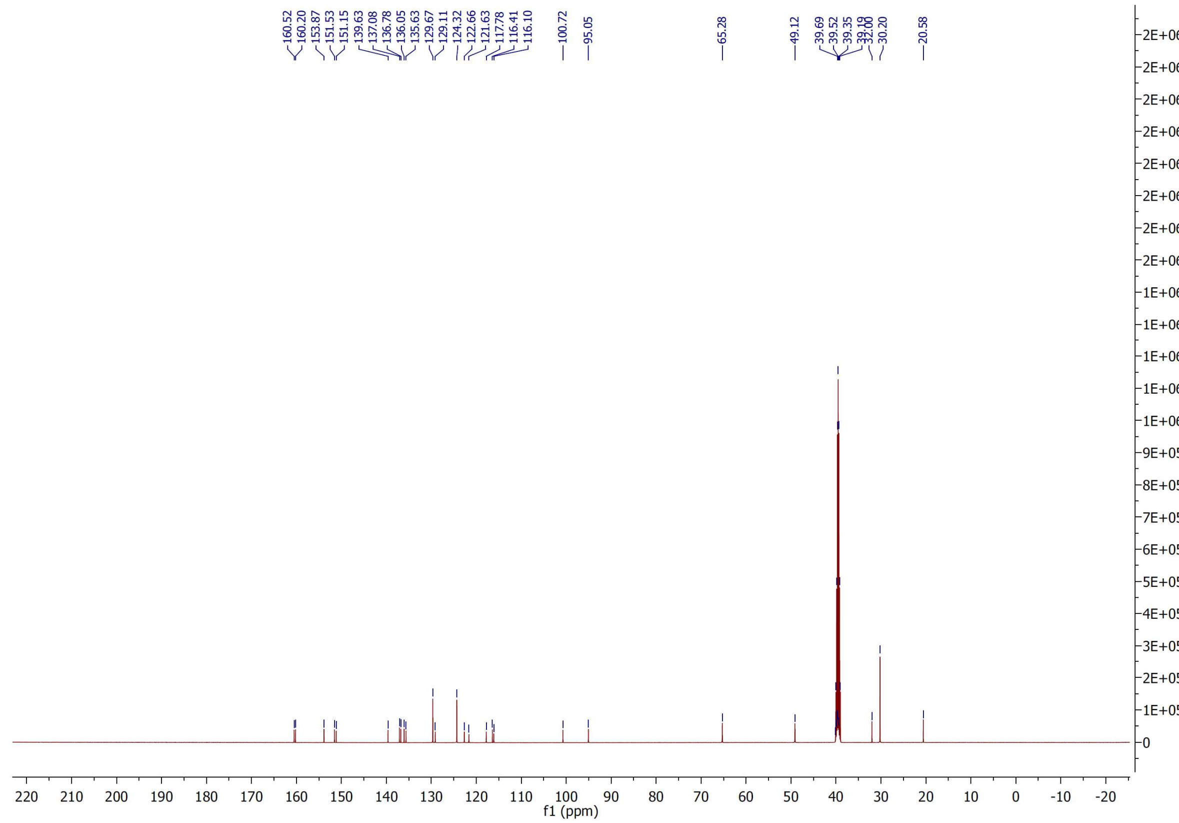

$^{1}H$ and $^{13}C$ NMR spectra of compound **31**

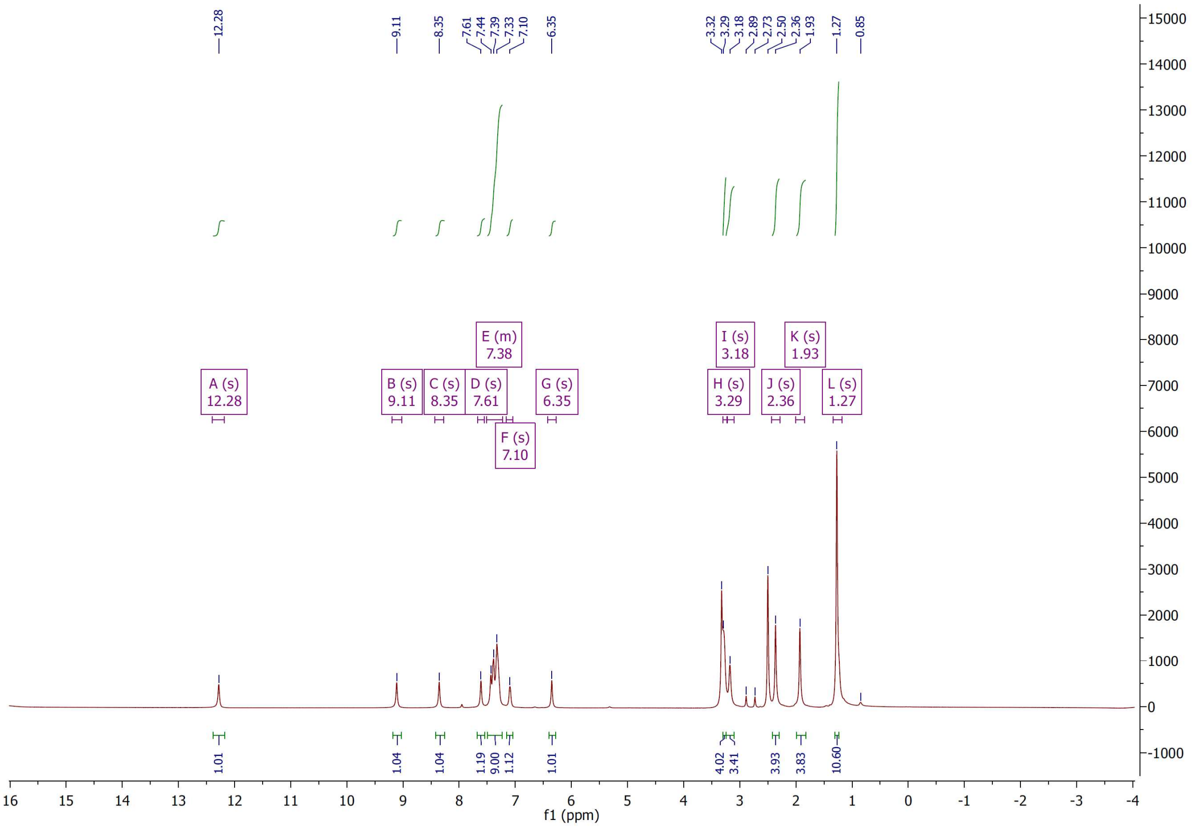


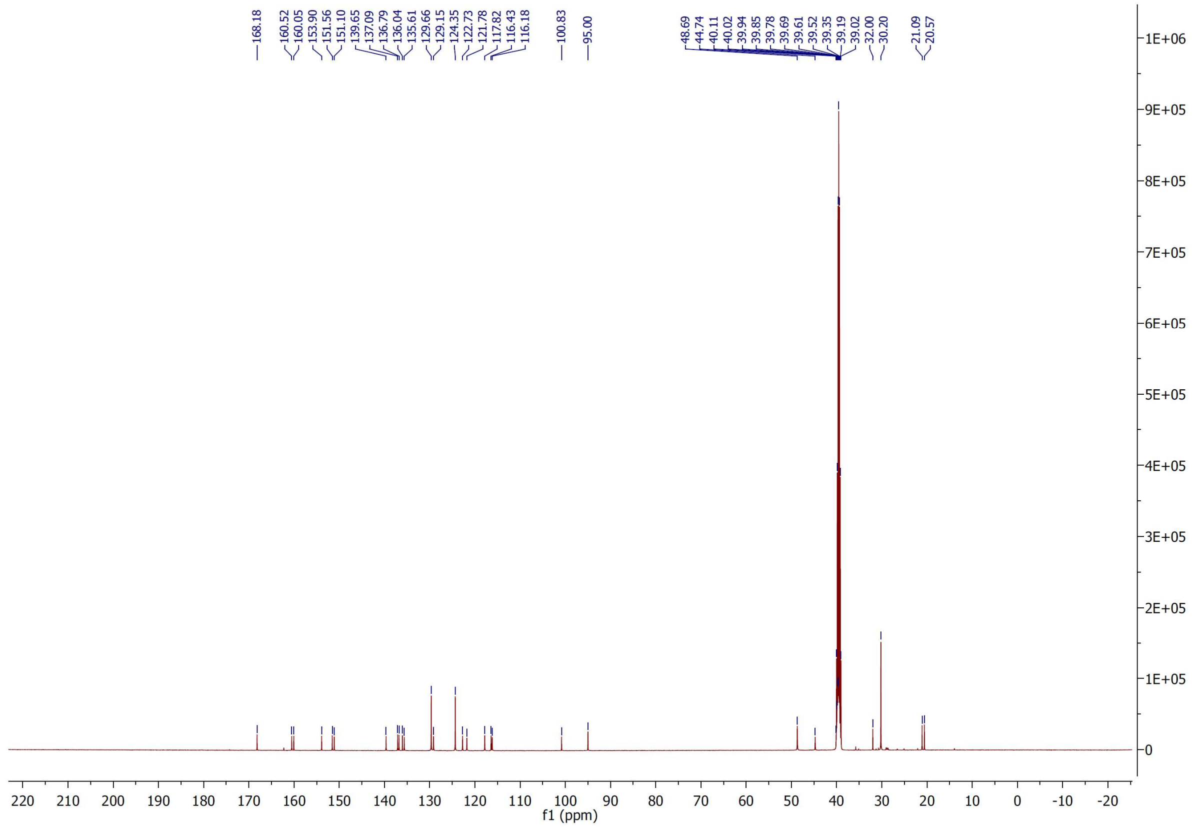

$^1H$ and $^{13}C$ NMR spectra of compound **32**

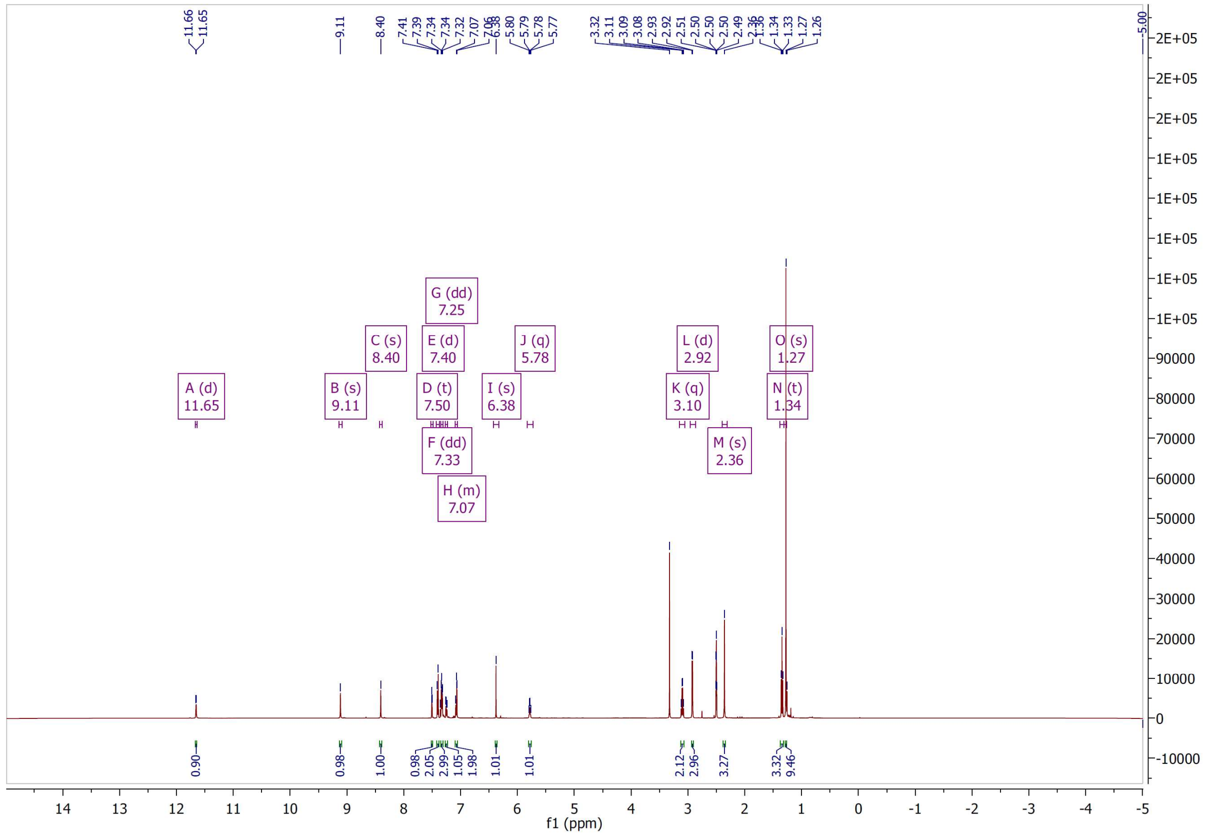

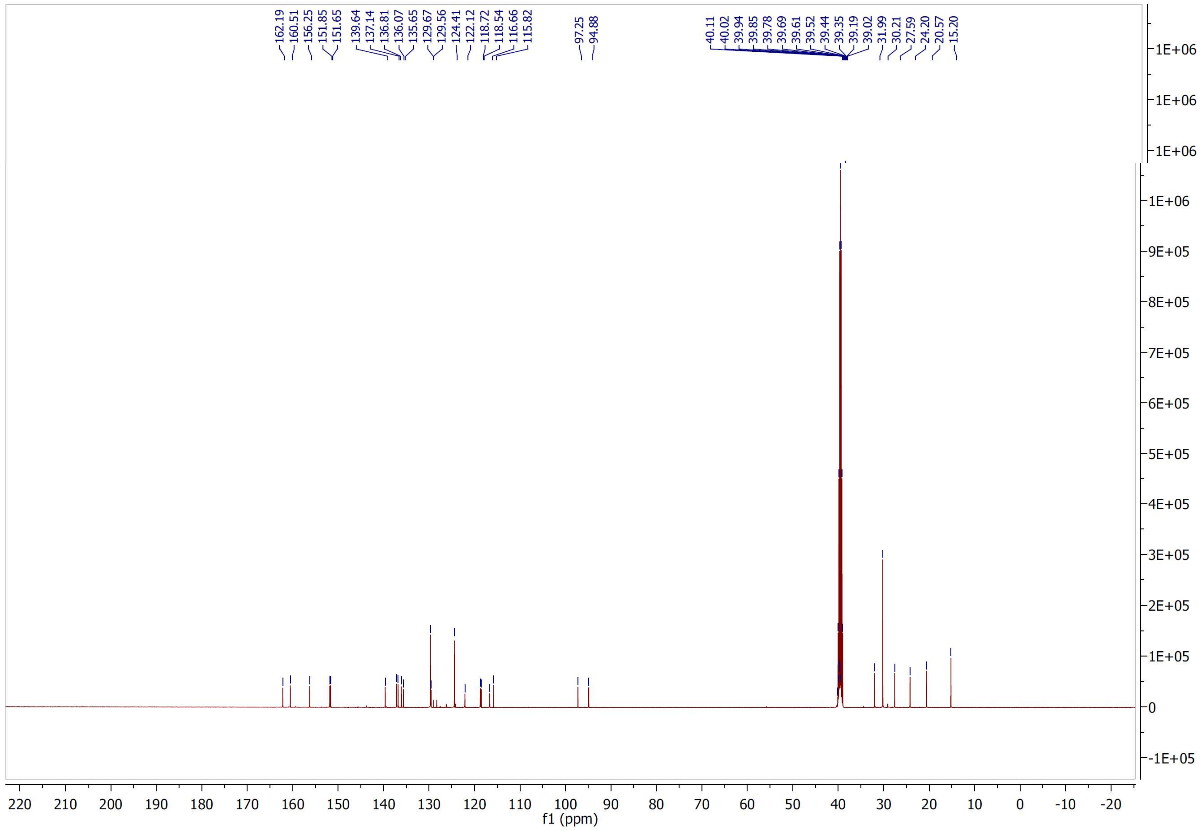

[1]H and [13]C NMR spectra of compound **33**

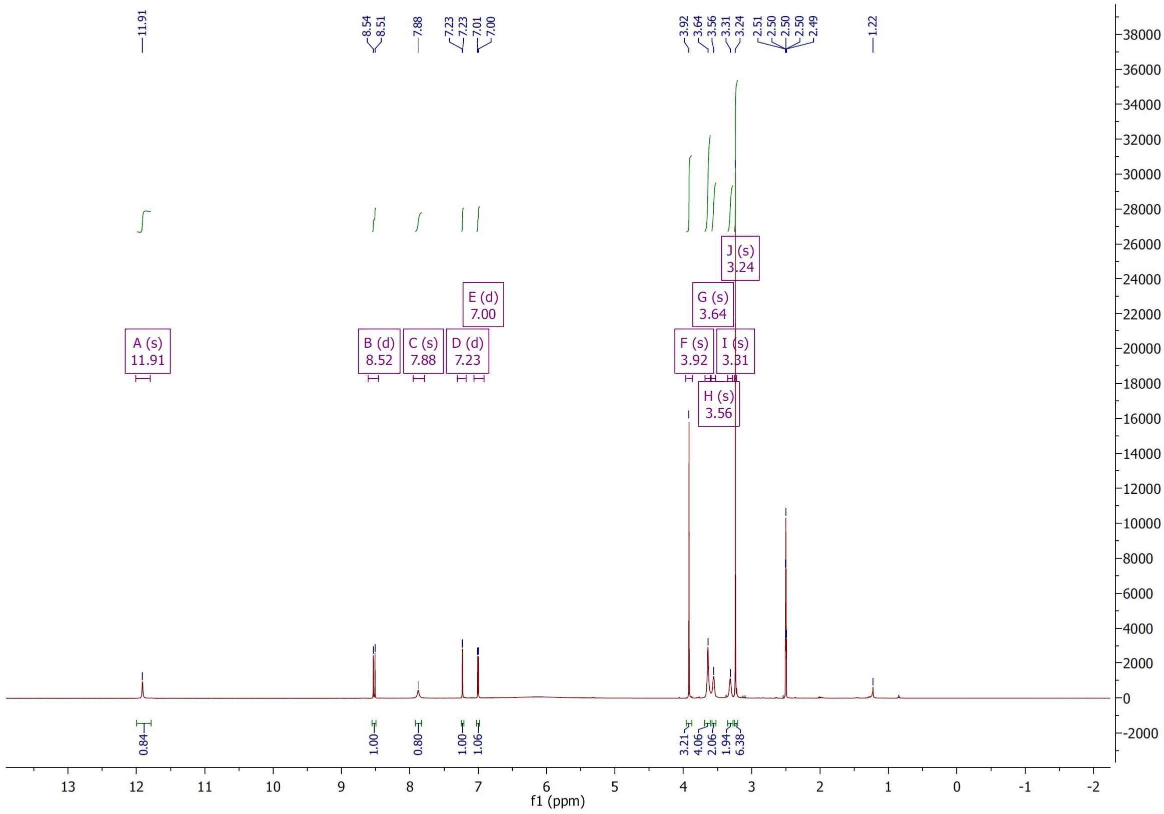


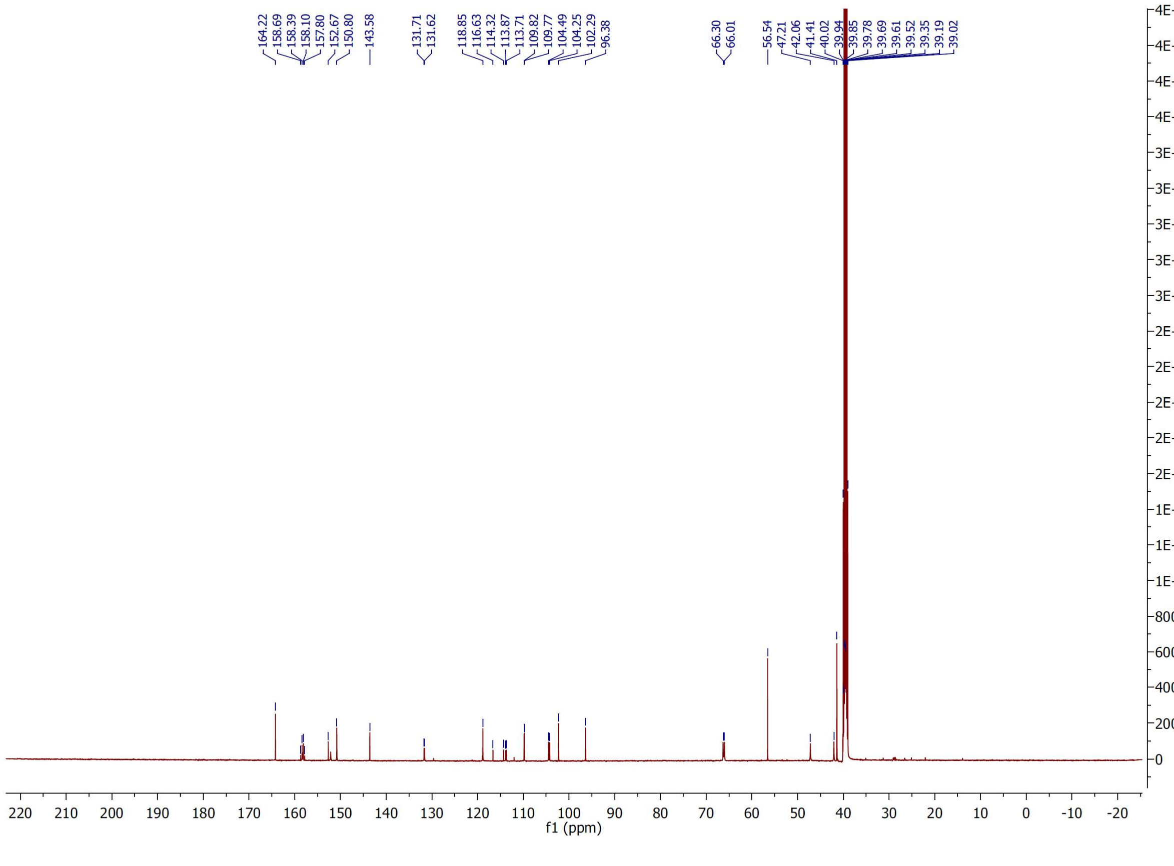

$^1H$ and $^{13}C$ NMR spectra of compound **34**

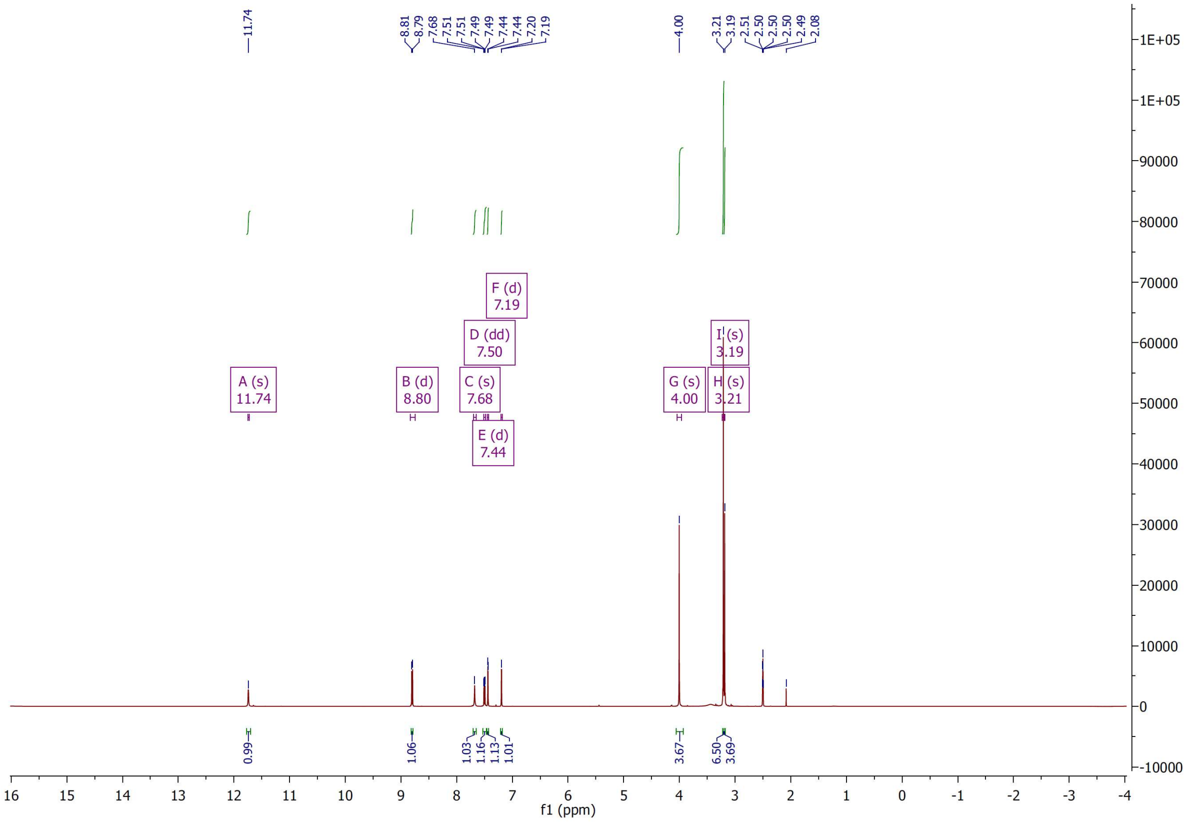


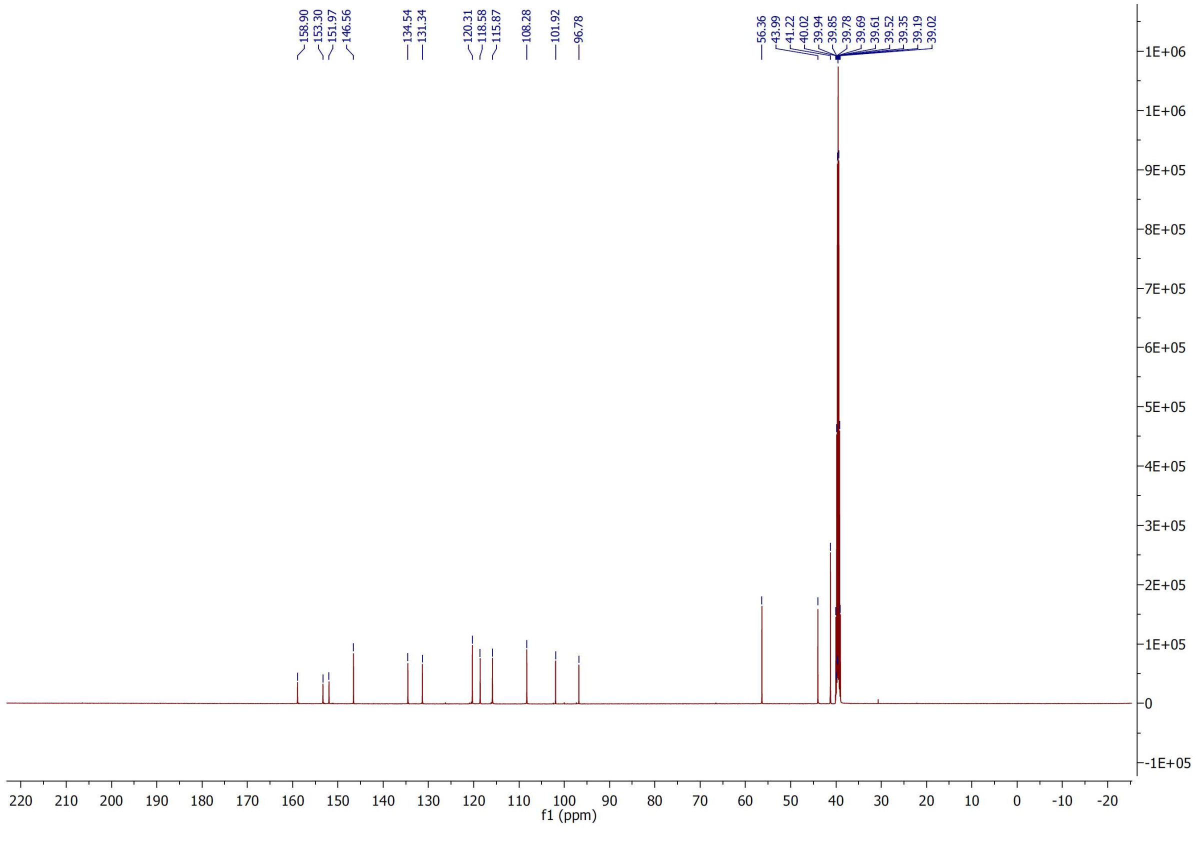

$^{1}$H and $^{13}$C NMR spectra of compound **35**

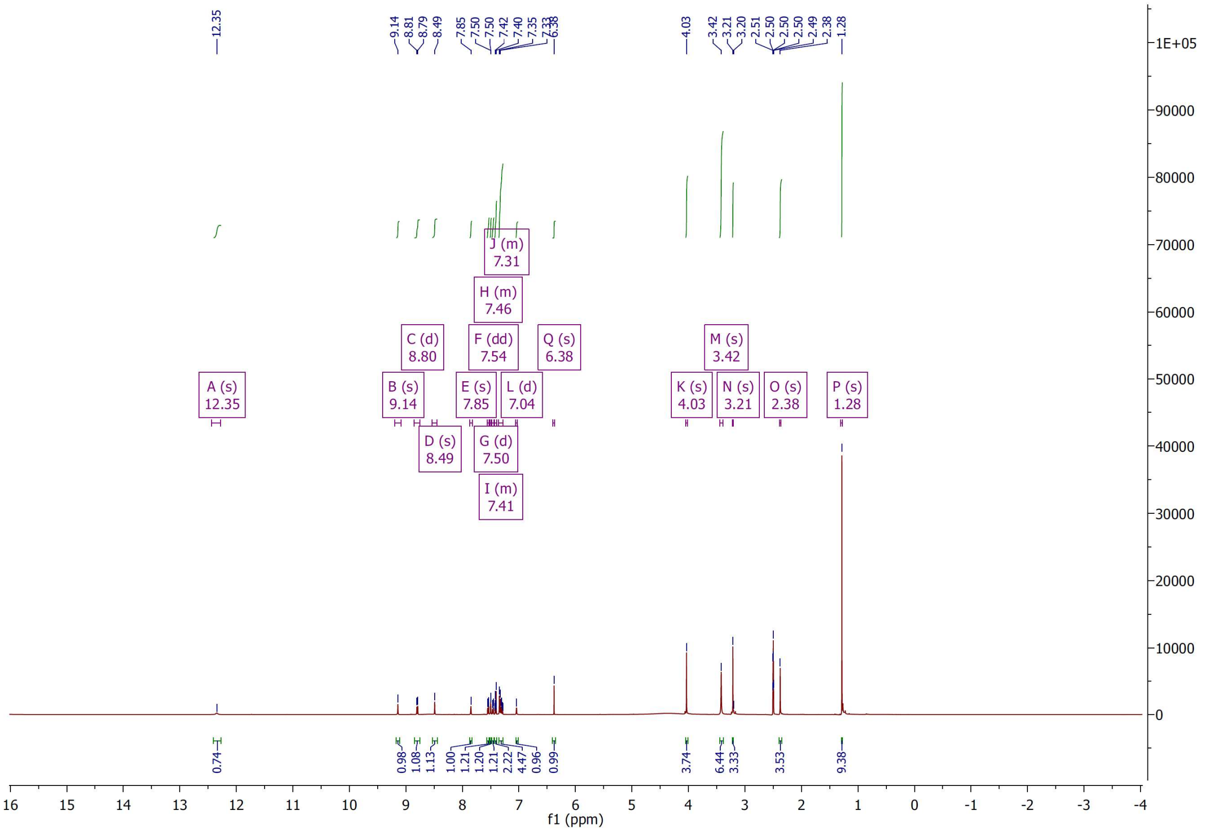

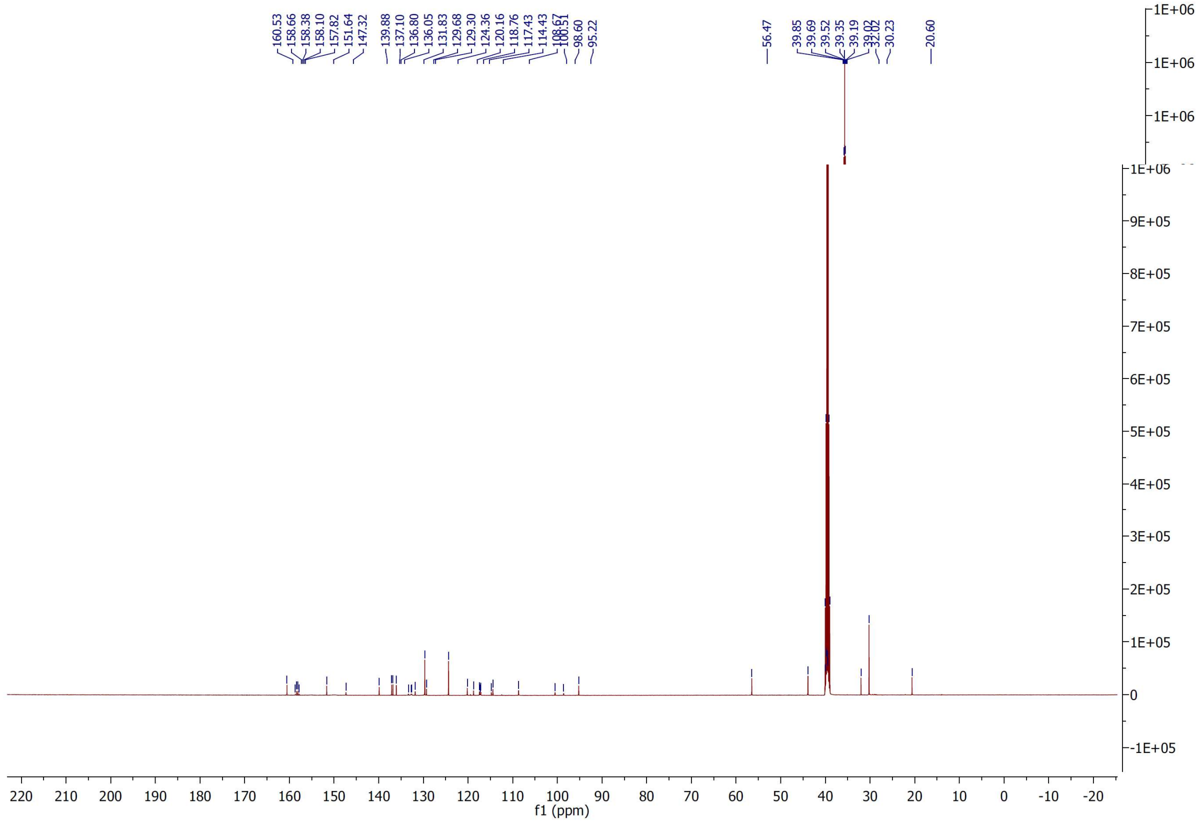

[1]H and [13]C NMR spectra of compound **36**

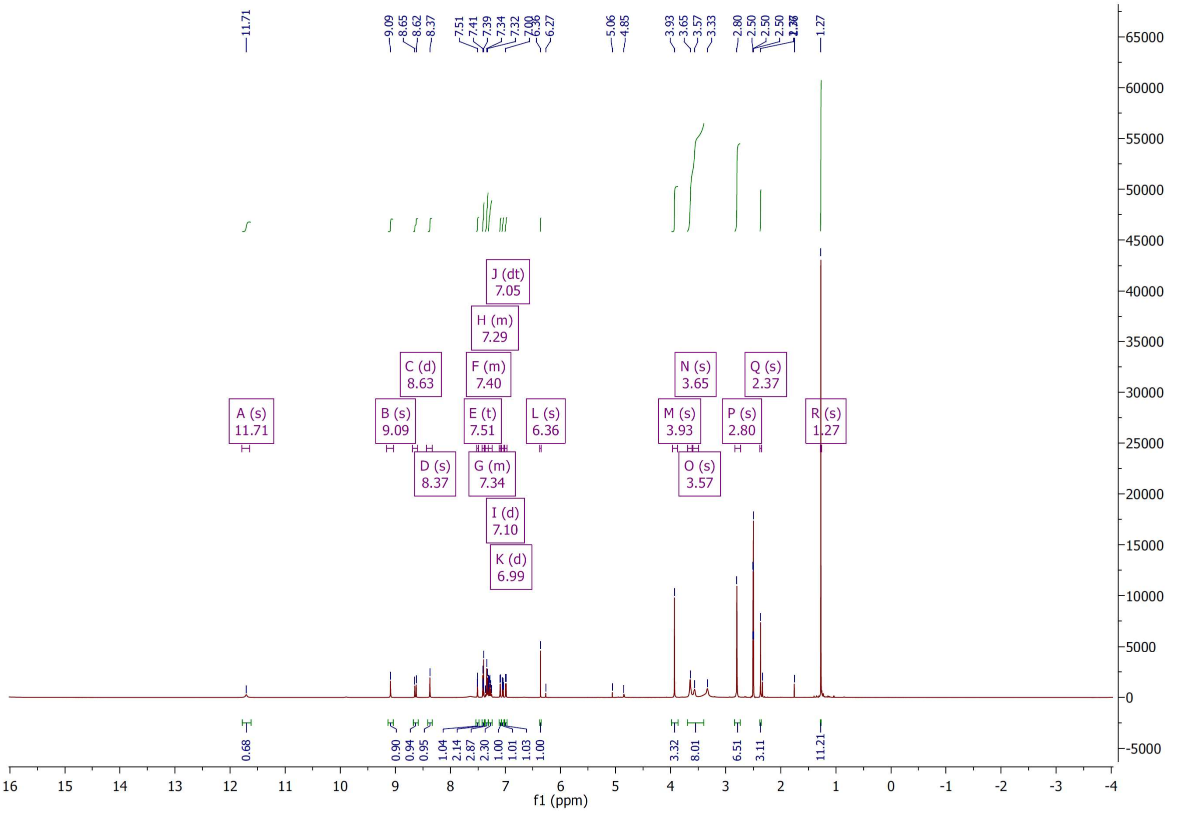


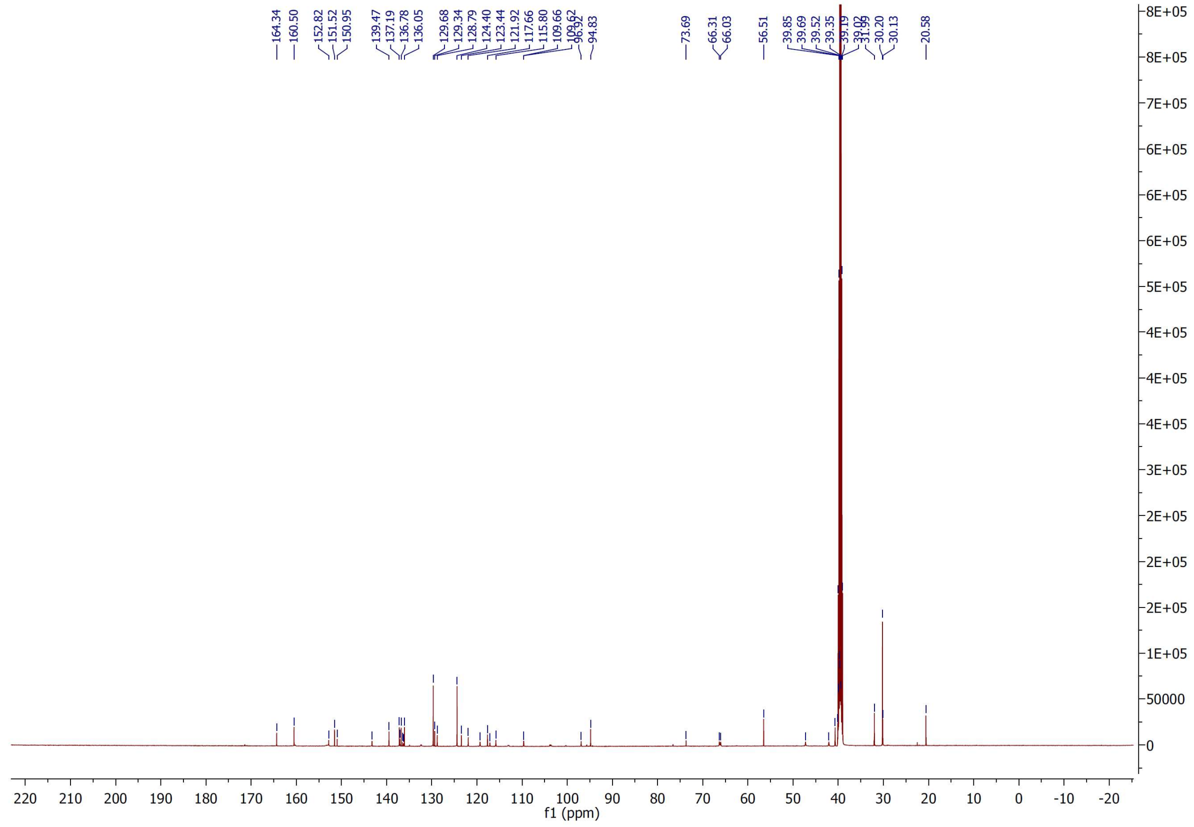

$^{1}H$ and $^{13}C$ NMR spectra of compound **37**

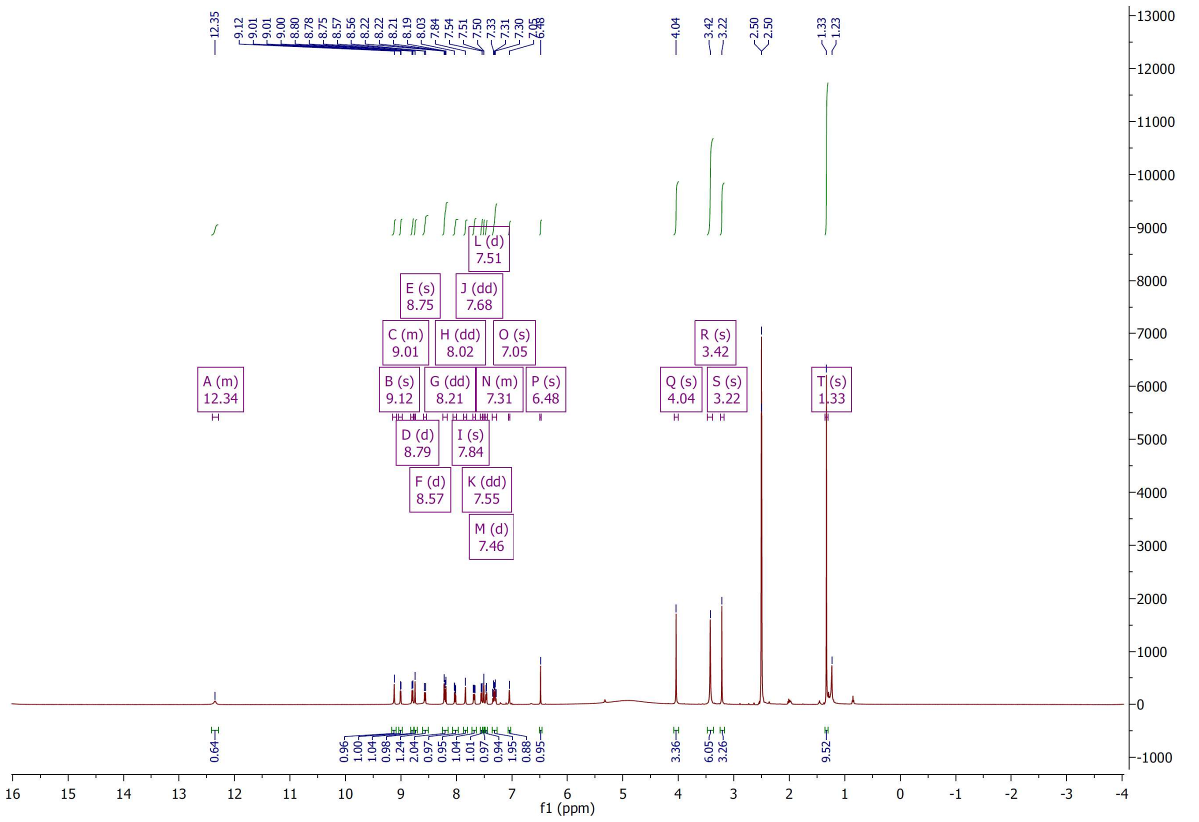

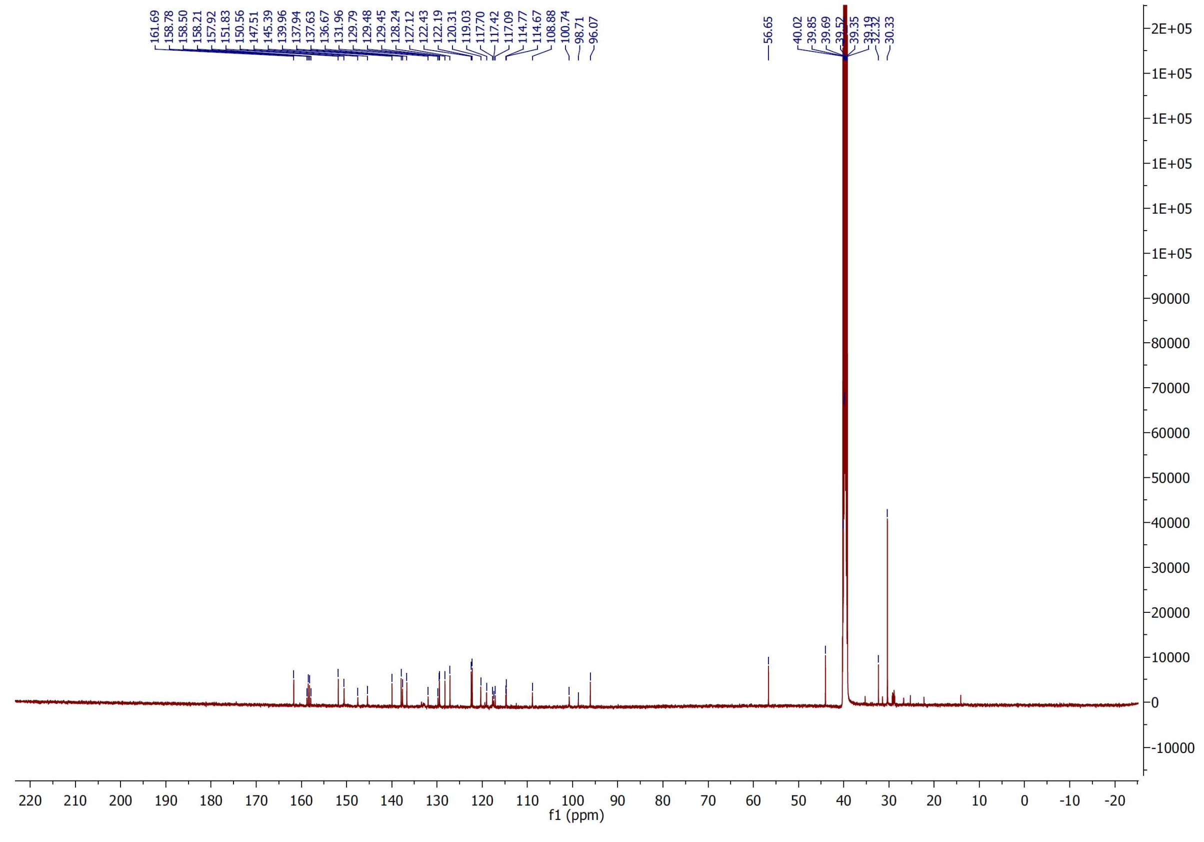

$^{1}H$ and $^{13}C$ NMR spectra of compound **47**

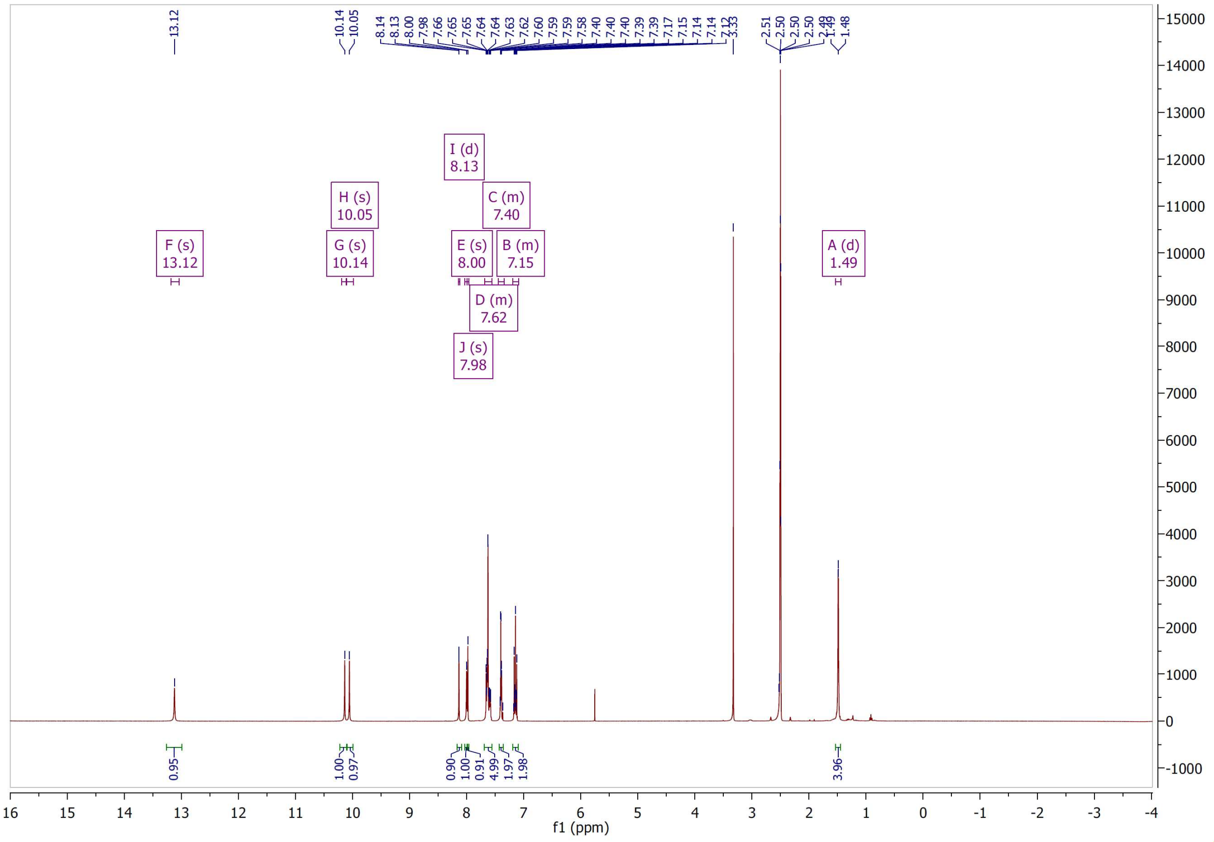

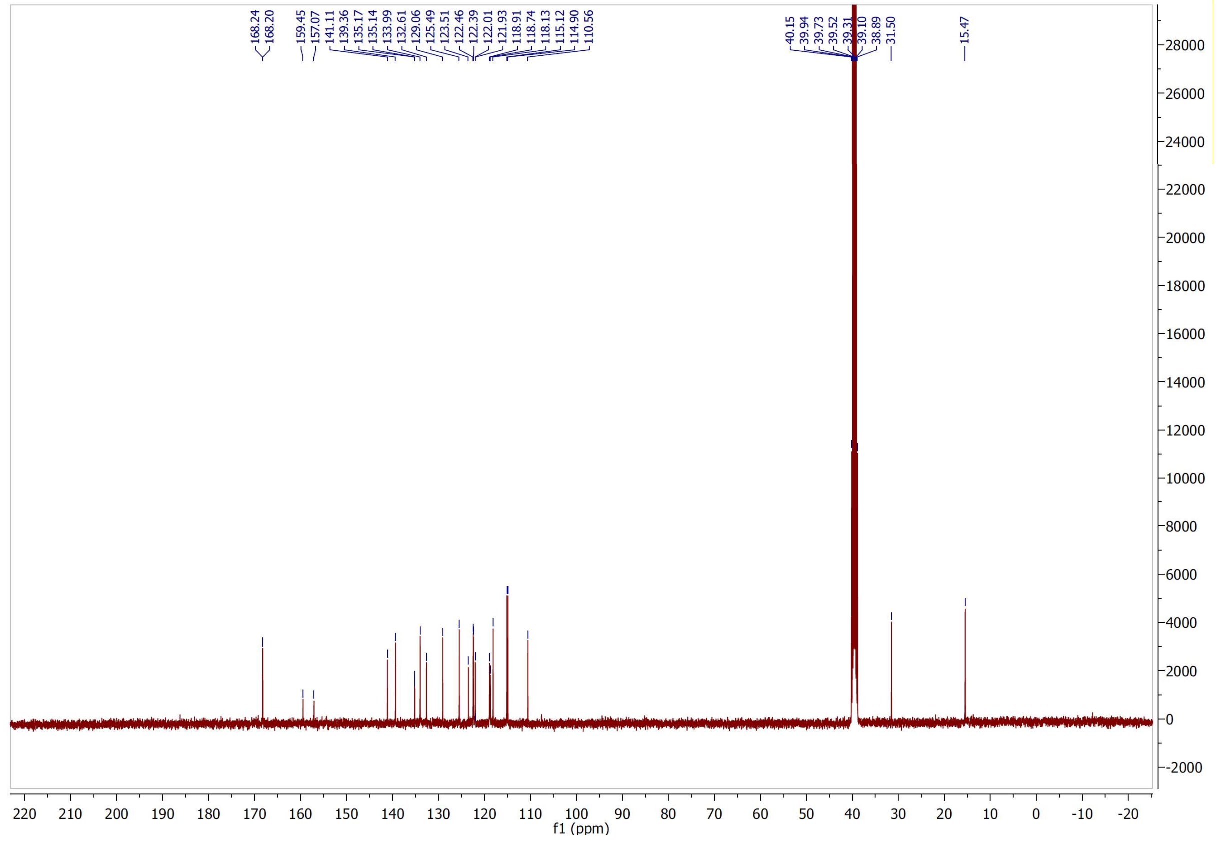

[1]H and [13]C NMR spectra of compound **48**

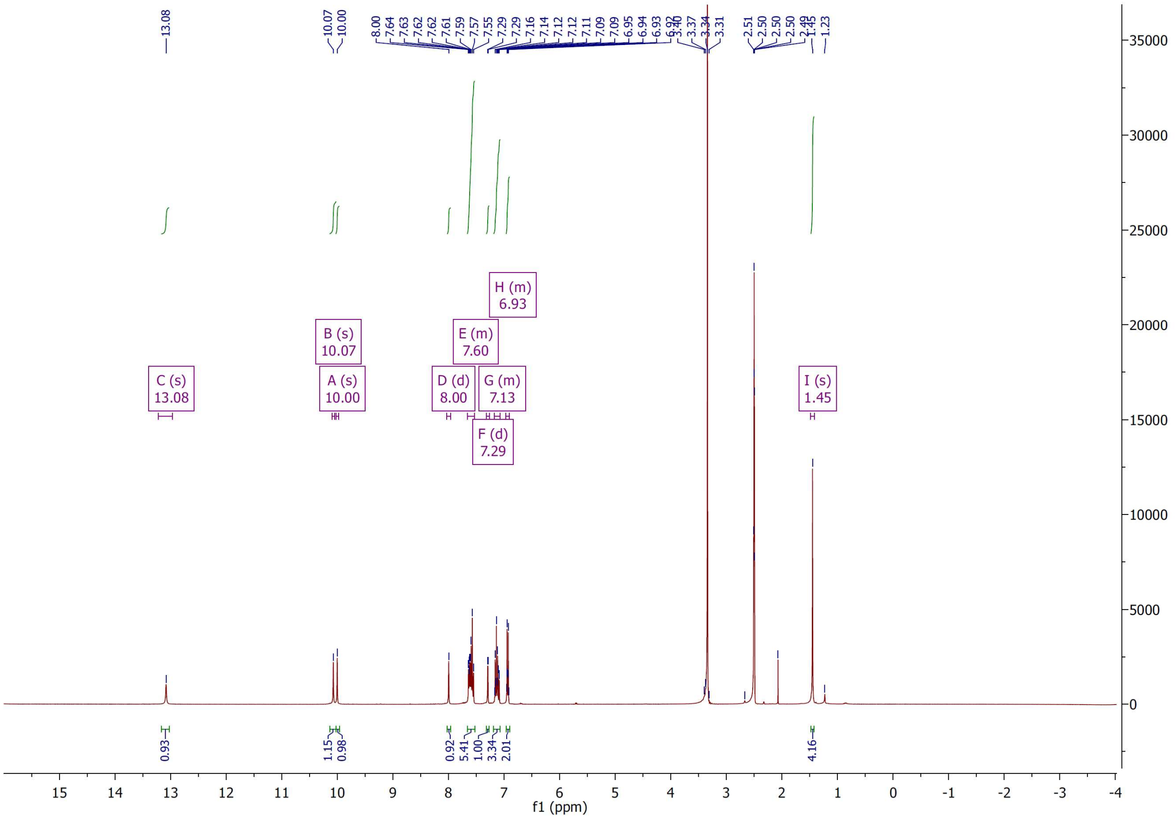

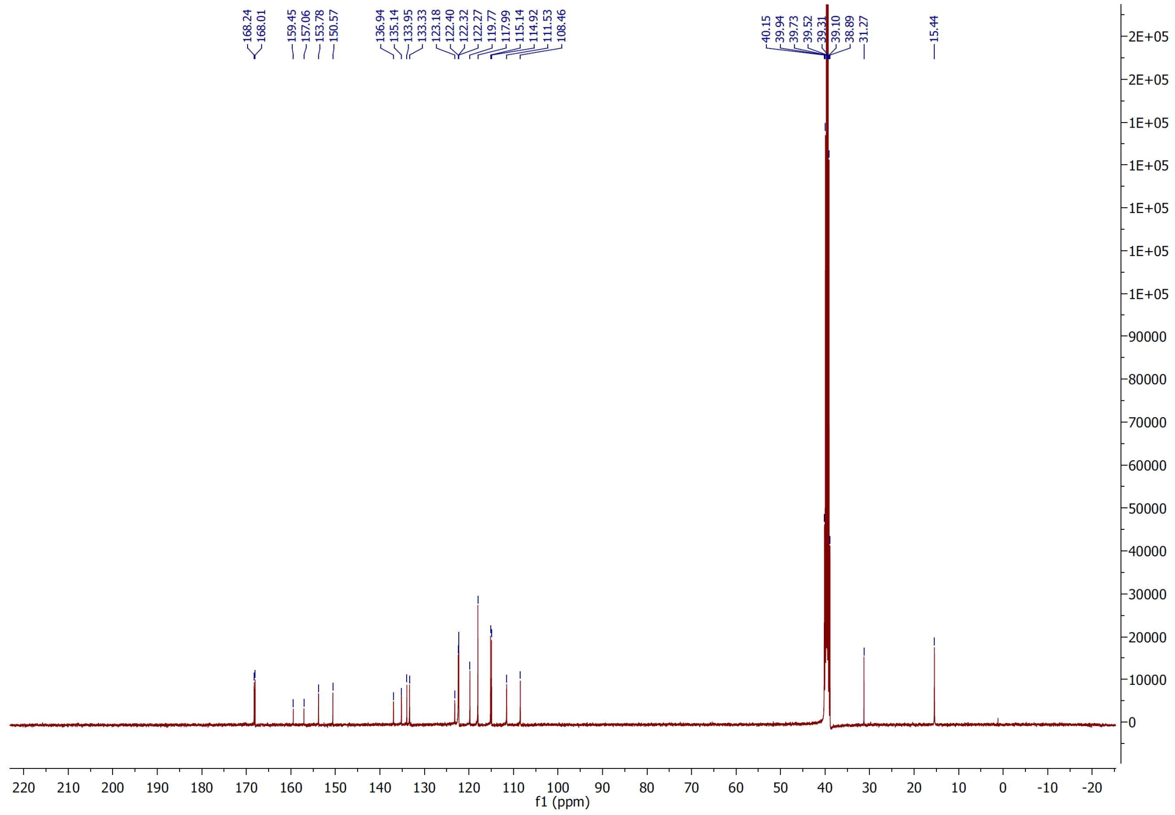

$^{1}H$ and $^{13}C$ NMR spectra of compound **49**

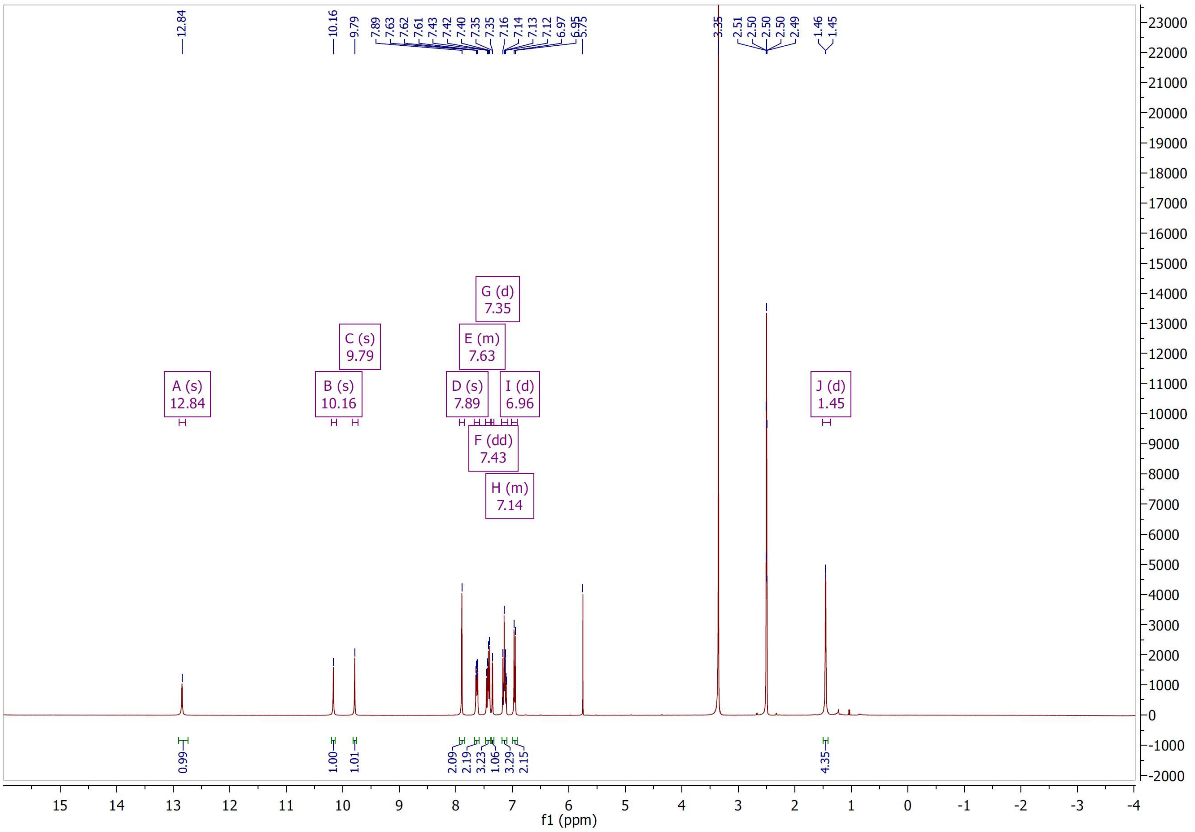


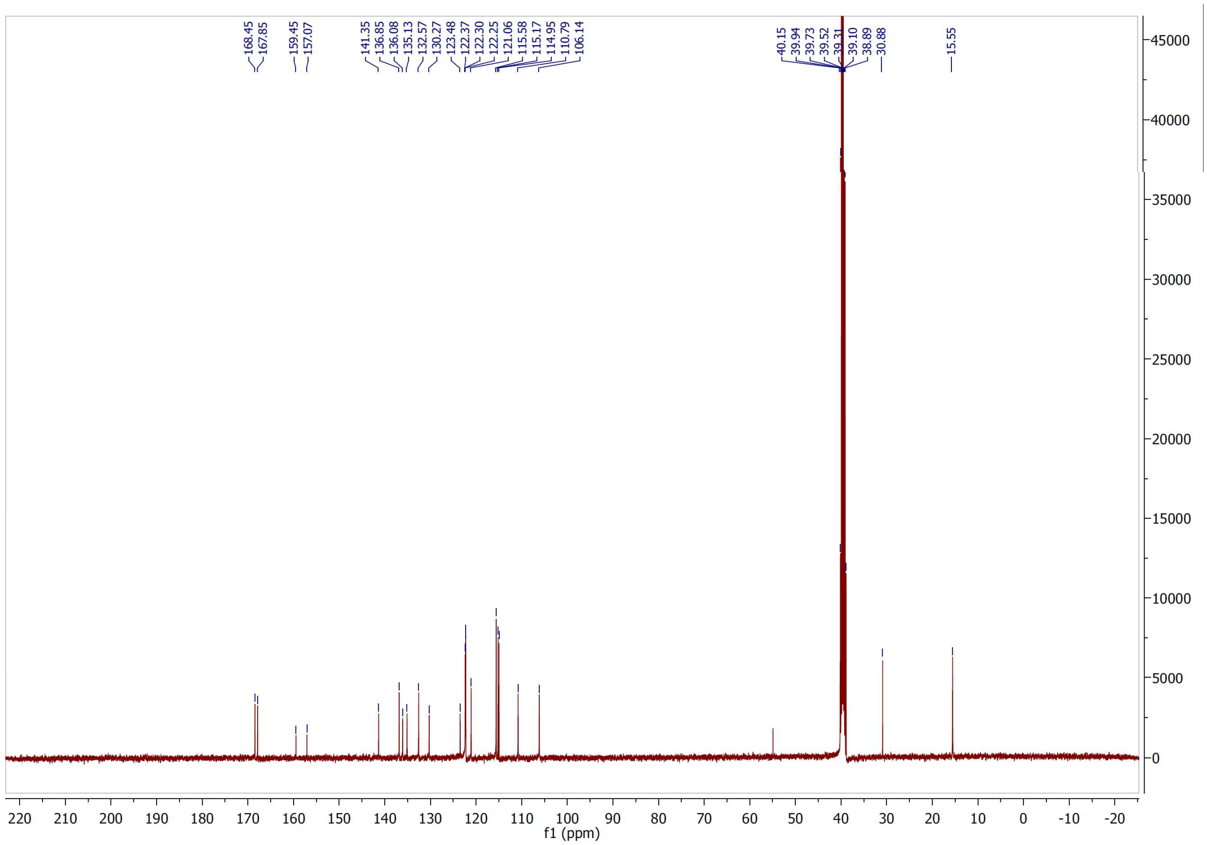

[1]H and [13]C NMR spectra of compound **50**

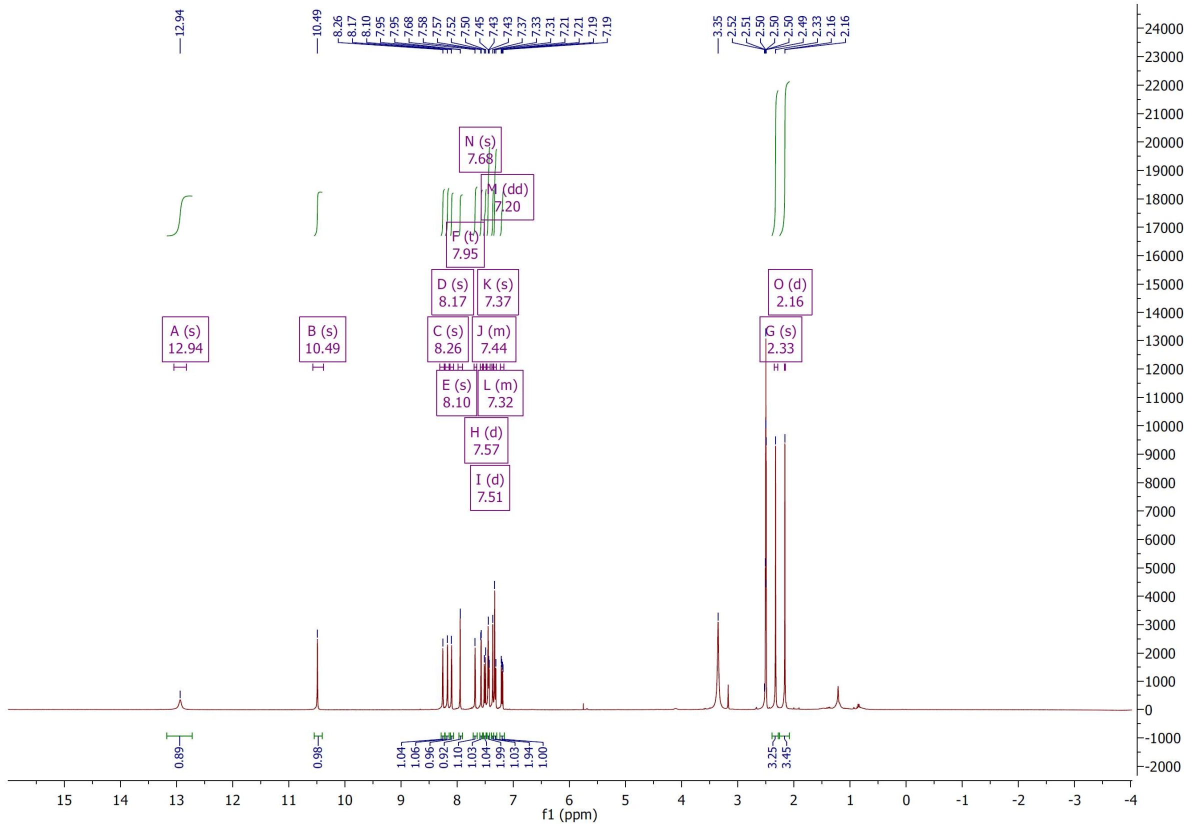

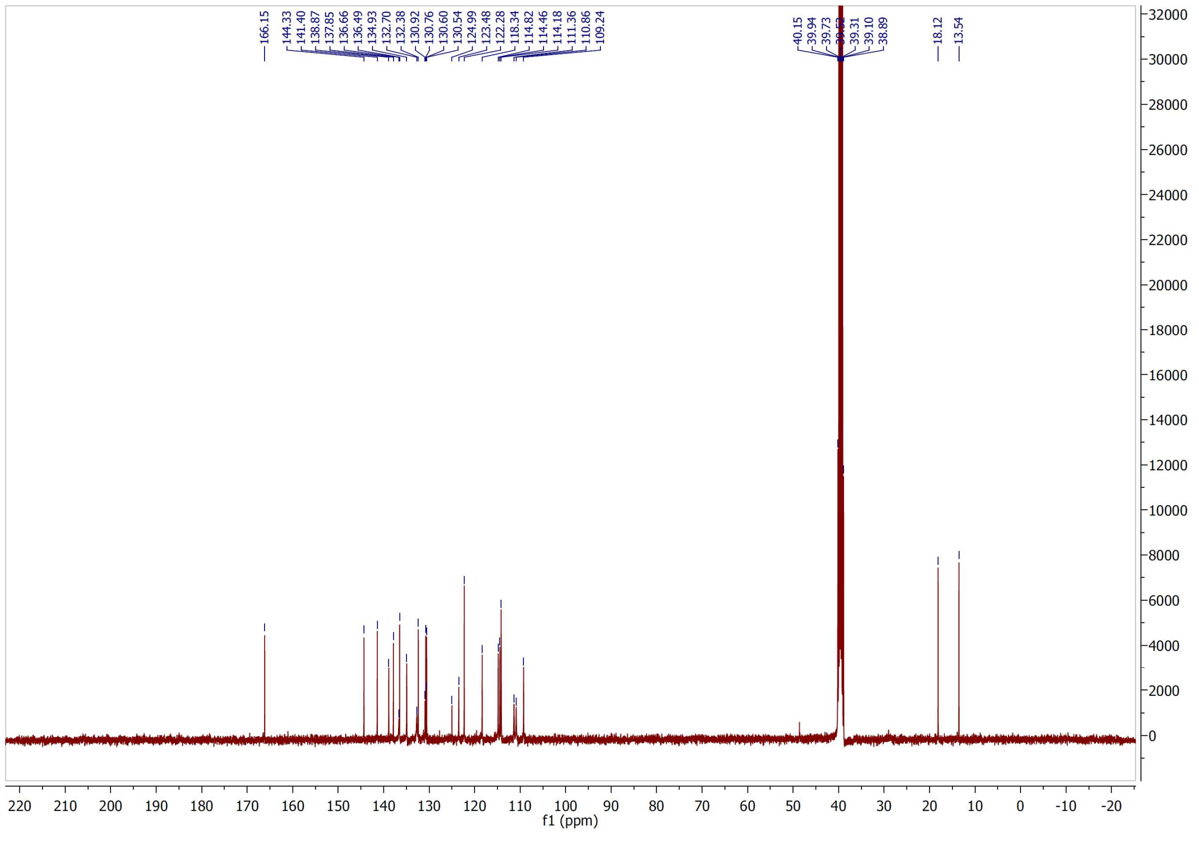

$^{1}H$ and $^{13}C$ NMR spectra of compound **51**

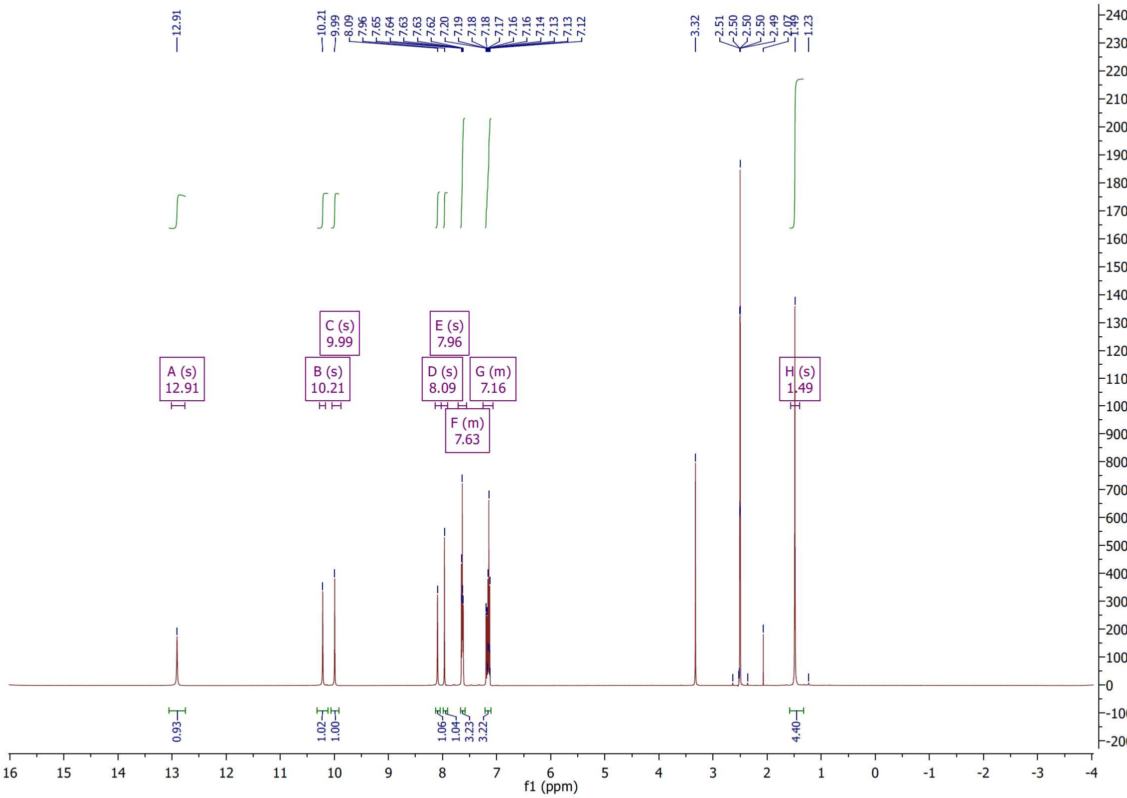

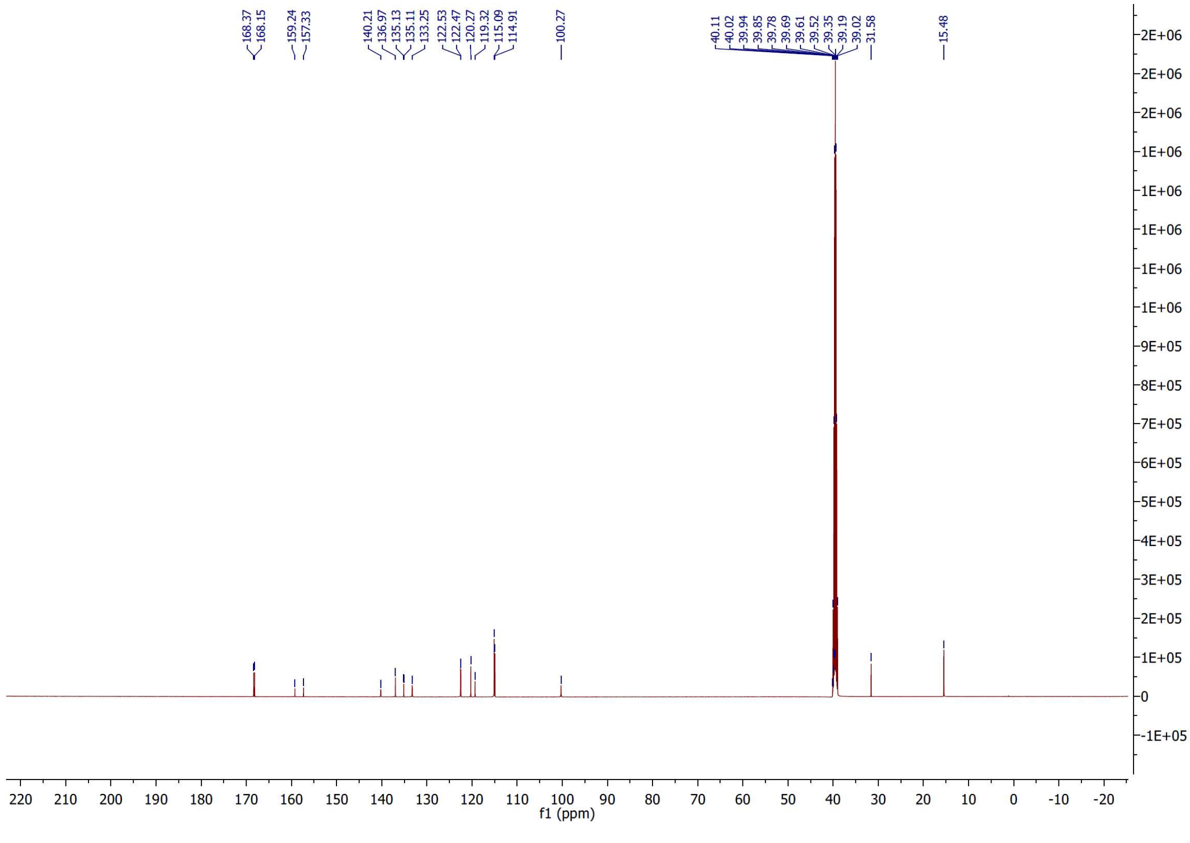

$^{1}H$ and $^{13}C$ NMR spectra of compound **52**

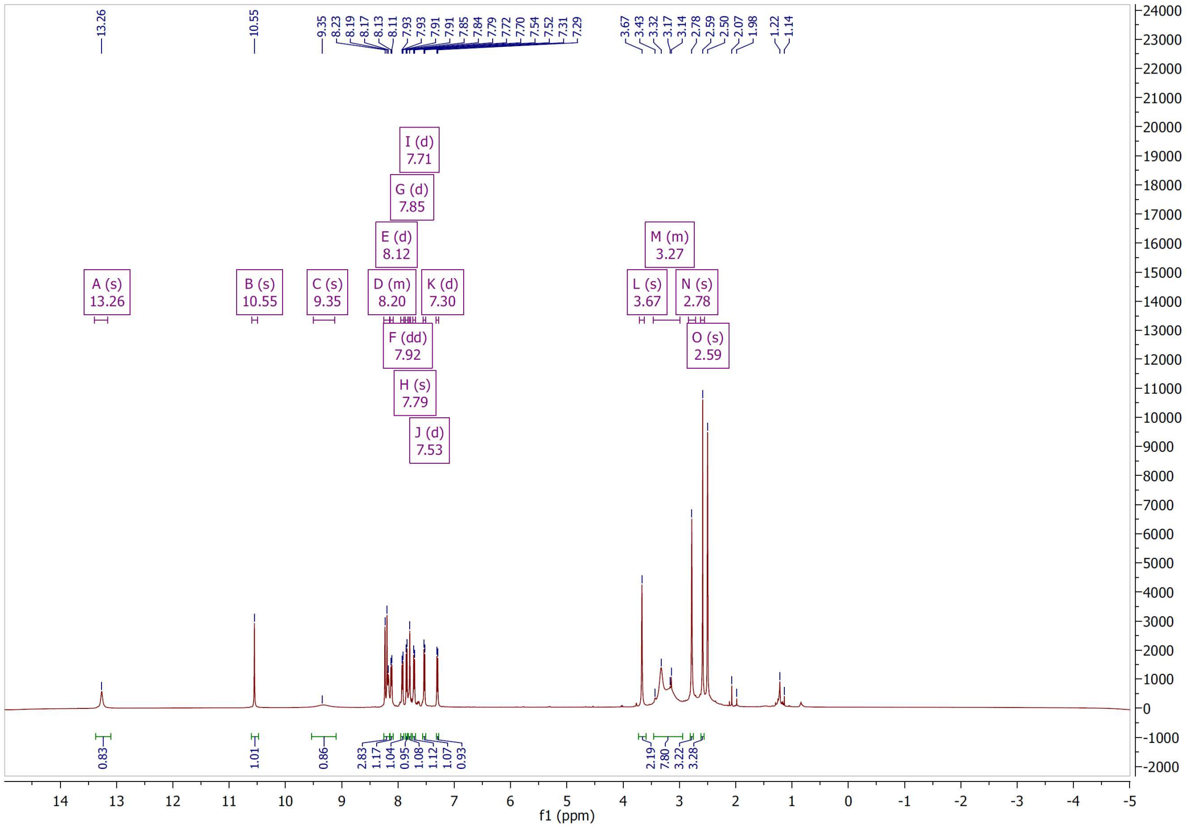

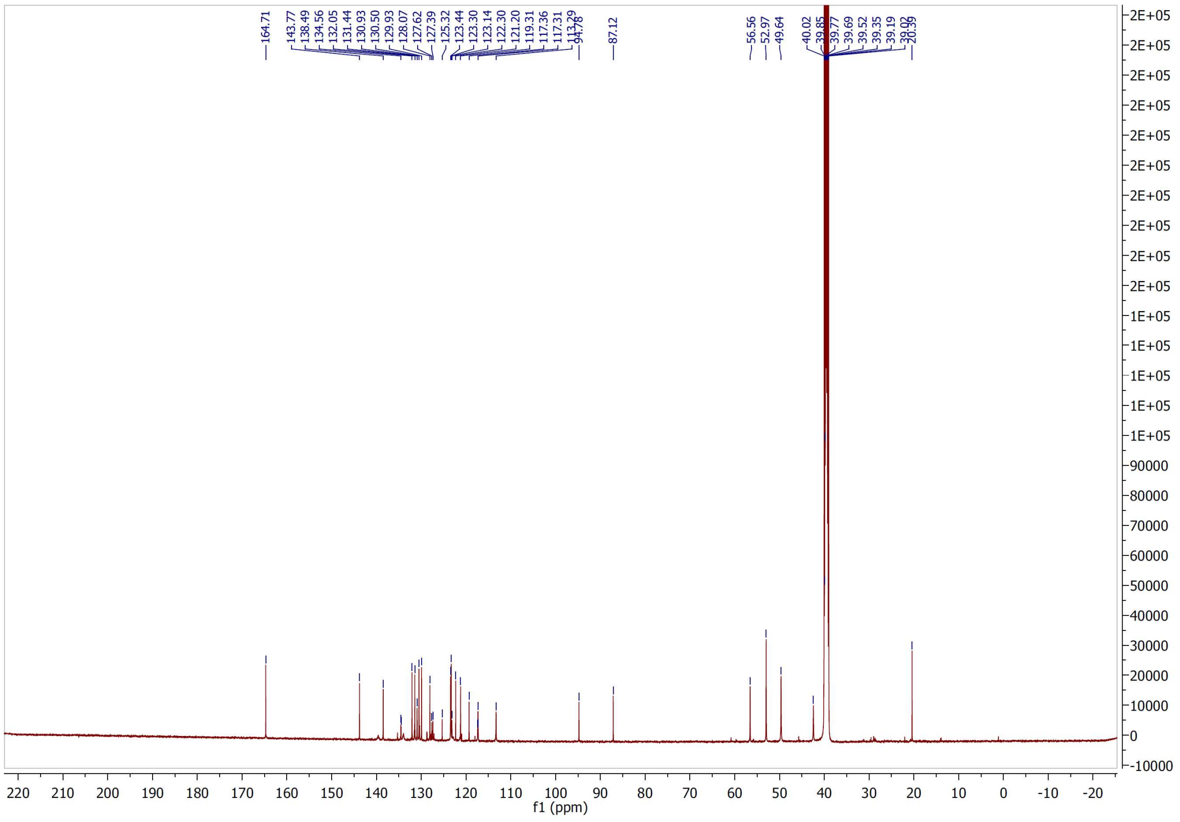

$^1H$ and $^{13}C$ NMR spectra of compound **53**

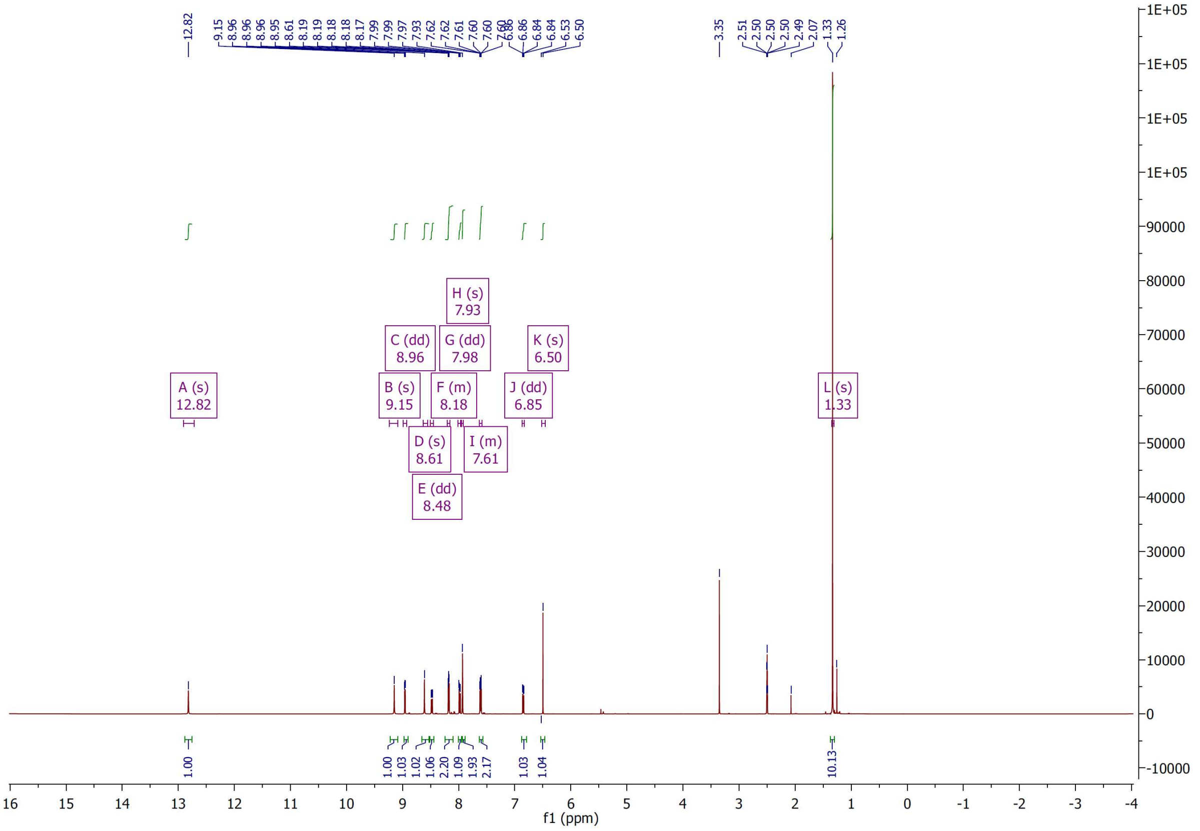

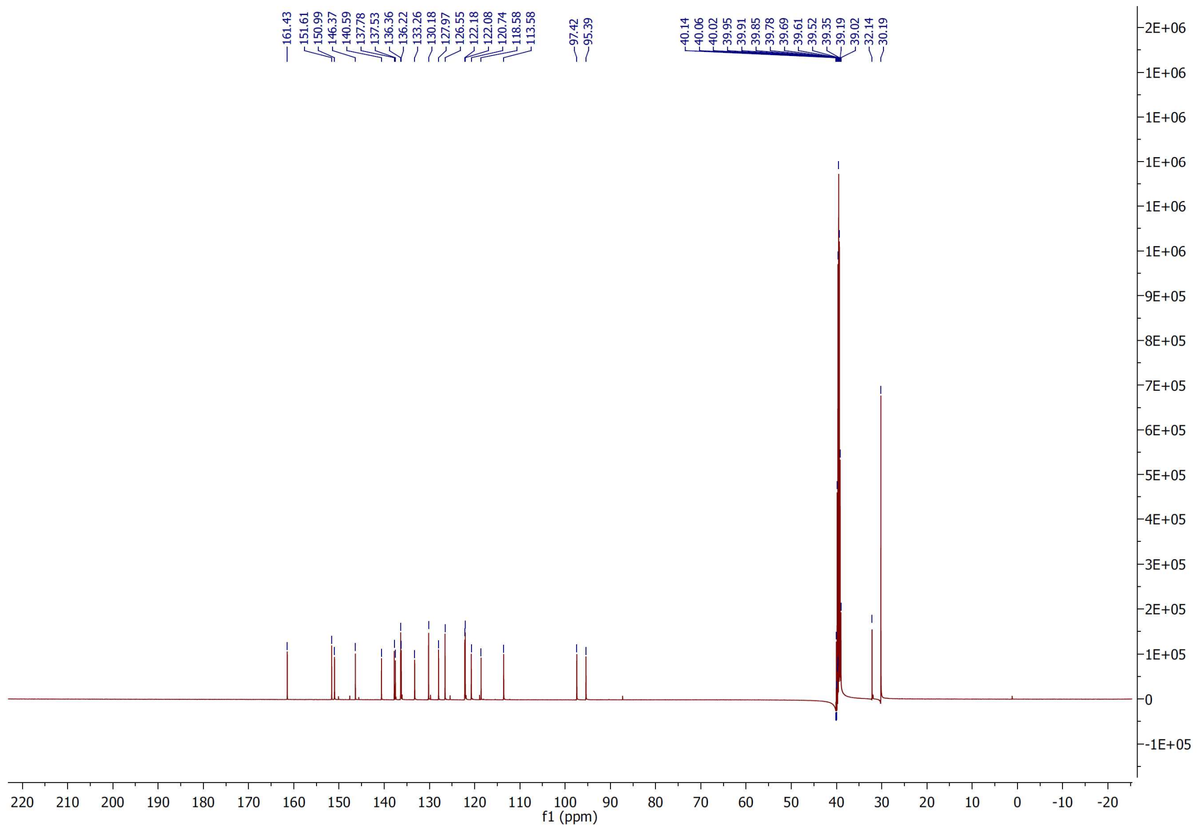

<sup></sup>

[1]H and [13]C NMR spectra of compound **54**

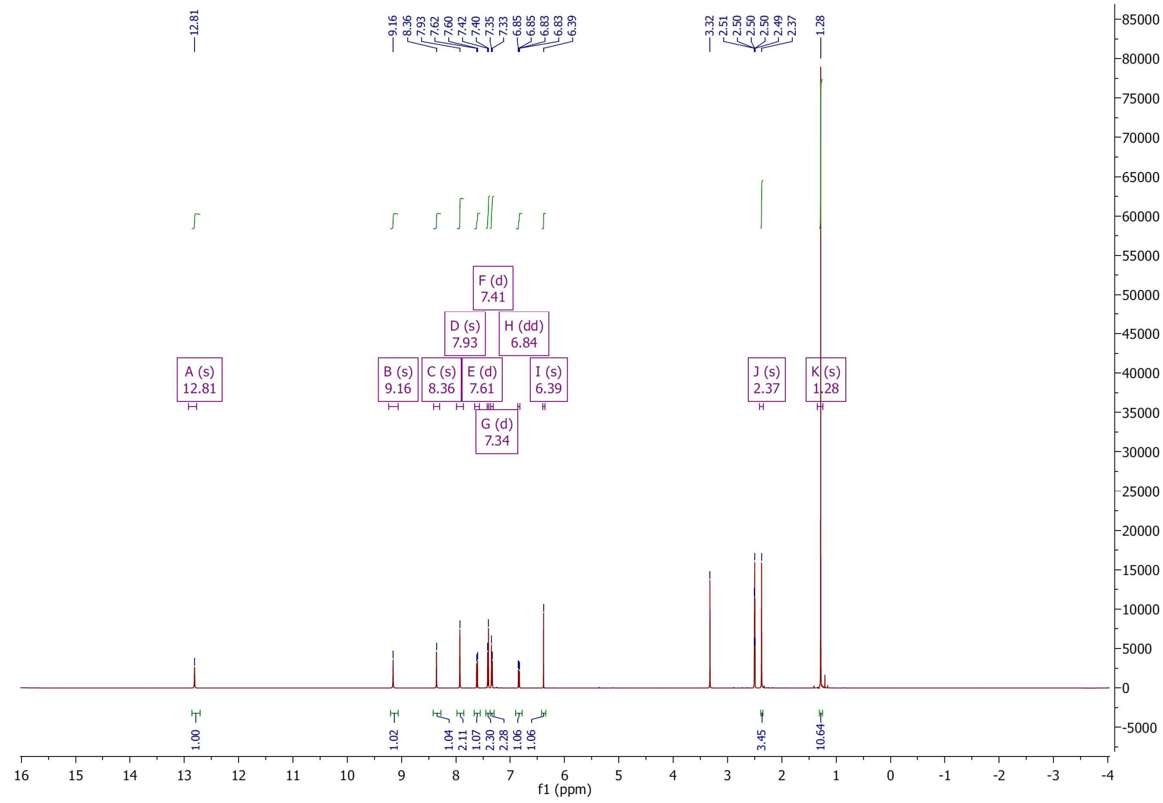

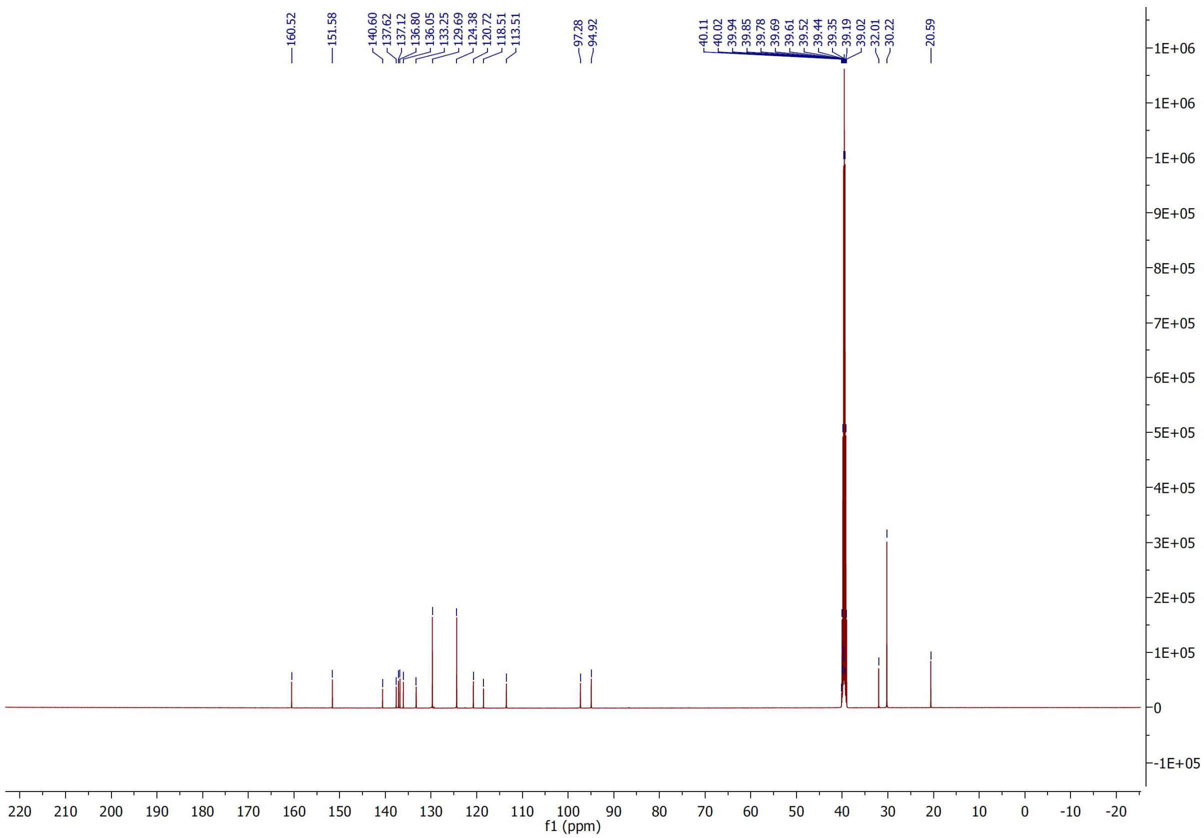

### HPLC/MS

HPLC purity and mass spectrum (ESI) **1**

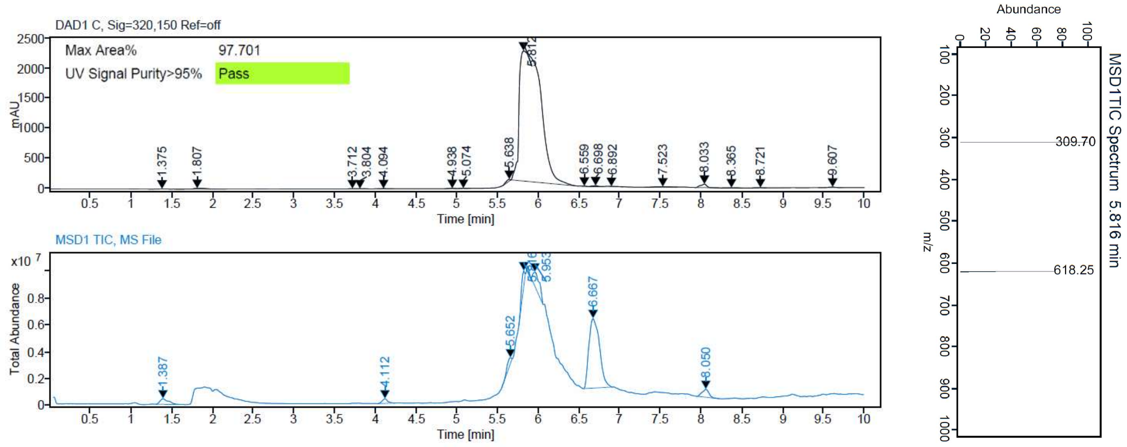


HPLC purity and mass spectrum (ESI) **2**

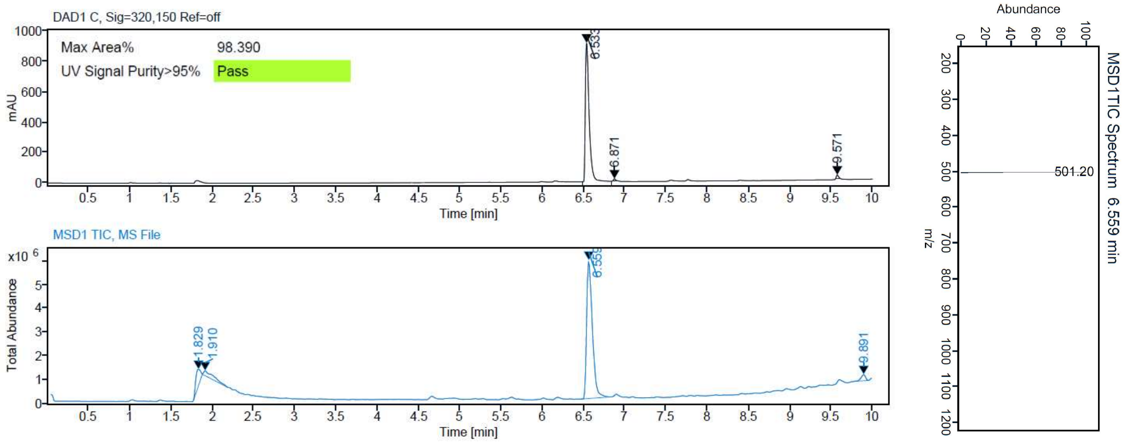


HPLC purity and mass spectrum (ESI) **3**

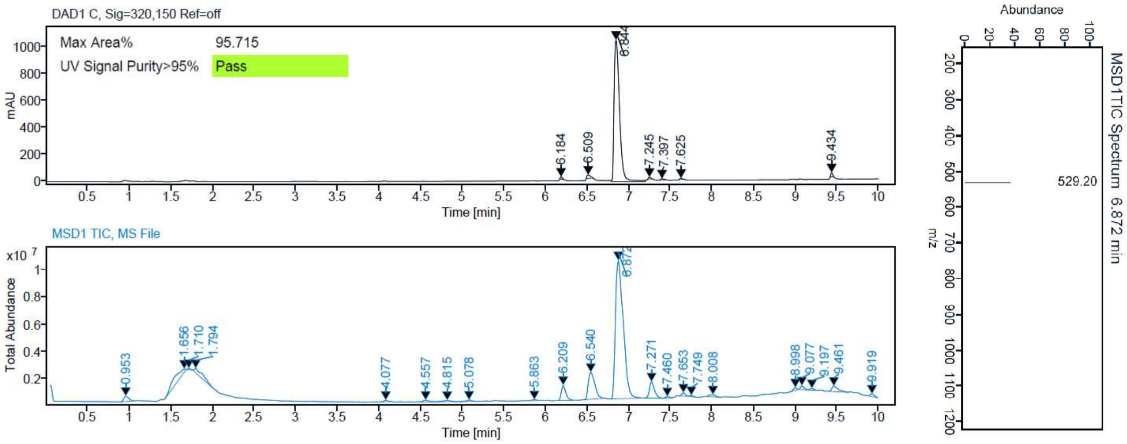

HPLC purity and mass spectrum (ESI) **4**

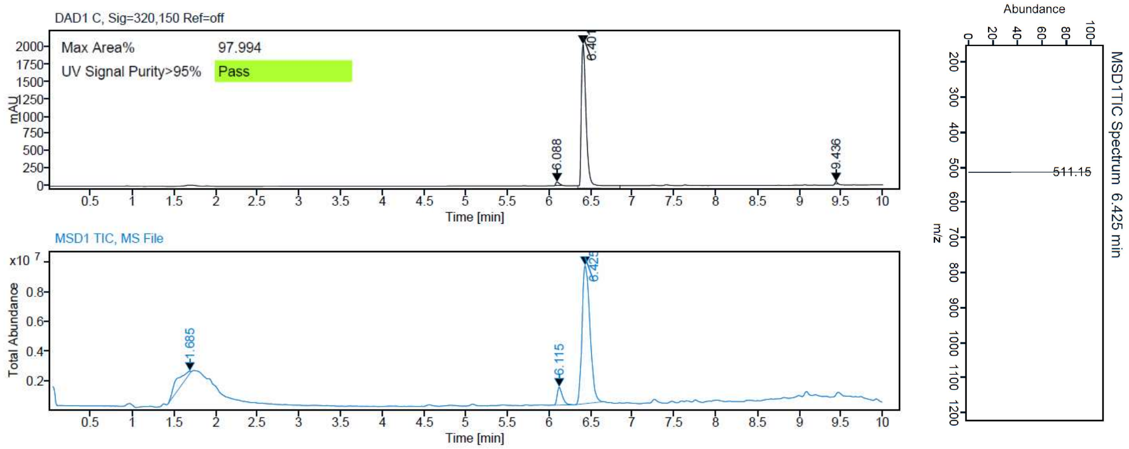


HPLC purity and mass spectrum (ESI) **5**

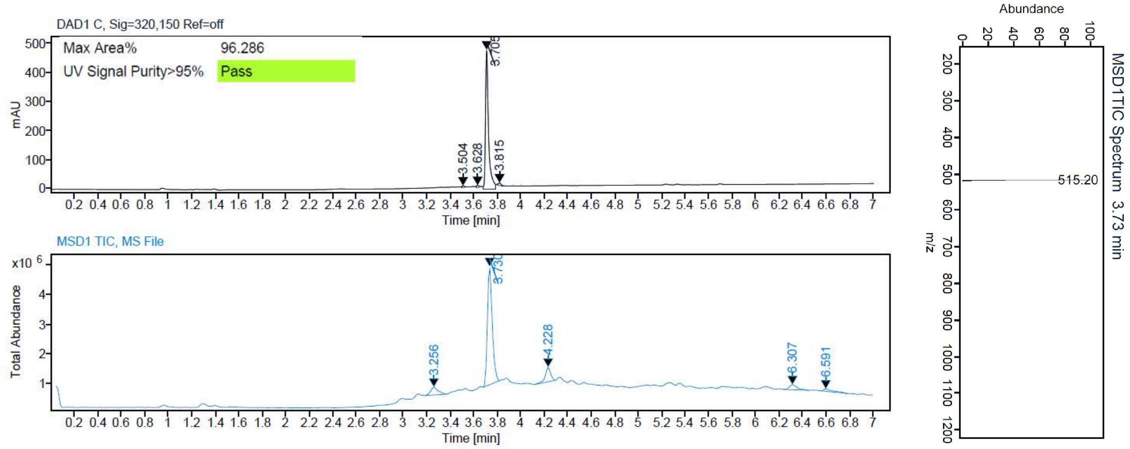


HPLC purity and mass spectrum (ESI) **6**

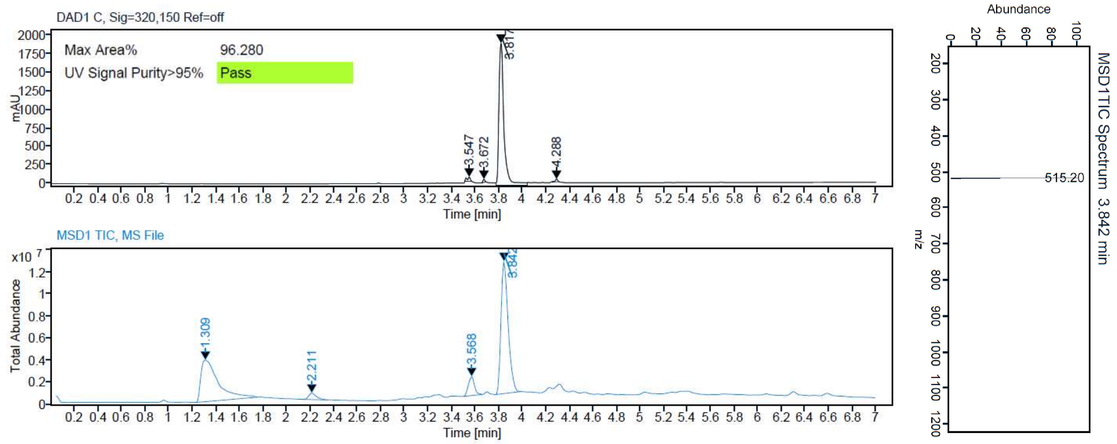

HPLC purity and mass spectrum (ESI) **7**

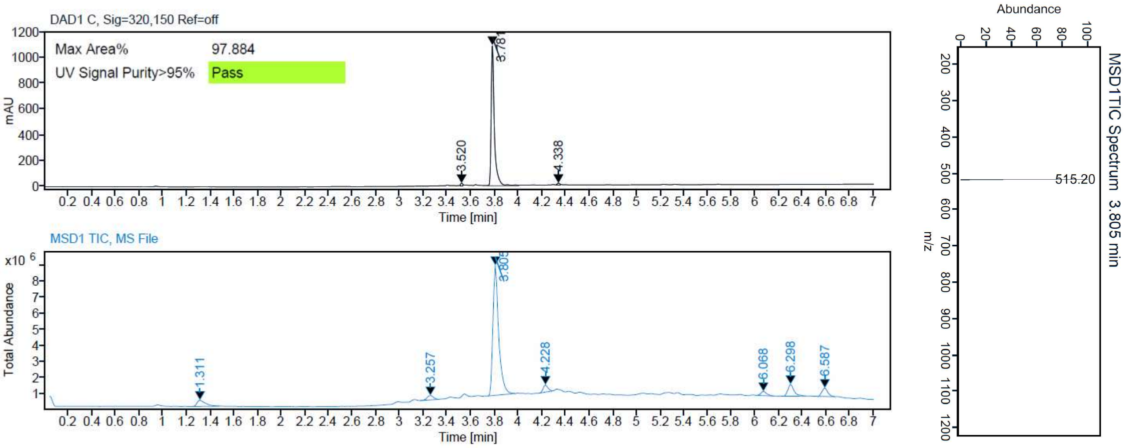


HPLC purity and mass spectrum (ESI) **8**

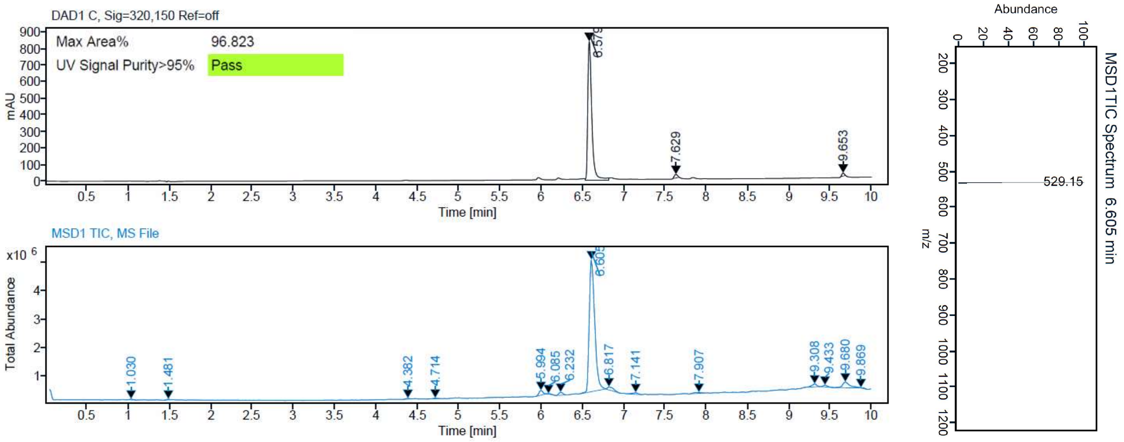


HPLC purity and mass spectrum (ESI) **9**

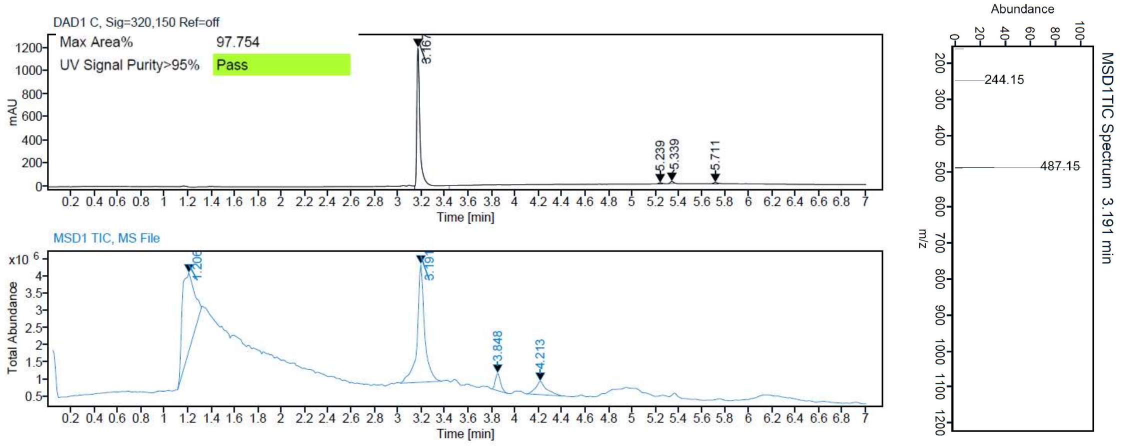

HPLC purity and mass spectrum (ESI) **10**

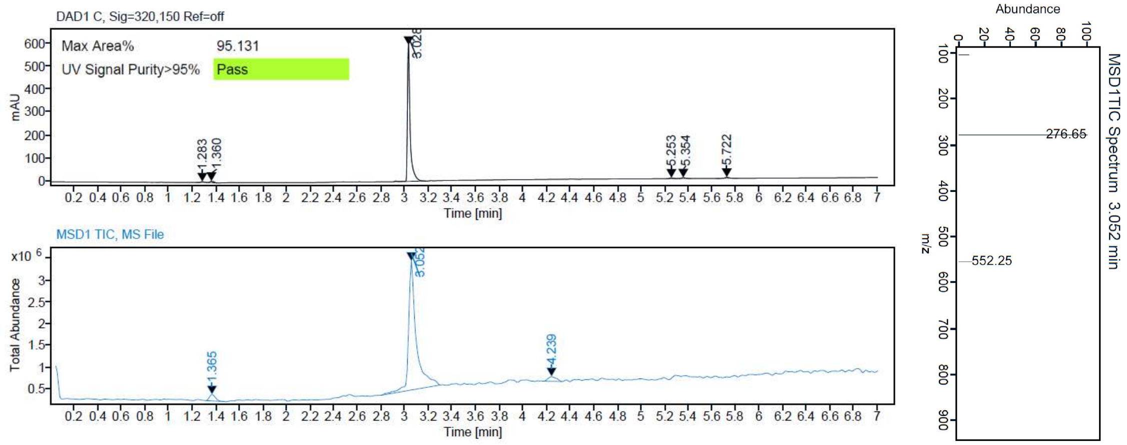


HPLC purity and mass spectrum (ESI) **11**

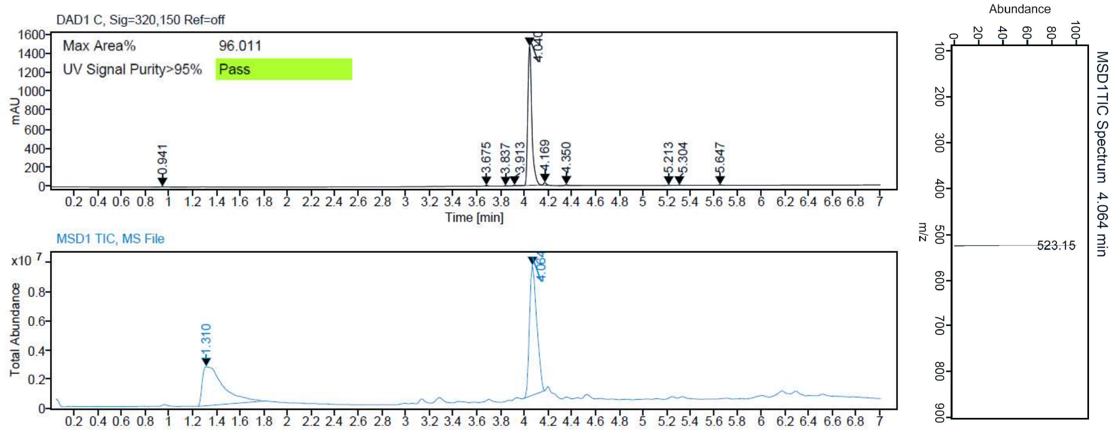


HPLC purity and mass spectrum (ESI) **12**

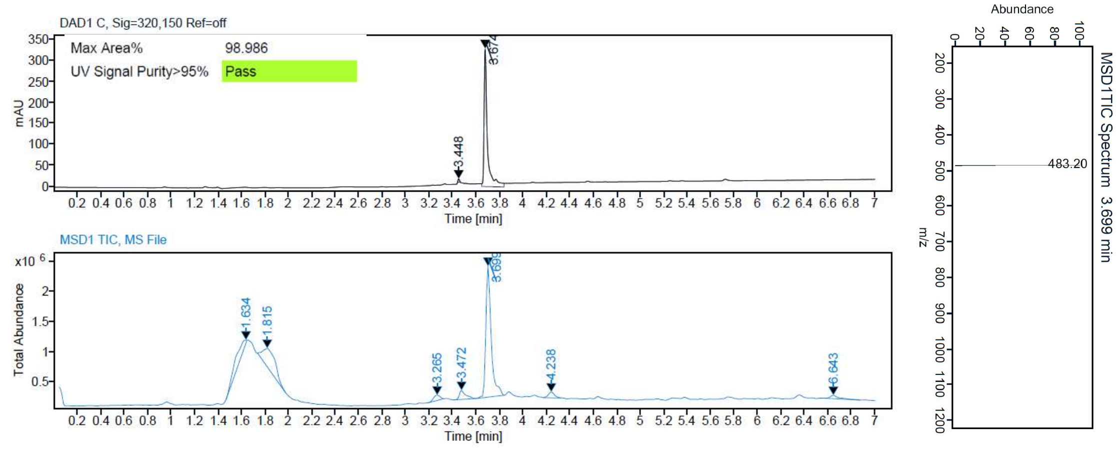

HPLC purity and mass spectrum (ESI) **13**

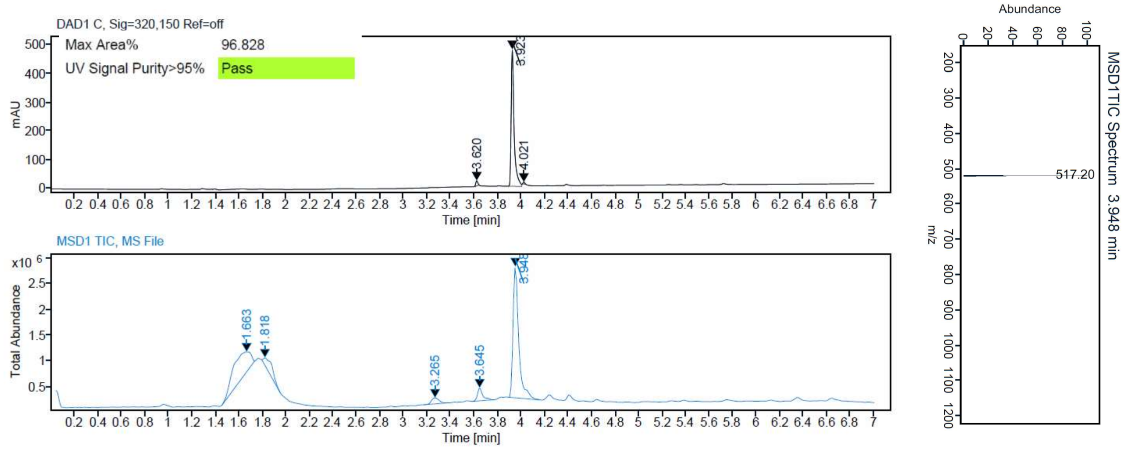


HPLC purity and mass spectrum (ESI) **14**

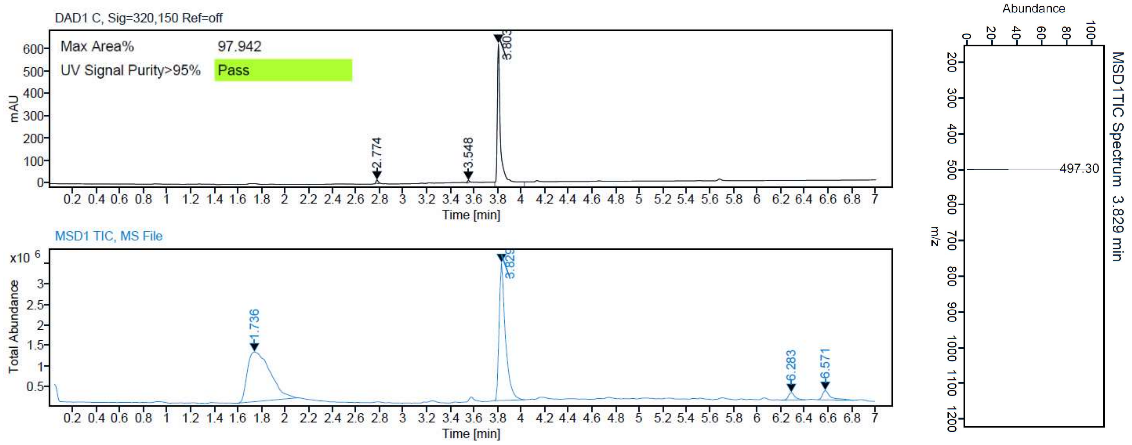


HPLC purity and mass spectrum (ESI) **15**

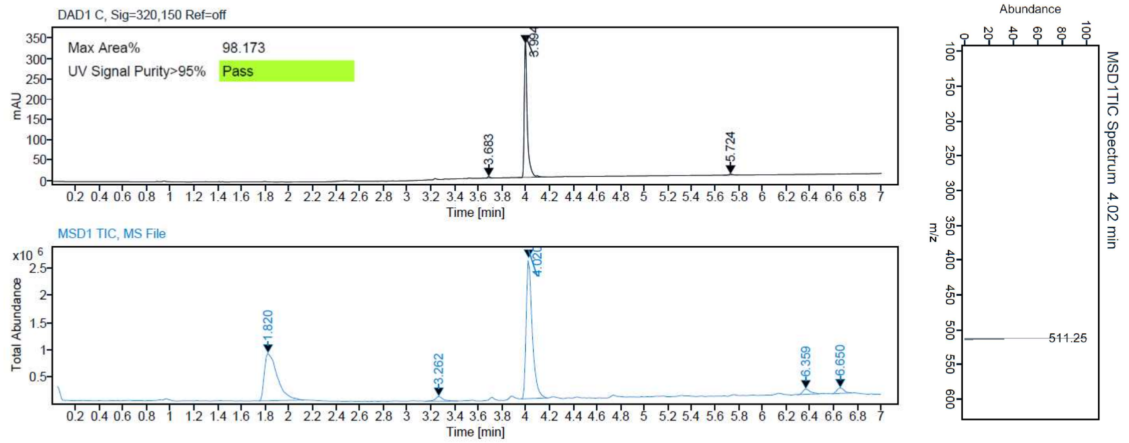

HPLC purity and mass spectrum (ESI) **16**

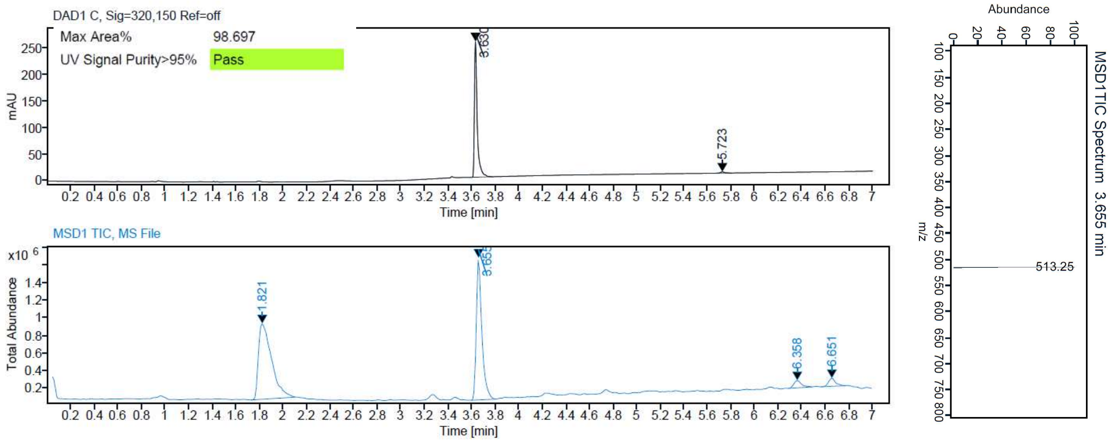


HPLC purity and mass spectrum (ESI) **17**

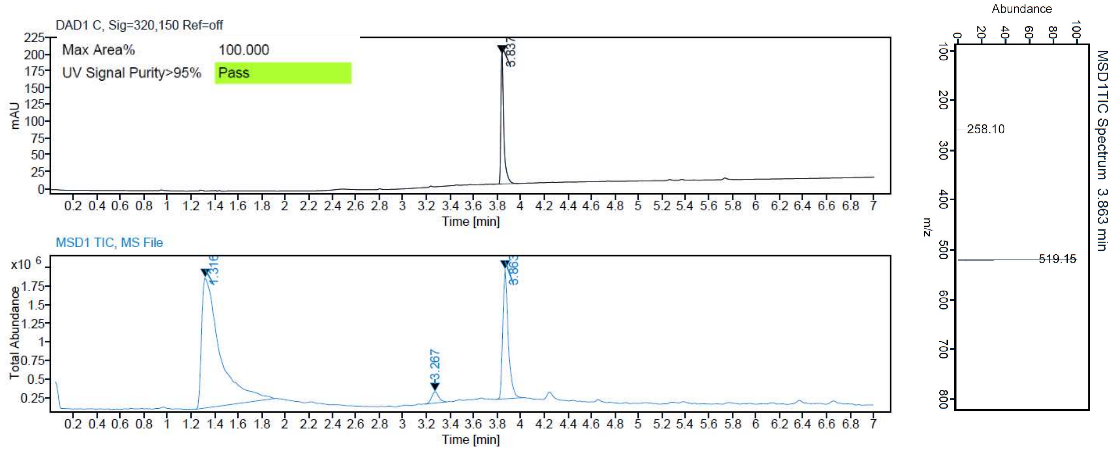


HPLC purity and mass spectrum (ESI) **18**

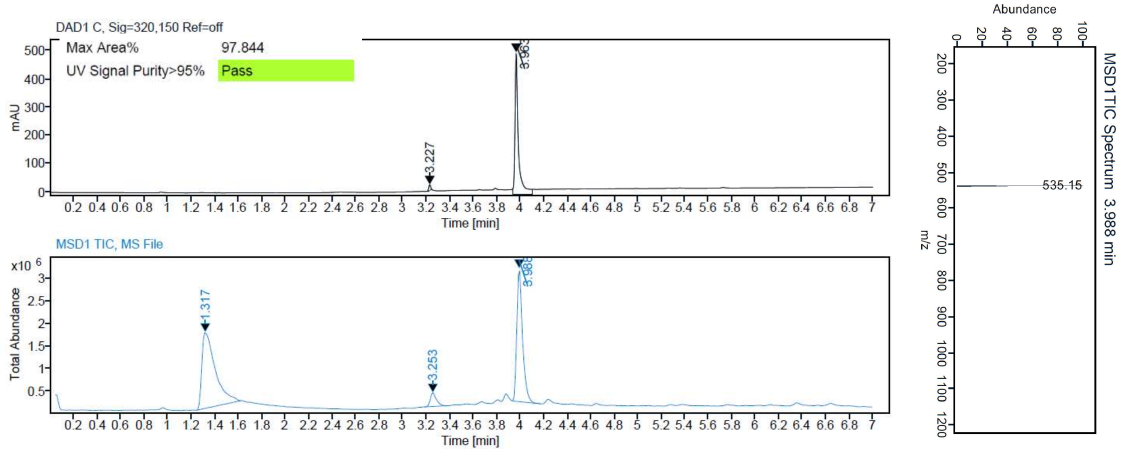

HPLC purity and mass spectrum (ESI) **19**

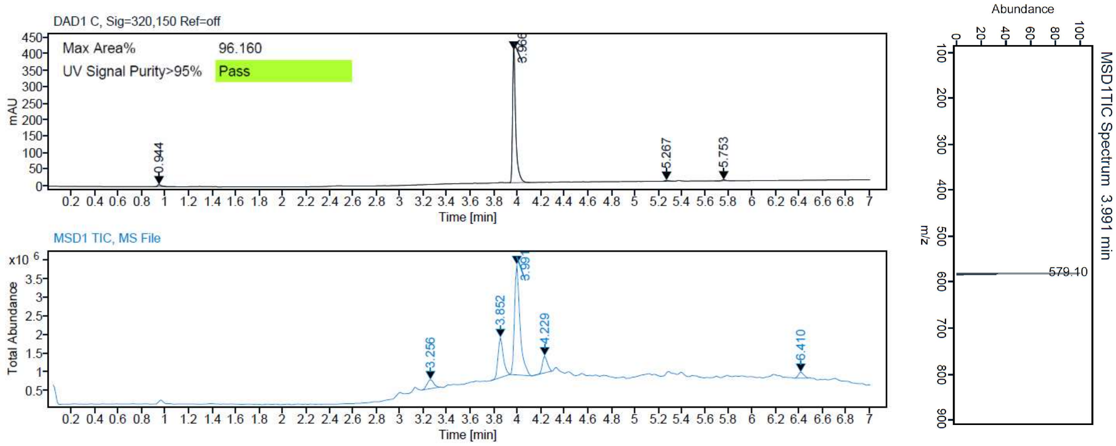


HPLC purity and mass spectrum (ESI) **20**

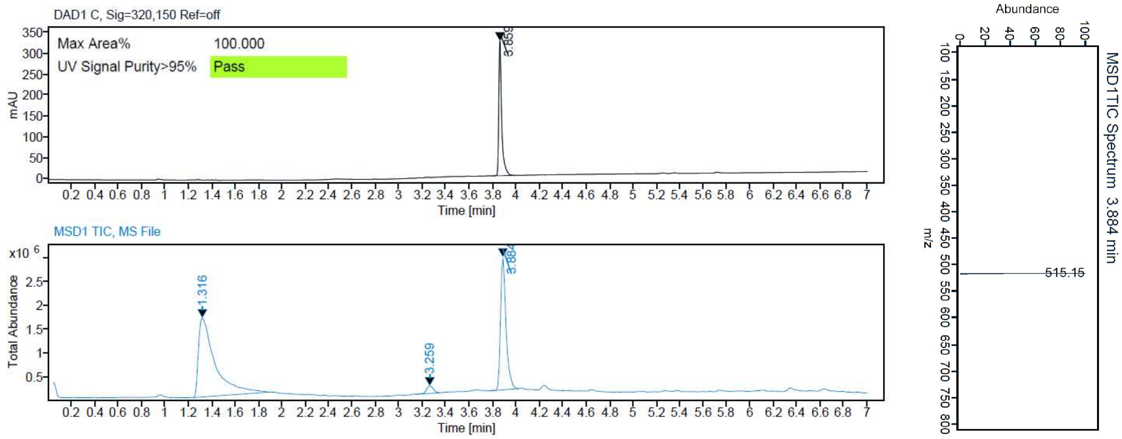


HPLC purity and mass spectrum (ESI) **21**

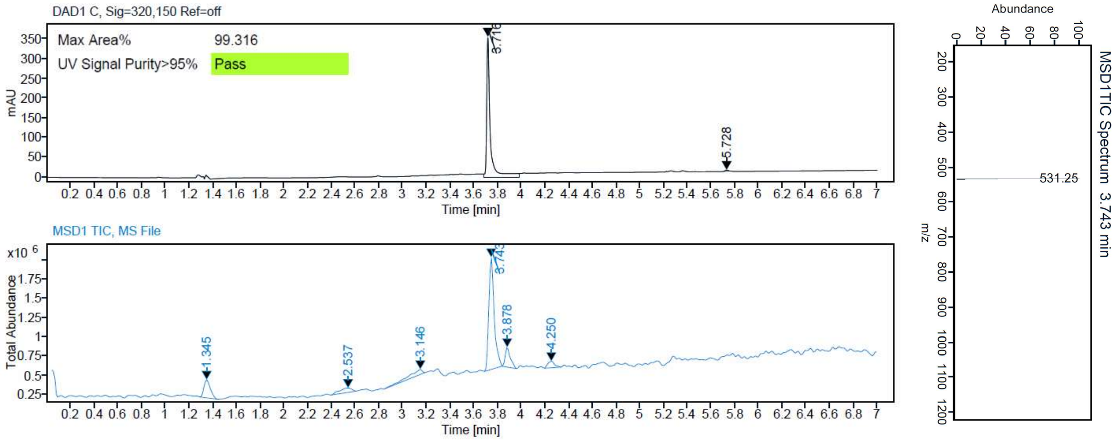

HPLC purity and mass spectrum (ESI) **22**

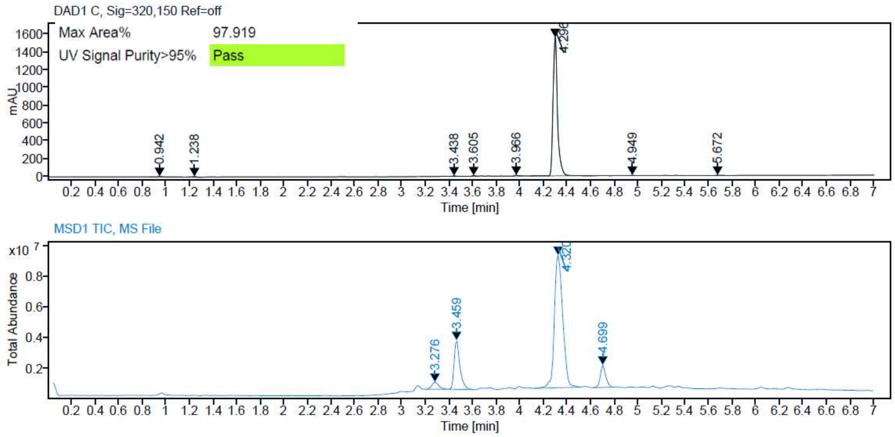


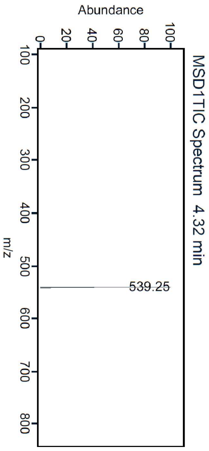


HPLC purity and mass spectrum (ESI) **23**

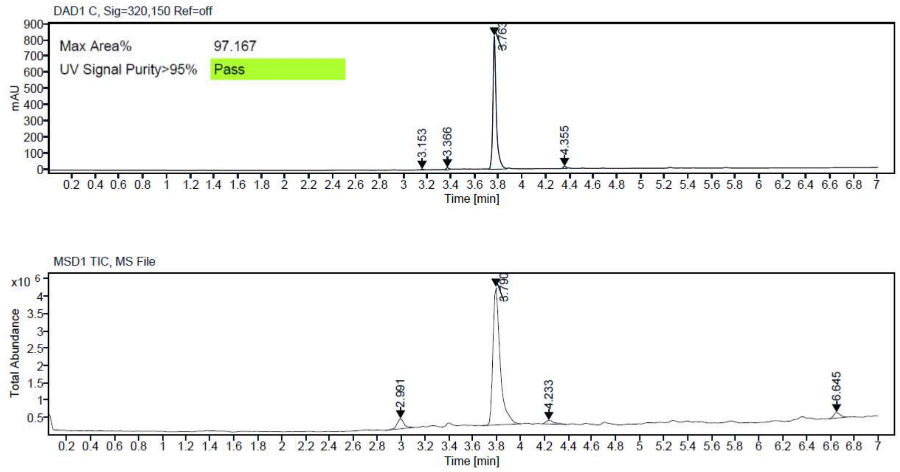


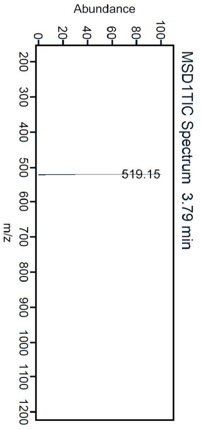


HPLC purity and mass spectrum (ESI) **24**

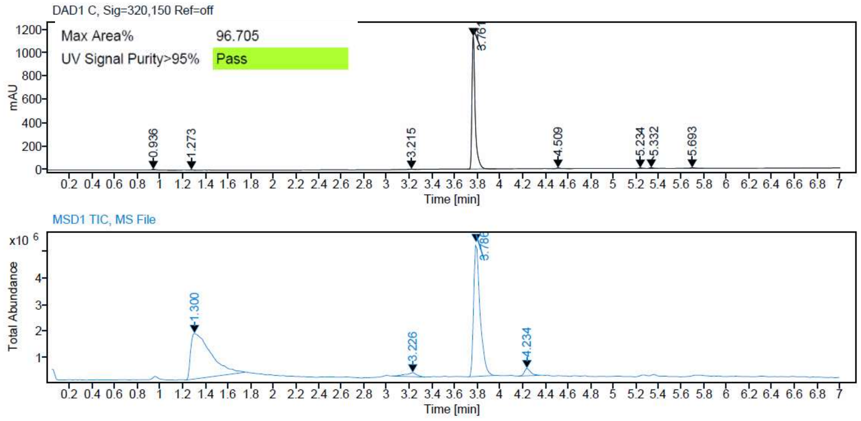


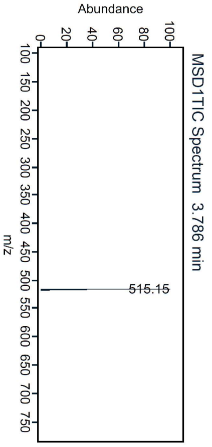

HPLC purity and mass spectrum (ESI) **25**

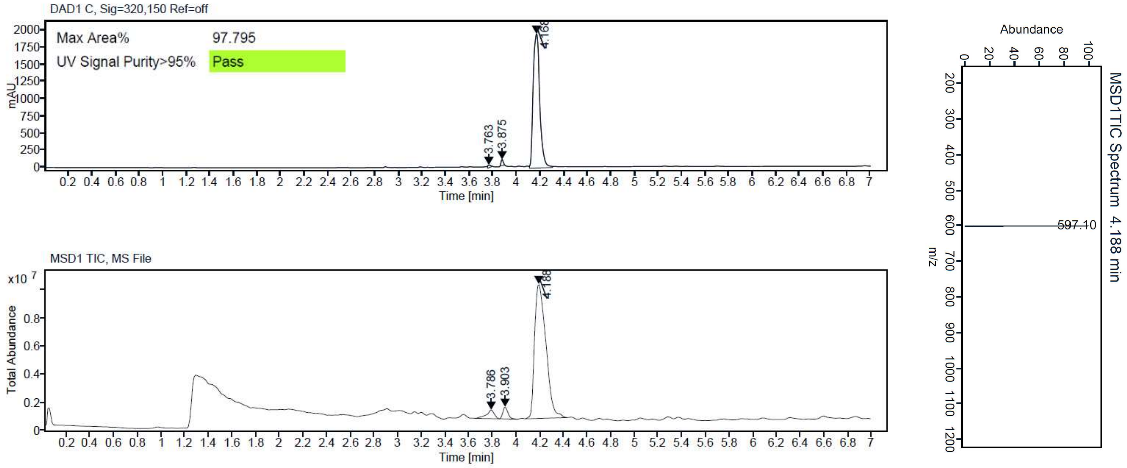


HPLC purity and mass spectrum (ESI) **26**

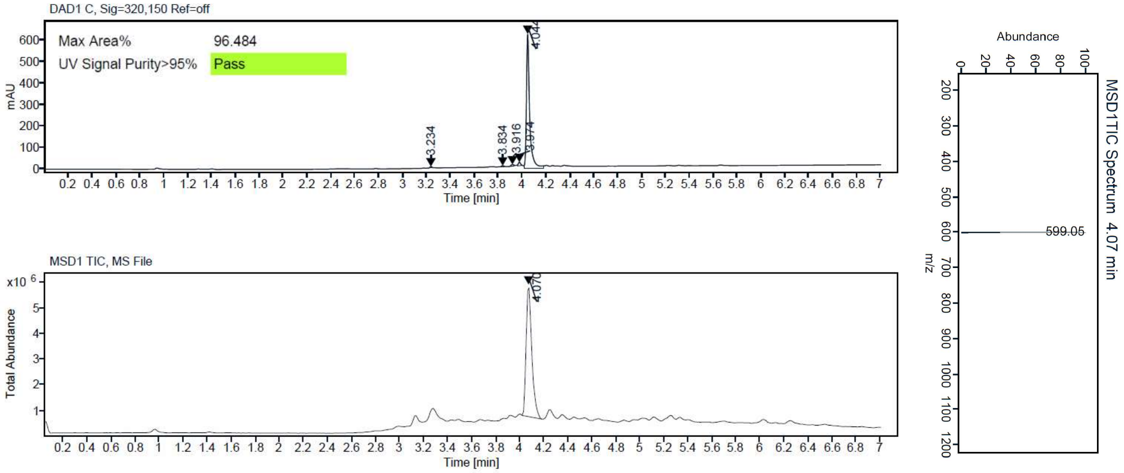


HPLC purity and mass spectrum (ESI) **27**

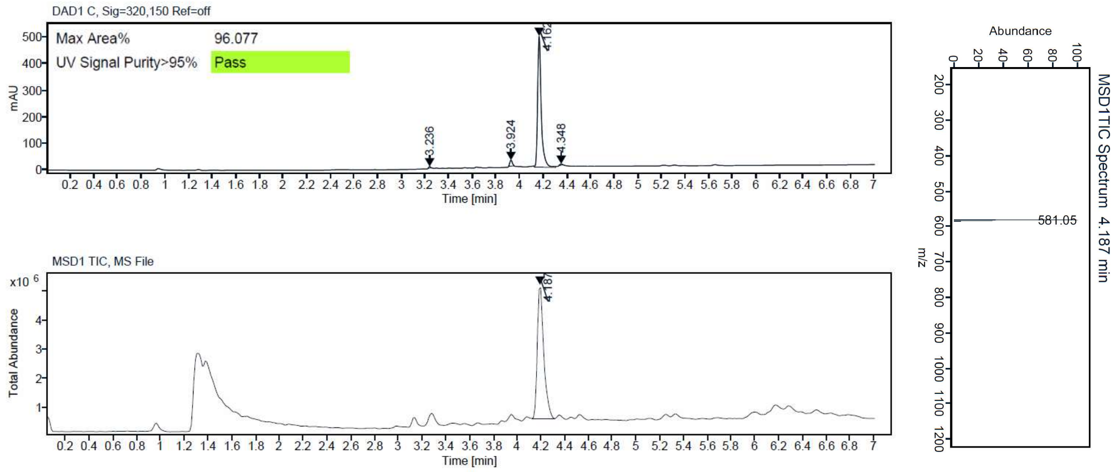

HPLC purity and mass spectrum (ESI) **28**

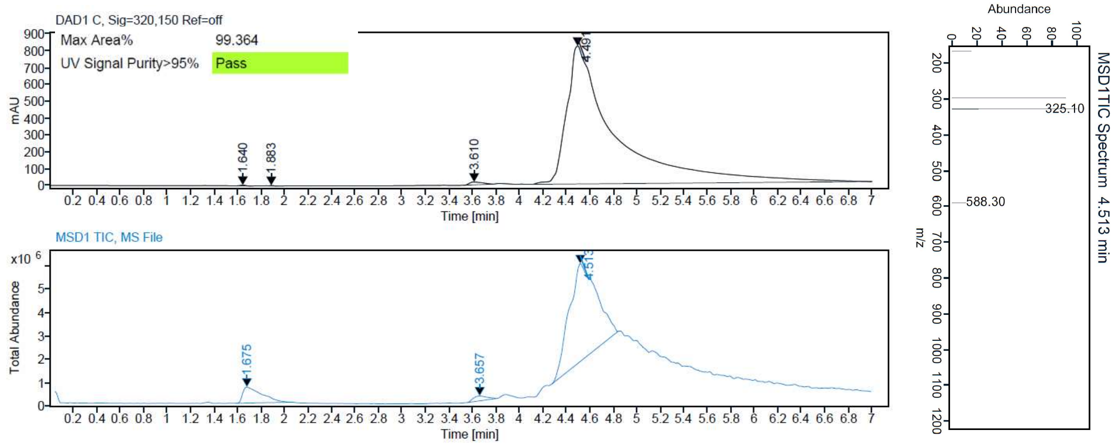


HPLC purity and mass spectrum (ESI) **29**

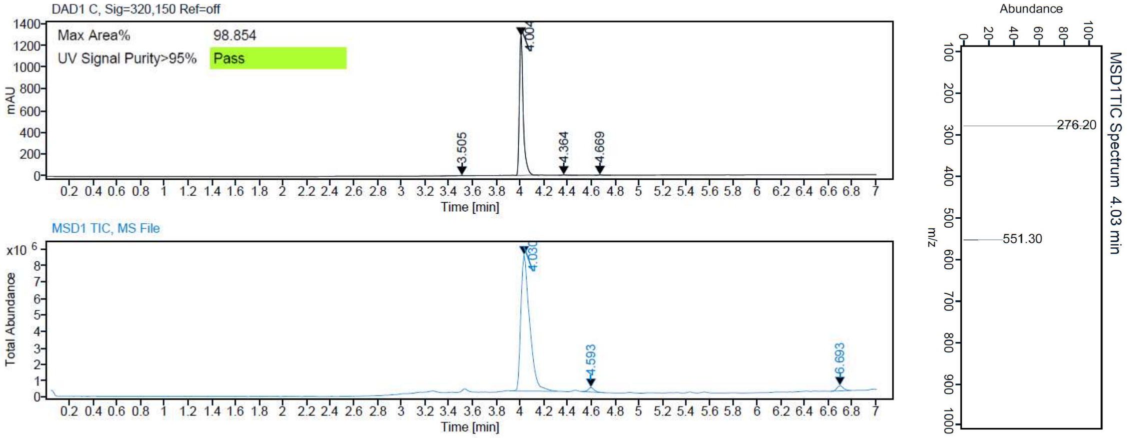


HPLC purity and mass spectrum (ESI) **30**

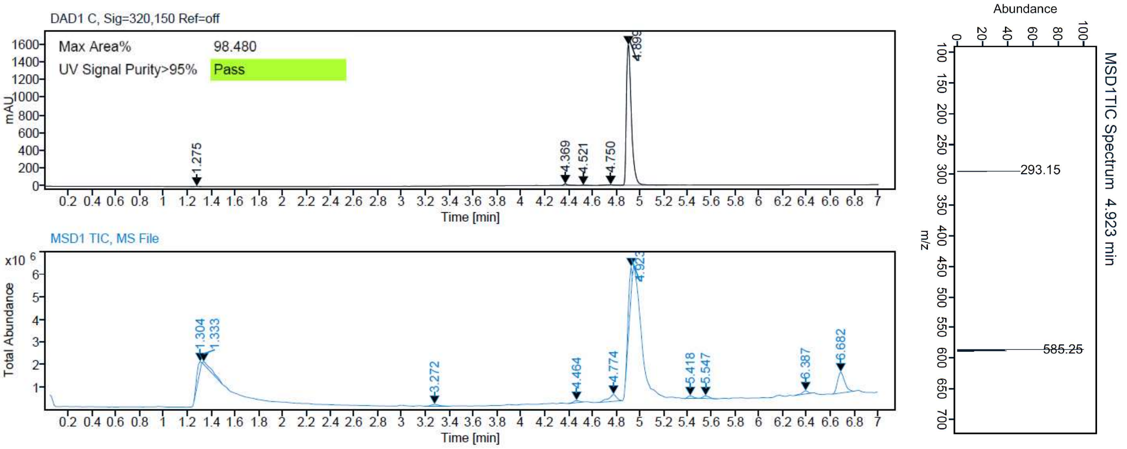

HPLC purity and mass spectrum (ESI) **31**

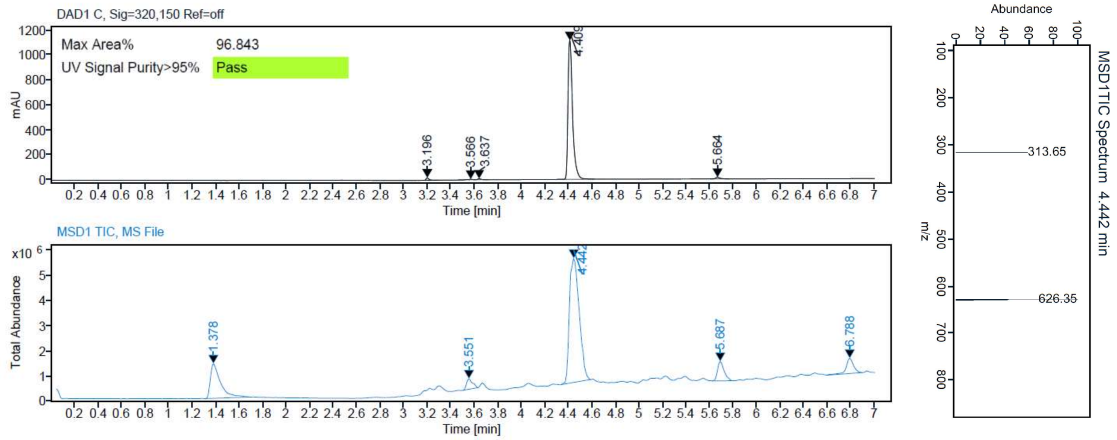


HPLC purity and mass spectrum (ESI) **32**

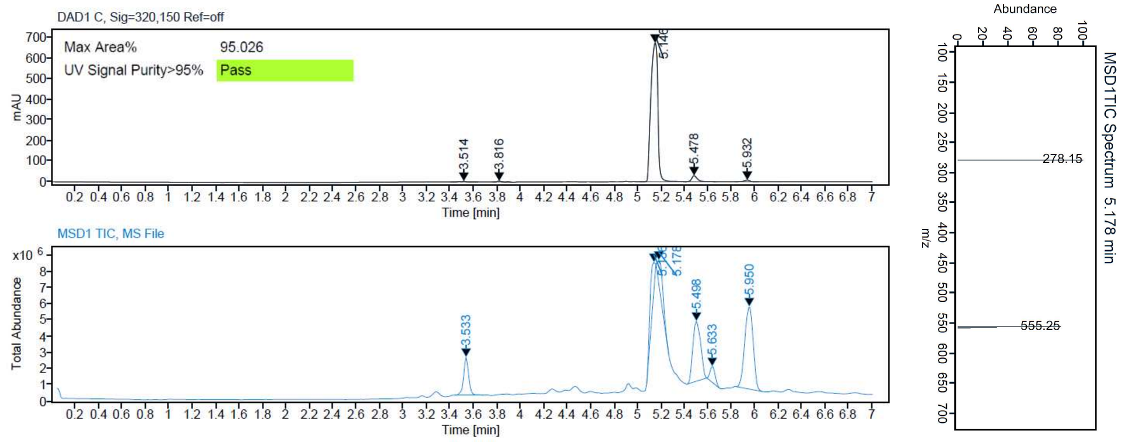


HPLC purity and mass spectrum (ESI) **33**

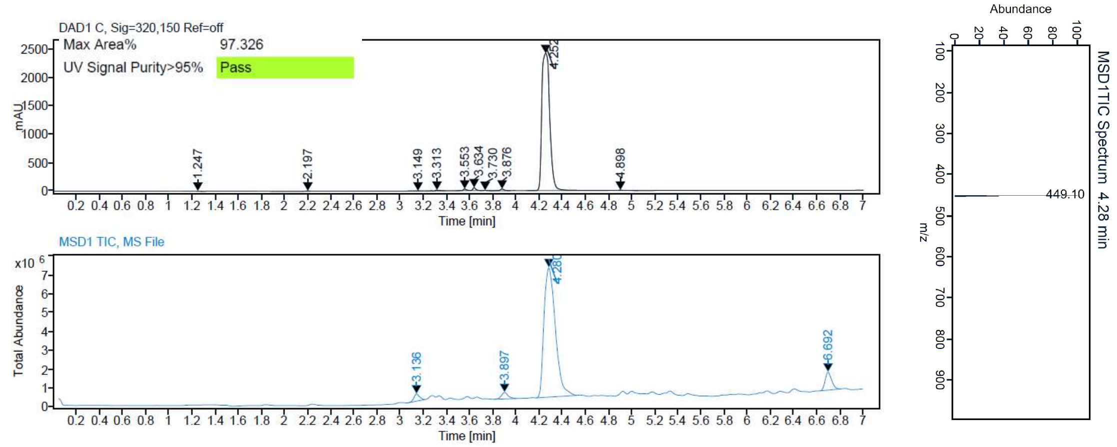

HPLC purity and mass spectrum (ESI) **34**

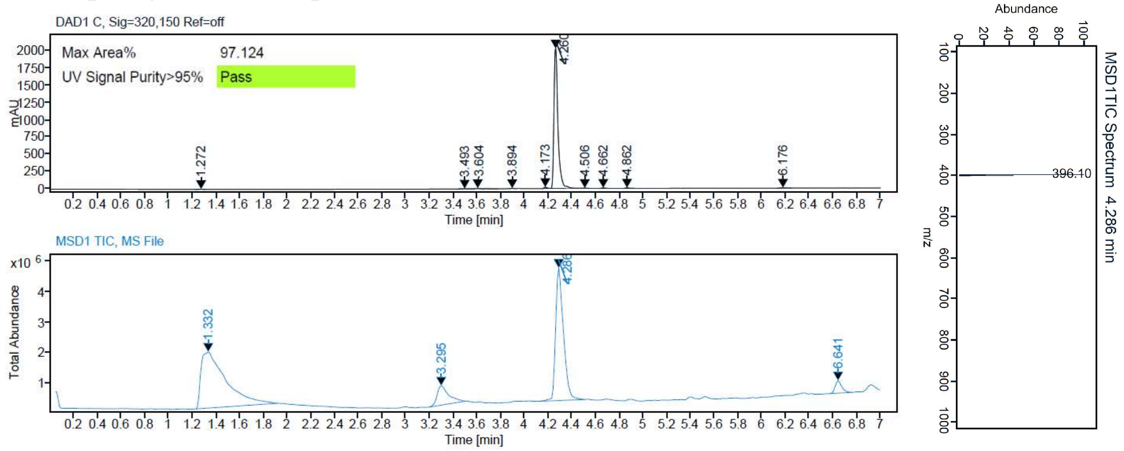


HPLC purity and mass spectrum (ESI) **35**

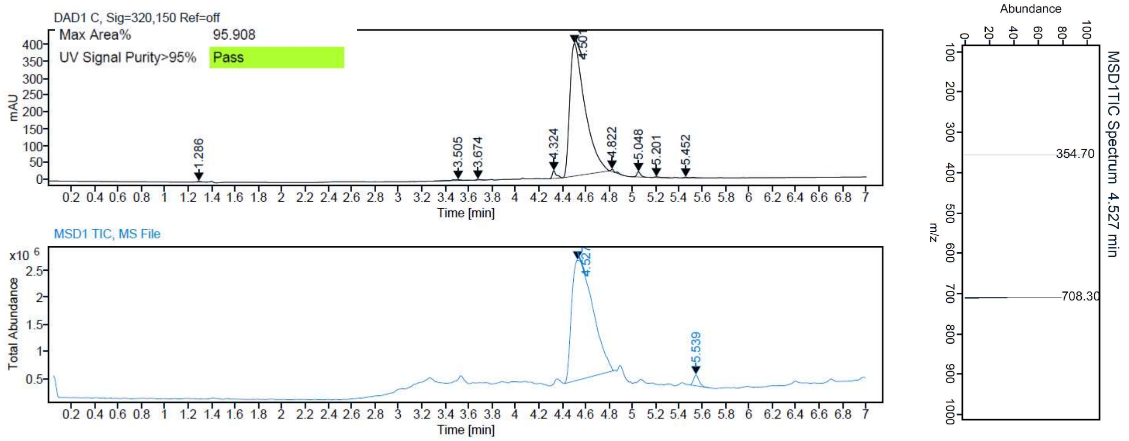


HPLC purity and mass spectrum (ESI) **36**

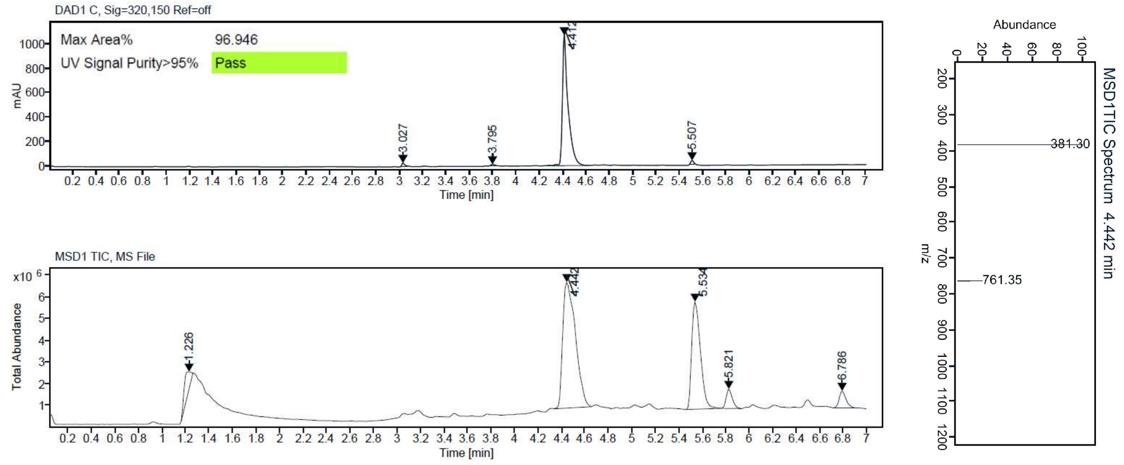

HPLC purity and mass spectrum (ESI) **37**

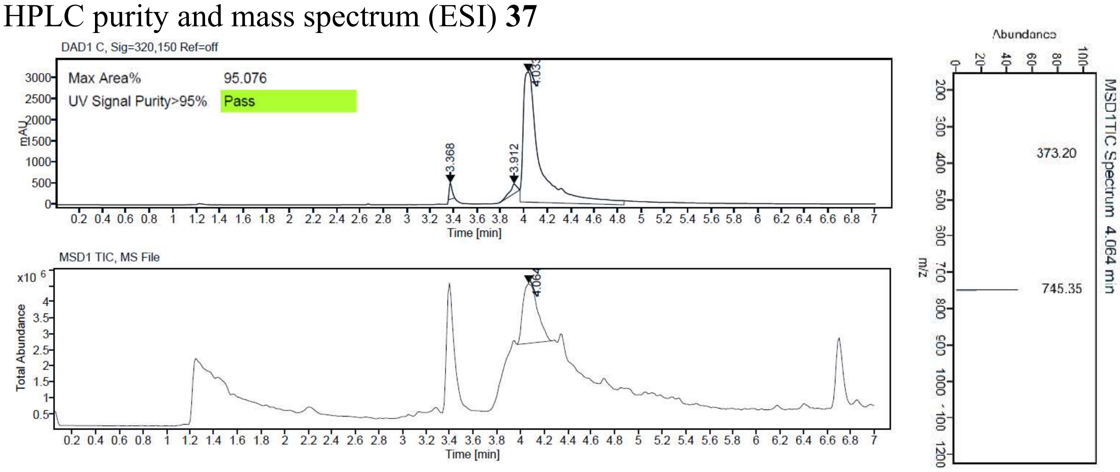


HPLC purity and mass spectrum (ESI) **47**

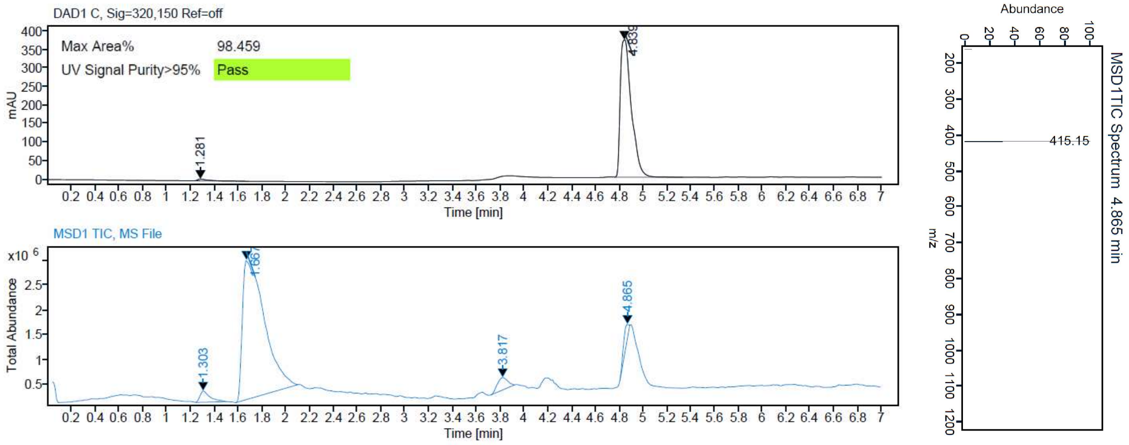


HPLC purity and mass spectrum (ESI) **48**

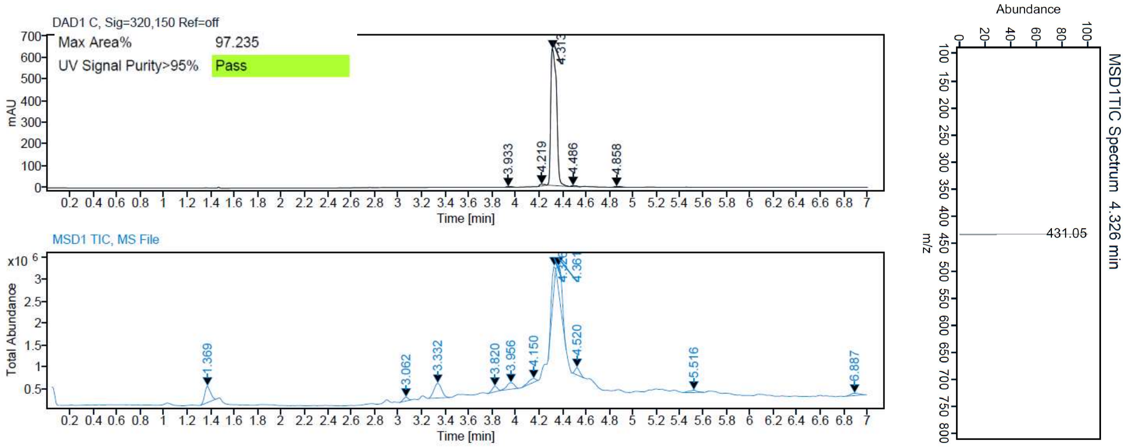

HPLC purity and mass spectrum (ESI) **49**

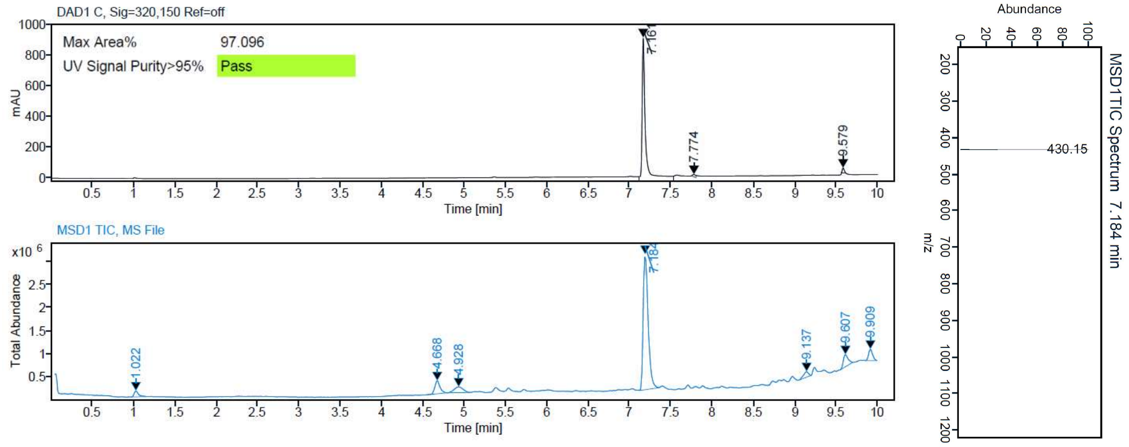


HPLC purity and mass spectrum (ESI) **50**

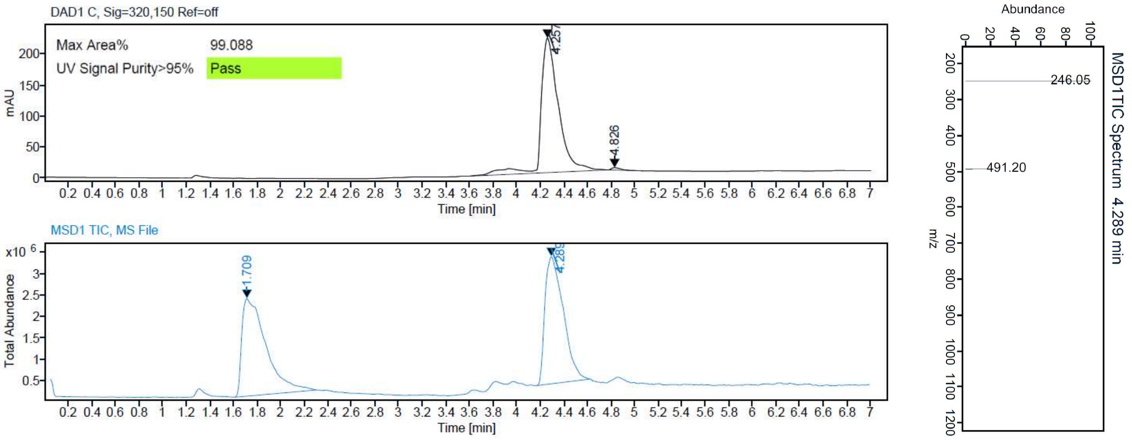


HPLC purity and mass spectrum (ESI) **51**

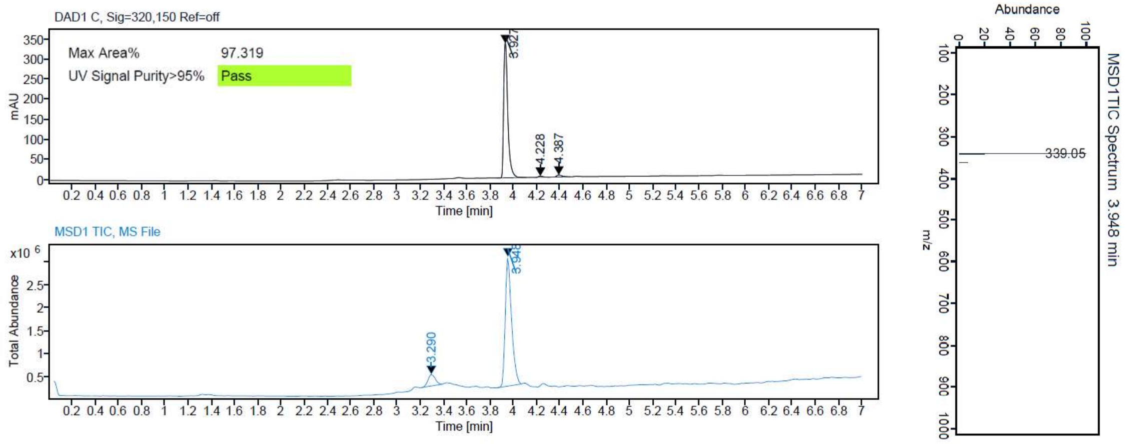

HPLC purity and mass spectrum (ESI) **52**

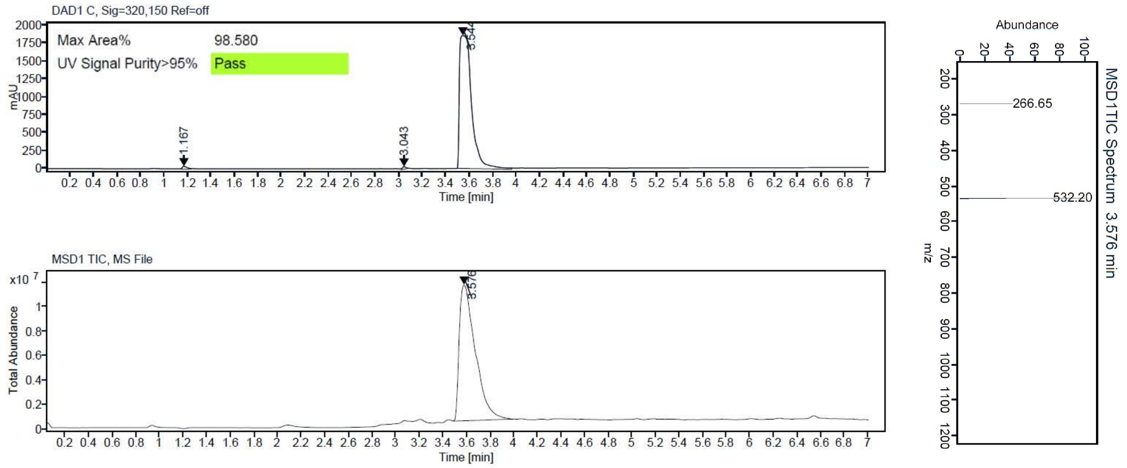


HPLC purity and mass spectrum (ESI) **53**

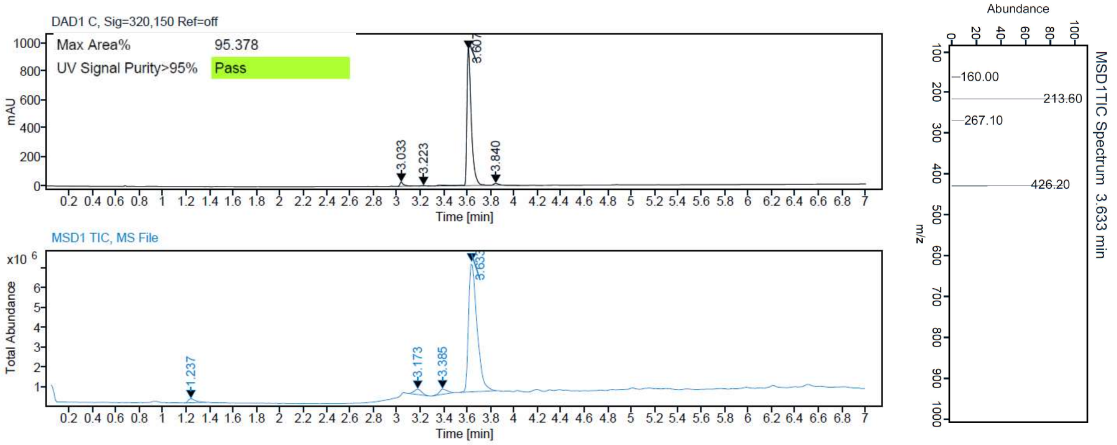


HPLC purity and mass spectrum (ESI) **54**

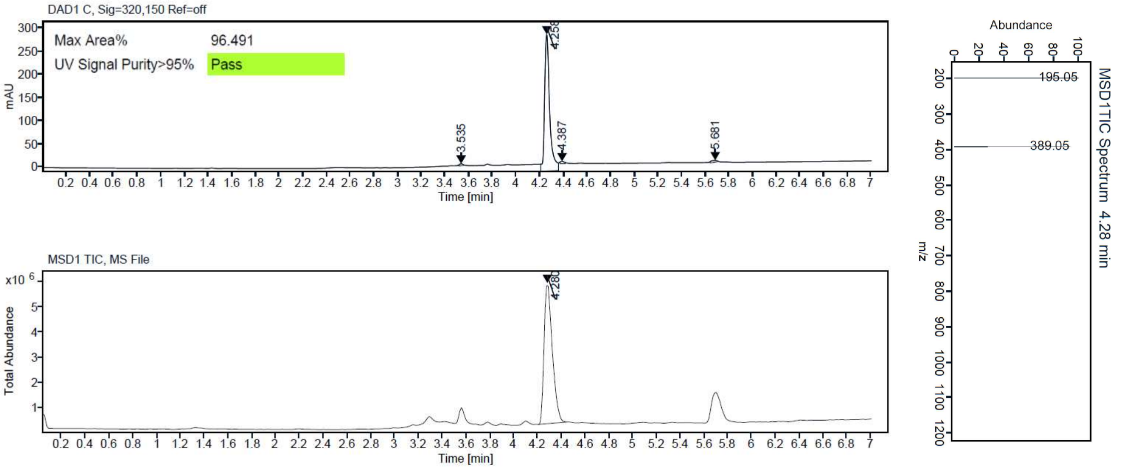

### HRMS

#### HRMS (MALDI) spectrum **1**

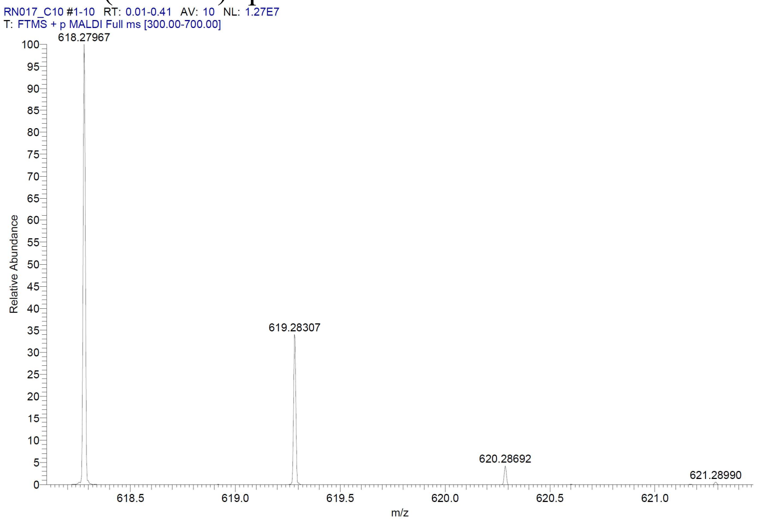


#### HRMS (MALDI) spectrum **2**

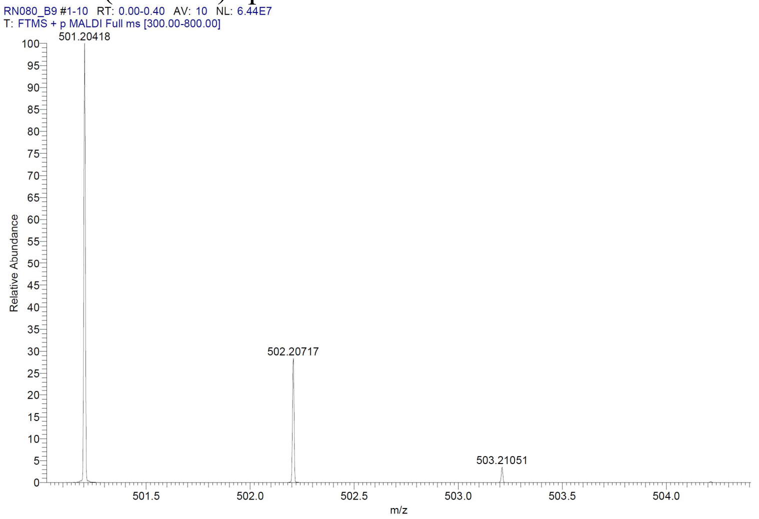


#### HRMS (MALDI) spectrum **3**

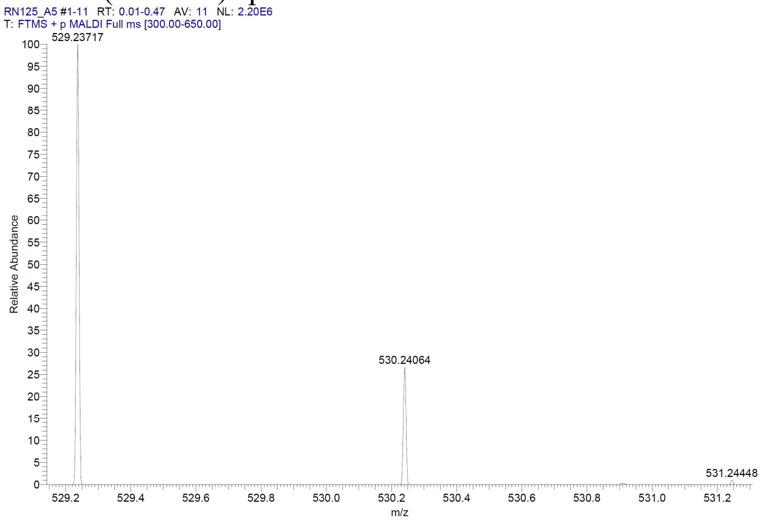

HRMS (MALDI) spectrum **4**

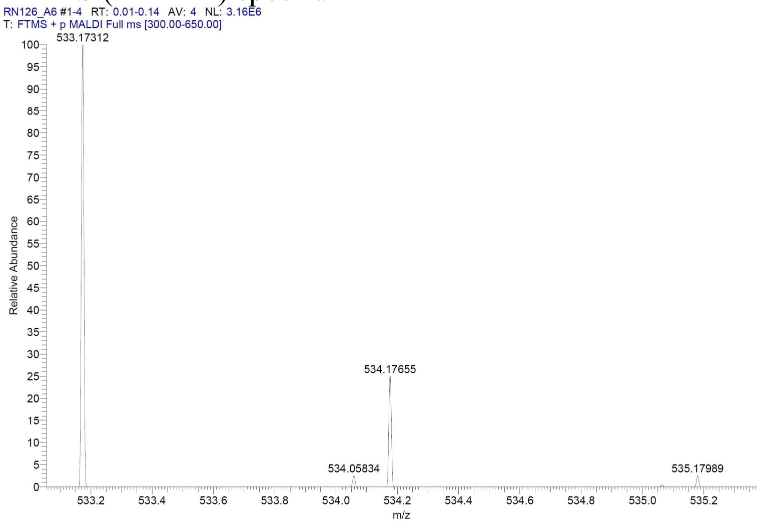


HRMS (MALDI) spectrum **5**

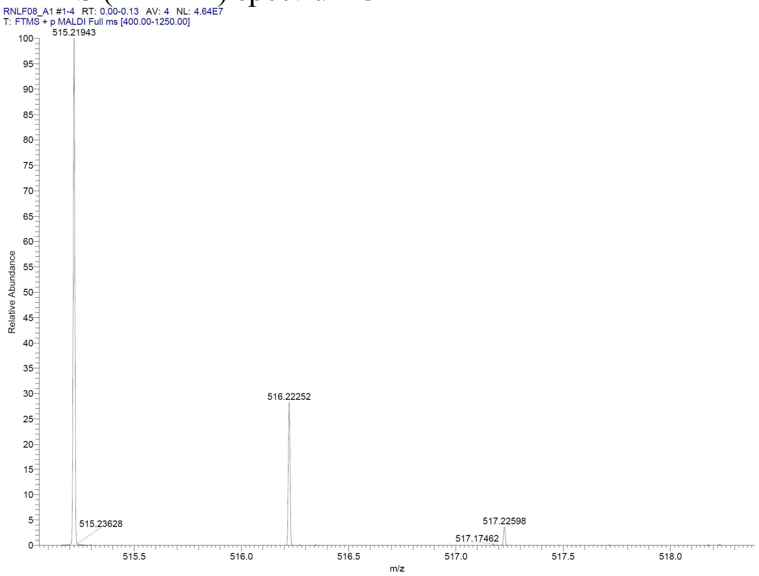


HRMS (MALDI) spectrum **6**

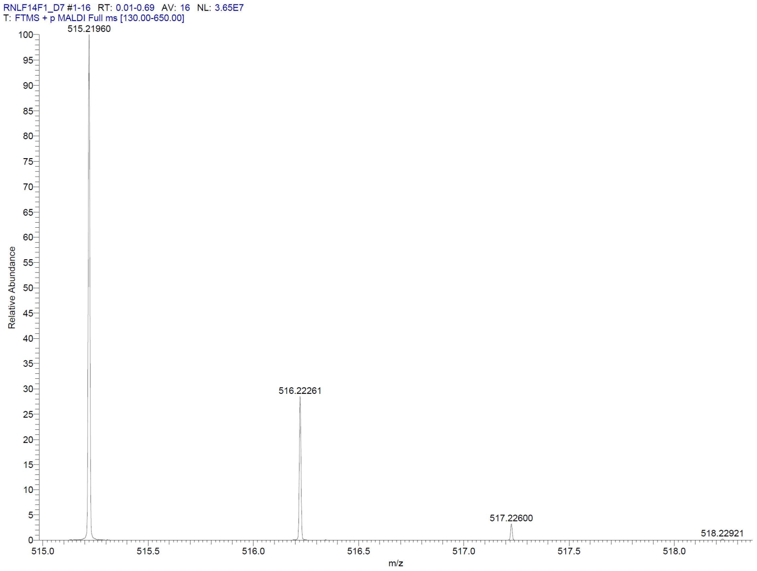


### HRMS (MALDI) spectrum **7**

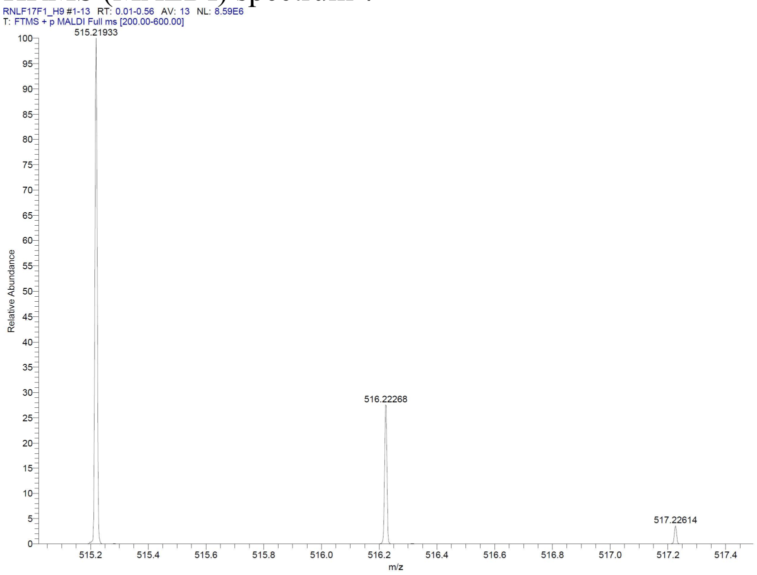


### HRMS (MALDI) spectrum **8**

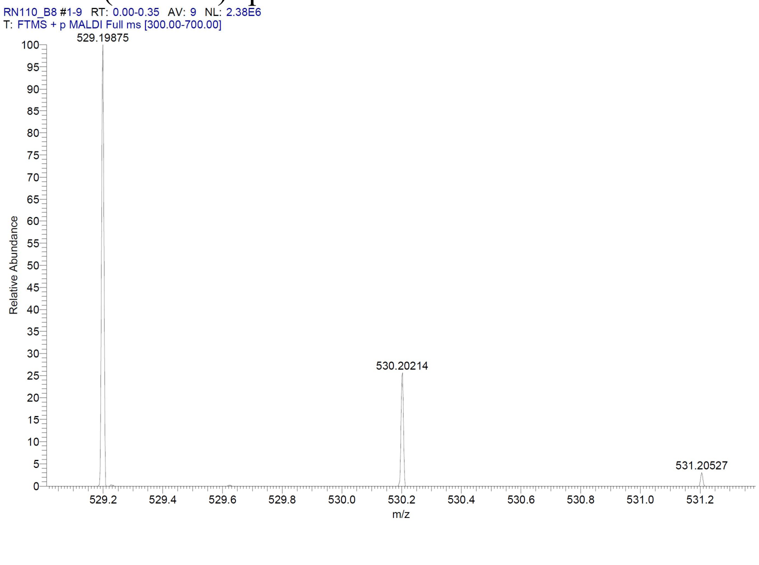

HRMS (MALDI) spectrum **9**

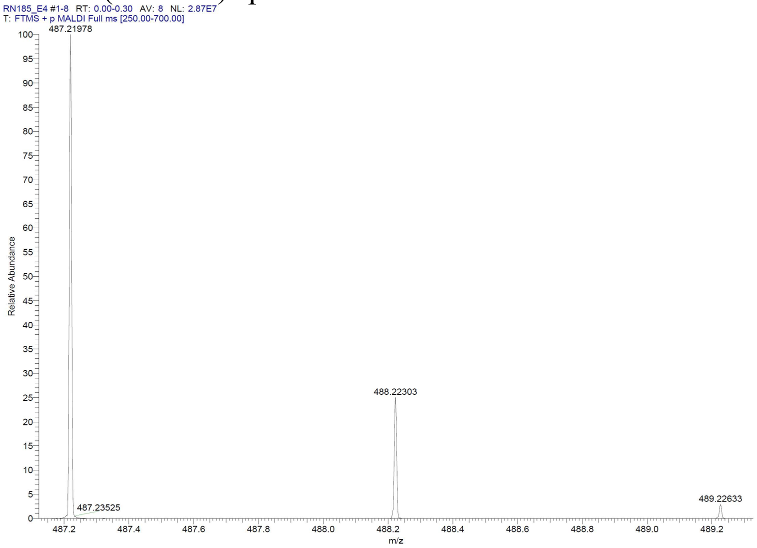


HRMS (MALDI) spectrum **10**

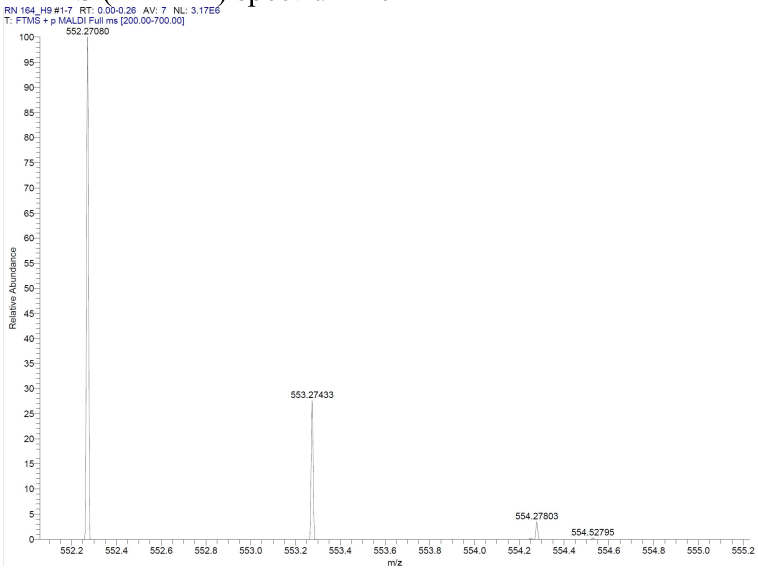

### HRMS (MALDI) spectrum **11**

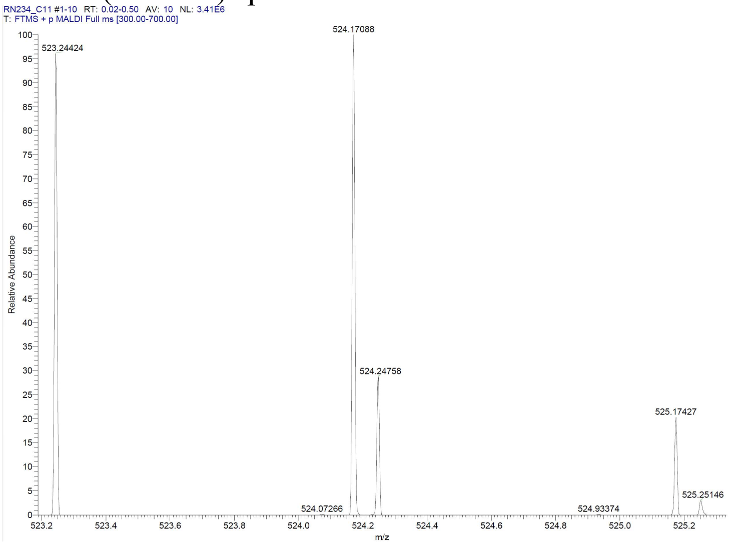


### HRMS (MALDI) spectrum **12**

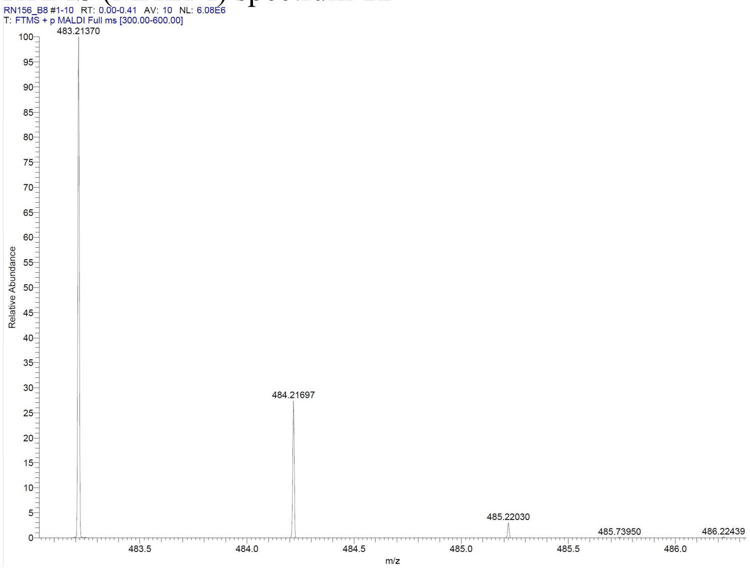

HRMS (MALDI) spectrum **13**

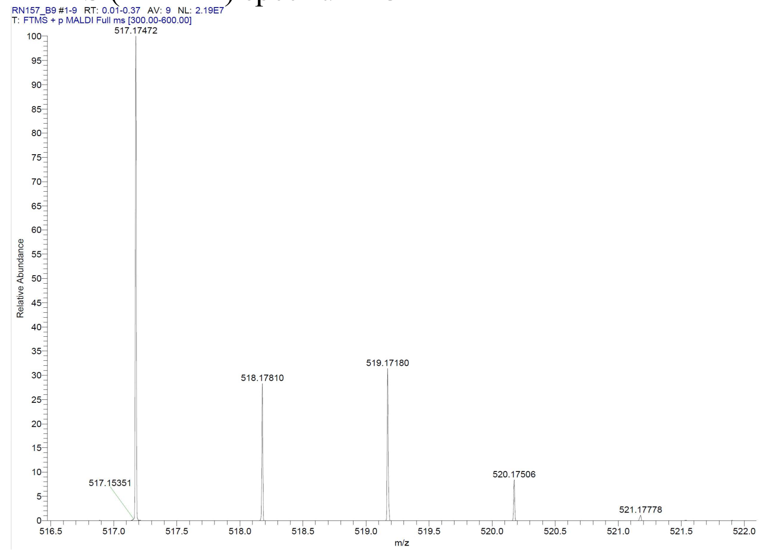


HRMS (MALDI) spectrum **14**

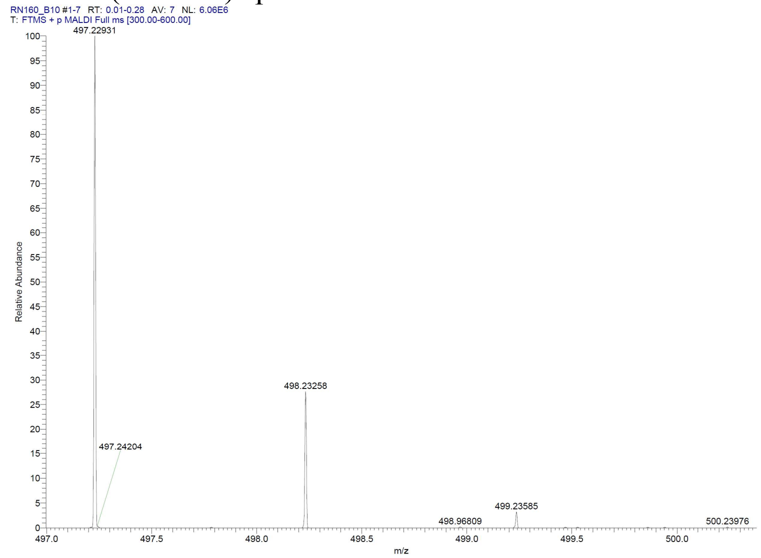

#### HRMS (MALDI) spectrum **15**

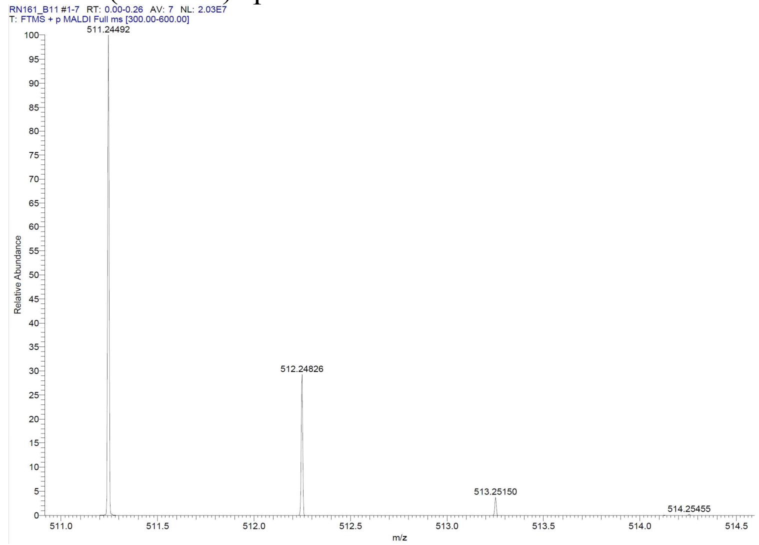


#### HRMS (MALDI) spectrum **16**

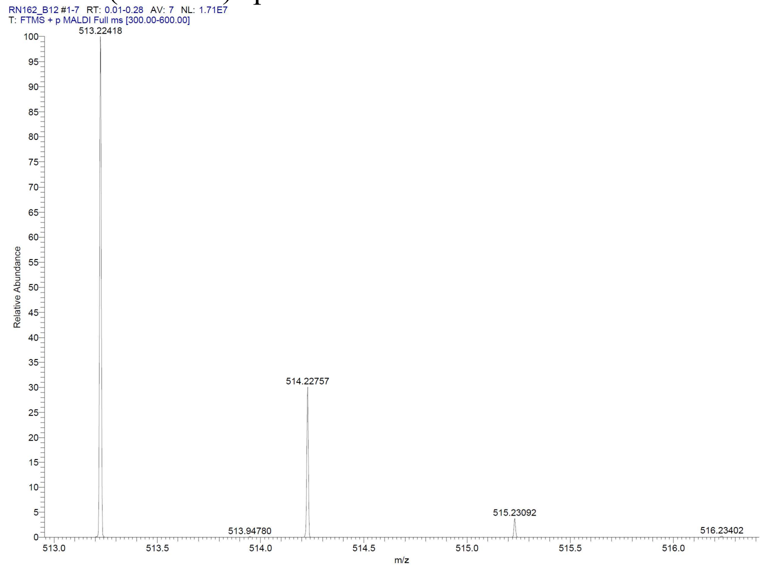

HRMS (MALDI) spectrum **17**

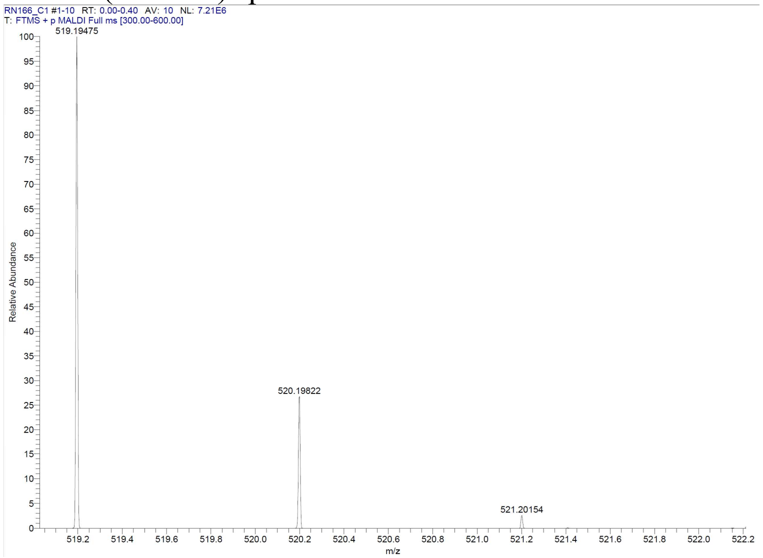


HRMS (MALDI) spectrum **18**

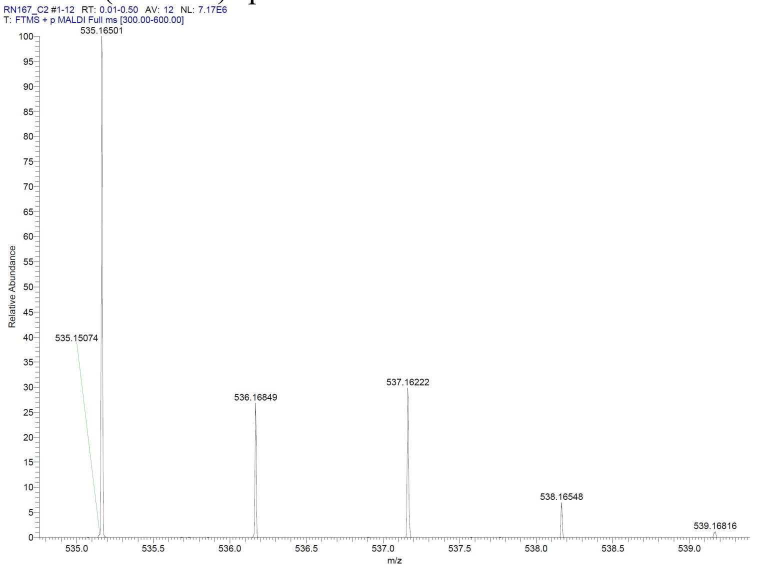

### HRMS (MALDI) spectrum **19**

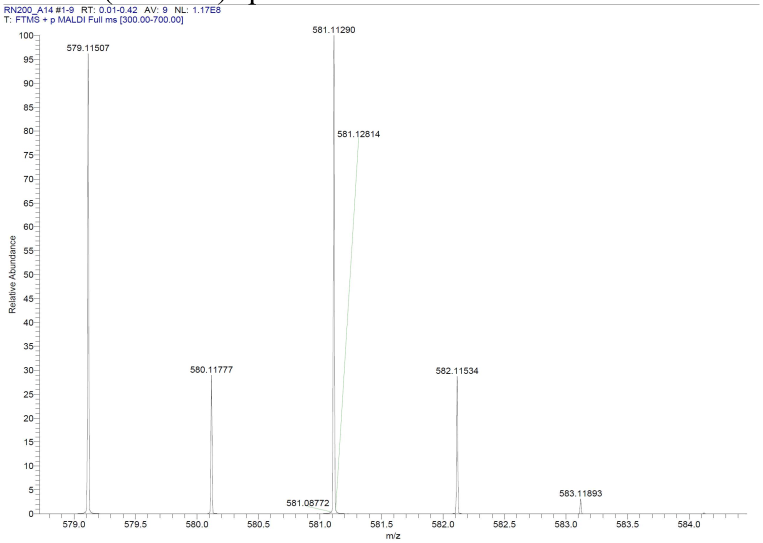


### HRMS (MALDI) spectrum **20**

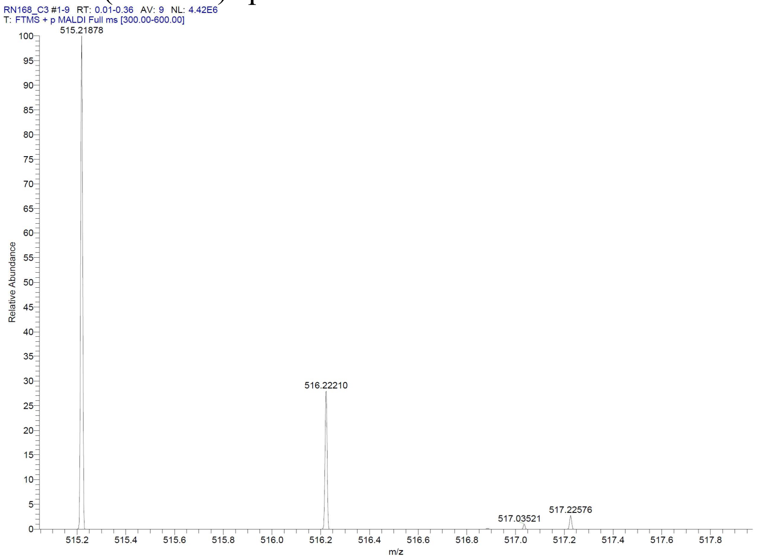

### HRMS (MALDI) spectrum **21**

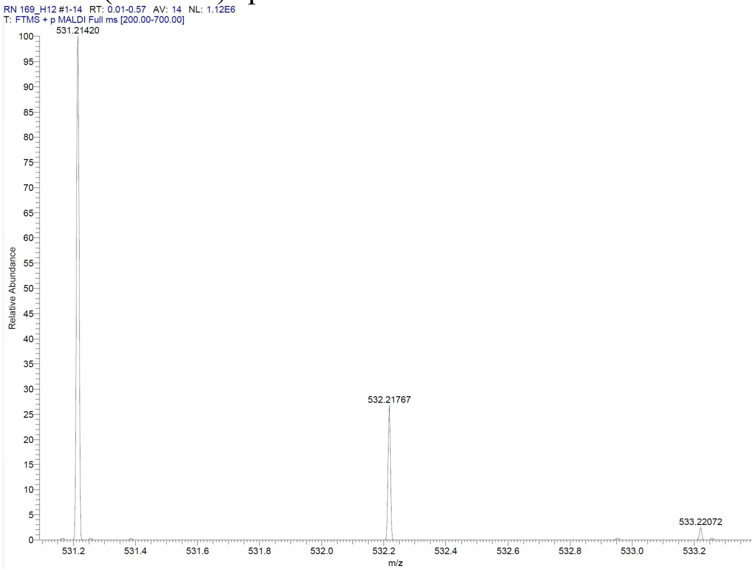


### HRMS (MALDI) spectrum **22**

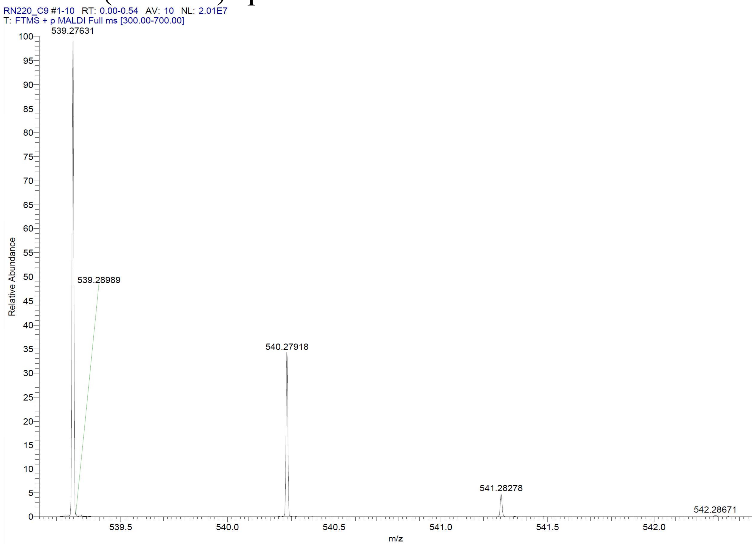

HRMS (MALDI) spectrum **23**

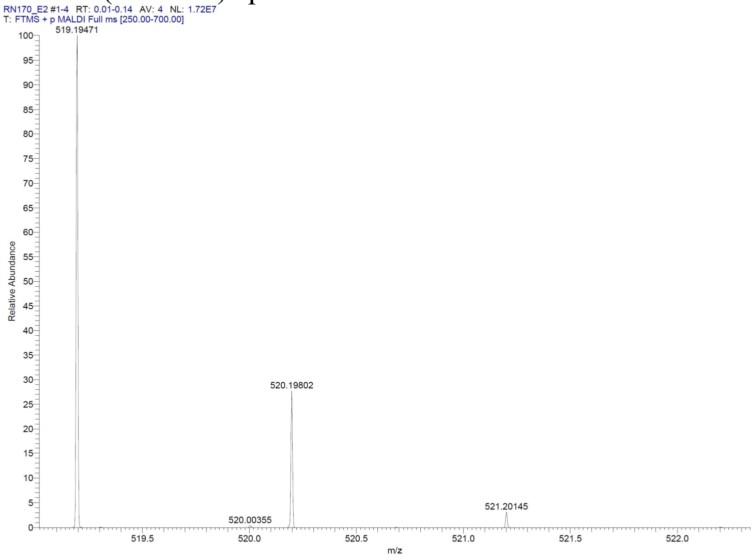


HRMS (MALDI) spectrum **24**

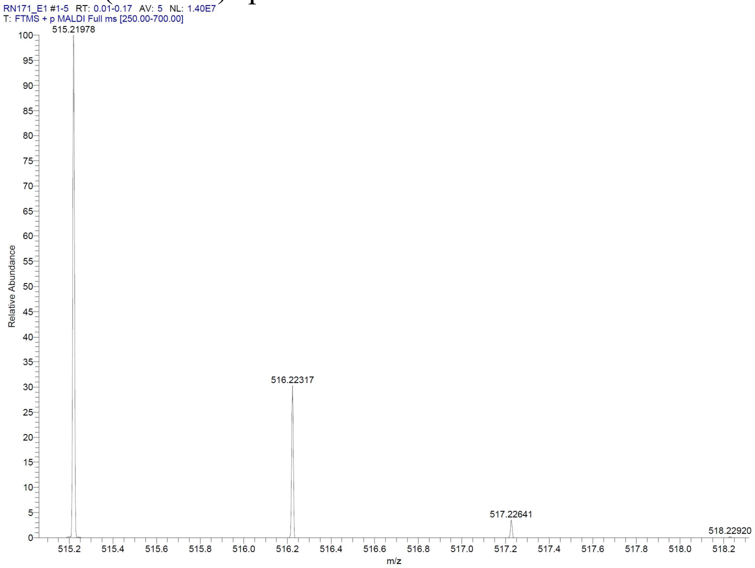

HRMS (MALDI) spectrum **25**

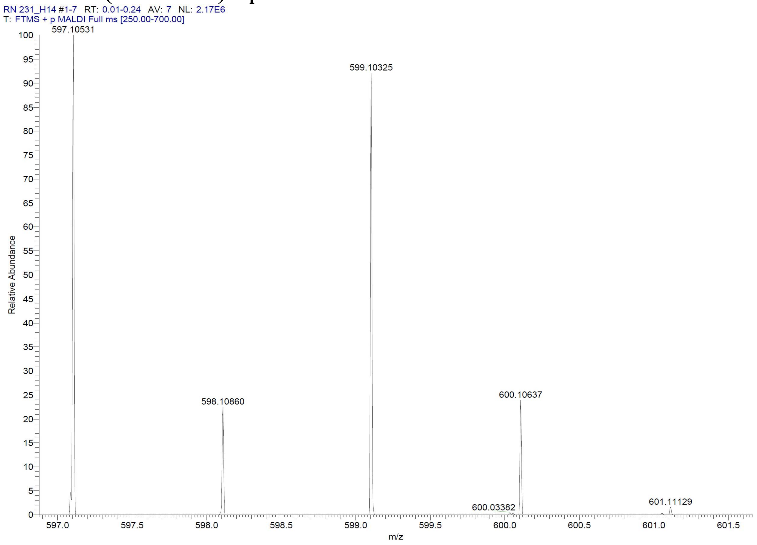


HRMS (MALDI) spectrum **26**

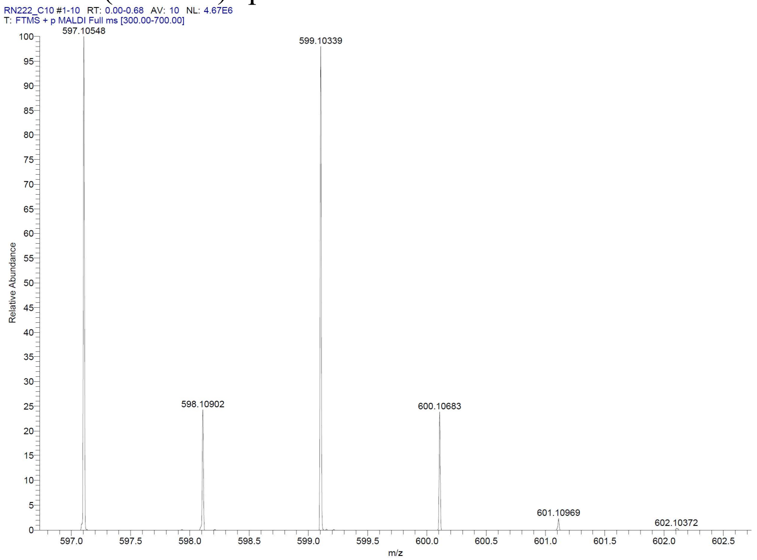

HRMS (MALDI) spectrum **27**

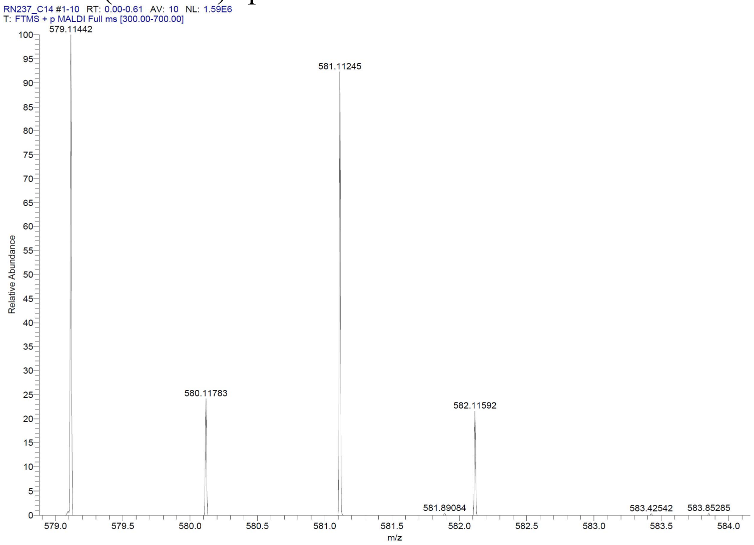


HRMS (MALDI) spectrum **28**

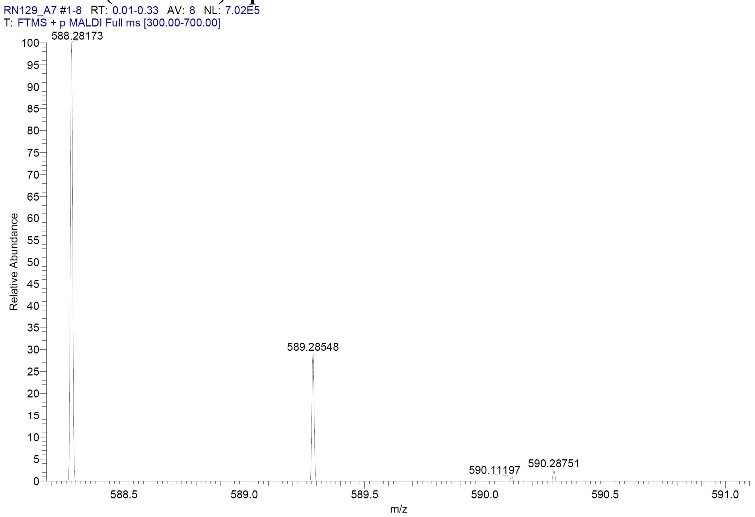


HRMS (MALDI) spectrum **29**

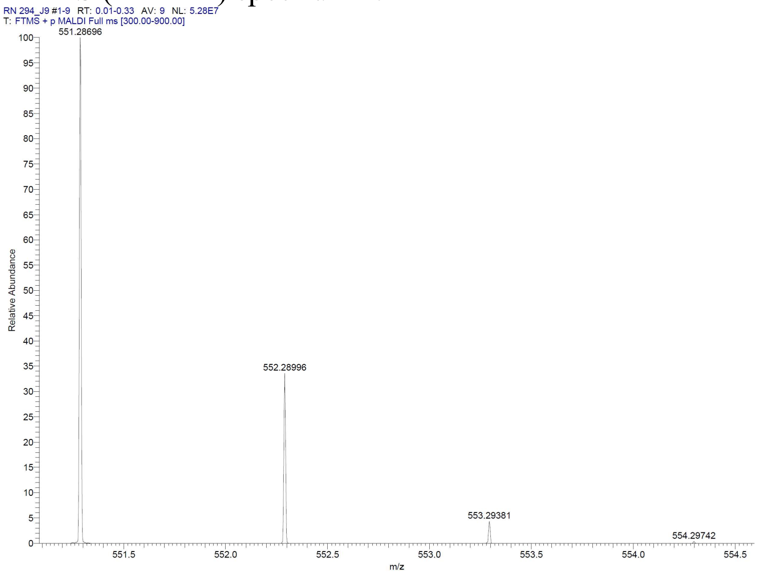

HRMS (MALDI) spectrum **30**

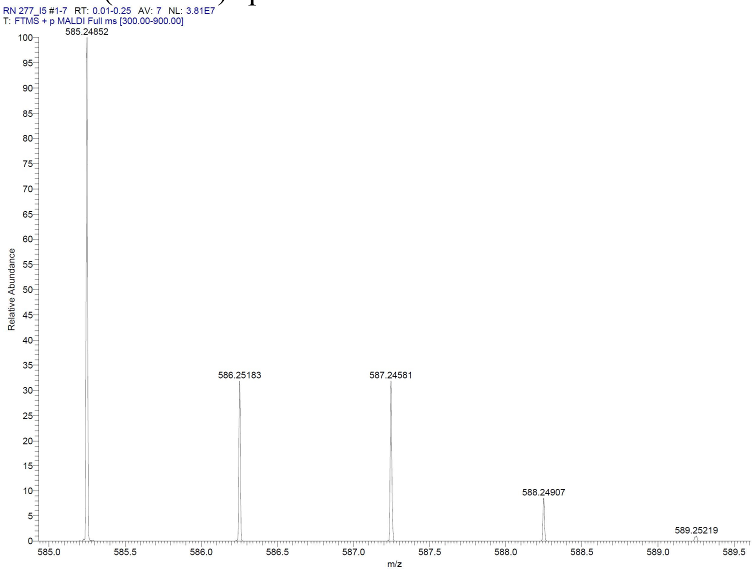


HRMS (ESI) spectrum **31**

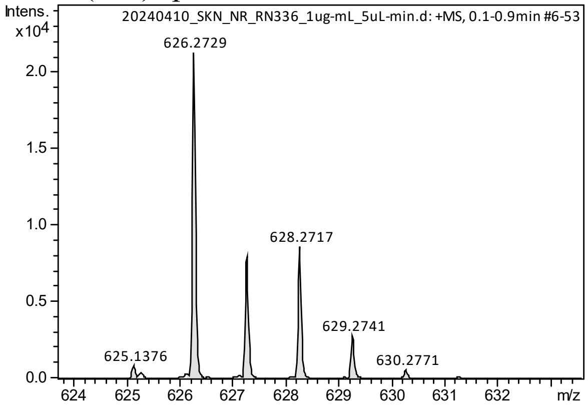


HRMS (ESI) spectrum **32**

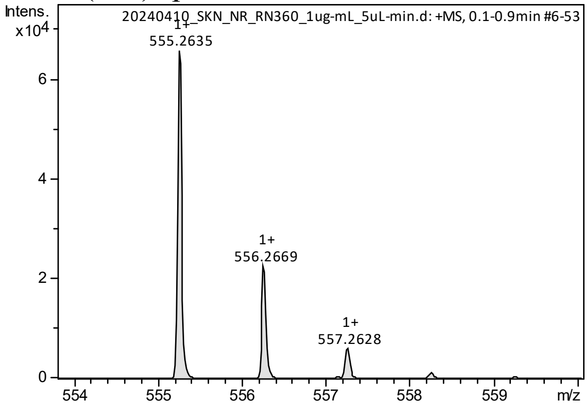

### HRMS (MALDI) spectrum **33**

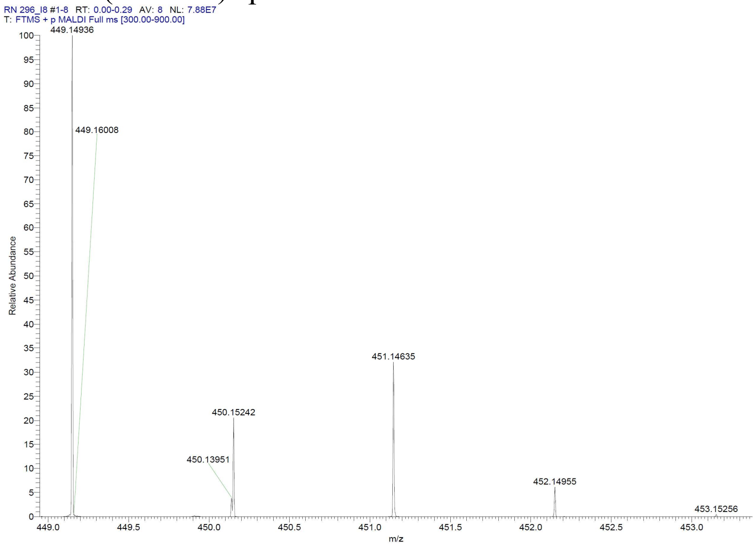


### HRMS (MALDI) spectrum **34**

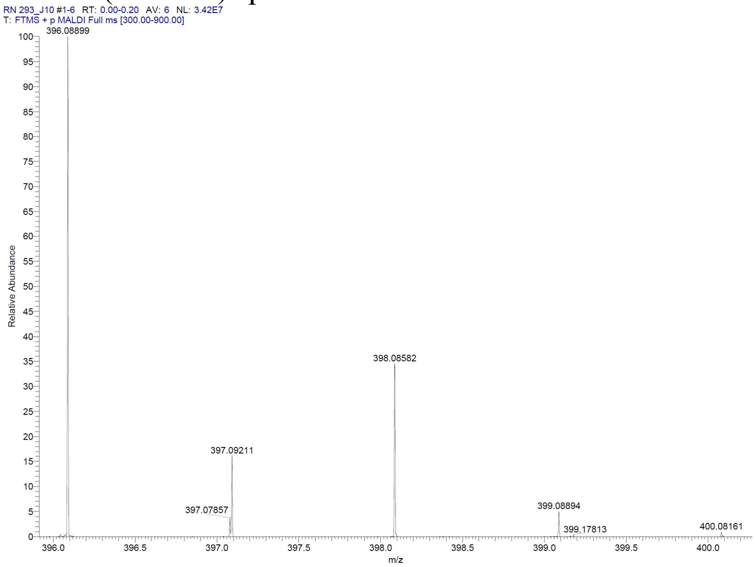

HRMS (MALDI) spectrum **35**

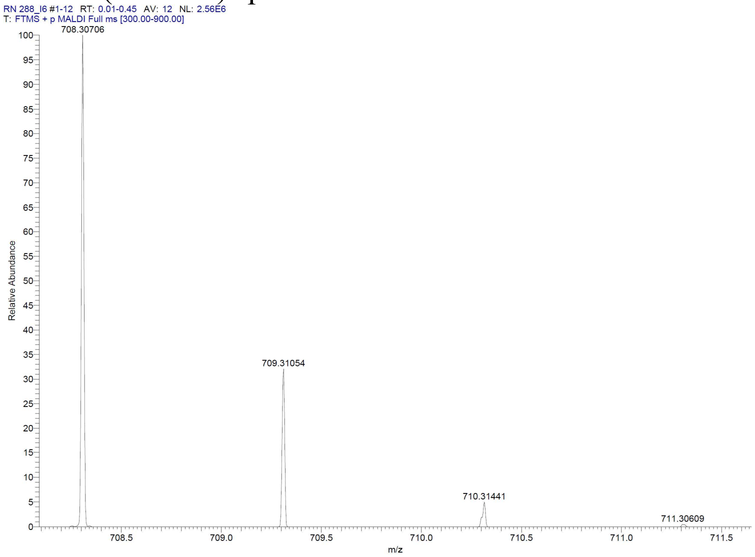


HRMS (ESI) spectrum **36**

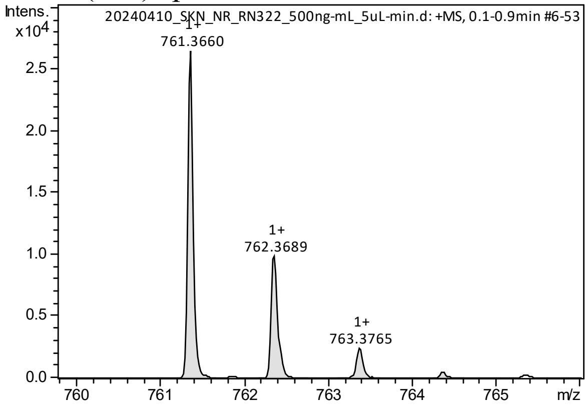


HRMS (MALDI) spectrum **37**

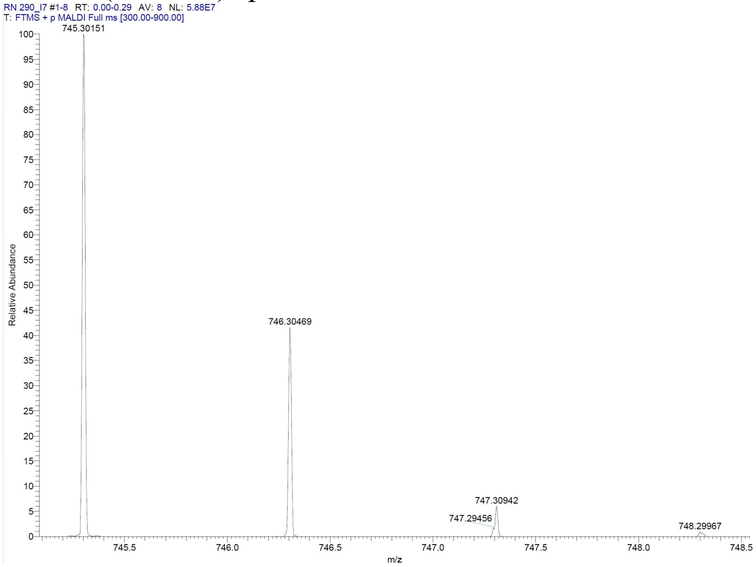

HRMS (MALDI) spectrum **47**

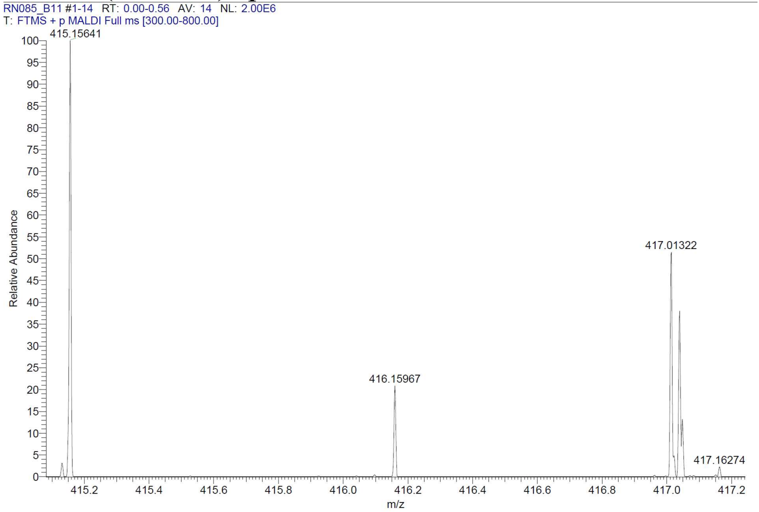


HRMS (MALDI) spectrum **48**

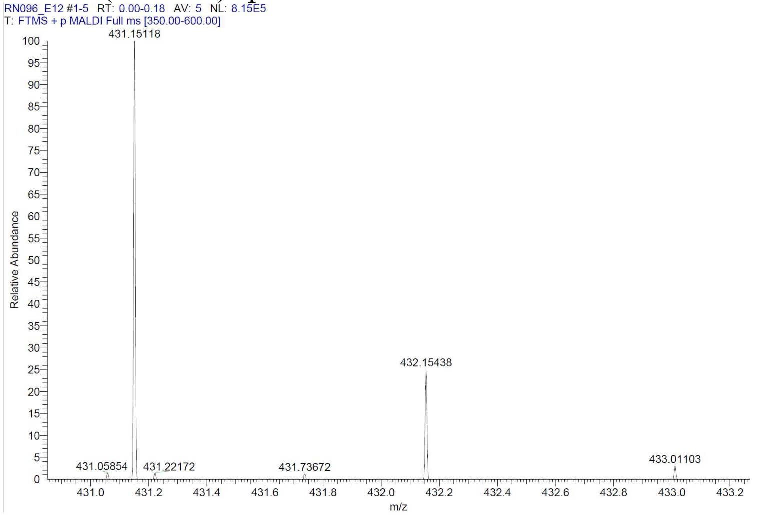


HRMS (MALDI) spectrum **49**

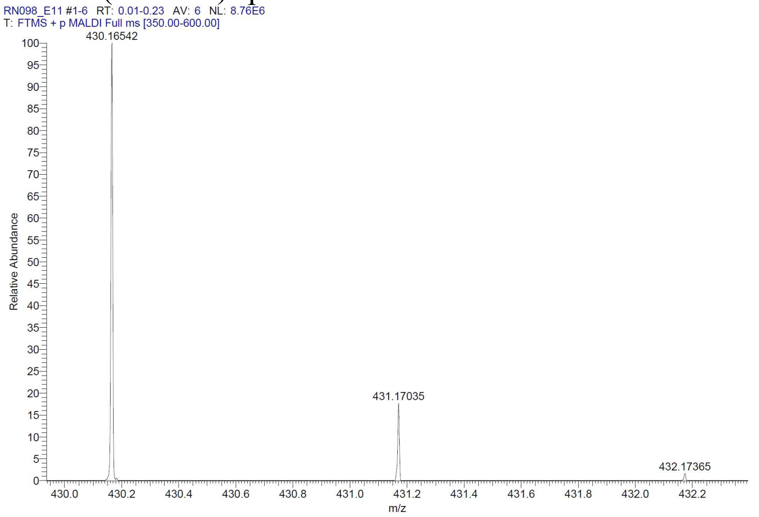

HRMS (MALDI) spectrum **50**

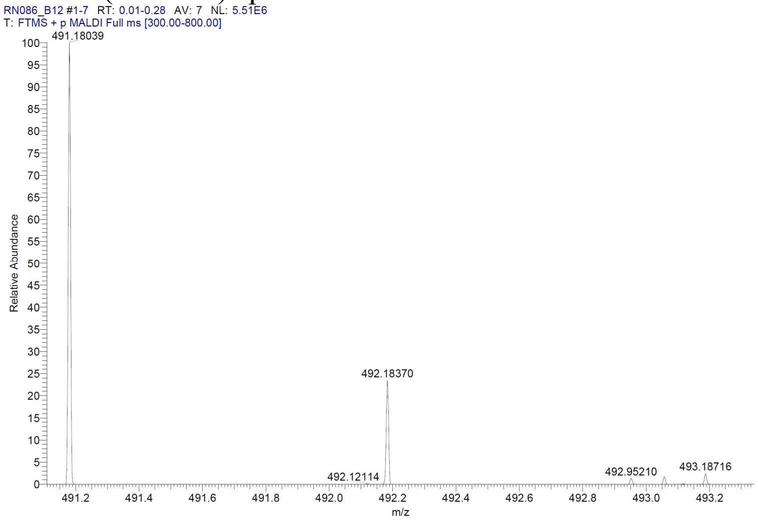


HRMS (ESI) spectrum **51**

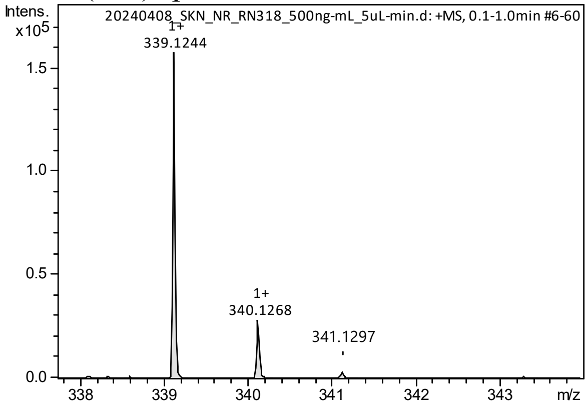


HRMS (ESI) spectrum **52**

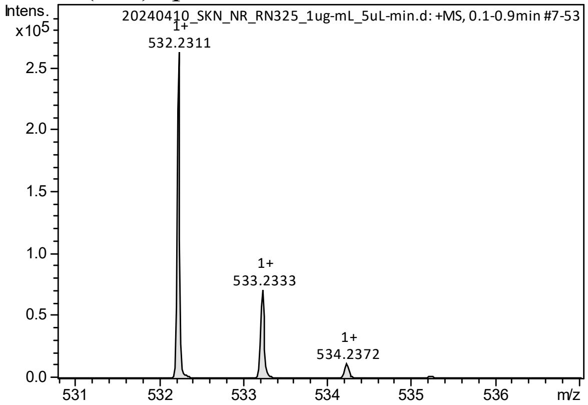

HRMS (ESI) spectrum **53**

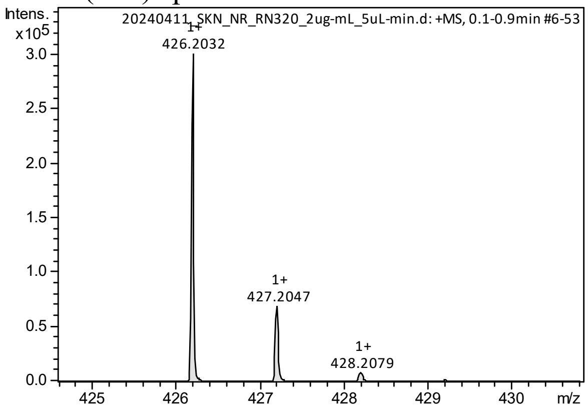


HRMS (ESI) spectrum **54**

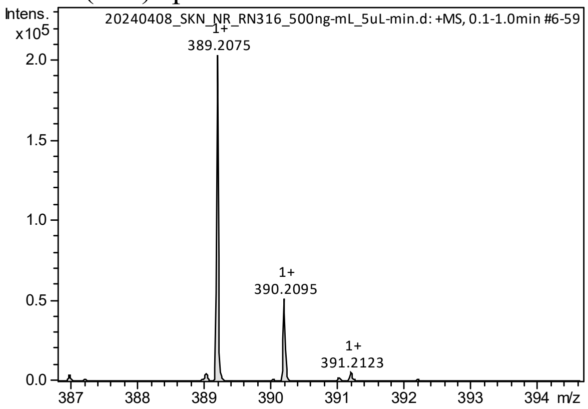
